# Dynamic Landscape Analysis of Cell Fate Decisions: Predictive Models of Neural Development From Single-Cell Data

**DOI:** 10.1101/2025.05.28.656648

**Authors:** Marine Fontaine, M. Joaquina Delas, Meritxell Saez, Rory J. Maizels, Elizabeth Finnie, James Briscoe, David A. Rand

**Affiliations:** Mathematics Institute, University of Warwick, Coventry CV4 7AL, UK; The Francis Crick Institute, 1 Midland Road, London NW1 1AT, UK; IQS, Universitat Ramon Llull, Via Augusta 390, 08017 Barcelona, Spain; Zeeman Institute for Systems Biology and Infectious Epidemiology Research, University of Warwick, Coventry CV4 7AL, UK

**Keywords:** Dynamical systems, single-cell genomics, machine learning, unstable manifolds, catastrophe theory, developmental biology, cell fate decisions, stem cell biology, neural tube, sonic hedgehog

## Abstract

Building a mechanistic understanding of cell fate decisions remains a fundamental goal of developmental biology, with implications for stem cell therapies, regenerative medicine and understanding disease mechanisms. Single-cell transcriptomics provides a detailed picture of the cellular states observed during these decisions, but building dynamic and predictive models from these data remains a challenge. Here, we present *dynamic landscape analysis* (DLA), a framework that applies dynamical systems theory to identify stable cell states, map transition pathways, and generate a predictive cell fate decision landscape from single-cell data. Applying this framework to vertebrate neural tube development, revealed that progenitor specification by Sonic Hedgehog (Shh) can be captured in a landscape with an unexpected topology in which initially divergent lineages converge to the same fate through multiple distinct routes. The model accurately predicted cellular responses and cell fate allocation for unseen dynamic signalling regimes. Cross-species validation using human embryonic organoid data demonstrated conservation of this decision-making architecture. By modelling the dynamic responses that drive cell fate decisions, the DLA framework provides a quantitative and generative framework for extracting mechanistic insights from high-dimensional single-cell data.

## INTRODUCTION

During embryonic development and tissue homeostasis, cells make sequential fate decisions to produce the diverse sets of specialised cell types that form functional tissues. These decisions are driven by gene regulatory networks (GRNs) that interpret external signals and coordinate cell-type-specific gene expression programmes ^1^. Understanding how signalling inputs and GRNs coordinate these fate decisions remains a fundamental challenge in biology.

Recent work has approached this problem by using dynamical systems theory to formalise Waddington’s pioneering metaphor ^2^ of cellular differentiation as balls rolling down a landscape of branching valleys ^3–25^. This view is based on the idea that cell states correspond to so-called attractors, which are the stable states of the dynamical system generated by the GRN. Transitions between cell states occur through bifurcations that destabilise the initial state. During these transitions, cells follow a path of steepest descent by traversing a saddle point and descending towards a new attractor. Mathematically, this path is defined as the unstable manifold of the saddle point, with the saddle point governing where cells choose between alternative fates ^17,19^. Fate decisions arise from signal-induced changes in attractor stability and landscape topology combined with stochasticity. This framework provides a precise mathematical description of cellular states and transition routes, offering powerful insights into developmental processes. In particular, it uniquely distinguishes between different bifurcation mechanisms that control cellular transitions, such as those governing branching decisions between alternative fates ^17,19^.

Despite these theoretical advances, there are still relatively few systems analysed in this way. Consequently it is unclear if the theoretically predicted bifurcations, which fall into a limited number of generic types ^17^, are sufficient to describe cell fate decisions. In part this is because the construction of quantitative landscape models from experimental data remains a challenge. This is especially true for models attempting to capture developmental dynamics from high dimensional scRNA-seq data. The challenge is compounded by the need for quantitative models in a low dimensional space that faithfully reproduce the underlying bifurcation mechanisms governing cell fate branching.

The relative lack of applications of the dynamical systems approach to scRNA-seq data impedes progress because comprehensive gene expression information is essential for understanding the regulatory mechanisms underlying cell fate decisions. Current practice ^26,27^ usually involves selecting all highly variable genes followed by dimensional reduction using linear and non-linear projections such as principal component analysis (PCA) and UMAP. However, this approach presents some critical obstacles for dynamical systems analysis of cell fate transitions. In particular, methods for identifying attractors and unstable manifolds require working in gene expression space, but applying these techniques to thou-sands of dimensions is computationally intractable, and may be unnecessary since experimental evidence indicates that cell fate decisions are governed by relatively small GRNs ^1^.

Here, we present a framework, which we call *dynamic landscape analysis* (DLA), which addresses these challenges. This comprises the tools to identify the structure of a dynamical landscape and to construct quantitative models of the landscape from single-cell data. Our approach finds cell states and transition pathways by identifying gene expression spaces of reduced dimension *d* ≈ 50-100 that preserve the structural details of higher-dimensional scRNA-seq data while explicitly including the genes governing transition dynamics. We avoid the use of nonlinear projections and use simple linear 3D visualisations that maintain direct connections to individual gene expression levels, enabling computationally tractable and biologically interpretable analysis.

We applied this to an in vitro system of ventral neural tube development. In the neural tube, progenitor cells organise into discrete gene expression domains linearly arrayed along the dorsal-ventral axis. The positioning of these domains is determined by Sonic Hedgehog (Shh) signalling from the ventral pole. This developmental patterning can be recapitulated in vitro using mouse embryonic stem cells exposed to defined Sonic Hedgehog (Shh) signalling regimes ^28–30^ (Fig. 2A). Profiling transcriptomes of individual cells during this differentiation process offers a high-resolution view of cell states and fate transitions.

Our analysis revealed both well-characterised progenitor states and previously unappreciated intermediates, situated within a complex landscape controlled by Shh signalling. We discovered an unexpected topology involving connections between multiple cell states where initially divergent lineages converge to the same set of fates through distinct routes. We used these insights to construct a mathematical model that accurately reproduced the observed cell fate decisions.

The model revealed several key features of neural tube patterning that suggest general principles. It identified the bifurcation mechanisms underlying branching decisions, confirming that simple, theoretically predicted bifurcations are ubiquitous. Moreover, model-guided experiments showed how Shh signalling level and timing control both the proportions of cell types and the reversibility of the bifurcations that destablise cell states.

The findings revealed a novel patterning mechanism. Conventionally, neural progenitor domains are thought to form along the dorsoventral axis of the neural tube through a French flag mechanism in which distinct domains arise sequentially at monotonic morphogen concentration thresholds. However, our results challenge this view. We found that two boundaries form through branching decisions, represented by flip bifurcations, with no direct differentiation pathway connecting the cell states on either side of these boundaries. This reframes our understanding of how morphogen gradients establish patterning thresholds. Rather than operating purely through simple concentration-dependent switches, Shh signalling organises spatial pattern hierarchically, thereby expanding the repertoire of developmental patterning mechanisms.

Finally, analysis of human embryonic organoid scRNA-seq data ^31^ demonstrates that this decision-making architecture is conserved between mouse and human. The dynamical landscape model accurately predicted cell state proportions despite species-specific differences in developmental timing and identified which decisions in the network were affected by notochord-derived Shh. This cross-species validation establishes the fundamental nature of the identified landscape topology in ventral neural progenitor specification.

## RESULTS

### Identification and Validation of Attractor Clusters in scRNA-seq Data

To prototype our approach, we took advantage of an in vitro system in which pluripotent mouse embryonic stem cells were differentiated into distinct neural progenitor subtypes in response to Sonic Hedgehog signalling ^30,32,33^ (Fig. 2A). We analysed a scRNA-seq dataset ^33^ comprised approximately 40,000 cells across ten time points, generated in response to continuous exposure to 500nM SAG, a Shh signalling agonist (Fig. 2B). Based on prior knowledge of ventral neural tube development^30,32–35^, we expected to identify neuromesodermal progenitors (NMPs) ^28^, the PreNeural state ^36^ linking NMPs to neural fates, mesoderm cells ^28^, and the ventral neural tube progenitor subtypes pMN, p3, and floor plate (FP) ^35^. We anticipated that p0/p1 and p2 neural progenitor states might be present but rare at this high SAG concentration ^30,33,37^.

#### Outward clustering identifies cell states through iterative gene selection

We developed an iterative gene selection and clustering approach based on the principle that genes relevant to the GRN should show differential expression between cell states or transition pathways (detailed in Appendix A Sect. A1.1). Since cell states are initially unknown, we bootstrapped the analysis using a small number of marker genes (typically three or four) that define known cell types, regions or pathways. For early time points (D3-D4), we used TBXT/BRA, Foxc2, Foxa2, and Olig2 which identify neuromesodermal progenitors, mesodermal tissues, and early neural lineages ^30^. We identified subsets of cells expressing just one of these markers and performed differential gene expression analysis between these subsets. This process generated gene modules of approximately 100 genes, determined by setting statistical significance thresholds. For later time points, we used Olig2, Nkx2.2, and Shh because these identify pMN, p3, and floor plate progenitors and performed a similar analysis (Fig. 2C-D).

Although the early and late gene modules produced in this way define relatively high-dimensional data spaces (which we call *effective gene spaces*), they are reduced in dimension by a factor of about 30 compared to the dimensionality of all highly variable genes. This allowed almost all analyses to be carried out in gene space, as advocated in ^38^, without resorting to low-dimensional projections, except for visualisation. This, for example, allows us to compute cell-fate differentiation paths in the expression space as opposed to tracing them on a 2D or 3D visualisation. The only exception was for the fine-resolution Leiden clustering for which we projected the data into a lower-dimensional PCA space ^39^. Following this, we merged fine-scale clusters into cell state clusters based on their molecular identities (Fig. 2C).

We confirmed that we had captured the genes that are differentially expressed between each cluster and the others. We checked for missed clusters by examining unassigned cells for potentially missed states and testing whether additional genes showed significant differential expression between cluster pairs. We also confirmed robustness by substituting alternative correlated marker genes and verifying that cluster structures remained consistent. After analysing transition pathways (described below), we expanded the gene set to include genes showing significant variation in transitions, along with expert-curated transcription factors. None of these additions substantially affected subsequent analyses. We termed this approach *outward clustering*.

#### Validation of attractor clusters

To validate whether the identified cell states (summarised in Fig. 1), which we termed *attractor clusters* (ACs), corresponded to attractors of the underlying gene regulatory network, we used multiple criteria. First, we tracked each AC over time to assess its stability at successive time points, identify any loss of stability, and characterise resulting transitions to adjacent ACs. We extracted groups of neighbouring ACs and their associated transitioning clusters, we term these *sub-landscapes*, and projected them into 2- or 3-dimensional linear discriminant analysis ^40^ (LDA) space to visualise the transitions and temporal dynamics (Fig. 2E-F). LDA allowed visualisation while preserving data geometry and optimising cluster separation. For example, the previously unappreciated Early p3 state together with the p3 and FP cells states and the transition routes make up such a sub-landscape where the Early p3 state loses stability causing cells to transition to either p3 or FP, consistent with published observations ^41^ (Fig. 2F). We further validated transitions using the transition index of Mojtahedi et al. ^16,42^. Secondly, Wilcoxon rank-sum tests confirmed that known progenitor marker genes distinguished ACs. Thirdly, we measured Kullback-Leibler divergence (KLD) in LDA space (Appendix A Table A1) which provides a lower bound for KLD in the full gene expression space. Fourthly, cell distributions within each AC were checked to be unimodal along linear discriminant components, consistent with approximately linear dynamics near attractors (Fig. 2E). Finally, we examined gene-gene correlations within and between ACs, with differing correlations providing evidence of distinct regulatory interactions in each cluster (Appendix A Sect. A1.3).

**Figure 1.**
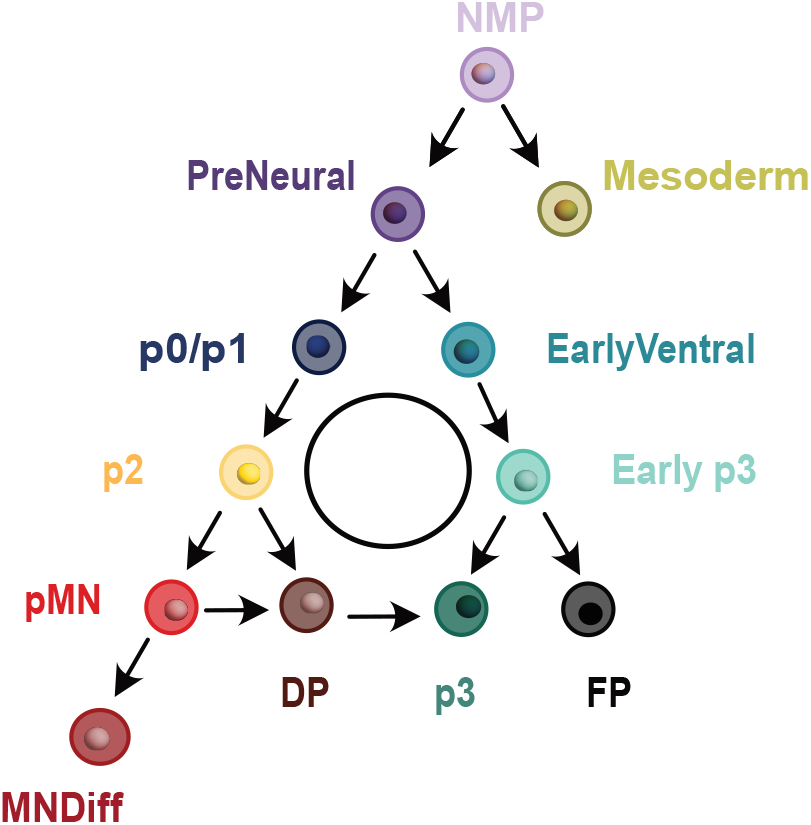
Neural Tube Decision Landscape. Schematic diagram summarising all the ACs associated with the cell states found in the scRNA-seq dataset at 500nM SAG across all time points (D3-D7). This is included to introduce the complete set of states in the landscape and the possible transitions between them. These will be discussed in later sections.

**Figure 2.**
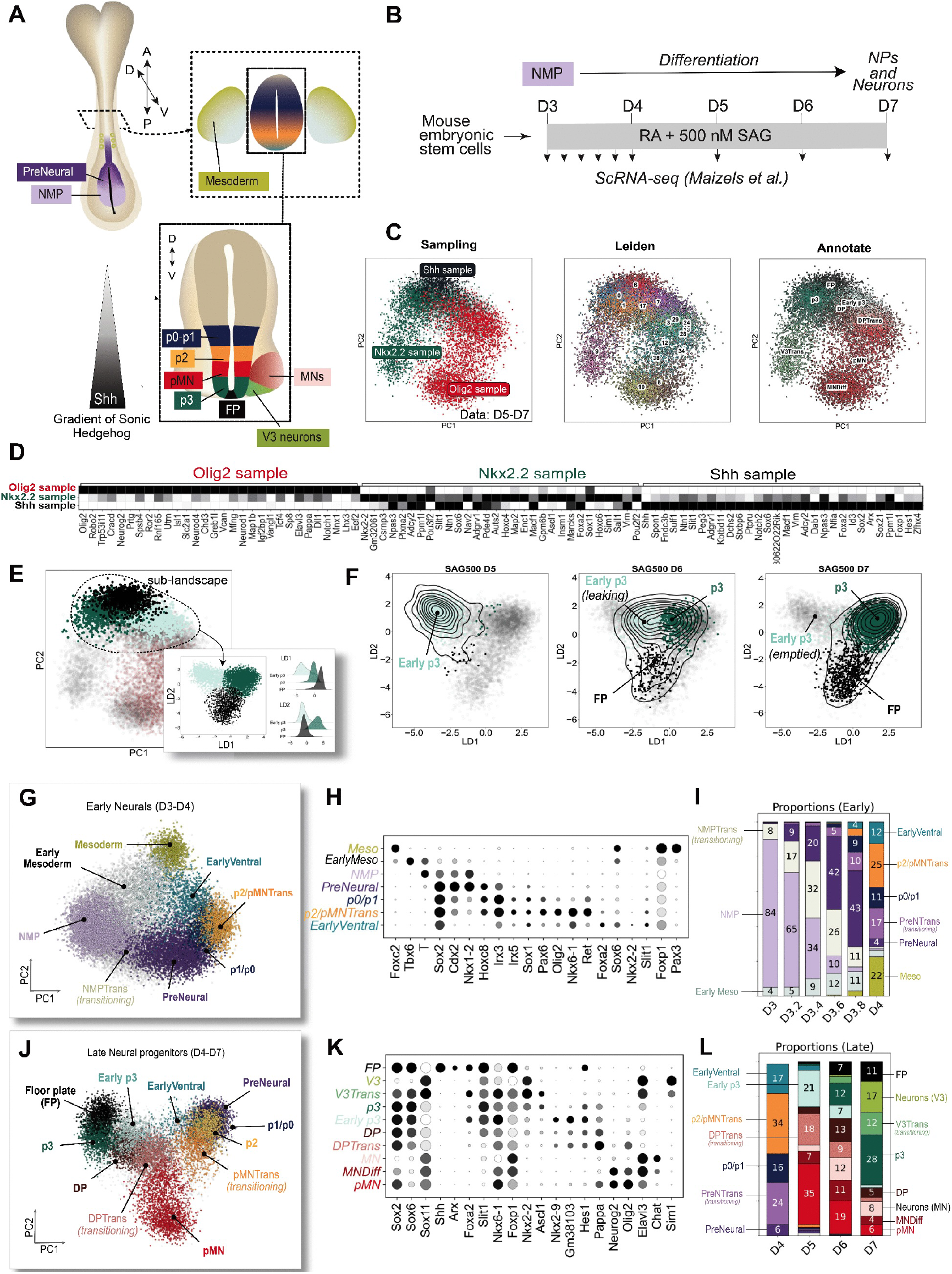
Attractor Clusters in Single-Cell RNA Sequencing Data Reveal Discrete Cell States During Neural Development. **A**. Schematic diagram of neural tube patterning regulated by the Shh gradient. **B**. Schematic of in vitro differentiation of mESCs and timing of the scRNA-seq data collection (D, Day). **C**. Example of the outward clustering methodology illustrated by analysis of the late time points (D5-D7). (Left) 2D PCA projection of cells that express one of the marker genes Olig2, Nkx2.2 and Shh. (Middle) PCA projection from *d*-dimensional effective gene space using the *d* most significantly differentially expressed genes between the 3 samples of cells (here *d* = 112). Cells are clustered using Leiden clustering with fine resolution and coloured accordingly. (Right) The fine Leiden clusters are merged based on their molecular identities. **D**. The most significantly differentially expressed genes distinguishing the 3 samples of cells shown in C (left panel). **E**. Extraction of the group of adjacent ACs, Early p3, p3 and FP, visualised in 2D LDA space. Histograms for each linear discriminant score (i.e the dot product of the data vector with the ith LDA eigenvector) show that ACs (such as FP) exhibit a well-defined approximately Gaussian structure in LDA space. **F**. Temporal progression of Early p3, p3, and FP identifies Early p3 as the (“head”) state from which cells transition to either p3 or FP. Grey dots represent all cells in the group (D5-D7), coloured dots indicate cells present at the indicated time points with the colour indicating their cluster assignment. Contour lines indicate the density of cells at each time point. **G.-I**. Identification of early ACs (D3-D4) and transitioning cells between them labelled -Trans. **G**. PCA projection of cells from D3-D4 samples showing the early ACs. **H**. Dot plot indicating the expression of markers used to identify the early ACs. The size of the circle represents the percentage of cells in the AC and the intensity of shading indicates the mean expression of the gene. **I**. The proportion of cells (in percent) in each AC and transitioning clusters at the early time points (D3-D4). **J.-L**. Identification of late ACs (D4-D7) and transitioning cells between them, labelled as -Trans. Similar to G.-I. but using cells from the D4-D7 samples. The D8 sample is not displayed, it predominantly contains MNs, V3 neurons, MNDiff cells, and p3 cells.

The identified cell states and several intermediates span early mesoderm specification to ventral neural progenitors ^28^(Fig. 1 and 2G-L).

At early timepoints, ACs included NMPs ^43^ (TBXT/Bra), mesodermal cells ^44^ (Foxc2), PreNeural cells ^36,45^ (Nkx1.2, Msx1, Sox2), transitioning NMPs (NMPTrans), and Early Mesoderm ^46,47^ (Tbx6). As differentiation progressed, ventral neural progenitor states emerged: pMN ^48^ (Olig2), p3^49^ (Nkx2.2), and FP^50^ (Foxa2, Arx, Shh). The p0/p1 (Pax6, Irx3) and p2 (Pax6, Irx3, Nkx6.1) ACs appeared at low numbers at D3.8 and D4, consistent with high SAG concentrations.

Additional states and genes that characterise them included MNDiff (Olig2, Neurog2), representing pMN progenitors undergoing motor neuron differentiation ^37,51^, and DP (Olig2, Nkx2.9, Hes1), representing an intermediate between p3 and pMN identities ^52^. The EarlyVentral population (Nkx6.1, Foxa2) at D3.8-D4 represents a common precursor of FP and p3^30^. At D5, we identified Early p3 (Nkx2.9), distinct from conventional p3 (Nkx2.2, Ascl1) observed at D6-D7^41,53,54^, and V3 neurons (Sim1, Nkx2.2), the differentiated p3 progeny ^49^.

Overall, this analysis demonstrates that outward clustering robustly identifies all known cell states and reveals previously undescribed developmental intermediates.

### Analysis of Cell State Transitions Using Unstable Manifold Mapping

We next set out to identify transition pathways and the bifurcations underlying branching trajectories. Cells change state through two related mechanisms: either an attractor is destroyed by collision with a saddle point (a *fold bifurcation*, also called a *saddle-node bifurcation*, Fig. 3A), or stochastic fluctuations drive escape over a nearby saddle. In the latter case it is expected that the attractor will be close to bifurcation with the saddle and we refer to both situations collectively as a *near-bifurcation escape*. In both cases, the cell’s transition to a target attractor is via a well-defined escape path determined by the saddle’s *unstable manifold*, although stochastic fluctuations can influence the dynamics along this path. This framework follows from dynamical systems theory and the hypothesis that GRN-driven stochastic dynamics underlie cellular differentiation and decision-making ^17–19^. Importantly, this principled approach avoids ad hoc choices inherent in other methods for identifying transition paths.

**Figure 3.**
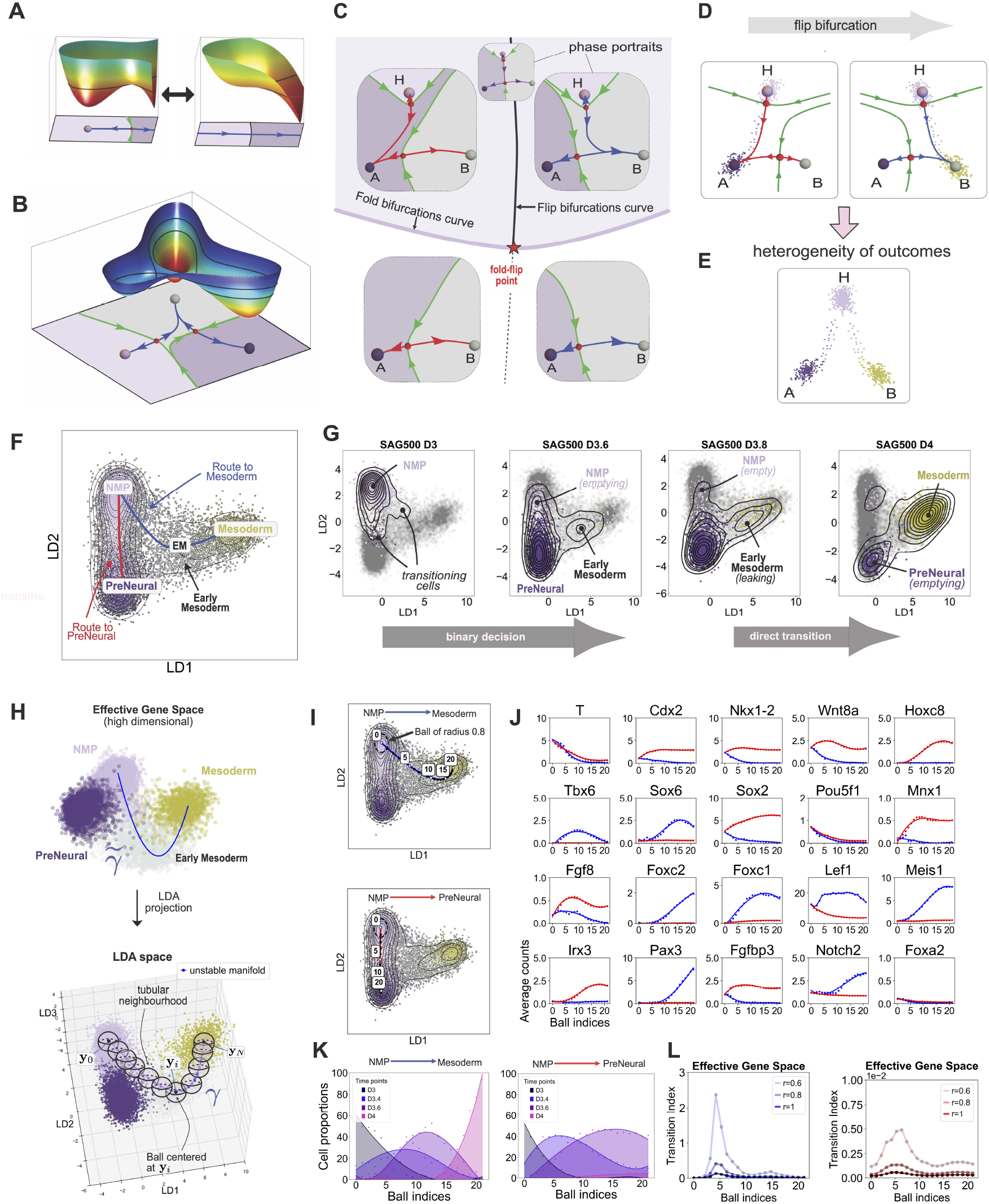
Analysis of Cell State Transitions Using Unstable Manifolds Between Attractor Clusters. **A**. Illustration of a fold bifurcation with a potential. (Left) Cells remain in the attractor which sits at the bottom of the potential well. There is a saddle close, over which some cells might escape. The blue curve is the approximated unstable manifold of the saddle. Escaping cells will, subject to noise, travel along this. (Right) A change in parameters leads to a collision with the saddle and the destruction of the attractor and saddle, causing all cells to escape. **B**. An illustration of the potential of a binary flip landscape. Cells start in the left hand attractor and, if they escape, they transition along the blue approximate unstable manifold to the grey attractor at the top of the illustration. A change in the parameters can result in the unstable manifold flipping so that cells transition to the bottom right attractor (dark purple) instead of the top attractor (grey). **C**. 2D Bifurcation set of a binary flip landscape. Fold and flip bifurcations curves divide the parameter space into region in which the landscape dynamics is distinct. When a change in parameters causes them to cross the fold curve, the head attractor (H) undergoes a fold bifurcation (i.e. it collides with a nearby saddle and disappears). **D**. When a change in parameters in C causes them to cross the flip curve, the system undergoes a flip bifurcation in which the unstable manifold of the saddle close to the H attractor changes its connection, flipping from attractor A to attractor B. Dots reprepresent cells escaping the H attractor and transitioning to a downstream attractor (A or B) along the unstable manifold. **E**. Heterogeneity of the cells in the head attractor H implies that their dynamics is driven by distinct landscapes, one with unstable manifold connecting to A and other connecting to B. **F**. Binary decision from NMP to PreNeural and Early Mesoderm/Mesoderm. The two approximated unstable manifolds (red and blue curves) for the landscapes of cells projected into LDA space. The red approximated unstable manifold outlines the route to PreNeural and the blue unstable manifold the route to Early Mesoderm and Mesoderm. **G**. Temporal progression of the NMP, PreNeural, Early Mesoderm and Mesoderm ACs in LDA coordinates. Grey dots represent all cells in the group, coloured dots indicate cells present at the indicated time points with the colours indicating their cluster assignment. Contour lines indicate the density of cells at each time point. This exemplifies the binary flip decision from NMP to PreNeural or to Early Mesoderm, followed by a direct transition from Early Mesoderm to Mesoderm. At D3 almost all cells (dots) are in the NMP AC. Cells in both PreNeural and Early Mesoderm (Tbx6^+^) are evident at D3.6, and increase at D3.8. There is then a direct transition from Early Mesoderm to Mesoderm at D3.8 and PreNeural empties as cells transition to other states (see Appendix A Fig. A1G). **H**. (Top) The unstable manifolds are estimated in the higher dimensional effective gene space. This avoids artificial ordering of the points along the curve that might occur if, for example, the LDA components were strongly determined by a single gene. This curve is then projected into the LDA space of interest (2D or 3D). (Bottom) Balls are placed along the projected curve in LDA space. Together they form a tubular neighbourhood of the unstable manifold. Raw counts (or, equivalently, log-normalised counts) of individual genes are averaged over the cells inside each ball to determine how gene expression changes along the unstable manifold. This construction ensures that the change in expression levels does not depend upon the particular LDA projection used. **I**. LDA plots of the binary flip decision from NMP to either PreNeural or Early Mesoderm using D3-D4 data showing the projected unstable manifolds. The positions of the balls are shown by the index of their centers. A ball of radius 0.8 centered at index 0 is shown (dashed black). **J**. The variation of key genes along the estimated unstable manifolds from I. Genes such as Cdx2, Sox2, Nkx1.2 and Tbx6 show diverging behaviour along the two routes consistent with the expected behaviour in a binary flip landscape. **K**. Temporal progression of cells along the projected unstable manifolds in I. Proportions of cells (dots) at stages D3, D3.4, D3.6, and D4 within each ball (of radius 0.8) along the unstable manifolds connecting NMP to Mesoderm (left) and NMP to PreNeural (right). We used a high smoothing factor to obtain smooth densities. In the right panel, there are very few cells at D4 (pink density) because cells at this stage have exited the PreNeural AC (see Appendix A Fig. A1G). **L**. Critical Transition index ^42^ computed in each ball along the approximated unstable manifolds show peaks as cells transition from one AC to another. Colour variations indicate different ball radii.

#### Direct transitions and binary decisions

When a near-bifurcation escape takes place, the simplest transition that occurs is that all the cells escaping an attractor follow the same path to the same downstream attractor (Fig. 3A). These *direct transitions* involve no branching and commit cells to a single fate. The next simplest outcome is that, depending on signals received by transitioning cells, they choose between two potential escape routes. The simplest branching architecture underlying this is the *binary flip sub-landscape*, which contains three attractors and two saddles (Fig. 3B). Cells enter at the head attractor (H in Fig. 3C-D) and remain there until near-bifurcation escape drives progression to one of two downstream attractors (A or B in Fig. 3C-D). The choice between these fates depends on the signals received and the cell’s state. Changes in signalling or cell state can redirect the escape route to the alternative downstream attractor (Fig. 3D), thereby changing cell fate. This landscape is constructed from two generic bifurcations: the *fold bifurcation* that can destroy the head attractor, and the *flip bifurcation* that switches the escape route between the two downstream states (Fig. 3C).

Populations of cells typically exit the head attractor via both routes, splitting into two groups with distinct gene expression trajectories. We identified the NMP, PreNeural, and Early p3 attractors as head attractors of such binary flip sub-landscapes, while the p2 attractor serves as the head of a slightly more complex but related architecture. Fig. 3E schematically illustrates a case in which cells divide into two types that culminate in distinct downstream attractors. For example, at the PreNeural attractor, cells with high Foxa2 expression take one route while those with lower expression take the other ^30^ (Appendix A Fig. A5).

The binary decision from the NMP head attractor to either neural or mesodermal fates ^28,29^ is a canonical example of a binary flip. LDA projection reveals clear separation between NMP, PreNeural, Early Mesoderm, and Mesoderm attractors, with transitioning cells forming bridges between populations (Fig. 3F). At D3.2, cells diverge from NMP along two distinct trajectories: one leading to PreNeural and another to Early Mesoderm (D3.4), which subsequently transitions directly to Mesoderm at D4 (Fig. 3G).

#### Transition analysis

To identify transition pathways given by unstable manifolds and validate the bifurcation structures, we developed a method of gene expression analysis using tubular neighbourhoods along the transition paths (Appendix A Sect. A2). We approximated the unstable manifolds in the effective gene space by fitting splines to the data between ACs (Fig. 3H-I). We then computed the mean gene expression level along these tubular neighbourhoods (Fig. 3J). This approach detects genes that vary significantly during transitions, including genes that are not differentially expressed between ACs but have nontrivial dynamics during the transition, thereby identifying potential GRN components. In addition to the genes defining the effective gene space, we tracked approximately 150 curated transcription factors relevant to neural development.

This approach was used to validate binary flip architectures, including escape from a head attractor immediately before branching, by revealing distinct expression trajectories that quantitatively distinguished exit routes from the head attractor. In every binary flip landscape, transitioning genes showed significantly different behaviour along the two exit routes. For example, for the head NMP attractor this revealed upregulation of Nkx1.2 and Irx3 for escape along the PreNeural pathway and of Tbx6 and Foxc2 for the Mesoderm pathway (Fig. 3J), providing strong support for the binary flip architecture.

The temporal progression of cells transitioning along the two unstable manifolds of this flip landscape is illustrated in Fig. 3K, showing how the distributions of all cells within the tubular neighbourhood at different time points progress along the unstable manifold. A broader variance in the distributions (e.g. at D3.4) supports the hypothesis that cells may experience delays and do not transition simultaneously.

The critical transition from ACs was further validated by tracking the transition index of Mojtahedi *et al*. ^16,42,55^ which tends to show a sharp increase at exit (e.g. Fig. 3L for exit from NMP). We applied similar analyses to the other branching sub-landscapes (details in Appendix A Sect. A2.4).

### The effect of changing Shh levels on the landscape

The route that a cell takes and its end fate depends critically upon the level of Shh signal. Therefore, we set out to understand how Shh signalling shapes the decision landscape by creating or destroying attractors (fold bifurcations) or altering the transition routes between them (flip bifurcations). To this end, we identified five markers -Sox2, Pax6, Olig2, Nkx6.1, and Nkx2.2 - sufficient to classify most ventral progenitor populations ^30^. Using these markers, we generated three independent time resolved flow cytometry datasets using four SAG concentrations (0, 10, 100, 500nM) at days D4, D5 and D6 of differentiation (Fig. 4A). We did not include the D3 sample that contained only NMPs. We selected neural progenitors (Sox2^+^) and used Gaussian Mixture Models to identify clusters in the remaining 4-dimensional marker gene space ^18^ (Appendix A Sect. A3). Using statistical validation criteria analogous to those applied to the ACs from the scRNA-seq data (Appendix A Sect. A3.3) we identified the major progenitor populations (Fig. 4A-C), analysed the local transitions between them, and tracked their proportions over time across SAG concentrations (Fig. 4D). The proportions and the molecular composition for the transitioning populations for each experimental replicate are presented in Appendix A Fig. A9B-C. Unlike the scRNA-seq dataset, MNs and V3 neurons were not identified in the flow cytometry because only Sox2^+^ neural progenitors were selected for analysis.

**Figure 4.**
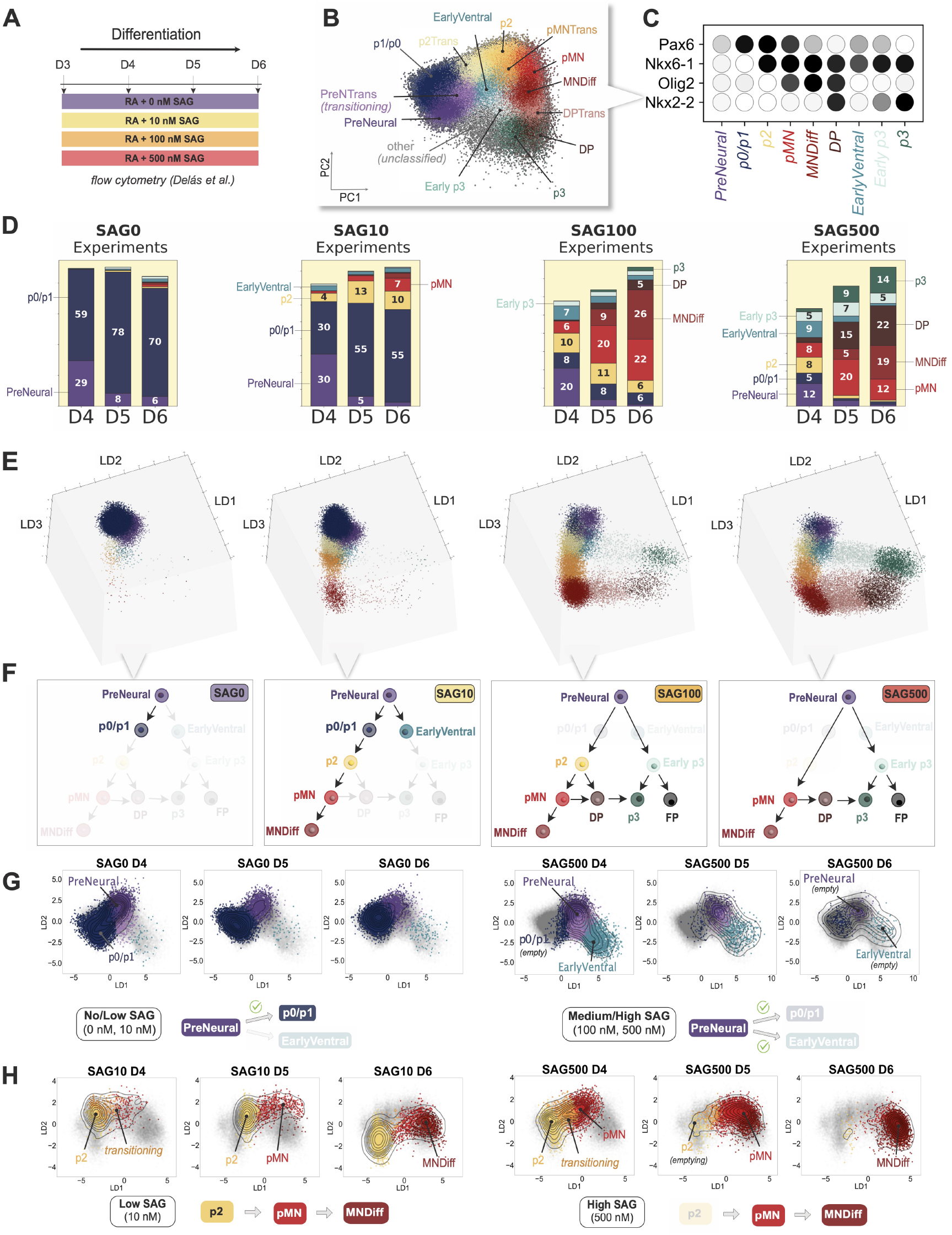
Flow Cytometry Analysis of Cell States Reveals SAG Concentration-Dependent Neural Progenitor Specification. **A**. Schematic of the signalling conditions used for flow cytometry. **B**. 2D PCA projection of the ACs found using flow cytometry data. Data for all SAG concentrations and time points are combined. **C**. Dot plot indicating the expression of markers used to characterise the ACs in B (for the other clusters see Appendix A Fig. A8B). The Floor Plate cluster (FP) was not identifiable in this dataset (see discussion in Appendix A Sect. A4.2). **D**. Percentage occupancy of the ACs as a function of Day (D) and SAG concentration (0, 10, 100, 500nM SAG). This shows the average proportions over 3 experimental replicates for 10nM and 100nM and 5 replicates for 0nM and 500nM (Appendix A Sect. A3). Blank spaces in bar plots correspond to the proportions of transitioning and unclassified cells, these are shown in Appendix A Fig. A8C for all replicates. **E**. 3D representation in LDA coordinates of the clusters at all time points (unclassified cells excluded). At 0nM SAG cells remain confined to the p0/p1 AC but start transitioning to p2 and pMN at 10nM SAG. The ventral cell branch (green/cyan dots) that includes EarlyVentral, Early p3, and p3 only appears at high SAG concentrations (100nM, 500nM). **F**. Schematic diagrams of attractors present at different SAG levels and the connections between them. This represents the consensus state because a landscape is associated with an individual cell, which are heterogeneous in state and signal. **G-H**. Temporal evolution of selected ACs (D4-D6) in LDA coordinates as a function of time and SAG concentration. Grey dots represent all cells in the group, coloured dots indicate cells present at the indicated time points with the colours indicating their cluster assignment. Contour lines indicate the density of cells at each time point. **G**. Binary flip decision from PreNeural to either p0/p1 or EarlyVentral. The figure shows 0nM, 500nM SAG (see Appendix A Fig. A13A for other concentrations). (Left) The p0/p1 AC is stable when SAG is absent (0nM) and there is no transition to EarlyVentral. (Right) The transition to EarlyVentral only appears at high SAG concentrations (100nM, 500nM). PreNeural, p0/p1, and EarlyVentral empty over three days at the highest SAG level (500nM). The attractors have bifurcated, but the unstable manifolds from PreNeural to pMN and Early p3 still pass through the regions in gene space formerly occupied by the bifurcated attractors. **H**. Direct transitions from p2 to pMN and from pMN to MNDiff. The figure shows 10nM, 500nM SAG (see Appendix A Fig. A14A for other concentrations). (Left) At 10nM SAG, the p2 AC is relatively stable but shallow as cells can escape and transition to pMN over time. (Right) The same transition at 500nM SAG where p2 has already bifurcated showing more rapid occupation of the pMN AC.

This approach proved particularly informative for revealing critical bifurcations that result from varying morphogen concentration (Fig. 4E-H). Changes in SAG concentration resulted in different proportions of cell types and affected the topology of the connections in the landscape (Fig. 4F). Although we can distinguish the Early p3 AC from the p3 AC, using the limited markers available we cannot distinguish FP, which is expected to co-express Sox2 and Nkx6.1 and to be present at D6. Nevertheless, the scRNA-seq data indicate it should be present at 500nM SAG. We therefore included it as part of the schematic diagram in Fig. 4F.

A summary of all the ACs and transitions from the flow cytometry and scRNA-seq data is shown in Fig. 5. The ventral branch comprising EarlyVentral, Early p3, and p3 states emerged from PreNeural only at the higher SAG concentrations, indicating the occurence of a flip bifurcation (Fig. 4G) as verified in the scRNA-seq analysis (Appendix A Fig. A5).

**Figure 5.**
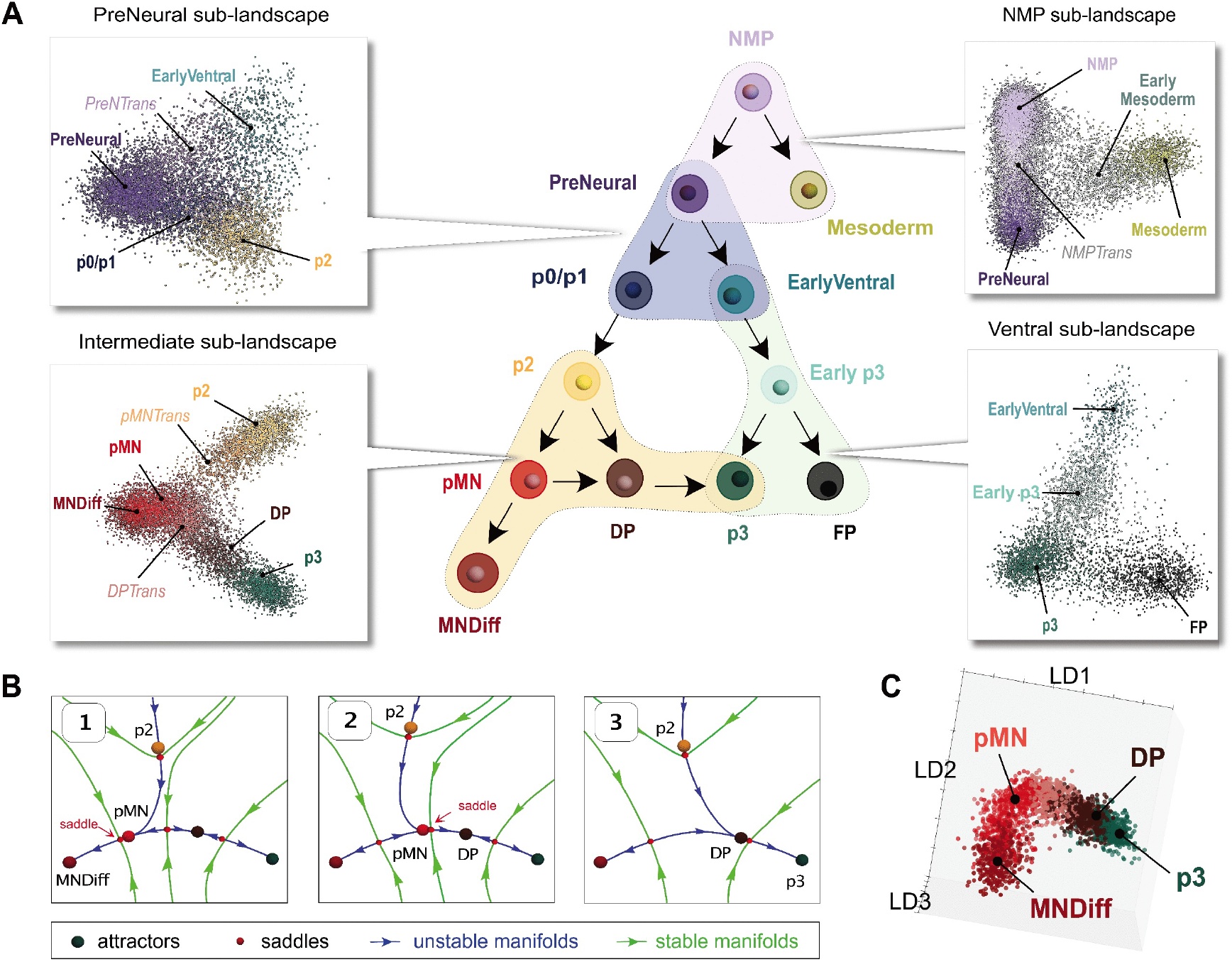
Sub-Landscapes Identification. **A**. Schematic diagram of all the possible transitions between cell states. A sub-landscape is defined as a group of neighbouring ACs and transitioning cells, projected in LDA space and highlighted on the diagram. A mathematical model should reproduce all the possible transitions. **B**. Distinct dynamical systems that model the transitions in the intermediate sublandscape. In Panel 1, p2 and pMN are close to their saddle and cells transition easily towards MNDiff. In Panel 2, the unstable manifold is directed towards DP and cells transition to DP via pMN. In Panel 3, the pMN attractor is bifurcated and DP is close to its right saddle, allowing a DP-to-p3 transition. **C**. The ACs MNDiff, pMN, DP and p3 form a smooth structure in LDA space, supporting our choice of using a linear landscape as in B to model the transitions from pMN to either MNDiff or DP.

The AC containing exclusively Pax6^+^ cells, labelled p0/p1, captures most cells at 0-10nM SAG (Fig. 4G). But, at higher concentrations, Nkx6.1 and Olig2 levels rapidly rise within the Pax6^+^ cells, leading to the emergence of p2 and pMN (Fig. 4H).

While the p2 AC is present and stable at 10nM SAG, it destabilises at higher SAG concentrations, with these cells instead adopting a pMN identity through up-regulation of Olig2 (Fig. 4H) or a DP identity through up-regulation of Olig2 and Nkx2.2. The pMN AC is populated by a few cells at 10nM SAG but increased occupation of pMN and MNDiff is apparent at 100-500nM SAG. By contrast, DP and p3, both expressing Nkx2.2, required a concentration above 100nM SAG to appear (Fig. 4D-F).

The time-resolved flow cytometry data confirmed the transition routes identified in the scRNA-seq data and revealed SAG-concentration dependent bifurcations (all remaining transitions are presented in Appendix A Sect. A4, Fig. A13-A14). Thus by combining transcriptional profiling with protein-level measurements, we established a comprehensive map of stable cell states and their connecting transition paths as a function of SAG concentration (Fig. 5).

### Mapping the Complete Neural Progenitor Landscape

The unstable manifold transition analysis proved robust to different projections and both data modalities, capturing the progression of cell fate during all neural progenitor transitions (Appendix A Sect. A2). Gene expression changes along transition routes found by both modalities revealed consistent patterns. Exit from a head AC is marked by the down-regulation of genes that define this AC and the simultaneous up-regulation of markers associated with the target ACs. Consistent with proposed multi-lineage priming mechanisms of fate commitment ^56^.

Together, the method captured the gradual gene expression changes during transitions and identified route-specific gene expression programs. This provided evidence for well-defined transition routes and critical fate decision points essential for a quantitative model of the developmental landscape.

Overall, we identified four sub-landscapes comprising ACs and transitioning cells, which we called respectively *NMP, PreNeural, intermediate* and *ventral* sub-landscapes (Fig. 5). We analysed each of these separately, following the approach used for the NMP sub-landscape (Appendix A Sect. A2). Each of the branching decisions shown, except those in the intermediate sub-landscape, is a binary flip bifurcation with exit paths selecting between the downstream states shown. The evidence for this is given in Appendix A Sect. A2.

The decisions in the intermediate sub-landscape are particularly interesting. PreNeural cells that transition past the p0/p1 AC proceed to p2 and enter this sub-landscape (Fig. 5 and Appendix A Fig. A6). The data indicate cells escaping p2 (or transitioning through it at higher SAG levels) proceed either to the pMN or DP ACs. This part of the sub-landscape can be regarded as a standard binary flip. However, the data indicated that the downstream states pMN and DP are part of a linear structure consisting of the four ACs MNDiff, pMN, DP and p3 with saddles between each of these (Fig. 5B). Here we describe the analysis of the gene expression along the approximate unstable manifolds (Appendix A Fig. A6C-E), and in later sections we fit a relatively simple mathematical model to the data that captures this more complex bifurcation.

Cells exiting p2 that progressed towards pMN upregulated Olig2 and Neurog2 whereas others showed coordinated upregulation of Olig2, Hes1, Rfx4 and Nkx2.9, indicating commitment to DP fate. The pMN-specified cells progressed through a Neurog2-high MNDiff intermediate state before terminal MN differentiation ^37^. On the other hand, we observed the progressive loss of DP cells by D7 with convergence towards p3 supporting a direct DP-to-p3 transition (Appendix A Fig. A7B). Lineage studies support this connection: while FP cells rarely derive from Olig2-expressing precursors, a subset of p3 progenitors descend from cells with an Olig2 expression history ^57^.

The observation that there is a flip between pMN and DP and escapes from pMN to MNDiff and from DP to p3 means there are three saddles the 1-dimensional unstable manifolds of which connect MNDiff to pMN, pMN to D3 and D3 to DP. Visualisation of the data (Fig. 5C) suggests that these three transition pathways link together smoothly resulting in the geometry shown in Fig. 5B. This is further justified by dynamical systems theory because the data show that the MNDiff, pMN and DP attractors and the saddles between them are close to each other (Appendix B Sect. B1.4). The elucidation of such uniquely detailed decision-making structure and the validation of this (see below) demonstrates the power of the approach.

### Constructing a Landscape Model

We next developed a mathematical model of the observed decision-making dynamics in response to different Shh signalling levels. Rather than modelling genegene interactions as in conventional GRN models, our approach constructs a quantitative model of the cell fate decisions at the population level including the bifurcation structure underlying these decisions. It can generate experimentally testable predictions because it can be used to simulate unseen situations such as changing morphogen conditions and missing time points. The model consists of deterministic differential equations, with parameters dependent on SAG concentration, and a space-dependent stochastic component.

The key step is the construction of the deterministic model component as this drives the decision-making. We constructed this by partitioning the system into overlapping local sub-landscapes corresponding to the binary flip and direct transition architectures identified above (Fig. 6). This enables the use of catastrophe and bifurcation theory ^58,59^ to find appropriate dynamical systems in normal form for each sub-landscape.

**Figure 6.**
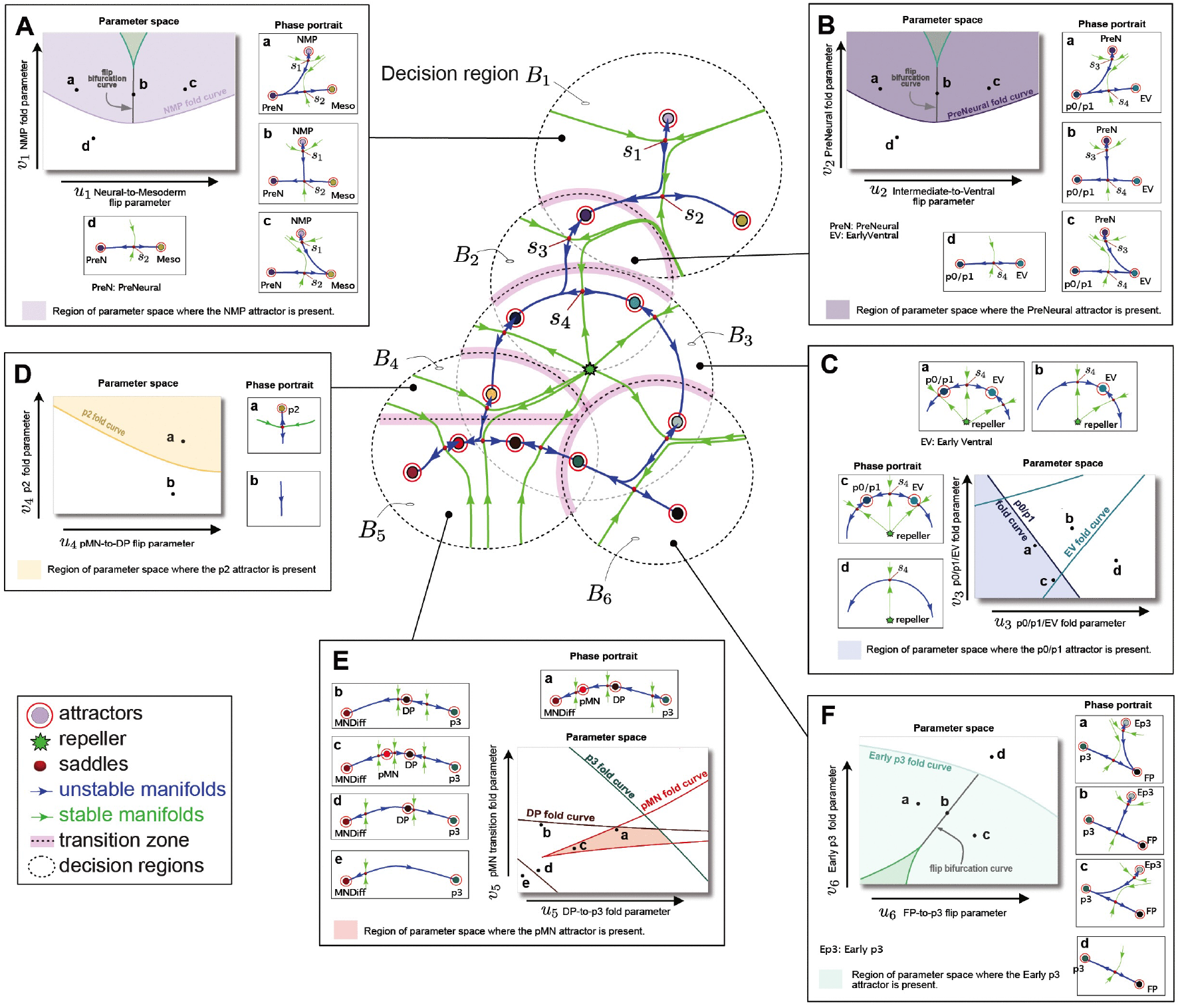
Constructing the full landscape model. The sub-landscapes used to construct the model showing one realisation of the unstable manifolds. Changing parameters allows unstable manifold flips or the bifurcation of attractors. The pink regions (transition zones) show where bump functions were used to glue the landscapes together. These divide the phase space into 6 domains and as the state crosses a transition zone its dynamics transition smoothly to the dynamical system governing the regions it has entered into. **A.-F**. Each panel shows the parameterised dynamics in the decision regions *B*_1_ − *B*_6_. These indicate the form of the bifurcation set which divides the parameter space into regions where the qualitative form of the dynamics is constant. They also show the configuration of the dynamics in each of these regions. As a bifurcation curve is crossed either a flip bifurcation occurs or one of the attractors appears or disappears (fold bifurcation). **A**. 2D Parameter space of the NMP sub-landscape with a fold parameter (y-axis) that controls the bifurcation of the NMP attractor and a flip parameter (x-axis) that controls the flip bifurcation from neural fates to mesodermal fates. **B**. Similar to A for the PreNeural sub-landscape. In this case the flip parameter controls the transition from intermediate (p0-p2-pMN) to ventral fates (EarlyVentral, Early p3, p3 and FP). **C**. Parameters that control the fold bifurcations of the p0/p1 and EarlyVentral attractors. **D**. Parameters that control the fold bifurcation of the p2 attractor. **E**. The dynamics of the MNDiff, pMN, DP and p3 attractors. The bifurcation set consists of two fold curves for the DP attractor (brown) meeting at a dual cusp (not shown) and two curves of fold points controlling the bifurcation of the pMN attractor (red), also meeting at a dual cusp (shown). This bifurcation structure is motivated by the experimental observation that cells can differentiate to motor neurons from pMN as well as transitioning to a DP fate by up-regulating Nkx2.9 and Hes1. This means that the saddles on both sides of the pMN attractor must interact with it. **F**. Parameters controlling the fold bifurcations of the Early p3 attractor and flips of the unstable manifold of the saddle near the Early p3 attractor that leads cells to either p3 or FP fates. The bifurcation set consists of a fold curve and a flip curve that meets the fold curve at a fold-flip point.

For systems with cell states that are non-periodic attractors, the possible topologies of the 1-dimensional unstable manifolds connecting the attractors are independent of the underlying state space dimension, provided it exceeds one ^17^. It is therefore reasonable that, for our system, a two-dimensional representation can capture the essential bifurcation structure and transition pathways present in the full high-dimensional gene expression space.

An advantage of using normal forms to describe the sub-landscapes is that it enables systematic understanding of how the parameters, which represent the effect of signalling, affect cell fate decision-making. Bifurcations drive cell fate decisions, hence understanding their organisation in parameter space is essential for establishing the genericity and robustness of the conclusions.

In 2-dimensional parameter space, bifurcation sets corresponding to each sub-landscape are made up of curves where fold or flip bifurcations occur. These curves divide the parameter space into regions with the same attractors and same connection topology in phase space. Crossing a bifurcation curve alters the decision-making structure either by creating or annihilating an attractor by interaction with a saddle, or by causing a flip bifur-cation.

The normal form approach provides precise control over which parameter drives each bifurcation type. The parameters function phenomenologically rather than representing specific biological rates and can be regarded as combinations or functions of the underlying biological parameters of the relevant GRN. They act as control variables that determine the system’s critical decision points. One parameter might control the timing of attractor destabilisation through a fold bifurcation. Another might govern the balance between alternative fates at a flip bifurcation. Thus bifurcation sets clarify how signalling levels translate into decision outcomes and reveals how population heterogeneity propagates through the decision landscape to generate outcome variability.

#### Box 1. Some details of the global landscape construction

**Modelling the individual sub-landscapes**

We define dynamical sub-landscapes as subsets of neighbouring attractors connected by 1-dimensional unstable manifolds within decision regions: attracting regions of phase space that constrain trajectories (*B*_1_-*B*_6_). The NMP sub-landscape (region *B*_1_) is a binary flip landscape containing the NMP, PreNeural, and Mesoderm attractors with two saddles (*s*_1_, *s*_2_), controlled by a flip and a fold parameter *u*_1_ and *v*_1_ (Fig. 6A). The PreNeural sub-landscape (*B*_2_) shares the PreNeural attractor with the NMP sub-landscape and has identical structure, with parameters *u*_2_ governing transitions towards intermediate (p0/p1) or ventral (EarlyVentral) fates and *v*_2_ bifurcating the PreNeural attractor (Fig. 6B). This connects to a circular sub-landscape with central repeller (*B*_3_), which allows p3 to be reached via two distinct pathways, where *u*_2_ can create heteroclinic connections between saddles *s*_3_ and *s*_4_. Parameters *u*_3_ and *v*_3_ control fold bifurcations of the p0/p1 and EarlyVentral attractors (Fig. 6C). The p0/p1 attractor connects to p2 in region *B*_4_, which mirrors the *B*_2_ structure (Fig. 6D). From p2, cells transition to pMN or DP, and continue through *B*_5_. This captures direct p2-to-pMN transitions, DP-to-p3 transitions, and pMN bifurcations to both MNDiff and DP (Fig. 6E). Finally, *B*_6_ contains a binary flip sub-landscape from Early p3 to either p3 or FP, with parameters *u*_6_ and *v*_6_ controlling the necessary fold and flip bifurcations (Fig. 6F). Details can be found in Appendix B Sect. B2.5.

**Connecting the sub-landscapes to obtain a global model**

The sub-landscapes are connected through simple topological relationships, with connections occurring along transition routes that avoid attractors and bifurcation-critical regions (Fig. 6 center and Fig. 7A). This design enables the sub-landscapes to be joined using differential topology principles ^60^ (Appendix B Sect. B2.4). The result is a differentiable dynamical system that depends smoothly on its parameters (Appendix B Sect. B2.5). The model qualitatively captures the key bifurcations observed in the experimental data, including fold and flip bifurcations.

**Dynamical stochasticity and cell-dependent dynamics**

To model the stochastic nature of the dynamics due to intrinsic noise arising from the randomness of biochemical reactions within a cell we include a spatially dependent diffusion term setting the noise level as an additional parameter. We allowed the noise to vary between the various regions of phase space, accounting for transition-specific noise variability (Appendix B Sect. B3.1). Finally, velocity parameters are included that allow the speed along trajectories to vary from region to region. This provides a highly flexible but minimally parameterised model that can accurately capture the behaviour in each local sub-landscape.

Non-genetic variability, arising from within-cell processes such as differences in transcription factor levels, plays an important role in determining cell fates alongside intrinsic noise arising from the randomness of biochemical reactions within a cell. To account for this non-genetic variability, we allow for variation in each cell’s parameters producing a distinct dynamic for each cell (Fig. 7B).

When we combine these various types of stochasticity we observe dynamics where cells starting in identical states can traverse widely different developmental trajectories (Fig. 7C). Moreover, the timing with which multiple cells follow the same trajectories tends to be highly variable because cells can remain trapped in shallow attractors for extended periods before escaping due to stochastic fluctuations, and then progressing relatively linearly along the unstable manifolds (Fig. 7C). We quantify this variation in timing below in Fig. 10D when we discuss the DP to p3 transition showing that it can be very large and represents a challenge to the notion of pseudotime.

**Figure 7.**
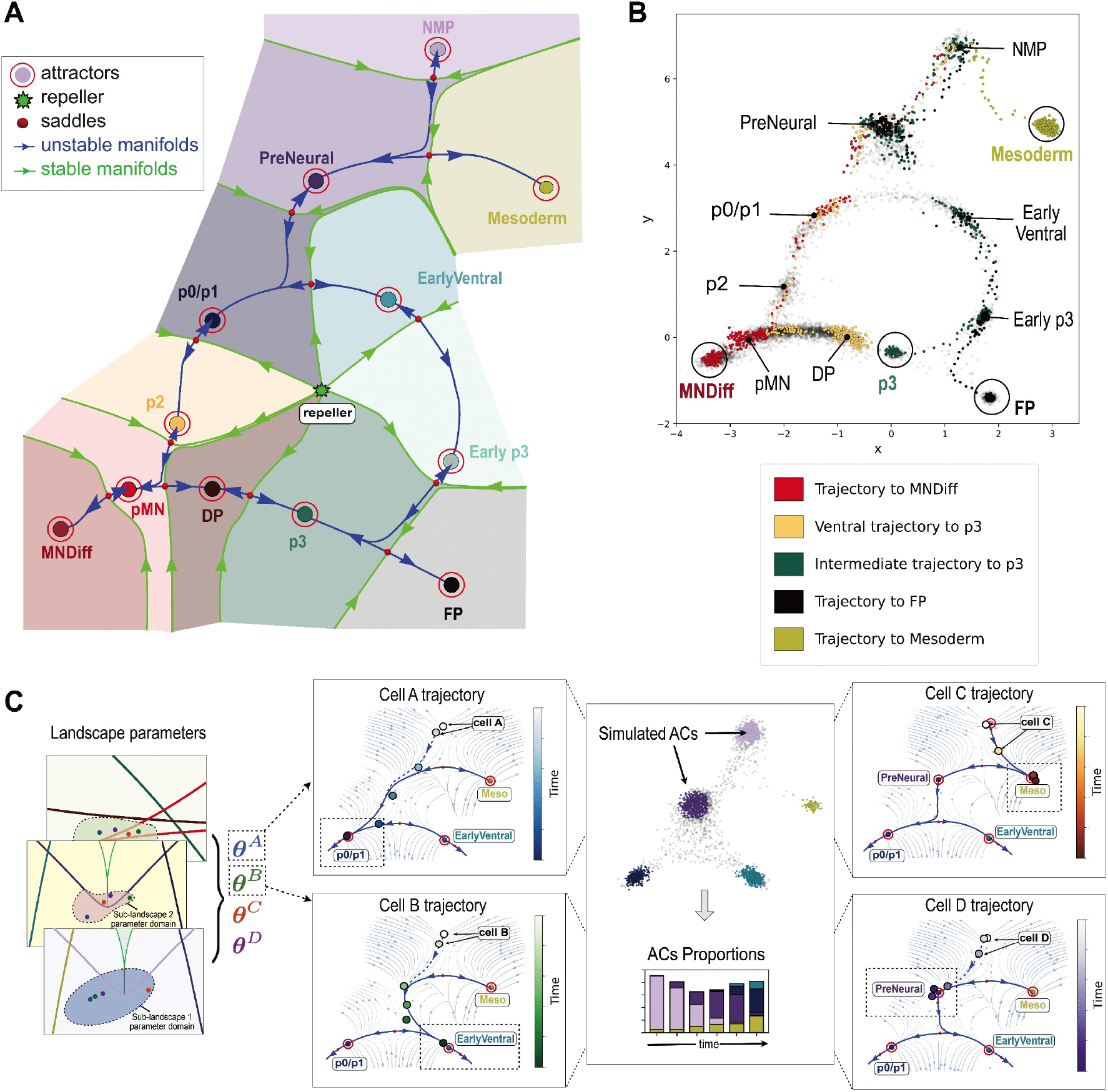
Global landscape simulation. **A**. Phase portrait of the global landscape for a fixed set of parameters when all attractors are present. The blue and green curves are respectively the unstable and stable manifolds of the various saddles. The green stable manifolds of the saddles separate the basins of attraction of the various attractors which are coloured accordingly. If the state of a reprogrammed cell falls in one of these coloured domains, it will converge towards the corresponding attractor and adopt that specific fate. One arbitrarily direction for each flip bifurcation is illustrated. **B**. Five stochastic single cell trajectories followed for 400 simulation timesteps coloured according to the eventual fate. Gray dots indicate other cell trajectories, included to help visualise the shape of the landscape. Notice that while pseudo-time makes sense for the transitioning segments of a trajectory, it does not make sense for a full trajectory because cells spend a random period of time near attractors. **C**. Schematic of the simulation procedure, illustrated for the first two binary decisions. For each cell *µ*, parameters for the individual sub-landscapes are selected within their respective parameter domains (two components in each 2D bifurcation panel represented by a dot coloured differently for each cell). Together, they form the components of a parameter vector ***θ***^*µ*^ of the global landscape (illustrated here for 4 cells labelled A, B, C and D with parameters ***θ***^*A*^, ***θ***^*B*^, ***θ***^*C*^, ***θ***^*D*^). These parameters define the configuration of attractors, saddles and unstable manifolds of the cell’s deterministic dynamics. Adding stochastic fluctuations to these dynamics, we simulate a stochastic trajectory for each cell, recording their positions in state space at discrete time points. Each of the panels on the left shows the deterministic (local) landscape for each cell determined by their respective parameters. In each panel, the evolving cell is coloured according to its time step. Different sets of parameters produce different landscape models resulting in distinct possible fates for each cell (dashed rectangles in each panel). If an attractor is present in a cell’s landscape (e.g PreNeural for cell D), the cell may remain trapped in this attractor as we see for the purple dots, indicating the position of the cell over time, staying near the PreNeural attractor. Expanding the parameter sampling allows the simulation of a large number of cells (central panel). Clustering the simulated cells identifies ACs (coloured according to the cell state they correspond to), and the proportion of cells in each AC at each time points were compared to the experimental data.

### Fitting the model to experimental data

We estimated model parameters for each SAG concentration using Approximate Bayesian Computation (ABC) particle filtering ^61–63 15,18,64^. This involves simulating each cell’s trajectory using the stochastic model on a finite time interval, calculating the proportions of simulated cells forming a *simulated AC* at regular intervals, and comparing these with their experimental counterpart (Fig. 7C). Parameters are iteratively sampled until convergence to a posterior parameter distribution (PPD) that optimally reproduces the experimental cell state proportions. Details are provided in Appendix B, Sect. B4 for the general fitting procedure, Sect. B5 for the fitting to the flow cytometry data, and Sect. B7 for the scRNA-seq data.

Fitting the model to the flow cytometry data (FACs) for all SAG levels (Fig. 8A) and the scRNA-seq data for SAG 500nM (Fig. 8B) gave good agreement with the experimental proportions of the ventral progenitors. FP cells could not be clearly distinguished in the flow cytometry data but the model provided an estimate (12%) of FP proportions at D6 for SAG 500nM which is consistent with those observed in the scRNA-seq data (7%) in Fig. 8B. Additional validation using flow cytometry is given in Appendix B Fig. B8E. Importantly, we observed that fitting on a limited set of time points still enabled accurate prediction of cell-state proportions at intermediate time points (Fig. 8C).

**Figure 8.**
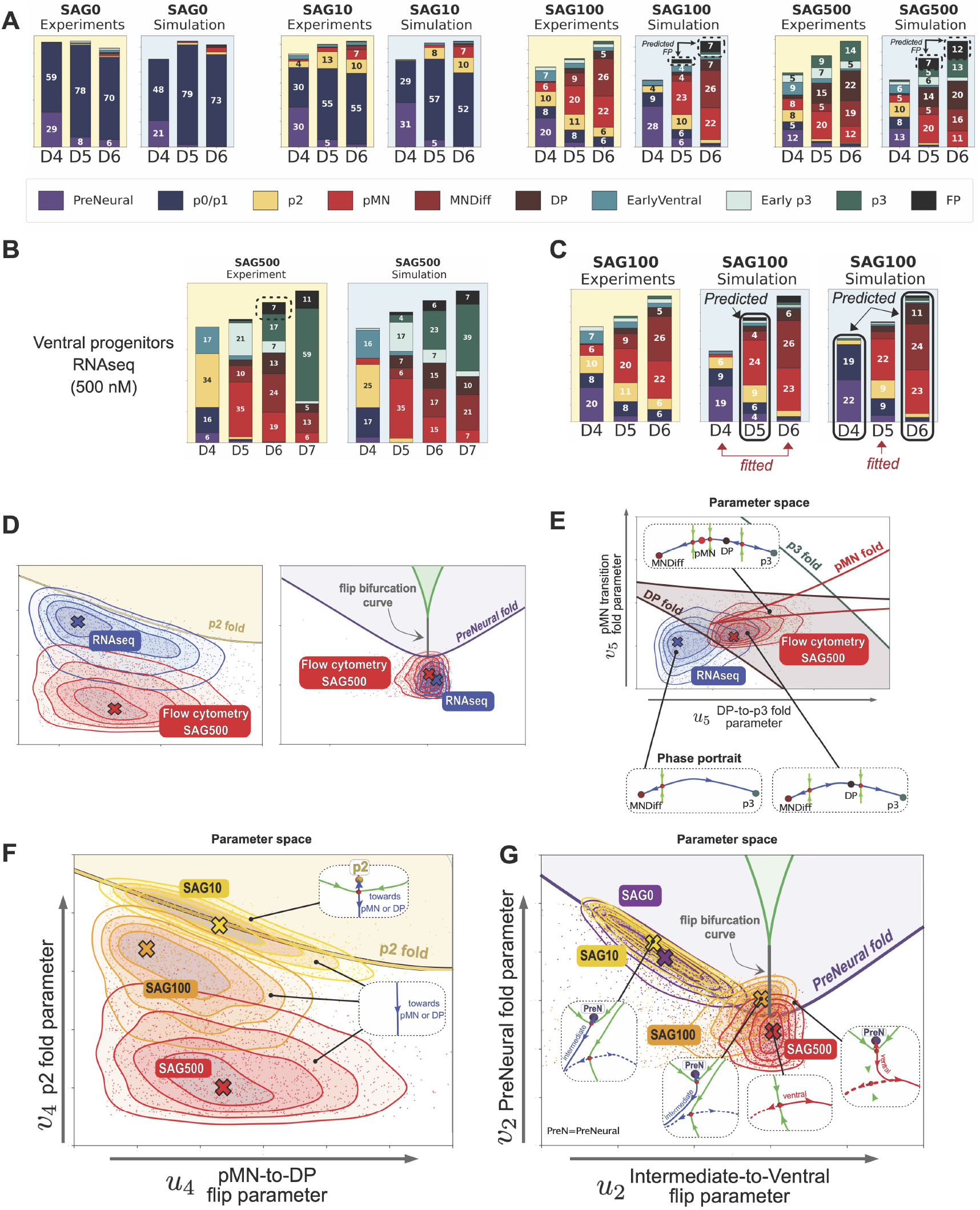
Parameter Estimation and Bifurcation Analysis of the Dynamic Landscape Model. **A**. Experimental proportions (yellow background) compared to the simulated proportions (blue background) after parameter fitting for each SAG concentration using the flow cytometry data. The experimental proportions were obtained by averaging the proportions for 3 individual experimental series for 10nM SAG, 100nM SAG and for 5 experimental series for 0nM and 500nM SAG (Appendix A Sect. A3). The predicted FP proportions (12%) obtained from simulations at 500nM SAG are consistent with those (7%) observed in the scRNA-seq at the same SAG concentration (see B below). **B**. Similar to A for the RNA-seq data at 500nM SAG. **C**. (Left) Experimental proportions for flow cytometry 100nM SAG. (Middle) Fitting the model to only the D4 and D6 experimental proportions accurately predicts the intermediate D5 proportions. (Right) Fitting the model to only the D5 experimental proportions predicts the proportions at D4 an D6. **D.-G**. Dots represent parameters accepted by the ABC particle filter. Contour lines are used to estimate the PPDs projected on the corresponding sub-landscape parameter space (blue: 500nM SAG RNA-seq, red: 500nM SAG flow cytometry, orange: 100nM SAG flow cytometry, yellow: 10nM SAG flow cytometry, purple: 0nM flow cytometry). The other curves show the bifurcation set where fold and flip bifurcations occur for the different sub-landscapes. Panels (E.-G.) contain the sub-landscape phase portraits that arise for parameters chosen within each region of bifurcation set. **D**. (Left) p2 sub-landscape bifurcation set. (Right) PreNeural sub-landscape bifurcation set. The flow cytometry and RNA-seq PPDs at 500nM SAG fall in the same regions of parameter space, delimited by fold and flip curves. The associated dynamics and decisions are thus qualitatively identical. **E**. Intermediate sub-landscape bifurcation set with the flow cytometry and RNA-seq PPDs at 500nM SAG. The shaded region indicates the set of parameters for which the DP attractor is present. As the blue PPD overlaps the DP fold bifurcations curve, we observe heterogeneity: some cells remain trapped in the DP attractor whereas others rapidly transition to p3. The flow cytometry PPD (red) remains largely within the shaded region, where the DP attractor is present, although the fraction of cells with parameters close to the DP fold curve still transition from DP to p3 due to stochastic fluctuations. **F**. Bifurcation set of the sub-landscape controlling the p2 transition. The yellow fold curve represents the parameter values where the p2 attractor bifurcates. At 10nM SAG, the PPD lies on both sides of the p2 fold curve, suggesting that the p2 attractor is present, but shallow, for half of the cells and bifurcated for the other half in which case cells transition to a downstream attractor. At higher SAG concentrations (100nM, 500nM), the PPDs lie outside the shaded region suggesting that the p2 attractor is bifurcated for all cells. The 0nM PPD (purple) is not shown, as the parameters did not converge due to the absence of p2 cells at this SAG concentration (the parameters are distributed across the entire region). **G**. Bifurcation set of the sub-landscape governing the binary decision from PreNeural to intermediate or ventral fates. Parameters on either side of the gray flip curve yield distinct sub-landscapes: one with an unstable manifold (blue) connecting PreNeural to an intermediate fate (p0/p1, p2, pMN), and another (red) to a ventral fate (EarlyVentral, Early p3, p3, FP). Crossing the purple fold curve causes the PreNeural attractor to bifurcate, with fate determined by the basin of attraction the cell is in. At low SAG concentrations (0–10 nM), PPDs lie left of the flip curve, indicating transitions primarily towards intermediate fates. At higher concentrations (100–500 nM), PPDs are near the fold–flip point, allowing heterogeneous transitions to both intermediate and ventral fates.

#### Comparison between flow cytometry and RNA-seq

Given longstanding questions about the the relationship between expression levels of mRNA and protein measurements, it was unclear whether model parameters fitted to both types of data would agree. We therefore compared the PPDs obtained by fitting the model to the scRNA-seq data with those of the flow cytometry at 500nM SAG by projecting them onto the bifurcation sets for each sub-landscape to reveal how signalling conditions control cell fate. We define two PPDs as qualitatively the same if they occupy the same region of the bifurcation set delimited by fold and flip curves. For most sub-landscapes, the SAG 500nM PPDs were qualitatively the same (Fig. 8D), This agreement additionally validates the robustness of the landscape structure across measurement modalities.

#### Model predicts two routes to p3

There was, however, one notable difference. The scRNA-seq PPD crossed the DP fold curve while the cytometry PPD did not (Fig. 8E), forcing DP cells to rapidly transition to p3. We hypothesised that this discrepancy reflected the substantially increased number of p3 cells in the scRNA-seq data on D7 Fig. 8B, as these include the V3 neurons that become abundant at D7. In comparison, the flow cytometry data extended only to D6 and the numbers of p3 cells were lower and did not include V3 neurons as only Sox2-expressing cells were selected. Simulations revealed that the substantial p3 population at late timepoints cannot be achieved solely through the PreNeural-to-p3 pathway via the ventral sub-landscape. This suggested that the PPD crossing the DP fold curve (Fig. 8E) reflects an alternative route: direct transition from DP to p3. The model predicts that cells can reach the p3 fate through two distinct developmental pathways, a prediction we validate below.

#### Bifurcation sets determine how molecular heterogeneity translates into decision heterogeneity

Bifurcation sets fundamentally shape how molecular heterogeneity affects cell fate outcomes. When noise levels are low, two cells, whose landscape parameters are in the same region of the bifurcation set, will make the same decisions with high probability. Conversely, nearly identical cells can make reliably different fate choices if a bifurcation curve divides them into distinct regions. This implies that heterogeneity in differentiation outcomes among molecularly similar cells arises from a cell population straddling bifurcation boundaries rather than from intrinsic noise (Fig. 3E). Understanding bifurcation set structure and how estimated parameters map onto it is therefore essential for interpreting decision heterogeneity in single-cell data.

We analysed the four PPDs obtained by fitting the model to flow cytometry data at constant SAG concentrations.

##### Trapping by p2

The bifurcation set controlling the direct transition from p2 contains a fold bifurcation curve (Fig. 8F). Crossing this curve destabilises the p2 attractor, forcing a cell to transition to pMN or DP. At 10nM SAG, parameters fall on both sides of the fold curve, creating heterogeneity where some cells remain in p2 while others transition. This reproduces the bistability in the p2 and the pMN cells population observed at D6 in Fig. 4H. At higher concentrations (100, 500nM), the PPDs lie below the fold curve, indicating that the p2 attractor has bifurcated for all cells and driving uniform differentiation to pMN. Even at low SAG concentration (10nM), this in vitro protocol fails to stabilise the p2 state. The model predicts that producing a persistent population of p2 cells is not possible by modulating Shh signalling alone.

##### The fold-flip point controlling the choice between the intermediate and ventral routes

In the PreNeural sub-landscape bifurcation set, a flip and a fold curve intersect at a fold-flip point (Fig. 8G). At low SAG (0, 10nM), the PPDs localise to the left of the flip bifurcation curve, biaising cells to intermediate fates. At high SAG (100, 500nM), the PPDs concentrate near the fold-flip point, generating heterogeneous outcomes between the two fates.

Other informative examples showing the relationship between bifurcation and PPDs mappings and validating agreement between RNAseq and flow cytometry data are discussed in Appendix B Sect. B5.1 and Sect. B7.3).

Overall, the fitted model accurately captured the experimental dynamics (Video S1-S3), and successfully reproduced the observed bifurcation behaviours. This framework revealed how individual bifurcations combine to generate complex decision structures and provides a qualitative and quantitative explanation for Shh signalling dependent patterning of ventral neural tissue and makes testable predictions.

### Model predictions of signalling induced landscape changes

We used the fitted model to develop and test various hypotheses. We first asked whether temporal changes in Shh signalling resulted in instantaneous or delayed landscape reconfigurations. We focused on three principal decision points: the transition from PreNeural to either the intermediate or ventral sub-landscapes; the transition from p0/p1 to p2 that leads to pMN; and the connection from DP to p3 responsible for the circular topology of the landscape.

To examine the first two decisions, we recast the model into a simplified version that focused on these transitions (Fig. 9A). This reduced model aggregated several related cell states (combining NMP/PreNeural, EarlyVentral/Early p3/p3, and pMN/MN/DP) while preserving the essential topology of a binary flip landscape followed by a linear transition pathway (Appendix B Sect. B8). In this framework, the ventral sub-landscape consists of a single state p3*, and the intermediate sub-landscape comprises two states: p2 and the combined pMN/MNDiff/DP state, relabelled pMN*. The model accurately reproduced the proportions of cell states observed in various SAG concentrations (Fig. 9C).

**Figure 9.**
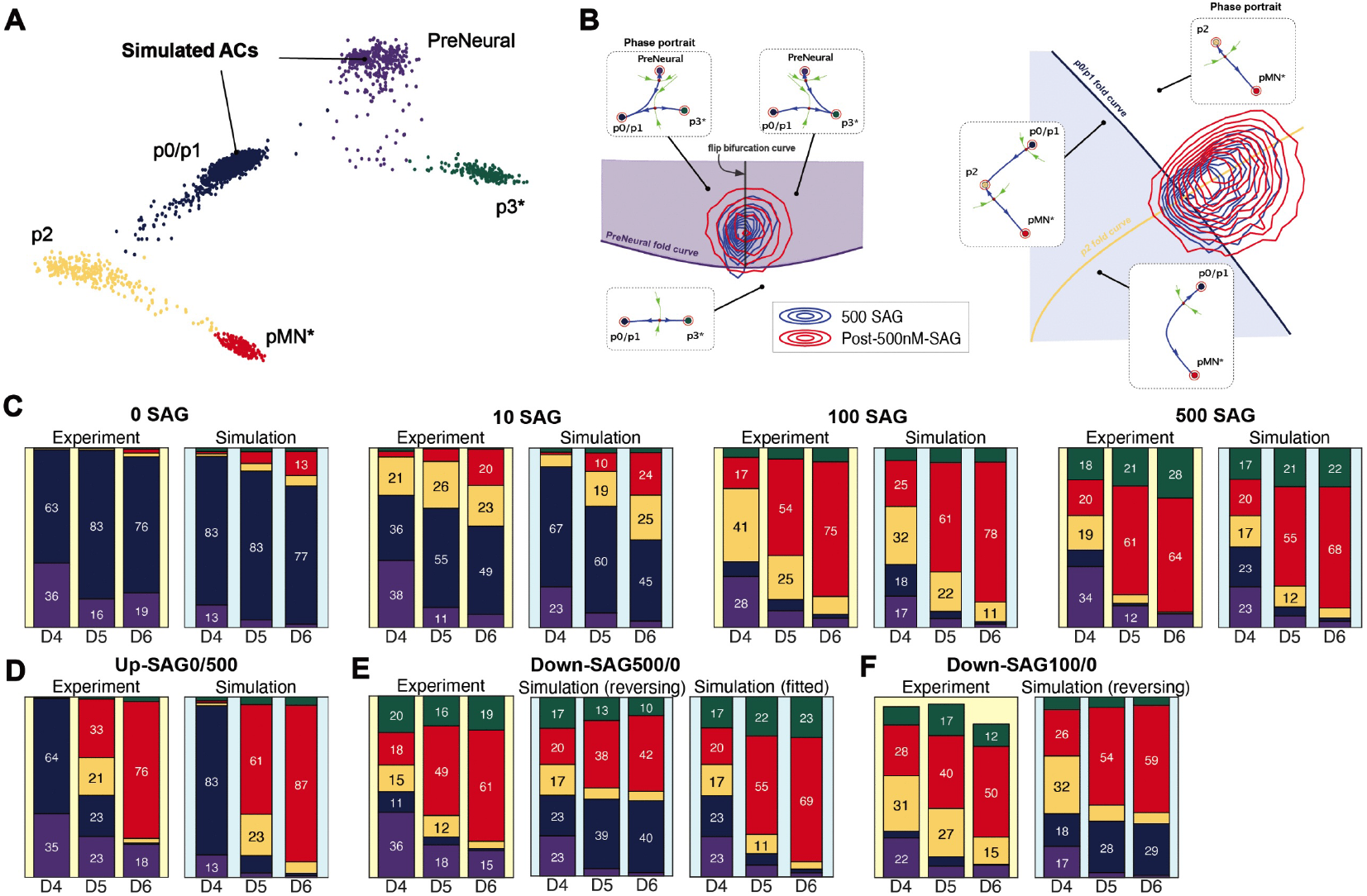
Experimental Validation of Model Predictions By Signalling Perturbations. **A**. Simulated data showing the attractors and transition routes. Dots represent simulated cells retained at 3 time points (D4, D5, D6). These are aggregated into simulated ACs coloured by the cell state to which they correspond. **B**. PPDs from model fitting for the 500nM SAG condition (blue) and the post-high-SAG condition (red) are close to identical, revealing that the two conditions are approximately equivalent. (Left) Projection of the PPDs on the bifurcation set of the binary flip sub-landscape. (Right) Projection of the PPDs on the bifurcation set of the linear sub-landscape. **C**. Comparison of average simulation results (blue background) for the constant SAG conditions with the corresponding experimental results (yellow background). Results are average over 10000 simulated proportions using parameters drawn from the fitted PPDs (Appendix B Sect. B8). **D**. As C but for the Up-SAG0/500 condition. The prediction aligns with the experimental data. It shows that delaying SAG application by 24h commits the cells to differentiate into pMN^*^ rather than p3^*^. **E**. As C but for the Down-SAG500/0 condition, modified as follows: (middle) uses the fitted PPDs (500nM and 0Nm SAG) obtained from C and assume instaneous response to signal change, (right) uses the refitted PPD. The latter shows better agreement with the experimental results demonstrating a lack of response to the change of signal. **F**. As for the first two panels of E for the Down-SAG100/0 condition where the fitted PPDs (100nM and 0nM SAG) obtained from C are used. We observe the same phenomenon as in E (middle panel), despite lower levels of the signal.

**Figure 10.**
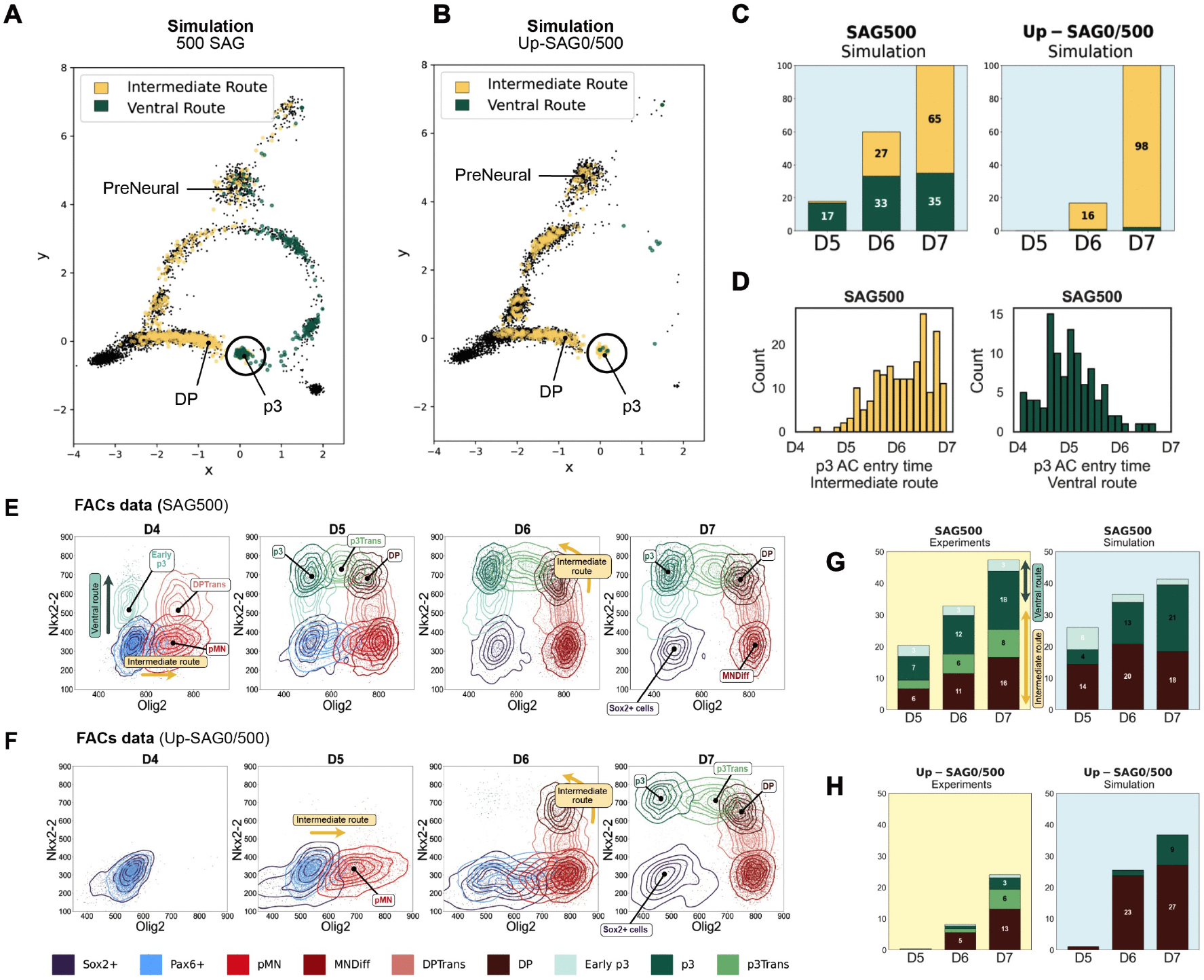
Validation of the DP-to-p3 Transition. **A**. Simulated cell trajectories using the PPD from model fitting (500nM SAG condition). Dots represent simulated cells retained at 4 time points (D4-D7). Cells reaching the p3 AC by D7 are coloured based on their transition history: yellow for cells that follow the intermediate route and green for those that follow the ventral route. Black dots represent cells that adopted alternative end fates (such as MNDiff or FP) or have not yet reached p3 by D7. **B**. Similar to A, but simulating a delayed pulse of SAG by switching the parameters from 0 nM to 500nM SAG after D4. Substantially fewer cells follow the ventral route as most cells had already taken the intermediate route by D4 when the signal changed. **C**. Proportions of simulated cells in the p3 AC for each day (D5 to D7) computed with respect to the total number of p3 cells at D7 and coloured by the route they originated from. (Left) 500nM SAG. (Right) Up-SAG0/500nM SAG condition. The majority of p3 cells under this condition originate from the intermediate route. **D**. Histograms showing cell counts (y-axis) by the time point (x-axis) at which simulated cells first reach the p3 AC, with colours indicating their route of origin: intermediate (yellow) or ventral route (green). The distributions reflect timing heterogeneity: cells that reach p3 via the intermediate route tend to arrive at late time points (D6-D7). By contrast, cells following the ventral route tend to reach p3 early (D4-D5) while few new cells arrive via this route after D6. **E.-F**. 2D Gene plots (Olig2-Nkx2.2 shown) of the clusters identified in the flow cytometry dataset at each time point. The dots represent individual cells, and contour lines represent the density of cells (plotted only when the cluster contains more than 150 cells). Without Nkx6.1 p0/p1 cannot be distinguished from p2 cells and both are combined into Pax6^+^ cells. PreNeural, EarlyVentral and FP are combined into Sox2^+^ cells. **E**. 500nM SAG: as early as D4, a group of cells follow the ventral route by up-regulating Nkx2.2 and reach p3 by D5. **F**. Up-SAG0/500nM SAG condition: with no SAG for 24h, cells are forced to take the intermediate route and most of them had up-regulated Pax6 by D4. When SAG is applied, they can only reach p3 by transiting through pMN (D5), then DP (D6), followed by a direct DP-to-p3 transition (D7). **G**. (Left) 500nM SAG: experimental proportions of the DP AC and p3Trans (intermediate route) and the Early p3 and p3 ACs (ventral route) at late time points (D5-D7). (Right) 500nM SAG: simulated proportions of the DP, Early p3 and p3 ACs at late time points (D5-D7). The p3 proportions can be further split into two type of cells depending on which route they followed. **H**. Same as G for the Up-SAG0/500nM condition. In the simulation (right), most p3 cells at D7 originate from the intermediate route as shown in B which is consistent with the experiment F.

#### Instantaneous cells response to signal increase

We used the model to simulate the effect of timed pulses of Shh signalling on neural progenitor proportions. First, we simulated the effect of 24h with no SAG followed by 48h with 500nM SAG. Under the hypothesis of instantaneous landscape responses, the increase in SAG would trigger immediate bifurcations, destabilising dorsal states (p0-p2). This predicted that, at D4, after 24h with 0nM SAG, cells would adopt a p0/p1 state, at which point an abrupt increase to 500nM SAG would destabilise this state, driving the cells to transition further into the intermediate sub-landscape and not into the usual ventral route to p3. Since the p2 state is absent at 500nM SAG, cells would rapidly adopt a pMN* identity. Importantly, because most cells are predicted to have left the PreNeural attractor and committed to an intermediate fate by D4, few PreNeural cells would be left to transition to the ventral sub-landscape. Consequently, under this hypothesis, the model predicts substantially fewer p3* cells and a purer population of pMN* at D5-D6 compared to constant 500nM SAG exposure.

We experimentally tested this hypothesis by culturing cells in the absence of SAG for 24h, from D3 to D4, and then adding 500nM SAG for the subsequent 48h (D4-D6). We analysed the proportions of cell types at D4, D5 and D6 using flow cytometry. The experimental data were in good agreement with the simulations, with the majority of cells adopting a pMN* identity at D6 (Fig. 9D). These results indicated that the response to raising levels of SAG resulted in fast predictable changes to the landscape.

#### Cells are unresponsive to signal withdrawal

We next simulated the opposite pulsing regime - 500nM SAG for 24h followed by 0nM SAG for 48h. Assuming cells respond immediately to the removal of the signal, triggering instantaneous bifurcations in the landscape, the model predicts the reestablishment of the p0/p1 state and cells that had not reached the p2 state by D4 would remain trapped in p0/p1. Consequently, we would expect a higher proportion of p0/p1 cells to persist at D6 compared to continuous exposure to 500nM SAG.

Experimental results showed a markedly different outcome from this prediction: pMN* cells dominated at D5 and D6 and p0/p1 cells were largely absent as in the constant exposure case (Fig. 9E). The almost complete absence of p0/p1 indicated an irreversible commitment to pMN* fates and that the cells did not respond to an instananeous signal withdrawal. This is consistent with previous experimental observations of bistability and hysteresis in neural progenitor specification ^65^.

The similarity in cell states proportions between constant 500nM exposure and SAG removal after 24h rules out the assumption of an instantaneous cellular response to signal withdrawal. To distinguish whether the response to withdrawing SAG is delayed or entirely absent, we fitted the model directly to the experimental data (Fig. 9E) and compared the resulting PPD ‘post-500nM-SAG’ with that from continuous 500nM SAG exposure. The PPDs were identical (Fig. 9B), indicating that the landscape remains unchanged after SAG withdrawal and the dorsal states (p0-p2) are not reestablished. Similar results from 100 nM to 0 nM SAG transitions demonstrate that 100 nM SAG is also sufficient for irreversible p0/p1 destabilisation (Fig. 9F). Further analysis, including an experiment using a shorter 12-hour pulse of 500 nM SAG,confirmed that the sustained landscape effect persists at full intensity throughout the post-withdrawal period (Appendix B Sect. B6).

#### Experimental validation of convergent topology

Finally, as explained above, the global model predicts that, at 500nM SAG, a substantial proportion of cells reach p3 via the intermediate DP route, arriving later than cells taking the direct ventral pathway. Consistent with this prediction, scRNA-seq data showed concurrent DP depletion and p3 accumulation between D6 and D7 (Fig. 8B).

To test this prediction, we simulated two signalling regimes: constant 500nM SAG versus 24 hours at 0nM followed by 72 hours at 500nM SAG (Fig. 10A-B). We tracked which route cells followed to reach p3. Under the second signalling regime, the vast majority of p3 cells took the intermediate DP-to-p3 route rather than the ventral route via EarlyVentral and Early p3 (Fig. 10C). Entry time analysis revealed distinct kinetics for the two pathways. Cells reached p3 earlier via the ventral route, whereas the intermediate route resulted in delayed arrival due to retention in shallow intermediate states (Fig. 10D).

To test these predictions, we performed experiments extending the time course to D7 (Fig. 10E-F) and analysed the effect of adding 500nM SAG at D4 after 24h without SAG. Under these conditions, the model predicts that PreNeural cells are forced to take the intermediate route as almost all cells initially transition to p0/p1. Subsequently, elevating SAG levels to 500nM SAG destabilises the p0/p1 and p2 attractors, making p3 accessible via the DP saddle. Flow cytometry data at D7 revealed that cells had transitioned from DP to p3, characterised by decreased Olig2 expression while maintaining Nkx2.2 (Fig. 10F). This is particularly clear at D7. In conditions in which SAG exposure is delayed by 24h, p3 cells emerge despite the near absence of Early p3 at earlier time points, ruling out the possibility of a ventral route.

We compared the simulated proportions of DP, Early p3 and p3 ACs for both experimental conditions (Fig. 10G-H), using the PPDs previously obtained for 0nM and 500nM (this experiment was not included in the fitting of the model). The model captured the increased proportion of DP cells and the passage from DP to p3 via the intermediate route (Fig. 10H) (Video S3-S4).

Together, these results further validate the model and demonstrate the complex relationship between signal dynamics and cell fate allocation. The analysis indicates that transient high-level Shh signalling induces irreversible commitment to ventral fates, while also providing support for the dual pathways to p3 identity. These findings establish how cells integrate and retain signalling information to make robust fate decisions, with the mathematical model providing a framework for understanding the underlying dynamical principles.

### Testing DLA on in vitro human data

To evaluate the generality of the landscape model, we investigated its applicability to an in vitro model of human trunk development^31^. This system generates multiple embryonic tissues, including neural progenitors, mesodermal cells, and notochord cells, which are a source of endogenous Shh signalling to the other tissues. The presence of the Shh producing notochord in these organoids results in the generation of ventral neural tube progenitors (p2, pMN, p3) and floor plate (FP). We analysed two experimental conditions - an *18h-delay* and *24h-delay* in TGF*β* addition - which produced different proportions of notochord cells and consequently varying levels of Shh signalling ^31^. The data comprised scRNAseq collected at 3 time points D3, D5, D7.

Using the clustering method described above (Fig. 11A-B), we identified the expected cell populations: NMPs, mesoderm, ventral neural progenitors, and notochord (Fig. 11C-G). When examining neural progenitors (SOX2^+^, TBXT^−^), we discovered remarkably similar ACs to the mouse dataset. Moreover, an LDA projection revealed the ACs were arranged in a similar pattern to the previous data: the PreNeural state leading to intermediate and ventral sub-landscapes and comparable transitions between ACs including a circular topology in which the two routes converge to the p3 AC (Fig. 11D-E).

**Figure 11.**
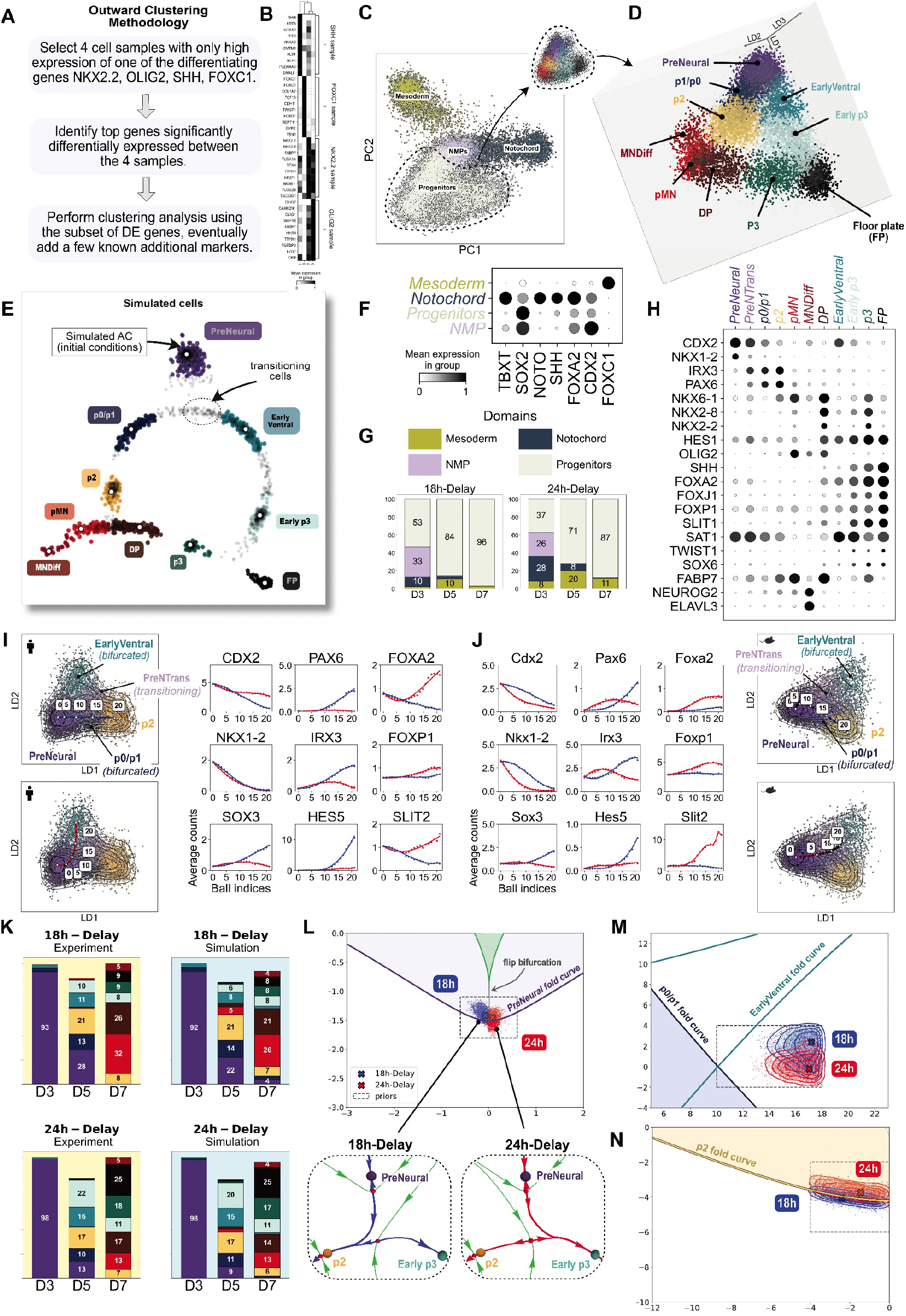
Analysis of Human Neural Progenitor Development Validates the Circular Topology of the Decision Landscape. **A**. Summary of the outward clustering methodology for the human organoid data. **B**. The most significantly differentially expressed genes found using the 4 samples defined in A. **C**. The four cell type domains including neural progenitors were extracted, annotated and separately projected using LDA from the effective gene space defined by the gene modules in B. **D**. 3D LDA projection of the neural progenitor ACs for all time points and experimental conditions. This reveals the circular topology of the transition pathways. **E**. Simulated neural progenitor ACs produced by the landscape model of this system after fitting in a similar fashion to that described above. Dots represent simulated cells retained at 3 time points corresponding to D3, D5, D7 using the parameters from the fitted PPDs for the 18h-Delay condition. Gray dots represent transitioning cells that were not assigned to any simulated AC. **F**. Dot plot indicating the expression of markers used to identify the tissue domains. The intensity of shading indicates the mean expression of each gene in each domain. **G**. Proportions of cells in each domain of C and F by time point in the two experimental conditions. **H**. Dot plot indicating the expression of markers used to identify the ACs and transitioning clusters of the Neural Progenitors domain in D. **I.-J**. The PreNeural sub-landscape extracted from the human dataset (I) and the mouse datasets (J) and transition analysis for marker genes that characterise the binary decision as cells exit PreNeural. Unstable manifolds defining routes between pairs of ACs where computed in effective gene space defined by the genes in B and projected onto 2D LDA space. Marker genes were analysed along both routes. Exit from PreNeural is marked by down-regulation of Cdx2 and Nkx1.2. A branching event occurs (around index 10), followed by up-regulation of markers characteristic of ventral states along the red route (leading to EarlyVentral), and markers such as Pax6 characteristic of the intermediate states p0/p1, and p2 along the blue route. **K**. (Top) Comparison of the experimental and simulated proportions after parameter fitting for the 18h-Delay condition. (Bottom) Similar for the 24h-Delay condition. This shows an increase of ventral states such as p3 and FP. **L**. (Top) Bifurcation set of the PreNeural sub-landscape shows the qualitative difference between the 18h-and the 24h-Delay conditions. The 18h-Delay PPD (blue) appears on the left of the flip curve and above the PreNeural fold curve. Under this condition, the PreNeural attractor is shallow for most cells and is connected to an intermediate fate. By contrast, the 24h-Delay PPD (red) lies to the right of the flip curve, favouring cell trajectories from PreNeural to a ventral fate and promoting the generation of more p3 and FP cells. (Bottom) Typical configurations of attractors, saddles and unstable manifolds for the cell landscape models for the PreNeural sub-landscape for parameters in each of the PPDs. In the 18h-Delay, cells are most likely to transition via the intermediate route towards p2 and to Early p3 via the ventral route in the 24h-Delay. As indicated by the PPDs in M, the attractors p0/p1 and EarlyVentral are bifurcated for all cells in both experimental conditions. Hence the connections from PreNeural lead directly to p2 (blue unstable manifold) or to Early p3 (red unstable manifold). **M**. Bifurcation set of the sub-landscape controlling the p0/p1 and EarlyVentral fold bifurcations. Under both conditions, the PPDs are in the region where the p0/p1 and the EarlyVentral attractors are bifurcated for all cells. **N**. Bifurcation set of the p2 transition shows that, under both conditions, the p2 AC is shallow as most parameters cluster near (and above) the p2 fold curve in the shaded region.

#### Conservation of the decision landscape topology

Despite differences in developmental timing, gene expression patterns characterising the ACs (Fig. 11H) and individual analysis of each sub-landscape showed striking conservation between mouse and human systems (Fig. 11I-J for the PreNeural sub-landscape and Appendix A Fig. A17-A16 for all the other sub-landscapes). We therefore fitted our landscape model to the two datasets *18h-delay* and *24h-delay* focusing on the neural progenitor landscape. We anticipated that, since the *18h-delay* condition contains less notochord and therefore less Shh signalling than the *24h-delay* condition, the *18h-delay* dataset would behave as if it had been exposed to lower SAG concentrations than the *24h-delay* dataset. Cells in the D3 PreNeural AC were used as initial conditions and comparisons were made to the proportions of cells in each simulated AC with the experimental cell states proportions. The model successfully reproduced the experimental data (Fig. 11K). Details of the fitting and the simulation can be found in Appendix B Sect. B9.

#### Using bifurcations to quantify the effect of notochord-derived Shh on the decision landscape

Analysis of the bifurcation sets revealed key differences between the *18h-delay* and *24h-delay* conditions. The 24h-delay generated more notochordal cells (Fig. 11G), resulting in higher proportions of floor plate and p3 progenitors and fewer pMN cells by D7 than the 18h-delay. Importantly, the bifurcation analysis showed that these differences primarily affected the PreNeural flip bifurcation (Fig. 11L), while other bifurcations remained unchanged between conditions (Fig. 11M-N). In both experimental conditions, the p0/p1 attractors were unstable across all cell landscapes, making the p0/p1 AC transient - mirroring the behaviour observed at high SAG concentrations in the mouse experiments (Fig. 11M). Most p2 attractors remained shallow, with the corresponding PPDs clustering just within the p2 bifurcation fold curve (Fig. 11N). This suggests that the increased number of notochordal cells in the 24-delay condition, and consequently the higher level of Shh signalling, primarily influenced the first neural progenitor cell fate decision, directing more cells towards the ventral sub-landscape, providing quantitative insight into how notochord-derived Shh signalling affects ventral patterning.

These findings demonstrate that the landscape model successfully captures neural patterning dynamics across species while revealing features of human development. The analysis identified how cell state transitions are affected by notochord-derived Shh: while the PreNeural flip bifurcation showed high sensitivity, other transitions displayed qualitative robustness to signalling variations. This raises important questions about how dorsal fates are established in the presence of strong ventralising signals from the notochord, suggesting the necessity of additional regulatory mechanisms. Future studies will need to explore how combinations of signals shape the early neural tube landscape.

## DISCUSSION

In this study, we established Dynamic Landscape Analysis (DLA), a systematic framework for constructing quantitative and predictive landscape models from single-cell data. By applying this approach to a model of neural tube development, we uncovered an extensive landscape with a rich bifurcation structure and an unexpected circular topology, where initially divergent lineages converge through multiple routes.

A key component of this approach was the identification and characterisation of attractor clusters (ACs). This follows a semi-supervised strategy, starting with known marker genes and iterative expansion to identify genes associated with the ACs and the transitions between them. This approach identified established neural progenitor states and less well-understood states, such as a double-positive population expressing both Olig2 and Nkx2-9, along with Hes1^52^. The method is characterised by greater control of dimensionality, motivated by the need to avoid noisy or irrelevant dimensions in the data and the extensive experimental work suggesting that cell fate decisions are controlled by a relatively small number of genes ^1^. While current techniques for analysing scRNA-seq data often start with highly variable genes (typically numbering in the thousands), the ratio of relevant genes to all features in such samples is very small. As the fraction of relevant features decreases, existing clustering and dimensionality reduction techniques often fail to discover the identity of relevant features because correlations between samples become dominated by noise and fluctuating irrelevant genes ^66^. Moreover, since understanding transition mechanisms and branching points is a key priority, optimal observability of transition routes is crucial and this requires the ability to compute cell-fate differentiation paths in the gene expression space as opposed to tracing them on a 2D or 3D visualisation. The relatively low dimensionality of our gene modules facilitates this.

A second important feature of DLA is the use of dimension reduction methods that preserve the geometry of the local landscape and the transitions. Our analysis of the bifurcation structures depends upon the dimension reduction projection being differentiable. Non-continuous or non-differentiable methods would eliminate critical geometric features, particularly unstable manifold smoothness. While neural networks can be used to modify t-SNE ^67^ and UMAP ^68^ to create smooth embeddings ^69,70^, LDA provides superior results for analysing related attractors because it maximises separation through a controlled, understandable linear embedding.

Finally, DLA was designed so that it can handle multiple datasets across different time points and morphogen levels without data integration, enable quantitative comparisons between such datasets, and accommodate multiple data modalities, including scRNA-seq and flow cytometry.

The availability of a fitted stochastic model that reproduces experimental data allows a better understanding of analytical methods for single cell data. For example, there has been much interest in trajectory inference methods which aim to infer a 1-dimensional graph-like structure underlying the dynamic process from which the cells are sampled ^71,72^. By projecting the cell states to this structure, their properties can be compared over pseudo-time, an inferred unit of progress along the graph-like structure. In this approach, the cell state branches correspond to nodes of the graph with one inward edge and two outward edges. Flip bifurcations have a similar branching topology, but cell states traverse it with radically different dynamics to that given by pseudo-time.

Our theory predicts that at the branching point there is a head attractor that cells pass through. Experimental results and models validate this. Cells can either remain in a strong attractor until a signal triggers a bifurcation, or linger near a shallow attractor for an extended period before escaping through stochastic fluctuations (Fig. 7B). Consequently, a global pseudo-time cannot accurately represent a cell’s passage through such an attractor. Simulations demonstrate this limitation, showing cells arriving at p3 via the intermediate route are approximately equally distributed across days D5, D6, D7, despite starting from the NMP state simultaneously (Fig. 10D). By contrast, pseudo-time aligns more closely with actual simulation time during transitions between attractors (Fig. 7B). The “stickiness” of attractors explains the difficulty in capturing transitioning cells in experimental data with limited temporal resolution.

Our analysis of the transitions between ACs revealed three transition types: fold bifurcations that drive a direct transition from one state to another (for example in the transition from p2 to pMN); binary flip and binary choice bifurcations that govern the branching decisions between alternative fates.

The bifurcation types provide insight into biological features of cell fate decisions. Fold bifurcations explain how an incremental change in signal strength can trigger a switch in cell identity once a threshold value is crossed. On the other hand, a binary flip bifurcation describes the concurrent production of two downstream states from a progenitor population. In this case, differences in cell state or signals received by a cell determine which of the two possible escape routes a cell chooses. Thus, at a population level, two fates can be generated simultaneously. An important implication of the flip bifurcation is that a cell first leaves the head attractor before determining which downstream fate it adopts. In other words, in these transitions, a cell commits to differentiate before specifying which fate it will choose. This could explain the phenomenon of multi-lineage priming, where cells simultaneously express low levels of genes associated with multiple potential fates before committing to a specific lineage ^56^.

A striking feature of our analysis is the circular topology of the neural progenitor landscape, where two developmental trajectories converge on the p3 identity. Experimental perturbations confirmed that p3 cells can be generated through both paths, with the route followed depending on the timing and level of Shh signalling. Applying our landscape model to human neural organoid development revealed consistency between human and mouse systems, suggesting that this circular topology represents a fundamental feature of vertebrate neural development. This finding reconciles previously contradictory findings of the p3 domain being generated through either a Foxa2^+^/Pax6^−^ route ^30^ or through a Pax6^+^/Olig2^+^ route ^57^. It also suggests greater complexity and redundancy in developmental programmes than can be explained by the conventional view of a strictly hierarchical ‘tree’ of cell type diversification. In development, distinct lineages derived from the same progenitor pool can later reconverge; hematopoiesis is one example ^73^.

While GRN models have been successfully constructed for some systems, there has been a notable lack of progress in others, and current single-cell computational approaches have been disappointing ^74^. GRN construction has been feasible in systems such as oscillators where there is an obvious dynamical phenotype that constrains the possible dynamics of the GRN with easy and informative readouts such as period, amplitude and phase that can be used to explore the effects of perturbations. The generative modelling approach we have developed could play an analogous role for other systems by providing similar tools such as a clear phenotype (the route and timing of a cell passing through the ACs and the proportion of cells doing this) and a precise prescription of the dynamical phenomena that the GRN model must display. Moreover, the simplest gene circuits that produce the local normal forms will likely be highly constrained and this will provide a limited set of gene interaction modules that can be searched for in the data ^17,19^.

Our approach to modelling allows us to readily identify the bifurcation set in the model parameter space. Combining this with the posterior parameter distribution (PPD) found by parameter optimisation provides a potentially powerful tool. It links the PPDs corresponding to different morphogen levels to dynamical behaviour. The relationship between the estimated PPD and the bifurcation curves predicts the behaviour of a heterogeneous group of cells exposed to a particular level of signal. For example, in cell states with stable identities, such as p0/p1 in low SAG concentrations, we find parameters clustering away from critical bifurcation curves so that even a heterogeneous cell population displays homogeneous behaviour. In other cases the PPD overlaps bifurcation curves in a way that suggests a cell population makes a mixture of alternative decisions (e.g. to transition, remain in place, or choose between options in a binary flip landscape). For different morphogen levels all are seen in the transition from the PreNeural state (Fig. 8F). More varied cellular outcomes can occur when the PPDs for different morphogen levels overlap more complex structures in the bifurcation curves, as is the case for the transition from p2 to pMN or DP (Appendix B Fig. B7A Panel 5). A key insight is that distinct fate choices for a group of initially near-identical or “micro-heterogeneous” cells ^75^ can emerge from the deterministic landscape dynamics (bifurcations) rather than solely from noise induced stochastic fluctuations affecting the trajectory of a cell ^76^.

By systematically characterising the decision points and the parameters that control cell fate decisions, we can design differentiation protocols that precisely manipulate bifurcation parameters towards desired cell type proportions. This systematic approach provides a deeper understanding of cellular decision making providing a precise and quantitative approach for designing *in vitro* protocols which, in time, could be applied to organoid models, stem cell therapies and regenerative medicine.

An unexpected insight arising from the analysis of the model is that it challenges the conventional hypoth-esis for neural progenitor patterning in which monotonically ordered morphogen thresholds directly determine all domain boundaries. Instead the analysis suggests that cells progress through a series of binary decisions, first choosing between the intermediate or ventral route and then patterning according to monotonic thresholds or further flips within the separate routes. Critically, the flip bifurcations at PreNeural and Early p3 attractors reveal that the pMN/p3 and p3/FP boundaries in the embryo arise from branching decisions rather than direct transitions between adjacent states. This involvement of flip bifurcations in establishing developmental boundaries introduces a hierarchical component to spatial patterning and expands the repertoire of morphogen-dependent pattern-forming mechanisms. It is consistent with recent evidence of distinct epigenetic regulatory states distinguishing p3 and pMN ^30^. We conjecture that this hierarchical decision-making architecture, coupled with the circular landscape topology, provides greater dynamical robustness compared to simple sequential monotonic thresholds.

Intriguingly, similar hierarchical decision-making dynamics have been observed in the learning process of generalised Hopfield networks, where memories progress through well-defined saddles before splitting into progressively specialised states ^77^. In both neural development and machine learning, these decision landscapes appear to follow reproducible, low-dimensional trajectories despite the high-dimensional nature of the underlying systems. This similarity between biological cell decision making and machine learning suggests that hierarchical binary decisions may represent a fundamental organising principle for robust decision-making across different complex systems.

A challenge for future work is to extend this DLA framework to describe how combinations of different signals shape developmental decision-making by including a model of the way such multidimensional signals change the parameters of the model, as was done for a simpler system ^18^. For these more sophisticated applications, integration of epigenetic data with transcriptional profiles may be required.

Taken together, DLA provides a framework for generating predictive mathematical models of cell fate decisions from single-cell data. With this framework, we have gained insight into how cells navigate fate decisions in neural development. By bridging the gap between high-dimensional single-cell data and mathematical models of cellular behaviour, DLA can advance our understanding of the principles governing cellular differentiation.

## SUPPLEMENTARY MATERIAL

### Supplementary documents

**Appendix A**: contains a detailed description of the data analysis (*AppendixA DataAnalysis*.*pdf*)

**Appendix B**: contains a detailed description of the modelling (*AppendixB Modelling*.*pdf*)

### Supplementary media files

**Video S1**: simulation 10nM SAG

**Video S2**: simulation 100nM SAG

**Video S3**: simulation 500nM SAG

**Video S4**: simulation Up–SAG0/500

### Supplementary table

**Table S1**: provides a summary of the flow cytometry experimental datasets, is included as an attached file *Table flow metadata*.*docx* with this publication.

## RESOURCE AVAILABILITY

## Data availability

The pre-processed flow cytometry data generated in this study have been deposited at https://crick.figshare.com/projects/Neural_Tube_Decision_Landscape/250460 and are publicly available as of the date of preprint publication, prior to peer review. The published sequencing data for mouse embryonic stem cells ^33^ and 3D human notoroids ^31^ analysed in this study can be found respectively in the GEO repositories GSE236520 and GSE255338.

## Acknowledgments

We gratefully acknowledge many very useful discussions with Eric Siggia and Dillon Cislo. We thank the Flow Cytometry Science Technology Platform at the Francis Crick Institute for their assistance in carrying out experiments.

## Author contributions

J.B., M.J.D. and D.A.R. conceived the project. M.J.D. and E.F. designed and performed flow cytometry experiments. R.M. generated and processed scRNA-seq data. M.F. and D.A.R. developed methodology. M.F. developed python software and analysed scRNA-seq data. M.F., M.S. and M.J.D. analysed flow cytometry data. Visualisation M.F. All authors interpreted data and designed experiments. M.F. and D.A.R. developed the mathematical models with contributions from M.S. M.F. constructed the global landscape model and performed related analysis. M.S. performed the pulses analysis and constructed the related model. M.F., J.B. and D.A.R. wrote the manuscript with additional material from M.S. All authors reviewed and edited the manuscript.

## MATERIALS AND METHODS

### Cell lines

Experiments were performed with the mouse embryonic stem cell line HM1^78^ maintained at 37°C with 5% CO 2.

### ES cell culture and differentiation

Mouse ES cells were maintained on a feeder layer of mitotically inactivated mouse embryonic fibroblasts (MEFs, derived and expanded in-house) in ES cell medium (Dulbecco, Modified Eagle Medium (DMEM) Knock Out (Gibco; 10829-018) supplemented with 1% Foetal Bovine Serum (Pan Biotech; P30-2602), Penicillin/Streptomycin (Gibco; 15140122), 2 mM L-Glutamine (Gibco; 25030024), 2 mM Non-essential amino acids (Gibco, Cat No. 11140-035), and 0.1 mM 2-mercaptoethanol (Gibco; 21985-023)) with 1000 U/ml LIF (Chemicon, Int ESG1107)). Media was changed every day and cells were passaged every other day at a density of 500,000 cells per 60mm dish.

For differentiations, cells were washed once with PBS and dissociated using 0.05% Trypsin-EDTA (Gibco; 25300054) for 4 min at 37°C and resuspended in 10 ml ES media. Cells were plated in 10 cm plates coated with 0.1% gelatin to remove feeder cells (‘panning’). Cells were incubated for 15 min at 37°C to allow feeder cells to attach. Without disturbing any attached cells, the cells in suspension were transferred to a second gelatinised 10 cm for another 15 min. The process was repeated a third time.

Differentiations were carried out as previously described ^30^, by plating ‘panned’ mES cells resuspended in N2B27 media (Advanced DMEM - F12 (Gibco, Cat. No. 21331-020) and Neurobasal medium (Gibco, Cat. No. A35829-01) (1:1), supplemented with 1xN2 (Gibco Cat no. 17502001), 1xB27 (Gibco Cat no. 17504001), 2 mM L-glutamine (Gibco, Cat No. 25030024), 40 *µ*g/ml BSA (Sigma-Aldrich, Cat No. A7979-50ML), and 0.1 mM 2-mercaptoethanol) at a density of 20,000 cells in 1.5 ml of media onto 6-well plates (Corning, Cat. No. 353046) precoated in Matrigel (Corning, Cat. No. 356231) diluted 1/100 in Advanced DMEM - F12. The media was supplemented on the different days as follows: Day 0 to Day 2, 10ng/ml bFGF (R&D, Cat. No. 100-18B) and 5 *µ*M LGK (Cayman Chemical Company, Cat. No. 1.800.364.9897); Day 2 to Day 3 for 20 h, 10 ng/ml bFGF, 5 *µ*M CHIR99021 (Axon Medchem, Cat. No. 1386), 10 *µ*M SB-431542 (Tocris, Cat. No. S0400), and 2*µ*M DMH1 (Adooq Bioscience, Cat. No. A12820); from Day 3 onwards, 100 nM RA (Sigma, Cat. No. R2625) and the indicated concentrations and timings of Smoothened Agonist SAG (Merck, Cat. No. 566660-5mg).

### Intracellular flow cytometry

Sample collection: 1*µ*l/ml of LIVE/DEAD Fixable Dead Cell Stain Near-IR fluorescent reactive dye (ThermoScientific, Cat. No. L34976) was added to cells in culture and incubated at 37°C for 30 mins. Cells were then washed twice with PBS (Gibco, Cat. No. 14190-094), and dissociated using 0.5ml Accutase (Gibco, Cat. No. 00-4555-56) per well of a 6 well plate incubated 5 min at 37°C. Cell were collected, centrifuged at 400g for 4 min and resuspended in 100 *µ*l of 4% paraformaldehyde (PFA) (ThermoScientific, Cat. No. 28908). PFA fixation was carried out for 10 min at room temperature. Cells were washed in PBS and resuspended in 500*µ*l PBS + 0.5% BSA.

#### Staining

1 million cells were stained for flow cytometry analysis. Pellets were resuspended in 0.1 ml of PBS supplemented with 0.5% BSA and 0.1% Triton-X100 (VWR Chemicals, Cat No. 28817.295) and the appropriate primary or directly-conjugated antibodies for 1.5 h protected from light at room temperature. Secondary antibodies were incubated under the same conditions for 40 min. Cells were washed in PBS supplemented with 0.5% BSA and 0.1% Triton-X100, and re-suspended in 300*µ*l of PBS with 0.5% BSA for analysis on a BD Fortessa analyser (Becton Dickinson).

The antibody panels used were as follows: Primary antibodies were Sox2-V450 (1:100)(BD Biosciences Cat no 561610), Pax6-PerCPCy5.5 (1:100)(BD Biosciences Cat no 562388), Nkx6.1-AlexaFluor647 (1:100)(BD Biosciences Cat no 563338), Goat Olig2 unconjugated (1:400) (R&D Cat no AF2418) and Nkx2.2-PE (1:100) (BD Biosciences Cat no 564730) followed by secondaries donkey anti-goat AlexaFluor488 (1:1000) (Thermo Fisher Scientific cat no A11055).

### Flow cytometry data pre-processing

The flow cytometry data were analysed using FlowJo™v10.8 Software (BD Life Sciences). Cells were selected for downstream analysis if they were negative for LIVE/DEAD Fixable Dead Cell Stain Near-IR (alive), and with uniform FSC, SSC distribution. The cells were also required to be SOX2+ (neural progenitors).

## References

1. Davidson, E. H. (2010). Emerging properties of animal gene regulatory networks. Nature 468, 911–920.

2. Waddington, C. The strategy of the genes. allen (1957).

3. Huang, S., Eichler, G., Bar-Yam, Y., and Ingber, D. E. (2005). Cell fates as high-dimensional attractor states of a complex gene regulatory network. Physical Review Letters 94, 128701.

4. Huang, S., Guo, Y. P., May, G., and Enver, T. (2007). Bifurcation dynamics in lineage-commitment in bipotent progenitor cells. Developmental Biology 305, 695–713. doi:10.1016/j.ydbio.2007.02.036.

5. Huang, S. (2012). The molecular and mathematical basis of waddington’s epigenetic landscape: A framework for post-darwinian biology? Bioessays 34, 149–157.

6. Corson, F., and Siggia, E. D. (2012). Geometry, epistasis, and developmental patterning. methods: model and numerical simulation. PNAS 0, 1–23.

7. Ferrell Jr, J. E. (2012). Bistability, bifurcations, and waddington’s epigenetic landscape. Current Biology 22, R458–R466.

8. Marco, E., Karp, R. L., Guo, G., Robson, P., Hart, H., Trippa, L., and Yuan, G. C. (2014). Bifurcation analysis of single-cell gene expression data reveals epigenetic landscape. Proceedings of the National Academy of Sciences of the United States of America 111, E5643–E5650. doi:10.1073/pnas.1408993111.

9. Bargaje, R., Trachana, K., Shelton, M. N., McGinnis, C. S., Zhou, J. X., Chadick, C., Cook, S., Cavanaugh, C., Huang, S., and Hood, L. (2017). Cell population structure prior to bifurcation predicts efficiency of directed differentiation in human induced pluripotent cells. Proceedings of the National Academy of Sciences of the United States of America 114, 2271–2276. doi:10.1073/pnas.1621412114.

10. Corson, F., and Siggia, E. D. (2017). Gene-free methodology for cell fate dynamics during development. eLife 6, 1–25. doi:10.7554/eLife.30743.

11. Wang, J., Xu, L., Wang, E., and Huang, S. (2010). The potential landscape of genetic circuits imposes the arrow of time in stem cell differentiation. Biophysical Journal 99, 29–39. doi:10.1016/j.bpj.2010.03.058.

12. Huang, S., Li, F., Zhou, J. X., and Qian, H. (2017). Processes on the emergent landscapes of biochemical reaction networks and heterogeneous cell population dynamics: differentiation in living matters. Journal of the Royal Society Interface 14, 20170097.

13. François, P., and Jutras-Dubé, L. (2018). Landscape, bifurcations, geometry for development. Current Opinion in Systems Biology 11, 129–136.

14. Cao, J., Zhou, W., Steemers, F., Trapnell, C., and Shendure, J. (2020). Sci-fate characterizes the dynamics of gene expression in single cells. Nature biotechnology 38, 980–988.

15. Camacho-Aguilar, E., Warmflash, A., and Rand, D. A. (2021). Quantifying cell transitions in c. elegans with data-fitted landscape models. PLoS Computational Biology 17, 1–28. doi:10.1371/journal.pcbi.1009034.

16. Freedman, S. L., Xu, B., Goyal, S., and Mani, M. (2021). Revealing cell-fate bifurcations from transcriptomic trajectories of hematopoiesis.

17. Rand, D. A., Raju, A., Sáez, M., Corson, F., and Siggia, E. D. (2021). Geometry of gene regulatory dynamics. PNAS 118.

18. Sáez, M., Blassberg, R., Camacho-Aguilar, E., Siggia, E. D., Rand, D. A., and Briscoe, J. (2022). Statistically derived geometrical landscapes capture principles of decision-making dynamics during cell fate transitions. Cell Systems.

19. Sáez, M., Briscoe, J., and Rand, D. A. (2022). Dynamical landscapes of cell fate decisions. Interface Focus 12, 20220002.

20. Raju, A., and Siggia, E. D. (2023). A geometrical perspective on development. Development, Growth & Differentiation 65, 245–254. doi:10.1111/dgd.12855.

21. Raju, A., and Siggia, E. D. (2024). A geometrical model of cell fate specification in the mouse blastocyst. Development 151, dev202467. doi:10.1242/dev.202467.

22. Howe, A. E. S., and Mani, M. (2024). Learning geometric models for developmental dynamics. bioRxiv. https://www.biorxiv.org/content/10.1101/2024.09.21.614191v2. doi:10.1101/2024.09.21.614191. Preprint.

23. Zhou, P., Wang, S., Li, T., and Nie, Q. (2021). Dissecting transition cells from single-cell transcriptome data through multiscale stochastic dynamics. Nature communications 12, 5609.

24. Yeo, G. H. T., Saksena, S. D., and Gifford, D. K. (2021). Generative modeling of single-cell time series with prescient enables prediction of cell trajectories with interventions. Nature communications 12, 3222.

25. Cislo, D. J., Delás, M. J., Briscoe, J., and Siggia, E. D. Reconstructing waddington’s landscape from data (2025). arXiv:in preparation.

26. Luecken, M. D., and Theis, F. J. (2019). Current best practices in single-cell rna-seq analysis: a tutorial. Molecular Systems Biology 15, e8746. doi:10.15252/msb.20188746.

27. Heumos, L., Schaar, A. C., Lance, C., Litinetskaya, A., Drost, F., Zappia, L., Lücken, M. D., Strobl, D. C., Henao, J., Curion, F. et al. (2023). Best practices for single-cell analysis across modalities. Nature Reviews Genetics 24, 550–572.

28. Gouti, M., Tsakiridis, A., Wymeersch, F. J., Huang, Y., Kleinjung, J., Wilson, V., and Briscoe, J. (2014). In vitro generation of neuromesodermal progenitors reveals distinct roles for wnt signalling in the specification of spinal cord and paraxial mesoderm identity. PLoS Biology 12, e1001937.

29. Gouti, M., Delile, J., Stamataki, D., Wymeersch, F. J., Huang, Y., Kleinjung, J., Wilson, V., and Briscoe, J. (2017). A gene regulatory network balances neural and mesoderm specification during vertebrate trunk development. Developmental Cell 41, 243–261.

30. Delás, M. J., Kalaitzis, C. M., Fawzi, T., Demuth, M., Zhang, I., Stuart, H. T., Costantini, E., Ivanovitch, K., Tanaka, E. M., and Briscoe, J. (2023). Developmental cell fate choice in neural tube progenitors employs two distinct cisregulatory strategies. Developmental Cell 58, 3– 17.e8.

31. Rito, T., Libby, A. R. G., Demuth, M., Domart, M.-C., Cornwall-Scoones, J., and Briscoe, J. (2025). Timely tgfβ signalling inhibition induces notochord formation from human pluripotent stem cells. Nature 637.

32. Sagner, A., Zhang, I., Watson, T., Lazaro, J., Melchionda, M., and Briscoe, J. (2021). A shared transcriptional code orchestrates temporal patterning of the central nervous system. PLoS Biology 19, e3001450. doi:10.1371/journal.pbio.3001450.

33. Maizels, R. J., Snell, D. M., and Briscoe, J. (2024). Reconstructing developmental trajectories using latent dynamical systems and time-resolved transcriptomics. Cell Systems 15, 411–424.e9.

34. Jessell, J. M. (2000). Neuronal specification in the spinal cord: inductive signals and transcriptional codes. Nature Reviews Genetics 1, 20–29.

35. Frith, T. J., Briscoe, J., and Boezio, G. L. Chapter five - from signalling to form: the coordination of neural tube patterning. In: M., M., ed. Vertebrate Pattern Formation vol. 159 of Current Topics in Developmental Biology (168–231). Academic Press (2024):(168–231).

36. Diez del Corral, R., Breitkreuz, D. N., and Storey, K. G. (2002). Onset of neuronal differentiation is regulated by paraxial mesoderm and requires attenuation of FGF signalling. Development 129, 1681– 1691. doi:10.1242/dev.129.7.1681.

37. Sagner, A., Gaber, Z. B., Delile, J., Kong, J. H., Rousso, D. L., Pearson, C. A., Weicksel, S. E., Melchionda, M., Gharavy, S. N. M., Briscoe, J., and Novitch, B. G. (2018). Olig2 and hes regulatory dynamics during motor neuron differentiation revealed by single cell transcriptomics. PLoS Biology 16, e2003127.

38. Wang, S., Sontag, E. D., and Lauffenburger, D. A. (2023). What cannot be seen correctly in 2d visualizations of single-cell ‘omics data? Cell systems 14, 723–731.

39. Traag, V. A., Waltman, L., and van Eck, N. J. (2019). From Louvain to Leiden: guaranteeing well-connected communities. Scientific Reports 9, 5233.

40. Belhumeur, P. N., Hespanha, J. P., and Kriegman, D. J. (1997). Eigenfaces vs. fisherfaces: Recognition using class specific linear projection. IEEE Transactions on Pattern Analysis and Machine Intelligence 19, 711–720. doi:10.1109/34.598228.

41. Schäfer, M., Kinzel, D., and Winkler, C. (2007). Discontinuous organization and specification of the lateral floor plate in zebrafish. Developmental Biology 301, 117–129. doi:10.1016/j.ydbio.2006.09.018.

42. Mojtahedi, M., Skupin, A., Zhou, J., Castaño, I. G., Leong-Quong, R. Y. Y., Chang, H., Trachana, K., Giuliani, A., and Huang, S. (2016). Cell fate decision as high-dimensional critical state transition. PLOS Biology 14, e2000640. doi:10.1371/journal.pbio.2000640.

43. Wilkinson, D. G., Bhatt, S., and Herrmann, B. G. (1990). Expression pattern of the mouse T gene and its role in mesoderm formation. Nature 343, 657– 659. doi:10.1038/343657a0.

44. Hiemisch, H., Monaghan, A. P., Schütz, G., and Kaestner, K. H. (1998). Expression of the mouse Fkh1/Mf1 and Mfh1 genes in late gestation embryos is restricted to mesoderm derivatives. Mechanisms of Development 73, 129–132. doi:10.1016/s0925-4773(98)00039-2.

45. Schubert, F. R., Fainsod, A., Gruenbaum, Y., and Gruss, P. (1995). Expression of the novel murine homeobox gene Sax-1 in the developing nervous system. Mechanisms of Development 51, 99–114. doi:10.1016/0925-4773(95)00358-8.

46. Takemoto, T., Uchikawa, M., Yoshida, M., Bell, D. M., Lovell-Badge, R., Papaioannou, V. E., and Kondoh, H. (2011). Tbx6-dependent sox2 regulation determines neural or mesodermal fate in axial stem cells. Nature 470, 394–398.

47. Javali, A., Misra, A., Leonavicius, K., Acharyya, D., Vyas, B., and Sambasivan, R. (2017). Coexpression of tbx6 and sox2 identifies a novel transient neuromesoderm progenitor cell state. Development 144, 4522–4529.

48. Novitch, B. G., Chen, A. I., and Jessell, T. M. (2001). Coordinate regulation of motor neuron subtype identity and pan-neuronal properties by the bHLH repressor Olig2. Neuron 31, 773–789. doi:10.1016/s0896-6273(01)00407-x.

49. Briscoe, J., Pierani, A., Jessell, T. M., and Ericson, J. (2000). A homeodomain protein code specifies progenitor cell identity and neuronal fate in the ventral neural tube. Cell 101, 435–445.

50. Echelard, Y., Epstein, D. J., St-Jacques, B., Shen, L., Mohler, J., McMahon, J. A., and McMahon, P. (1993). Sonic hedgehog, a member of a family of putative signaling molecules, is implicated in the regulation of CNS polarity. Cell 75, 1417–1430. doi:10.1016/0092-8674(93)90627-3.

51. Mizuguchi, R., Sugimori, M., H, T., H, K., Nagao, M., Yoshida, S., Nabeshima, Y., Shimamura, K., and Nakafuku, M. (2001). Combinatorial roles of olig2 and neurogenin2 in the coordinated induction of pan-neuronal and subtype-specific properties of motoneurons. Neuron 31, 757–771. doi:10.1016/S0896-6273(01)00413-5.

52. Jang, S., Gumnit, E., and Wichterle, H. (2024). A human-specific progenitor sub-domain extends neurogenesis and increases motor neuron production. Nature Neuroscience 27, 1945–1953. doi:10.1038/s41593-024-01739-8.

53. Pabst, O., Herbrand, H., and Arnold, H.-H. (1998). Nkx2-9 is a novel homeobox transcription factor which demarcates ventral domains in the developing mouse cns. Mechanisms of Development 73, 85–93. doi:10.1016/S0925-4773(98)00035-5.

54. Holz, A., Kollmus, H., Ryge, J., Niederkofler, V., Dias, J., Ericson, J., Stoeckli, E. T., Kiehn, O., and Arnold, H.-H. (2010). The transcription factors nkx2.2 and nkx2.9 play a novel role in floor plate development and commissural axon guidance. Development 137, 4249–4260. doi:10.1242/dev.053819.

55. Toh, K., Saunders, D., Verd, B., and Steventon, B. (2022). Zebrafish neuromesodermal progenitors undergo a critical state transition in vivo. iScience 25, 105216.

56. Hu, M., Krause, D., Greaves, M., Sharkis, S., Dexter, M., Heyworth, C., and Enver, T. (1997). Multilineage gene expression precedes commitment in the hemopoietic system. Genes & Development 11, 774–785. doi:10.1101/gad.11.6.774.

57. Dessaud, E., Yang, L., Hill, K., Cox, B., Ulloa, F., Ribeiro, A., Mynett, A., Novitch, B. G., and Briscoe, J. (2007). Interpretation of the sonic hedgehog morphogen gradient by a temporal adaptation mechanism. Nature 450, 717–720. doi:10.1038/nature06347.

58. Thom, R. (1969). Topological models in biology. Topology 8, 313–335. doi:10.1016/0040-9383(69)90018-4.

59. Thom, R. Structural Stability and Morphogenesis: An Outline of a General Theory of Models. CRC Press (1989).

60. Hirsch, M. W. Differential Topology vol. 33. ‘Springer Science & Business Media’ (2012).

61. Toni, T., Welch, D., Strelkowa, N., Ipsen, A., and Stumpf, M. (2009). Approximate bayesian computation scheme for parameter inference and model selection in dynamical systems. Journal of The Royal Society Interface 6, 187–202.

62. Klinger, E., Rickert, D., and Hasenauer, J. (2018). pyabc: distributed, likelihood-free inference. Bioinformatics 34, 3591–3593.

63. Schälte, Y., Klinger, E., Alamoudi, E., and Hasenauer, J. (2022). pyabc: Efficient and robust easyto-use approximate bayesian computation. Journal of Open Source Software 7, 4304.

64. Camacho-Aguilar, E. New mathematical methods for the study of stem cell differentiation. Ph.D. thesis University of Warwick Warwick, UK (2018). https://wrap.warwick.ac.uk/id/eprint/108668/.

65. Balaskas, N., Azevedo, G. R., Panov, J. K., Marti, E., Briscoe, J., and Anderson, K. R. M. (2012). Gene regulatory logic for reading the sonic hedgehog signaling gradient in the vertebrate neural tube. Cell 148, 273–284. doi:10.1016/j.cell.2011.10.047.

66. Melton, S., and Ramanathan, S. (2021). Discovering a sparse set of pairwise discriminating features in high-dimensional data. Bioinformatics 37, 202–212.

67. van der Maaten, L., and Hinton, G. (2008). Visualizing data using t-sne. Journal of Machine Learning Research 9, 2579–2605.

68. McInnes, L., Healy, J., Saul, N., and berger, L. G. (2018). Umap: Uniform manifold approximation and projection. Journal of Open Source Software 3, 861. https://doi.org/10.21105/joss.00861. doi:10.21105/joss.00861.

69. Van Der Maaten, L. Learning a parametric embedding by preserving local structure. In: Artificial intelligence and statistics. PMLR (2009):(384–391).

70. Sainburg, T., McInnes, L., and Gentner, T. Q. (2021). Parametric umap embeddings for representation and semisupervised learning. Neural Computation 33, 2881–2907.

71. Deconinck, L., Cannoodt, R., Saelens, W., Deplancke, B., and Saeys, Y. (2021). Recent advances in trajectory inference from single-cell omics data. Current Opinion in Systems Biology 27, 100344.

72. Saelens, W., Cannoodt, R., Todorov, H., and Saeys, Y. (2019). A comparison of single-cell trajectory inference methods. Nature biotechnology 37, 547– 554.

73. Adolfsson, J., Månsson, R., Buza-Vidas, N., Hultquist, A., Liuba, K., Jensen, C. T., Bryder, D., Yang, L., Borge, O.-J., Thoren, L. A. M., Anderson, K., Sitnicka, E., Sasaki, Y., Sigvardsson, M., and Jacobsen, S. E. W. (2005). Identification of Flt3^+^ lympho-myeloid stem cells lacking erythro-megakaryocytic potential: a revised road map for adult blood lineage commitment. Cell 121, 295– 306.

74. Badia-i Mompel, P., Casals-Franch, S., Aguilar-Mogas, A., Piulachs, M.-D., Guigó, R., Pérez-Brocal, V., Moya, A., Montañez, R., Pérez-R., I., and Conesa, A. (2022). Comparison and evaluation of methods to infer gene regulatory networks from single-cell rna-seq data. bioRxiv. doi:10.1101/2022.12.20.521206.

75. Kamenev, D., Kameneva, P., and Adameyko, I. (2025). The role of microheterogeneity in cell fate decisions in neural progenitors and neural crest. Current Opinion in Neurobiology 65, 101316. doi:10.1016/j.conb.2025.101316.

76. Balázsi, G., van Oudenaarden, A., and Collins, J. J. (2011). Cellular decision making and biological noise: From microbes to mammals. Cell 144, 910–925. doi:10.1016/j.cell.2011.01.030.

77. Boukacem, N. E., Leary, A., Thériault, R., Gottlieb, F., Mani, M., and François, P. (2024). Waddington landscape for prototype learning in generalized hopfield networks. Physical Review Research 6, 033098. doi:10.1103/PhysRevResearch.6.033098.

78. Magin, T. M., McWhir, J., and Melton, D. W. (). A new mouse embryonic stem cell line with good germ line contribution and gene targeting frequency. 20, 3795–3796. doi:10.1093/nar/20.14.3795.

