## Appendix A for "Dynamic Landscape Analysis of Cell Fate Decisions: Predictive Models of Neural Development From Single-Cell Data"

#### Data Analysis

**Overview:** This appendix provides comprehensive methodological details for the data analysis in the main text. Section A1 describe the "outward clustering" approach for identifying attractor clusters from scRNA-seq data and comprises a detailed analysis of all the found clusters. Section A2 presents the transition analysis methodology for characterising gene expression changes along unstable manifolds. Section A3 details the flow cytometry (FACs) analysis and cluster validation. Section A4 analyses how the decision landscape responds to SAG perturbations. Section A5 validates the landscape topology using human scRNA-seq data to demonstrate cross-species conservation. All gene modules required for reproducibility are provided in Appendix A. The additional supplementary figures are organised to cover mouse scRNA-seq attractor clusters at early and late stages (Fig. A1-A2), transition analysis for key sub-landscapes (Fig. A4-A7), FACs analysis and validation (Fig. A8-A10), SAG perturbation responses (Fig. A12-A14), and human data cross-species validation (Fig. A15-A17).

**Connection to Main Paper:** This analysis complements the scRNA-seq data analysis (**Main Fig. 2-3**), the flow cytometry analysis (**Main Fig. 4**) and the Human data validation (**Main Fig. 11**).

**A Appendix: Gene modules** **29**

### Part I
### Data analysis

#### A1. scRNA-seq analysis

##### A1.1. Outward clustering and gene module selection in scRNA-seq data

We begin with a small set of marker genes and iteratively work from these to find clusters and genes that define and differentiate the found clusters. We reason that, if we knew the ACs and the transition pathways between them, we could identify key genes by finding those that are differentially expressed between these ACs and adding to these the genes with nontrivial variation along the paths between the ACs. Since neither the ACs nor the transition pathways are known a priori, we use a bootstrap approach that uses samples of cells characterised by specific markers to start the process and then when we have candidate ACs use the transition pathways to identify the further transitioning genes.

Our approach starts with a small number $N < 5$ of known marker genes characteristic of cell states or specific transition routes in the system. For each time point $t$ and for each marker gene $g_j$ we extract from the relevant dataset the sample $S_{j,t}$ of all cells that express only that marker gene $g_j$ and no other from the list (or they can have low expression). Empirically we found log-normal gene expression $g_j$ in cells in $S_{j,t}$ to be approximately normal and we remove from $S_{j,t}$ cells that are either in the left or right 5% tails. Here we log-normalised the data to account for cell size[4,19], other types of normalisation can be used to account of the negative binomial nature of raw counts in scRNA-seq data[13].

We proceed by finding the genes that are differentially expressed between the different subsets $S_{j,t}$ of cells using a threshold on the ranking provided by Z-score of the Wilcoxon rank-sum test implemented in Scanpy[34]. We take the union $G$ of these genes over the different time points and manually optimise the size of $G$ by varying the threshold for the level of significance in the differential expression analysis so that we obtain sufficiently many genes in the list (around $d = 100$ genes). The size $d$ of the module is chosen to be large enough to preserve data quality (so that the analysis is not dominated by noise caused by dropouts or irrelevant genes), yet small enough that adding more genes does not affect the global and local PCA projections of the clusters, retaining the topology of the transitions (see below).

We now restrict the data to this set of $d$ genes and call the resulting gene expression space the *effective gene space*.

To facilitate the clustering algorithm, PCA is then used to further reduce the dimension of the effective gene space to a $p$-dimensional space where $p$ is chosen after inspection of the singular value elbow plot to choose the number of principal components to project onto (typically in our examples $p = 7$). Leiden clustering[31] at high resolution is then used to find clusters in this $p$-dimensional space. By high resolution, we mean that the clustering produces more than twice the number of expected cell types as found in[20]. The resulting clusters are then merged based on shared marker gene expression profiles to form larger clusters with a coherent gene expression that differentiates them from the neighbouring clusters. This process relies on the geometry of the data rather than the explicit expression values and consequently is robust to individual marker gene drop-outs. It produces well-distributed clusters that line up along distinct transition routes.

Once AC candidates have been identified we can further expand the module $G$, using the differentially expressed genes between the found ACs in place of the one-marker samples and reiterate the process. For the data discussed below we found that, after optimisation, such iteration was unnecessary as it reproduced the same ACs and the topology of their projection in PCA space was unaffected.

In a later step we will add genes that are not differentially expressed between ACs but which have non-trivial variation along the transition pathways.

##### A1.2. Choice and use of marker genes

For the neural system under consideration[20], we divided the dataset into an *early subset* comprising data for 6 time points between days D3 and D4 (22358 cells) and a *late subset* with

4 time points between days D5 and D8 (24885 cells). The reason for splitting the dataset is that markers such as Nkx2.2 (used to compute genes in $G_{late}$) are not expressed in cells at early time points and are therefore not informative at early stages.

To find the gene module $G_{early}$ (Table A2), associated with the early dataset, we started with the $N = 4$ marker genes, namely TBXT/BRA, Foxc2, Foxa2, Olig2. This choice is chosen because we expected to distinguish the starting NMP population (TBXT/BRA) as well as Mesoderm (Foxc2) and early neural lineages[7,11,12]. As cells were exposed to 500nM SAG, we anticipated that Olig2, a marker of pMN[25] would be the dominant neural progenitor marker. Foxa2 marks the ventral cell lineage producing progenitors such as p3 and FP[7]. We found a similar module of genes by replacing these markers by equivalent ones, e.g Olig2 by Neurog2 or Foxa2 by Sox6 or Foxp1, Foxc2 by Pax3.

To produce the gene module $G_{late}$ (Table A3), associated with the late dataset, we found that it was sufficient to use only $N = 3$ marker genes Olig2, Nkx2.2 and Shh which are themselves markers for the known progenitor states pMN, p3 and FP respectively, expected to be generated in the 500nM SAG experimental condition[20]. Building on the analysis of the early time points, we included a module of differentially expressed genes between the Foxa2 and Olig2 samples. However, this yielded only two extra significantly differentially expressed transcription factors: Foxa2 and Sox6. To cluster the late dataset we used the module of genes obtained as described above without including any extra genes.

For further analysis, the module can be inflated by further differential expression analysis (using the full set of genes) between the found clusters. For example the so-called DP cluster was found to express Gm38103 (Nkx2.9), Hes1 and Rfx4 which distinguished it from other clusters and these markers can be added to $G_{late}$ if not already present.

##### A1.3. Attractor clusters and quality criteria

We then restricted the early and late datasets independently using their respective gene modules $G_{early}$ and $G_{late}$ to perform Leiden clustering in PCA space as above. Leiden clusters were merged based on similarities in gene expression to form larger clusters, that were then classified between ACs and transitioning clusters. All identified clusters are presented in Fig. A1-A2.

To test the quality of these clusters we applied linear projection techniques, specifically linear discriminant analysis[3] (LDA) (see Sect. A1.4 below). Using well-understood linear projections like this has advantages over non-differentiable methods such as t-SNE[32] and UMAP[22]. For example, it gives an explicit linear transformation that can be understood and interpreted in terms of gene expression, it indicates which genes drive cluster separation[1], and unlike t-SNE and UMAP it is deterministic and requires no parameter adjustment (see e.g discussion in[33]). Most importantly, it preserves the geometry of the local landscape and the transitions. For example, for a binary flip decision, it preserves both the triangular structure of the ACs involved and the transition paths given by the unstable manifolds (Fig. A1A-B). A useful property for understanding the separation of clusters is that if two clusters are distinct in the LDA projection (e.g. in terms of Kullback–Leibler divergence, the natural measure of this difference) then they are also distinct in the higher dimensional space.

We evaluated the quality of the clusters using three **criteria**:

1. *Temporal progression:* Neighbouring clusters were extracted and their temporal dynamics analysed in LDA space (Fig. A1D). The persistence of a cluster over a significant time interval and its potential breakdown as its cells transitions towards a neighbouring cluster shows consistency with bifurcation theory that an attractor either bifurcates or is close enough to a saddle to allow cells to pass over due to stochastic fluctuations.

2. *Unimodality in LDA coordinates:* Since the cells in the ACs are genetically homogeneous we expect the raw counts of the characteristic marker genes within an AC to follow a negative binomial distribution, a well-known property of gene expression in scRNA-seq data[18]. In our examples, all clusters satisfy this property regardless of their classification as ACs or transitioning clusters. We assume that the dynamics can be locally approximated within a subspace of gene expression space, with the dimensions corresponding to combinations of genes given by the LDA components (LDs). For an AC we expect that the cluster is well structured in LDA space and we therefore checked that, for each linear discriminant component, the distribution across cells in the AC of the corresponding LDA score was unimodal (Fig. A1E), indicating well-defined gene expression profiles for each AC consistent with the idea that dynamics in the vicinity of attractors are approximately linear[14,17].

3. *Cluster separability:* In LDA space, the ACs, labelled A and B, can be approximated by multivariate Gaussian distributions $\alpha_A, \alpha_B$. To quantify whether these are well separated, we use the explicit formulation of the *Kullback-Leibler (KL) divergence*: for two multivariate Gaussian distributions $\alpha_A, \alpha_B$ with sample means $\mu_A, \mu_B$ and sample covariance matrices $\Sigma_A, \Sigma_B$ in dimension $m$ (which is the dimension of the LDA space), the KL-divergence between $\alpha_A$ and $\alpha_B$ is explicitly given by

$$D_{KL}(\alpha_A, \alpha_B) = 1/2 \left(\mathrm{tr}(\Sigma_B^{-1}\Sigma_A) - m + (\mu_B - \mu_A)^t \Sigma_B^{-1} (\mu_B - \mu_A) + \ln\left(\frac{\det \Sigma_B}{\det \Sigma_A}\right)\right). \quad (1)$$

We then divide this quantity by the differential entropy

$$H(\alpha_A) = \frac{1}{2} \log\left((2\pi e)^m \det(\Sigma_A)\right) \quad (2)$$

(see Table A1).

4. *The critical transition index:* The transition index introduced by Mojtahedi et al.[24] quantifies the fluctuations in gene expression among cells as they approach a transition point (see Sect. A2.3 below).

5. *Local gene-gene correlations:* Distinct cell identities were further validated by analysing the gene-gene correlations within each cluster (ACs and transitioning) by using the raw counts rather than the normalised counts (Fig. A1F). Different correlations for a fixed pair of genes between clusters provide evidence for different regulatory interactions.

Regarding point 3 above, it is usual to normalise the KL divergence $D_{KL}(\alpha_A, \alpha_B)$ by dividing by the entropy $H(\alpha_A)$ to get a scale-invariant measure that expresses the divergence as a fraction of the uncertainty in $\alpha_A$, and therefore gives an answer that can be compared across systems and datasets. Having the ratios $D_{KL}(\alpha_A, \alpha_B)/H(\alpha_A)$ and $D_{KL}(\alpha_B, \alpha_A)/H(\alpha_B)$ of the order of 1 (e.g. $> 0.5$) is regarded as implying that the distributions are identifiable i.e. given a sample there is a high probability of identifying which of the distributions it came from [5,6].

Due to the continuous exposure to a high SAG concentration (500nM), cells occupying ACs such as p0/p1 or p2 were relatively rare. Although small clusters associated with these states were found, they only appeared briefly at D3.8 and D4 (Fig. A1G) and we hypothesised that these ACs were bifurcated with cells transitioning through them without stabilising (an observation that was validated by fitting a model (Appendix B Fig. B9F Panel 3). The behaviour of these cluster was verified below using lower doses of SAG and single cell flow cytometry data.

##### A1.4. Linear discriminant analysis (LDA)

To analyse the transitions and further validate that ACs correspond to stable cell states, we used Linear Discriminant Analysis [3] (LDA) as implemented in the Python package *sklearn.lda* [26]. Unlike Principal Component Analysis, LDA requires the dataset already to be separated into clusters. Projected clusters are separated in an optimal way by maximising the dispersion of cluster means relative to the global mean while minimising the dispersion of the data points within each individual cluster around its mean. This method is most effective if restricted to a small number of $k = 3, 4$ clusters $L = \{C_i\}_{i=1}^k$ that we refer to as a *sub-landscape*. Sub-landscapes are derived from the reduced dataset with columns comprising the $d$ informative genes $g \in G$ of the effective gene space. Each cluster $C_i$ of $L$ contains $q_i$ barcodes and can be either an AC or a transitioning cluster. The set of rows of $L$ corresponding to the barcodes $\mathcal{B}$ has size $q = \sum_{i=1}^k q_i$. LDA maps each barcode $\mathbf{x} \in \mathcal{B}$ in effective gene space $\mathbb{R}^d$ to an element $\mathbf{y}$ in lower dimensional space $\mathbb{R}^m$ with $m < d$ by means of a linear projection $\mathbf{y} = Q\mathbf{x}$. The projection matrix $Q$ is found such that it maximises the ratio

$$J(Q) = \frac{\det(QS_bQ^t)}{\det(QS_wQ^t)} \tag{3}$$

between the within-class scatter $S_w$ and the between-class scatter $S_b$ for the projected data. The matrix

$$S_w = \sum_{i=1}^{k} \sum_{\mathbf{x} \in C_i} (\mathbf{x} - \mathbf{m}_i)(\mathbf{x} - \mathbf{m}_i)^t$$

evaluates the spread of the data in each cluster $C_i$ around their mean $\mathbf{m}_i = \frac{1}{q_i} \sum_{\mathbf{x} \in C_i} \mathbf{x}$, while the matrix

$$S_b = \sum_{i=1}^{k} q_i(\mathbf{m} - \mathbf{m}_i)(\mathbf{m} - \mathbf{m}_i)^t$$

evaluates the spread of the clusters means around the global mean $\mathbf{m} = \frac{1}{q} \sum_{\mathbf{x} \in \mathcal{B}} \mathbf{x}$. Provided the matrix $S_w$ is invertible, the ratio (3) is optimised by taking the $m = k - 1$ largest eigenvalues of $S_w^{-1} S_b$ [9]. The optimal matrix is the matrix $Q = [v_1| \dots |v_m]^t$ the rows of which are the $m$ corresponding eigenvectors. See [10] for more information.

**NMP sub-landscape**

| A \ B | NMP | NMPTrans | PreNeural | EarlyMeso | Meso |
|---|---|---|---|---|---|
| NMP | 0.0 | **0.9** | 1.2 | **2.0** | 3.8 |
| NMPTrans | **0.9** | 0.0 | **0.6** | 1.4 | 4.9 |
| PreNeural | 1.6 | **1.0** | 0.0 | 2.8 | 3.9 |
| EarlyMeso | **3.6** | 3.2 | 4.4 | 0.0 | **0.9** |
| Meso | 6.0 | 7.6 | 6.1 | **1.2** | 0.0 |

**PreNeural sub-landscape**

| A \ B | PreNeural | PreNTrans | EarlyVentral | p0/p1 | p2 |
|---|---|---|---|---|---|
| PreNeural | 0.0 | **0.5** | 1.5 | 0.6 | 1.7 |
| PreNTrans | **0.6** | 0.0 | **0.5** | **0.5** | 0.9 |
| EarlyVentral | 2.6 | **0.7** | 0.0 | 1.4 | 1.6 |
| p0/p1 | 0.6 | **0.3** | 0.7 | 0.0 | **0.4** |
| p2 | 2.3 | 0.6 | 0.6 | **0.5** | 0.0 |

**Intermediate sub-landscape**

| A \ B | p2 | pMN | MNDiff | DPTrans | DP |
|---|---|---|---|---|---|
| p2 | 0.0 | **4.0** | 7.9 | **2.8** | 5.2 |
| pMN | **4.9** | 0.0 | **0.7** | 1.4 | 2.4 |
| MNDiff | 14.6 | **0.7** | 0.0 | 4.5 | 4.7 |
| DPTrans | **3.2** | 0.6 | 3.3 | 0.0 | **0.5** |
| DP | 8.5 | 1.7 | 3.9 | **0.9** | 0.0 |

**Ventral sub-landscape**

| A \ B | EarlyVentral | Early p3 | p3 | FP |
|---|---|---|---|---|
| EarlyVentral | 0.0 | **1.7** | 11.5 | 10.5 |
| Early p3 | **0.9** | 0.0 | **2.1** | **2.0** |
| p3 | 5.9 | **1.3** | 0.0 | **1.5** |
| FP | 8.0 | **3.3** | **3.1** | 0.0 |

**Table A1: ACs separation using normalised KL divergence.** Quantification of the separation of the ACs in each sub-landscape for the time points D4-D8. Each entry corresponds to the normalised KL-divergence $D_{KL}(\alpha_A, \alpha_B)/H(\alpha_A)$ (1) between the distributions $\alpha_A, \alpha_B$ of data points in clusters $A, B$ in $m$-dimensional LDA space. Here $H(\alpha_A)$ is the entropy of $\alpha_A$ (2).

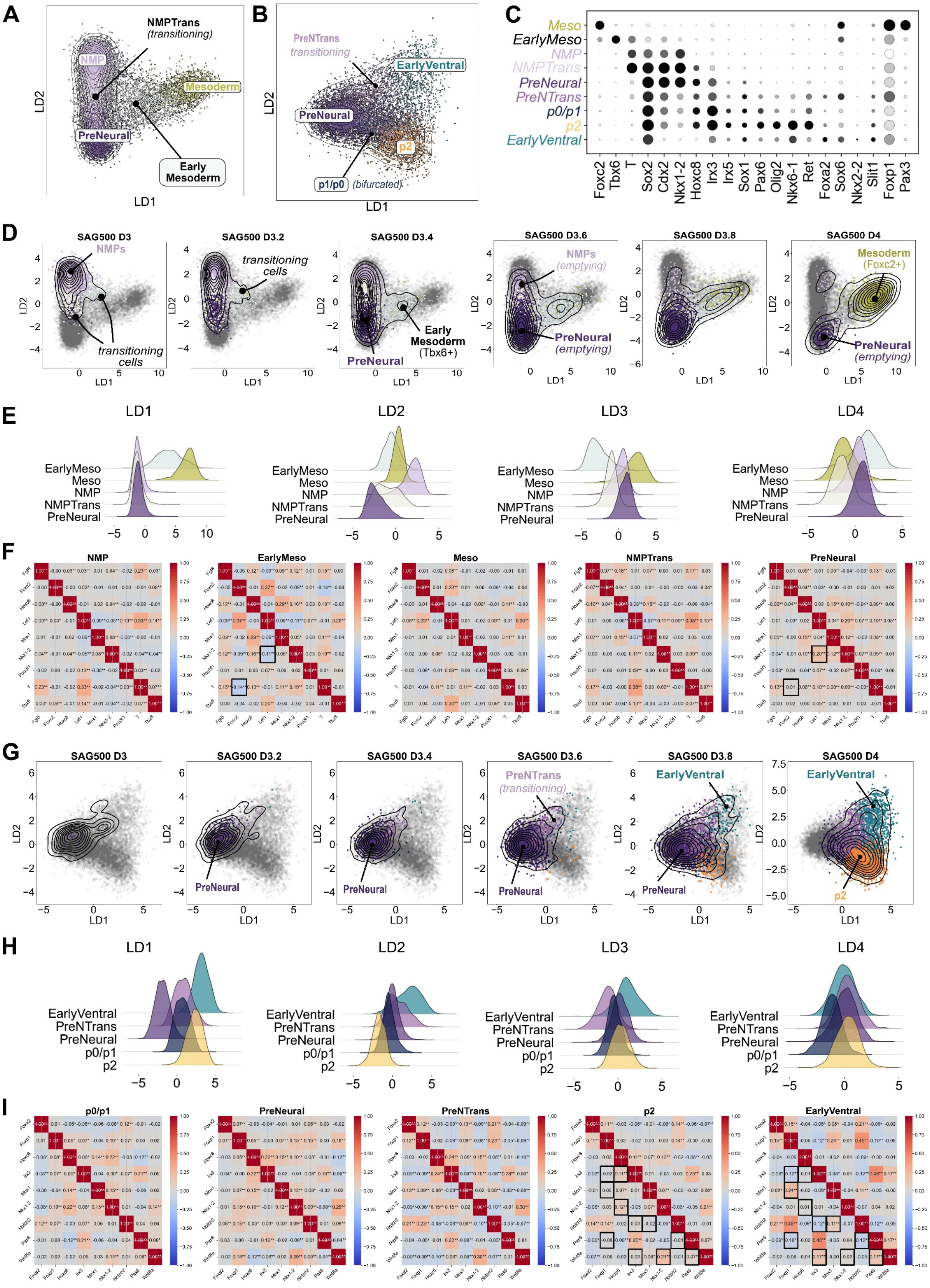

**Attractor Clusters in Early time points scRNA-seq Data**

**Figure A1:** (Caption next page)

**Figure A1:** **A.** 2D LDA projection of the *NMP sub-landscape* that includes the ACs NMP, PreNeural, Early Mesoderm and Mesoderm and the transitioning cluster NMPTrans. **B.** 2D LDA projection of the *PreNeural sub-landscape* that includes the ACs PreNeural, p0/p1 and EarlyVentral and transitioning cells PreNTrans. As the p0/p1 state appears bifurcated we also show p2. **C.** Dot plot indicating the expression of markers used to identify the early ACs (D3-D4). PreNTrans appears to express simultaneously low levels of the markers characteristic of both branches before committing to a specific lineage. **D.** Temporal progression of the ACs in the NMP sub-landscape in 2D LDA space. The NMP AC empties by D3.6 and cells commit to either PreNeural or Mesoderm, the latter involves passing through the intermediate Early Mesoderm AC characterised by Tbx6 expression[15,30]. Grey dots represent the cells at all time points, coloured dots indicate cells present in the indicated samples with the colour indicating their cluster assignment, contour lines the density of cells at the indicated time points. **E.** Unimodal distributions of the LDA scores within each AC within the NMP sub-landscape. The Early Mesoderm AC has broad variance when projected onto the first LD, as does NMPTrans onto the second LD. The Mesoderm AC appears more stable as all components are unimodal with relatively small variance. Fitting a mathematical model to these data further indicated that NMP and PreNeural have just bifurcated at this high SAG concentration (Appendix B Fig. B5E) which explains the rapid leakage of cells from these ACs. A distinction between the means of the various LDA scores is further indication that the clusters are distinct and well-separated. Here LD1 and LD2 capture most of the clusters separation. **F.** Gene-gene correlations (using the raw counts) within each AC in the NMP sub-landscape for all time points using selected pairs of differentially expressed genes. Statistical significance is given by a p-value $< 0.01$ (**) and a p-value between $0.01$ and $0.05$ (*). Distinct correlations for a pair of genes between two ACs suggest a difference in the regulatory interactions. For example the correlation between the pair Lef1-Nkx1.2 is negative in Early Mesoderm ($-0.11^{**}$) but positive in PreNeural ($0.20^{**}$). **G.** Temporal progression of the ACs in the PreNeural sub-landscape in 2D LDA space. It shows leakage of cells from PreNeural from which two distinct branching pathways appear at D3.8, one leading to EarlyVentral, the other to p0/p1 and p2. Grey dots represent the cells at all time points, coloured dots indicate cells present in the indicated samples with the colour indicating their cluster assignment, contour lines the density of cells at the indicated time points. **H.** Unimodal distributions of the LDA scores within each AC within the PreNeural sub-landscape. The PreNTrans cluster is bimodal in the second LD and was initially split into two small Leiden clusters. This cluster is likely to contain a mix of transitioning cells and was then classified as such. The two routes are defined by cells exiting PreNeural, leading either to the target ACs p0/p1 and p2 (intermediate route) or to EarlyVentral (ventral route). **I.** Gene-gene correlations (using the raw counts) within each AC in the PreNeural sub-landscape for all time points using selected pairs of differentially expressed genes. Markers such as Pax6, Irx3 characterise the intermediate route whereas markers such as Foxa2, Foxp1, Notch2 characterise the ventral route. These markers pairs exhibit distinct correlations in the target ACs: for example Irx3-Notch2 ($0.03$ in p2 and $-0.12^{**}$ in EarlyVentral) or Irx3-Foxp1 ($0.07^{*}$ in p0/p1, $-0.02$ in p2 and $-0.12^{**}$ in EarlyVentral).

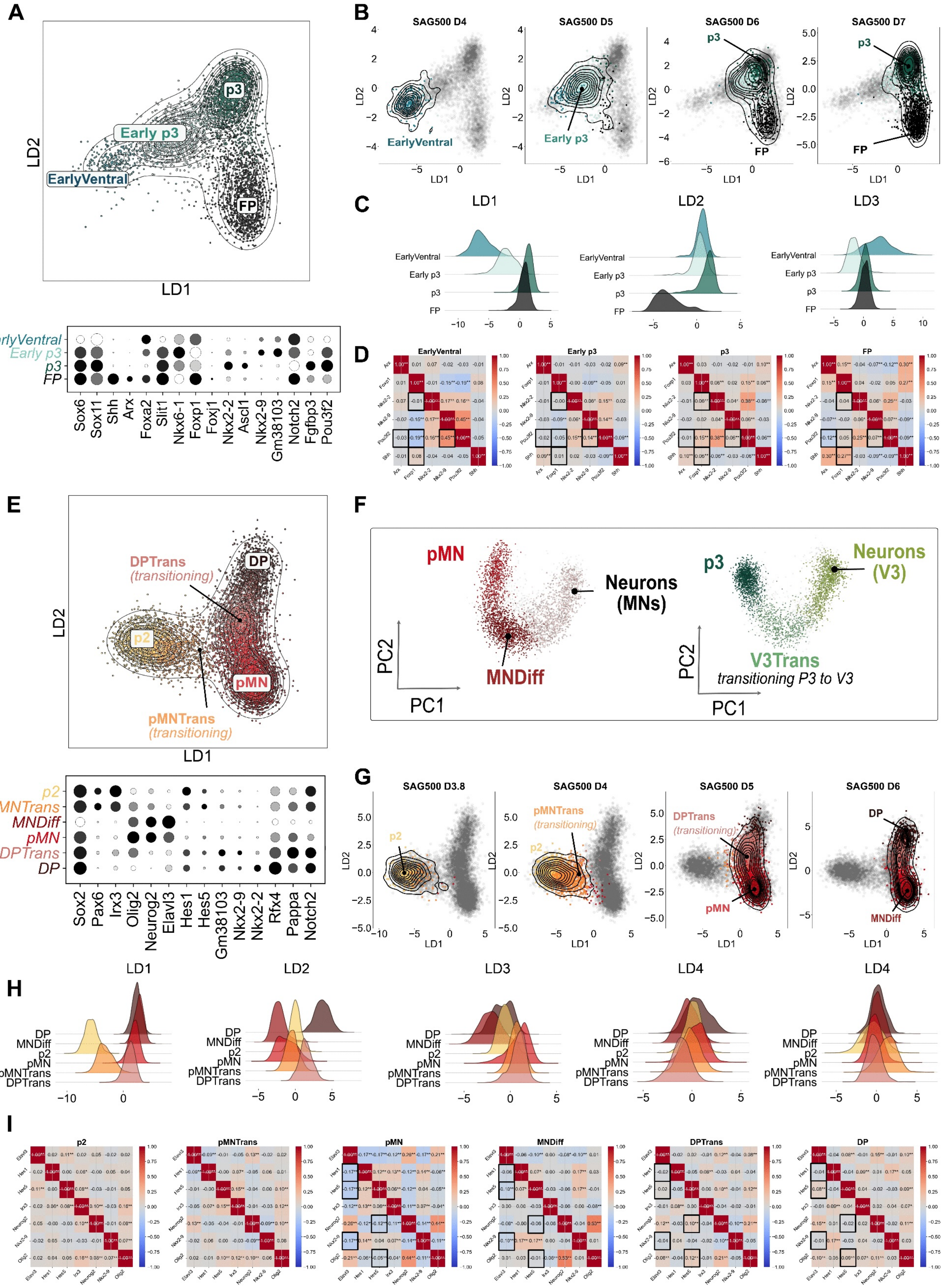

**Attractor Clusters in Late Time Points scRNA-seq Data**

**Figure A2:** (Caption next page)

**Figure A2:** **A.** The *ventral sub-landscape* includes the ACs EarlyVentral (Foxa2, Nkx6.1, Foxp1), Early p3 (Nkx2.9), p3 (Nkx2.2, Ascl1) and FP (Shh, Arx, Foxa2). **B.** The temporal progression of the ACs in 2D LDA space shows a direct transition from EarlyVentral to Early p3 followed by a binary decision to either p3 or FP. Grey dots represent the cells at all time points, coloured dots indicate cells present in the indicated samples with the colour indicating their cluster assignment, contour lines the density of cells at the indicated time points. **C.** Unimodal distributions of the LDA scores within each AC. The transient appearance and wide variance of EarlyVentral in the first LD suggests that this AC may be bifurcated (later confirmed by fitting a model Appendix B Fig. B9F Panel 3). The wide variance of FP in the second component arises from merging with FP an unclassified Leiden cluster that showed moderate expression of Shh and Nkx2.2 (we attributed this to stochastic variations in the mRNA counts). **D.** Gene-gene correlations within each AC for all time points using selected differentially expressed genes. The statistical significance is given by a p-value $< 0.01$ (**) and a p-value between $0.01$ and $0.05$ (*). Distinct correlations for a pair of genes between two ACs indicates a difference in the regulatory interactions. For example the correlation Foxp1-Pou3f2 is equal to $-0.19^{**}$ in EarlyVentral, $0.15^{**}$ in p3 and $0.05$ in FP. **E.** The *intermediate sub-landscape* comprises the ACs p2 (Irx3, Pax6, Nkx6.1), pMN (Olig2), MNDiff (Olig2, Neurog2), DPTrans (Olig2, Nkx2.9) and DP (Olig2, Nkx2.2, Hes1). **F.** 2D PCA representation of the Neurons ACs (MN and V3) that mark the terminal differentiation of pMN and p3 cells together with the intermediate populations MNDiff and V3Trans. **G.** Temporal progression of the ACs in the intermediate sub-landscape in 2D LDA space. Grey dots represent the cells at all time points, coloured dots indicate cells present in the indicated samples with the colour indicating their cluster assignment, contour lines the density of cells at the indicated time points. **H.** Unimodal distributions of the LDA scores within each AC. The second component LD2 further distinguishes the two ACs pMN and DP. However, LD3 is required to separate pMN from MNDiff. **I.** Gene-gene correlations further support the distinction between the two Olig2 lineages, one with pMN and MNDiff, the other with DPTrans and DP (for example Olig2-Hes5 and Neurog2-Hes5 show statistically significant differences within these ACs).

#### A2. Probing expression changes during transition using scRNA-seq

##### A2.1. Overview

This section describes our geometric approach for identifying and characterising the gene expression changes along transitions between ACs. The results validate that observed branching events are binary flip bifurcations, as different sets of genes are up- or down-regulated along alternative pathways. We focus on *sub-landscapes*, which refer to small groups of neighbouring ACs and transitioning cells. LDA (Sect. A1.4) is used to visualise the local dynamics within each sub-landscape. Fig. A4-A6 provide a comprehensive analysis of all sub-landscapes.

##### A2.2. Computational Approximation of gene dynamics along unstable manifolds

We approximate transition routes in LDA space that correspond to unstable manifolds projected from high-dimensional effective gene space. We construct a tubular neighbourhood of these manifolds through transitioning cells. As we assume a heterogenous population of cells, each will be driven by slightly different dynamical systems hence we compute distinct candidates of unstable manifolds.

**Algorithm steps (illustrated in Fig. A4C):**

1. For each AC connected by a route (e.g EarlyVentral, Early p3, p3), fit a Gaussian density kernel to each AC in $m$-dimensional LDA space, find the high density point and then corresponding point in $d$-dimensional effective gene space (defined by the gene module).
2. Fit a piecewise polynomial curve between the relevant high density points (a spline, using Scipy *splrep* that minimises a penalised least squares functional) between these points component by component to produce a curve $\widetilde{\gamma} : [0, 1] \to \mathbb{R}^d$ in effective gene space. The order of the spline (linear, quadratic, cubic) depends on the number of clusters traversed by it.
3. Project $\widetilde{\gamma}$ into a curve $\gamma : [0, 1] \to \mathbb{R}^m$ in LDA space using the projection matrix $Q$ of Sect. A1.4.
4. Divide $\widetilde{\gamma}$ into $N+1$ intervals, marked by points $\{\mathbf{x}_i\}_{i=0}^N$ projecting to $\mathbf{y}_i = Q\mathbf{x}_i$ in LDA space.
5. For each $i$, define the $L_2$-ball of radius $r$ centered at $\mathbf{y}_i$ by
$$B_r(\mathbf{y}_i) = \{\mathbf{y} \in \mathbf{R}^m : \|\mathbf{y} - \mathbf{y}_i\| < r\}$$
where $\| \cdot \|$ is the $L_2$-norm. The union of these balls defines a tubular neighbourhood of $\gamma$ and therefore all data points in the effective gene space that map into $B_r(\mathbf{y}_i)$ are close to $\widetilde{\gamma}$ if $r$ is not too large.
6. For each ball $B_r(\mathbf{y}_i)$ covering $\gamma$, we compute the average raw counts (or log-normalised counts in Fig. A3) for each gene $g$ across cells in the ball. If flow cytometry data are used, we use the protein concentration instead of the mRNA counts.
7. We then fit cubic splines to obtain gene expression curves along $\gamma$ (Fig. A3).

The method is robust as the results show small dependence on the balls radius $r$ (tested for $r$ ranging from 0.4 to 1 in Fig. A3). If $r$ is too small, the drop-outs cause larger variations. What matters is to take sufficiently many balls to cover $\gamma$ entirely, without comprising cells that transition through a different pathway.

##### A2.3. The Critical Transition Index

Given a ball cover of an approximated unstable manifold $\gamma$, we can compute the *Critical Transition Index* of Mojtahedi et al.[24] in each ball and plot its variation along $\gamma$. This index is an indicator of a cell state transition. For each ball $B$, the index is defined as

$$I_c(B) = \frac{\langle |R(\mathbf{g}_i, \mathbf{g}_j)| \rangle_{i,j}}{\langle R(\mathbf{x}_k, \mathbf{x}_l) \rangle_{k,l}}$$

where $\mathbf{g}_i$ is the vector of expressions of the $i$th gene accross all cells, $\mathbf{x}_k$ is the vector of gene expression for cell $k$ and $R$ is the Pearson's correlation coefficient between two vectors. The brackets $\langle\cdot,\cdot\rangle$ denotes the mean over all pairs indicated and $|\cdot|$ denotes the absolute value. The index compares variability of a gene across a set of cells with the variability of distinct genes within one single cell. As expected, we note an increase of the index as cells exit an attractor when transitioning along an unstable manifold (Fig. A4E top).

Note that whereas in[24], the transition index is computed with respect to pseudo-time ordering of the cells, we compute it along the balls that cover $\gamma$.

Alternatively, we could compute the index directly in LDA space by replacing the $\mathbf{x}_i$ by $\mathbf{y}_i$ and the $\mathbf{g}_j$ replaced by the vector made up from the $j$th LDA scores across all cells (Fig. A4E bottom).

##### A2.4. Example applications to key sub-landscapes

**Ventral sub-landscape (Fig. A4):** contains four ACs: EarlyVentral (Foxa2, Foxp1), Early p3 (Nkx2.9, Foxp1), p3 (Nkx2.2) and FP (Shh, Foxa2, Arx) in 3D LDA space. Two types of transitions are observed (Fig. A4B): a direct transition from EarlyVentral to Early p3 (fold bifurcation destroying EarlyVentral), and a subsequent binary decision from Early p3 to either p3 or FP (binary flip bifurcation).

**PreNeural sub-landscape (Fig. A5):** contains the decision from PreNeural at D3.8 towards either an intermediate fate (Pax6) or a ventral fate (Foxa2)[7]. The ventral branch is dependent on high concentration of SAG as validated by FACs experiments (Sect. A4.1). The exit of PreNeural is marked by downregulation of Nkx1.2 and Cdx2 and up-regulation neural progenitor genes (Fig. A5D). Cells along the ventral route up-regulate Foxa2, Slit1, whereas cells taking the intermediate route up-regulate Pax6, Irx3, Olig2. The Critical Transition Index peaks as cells exit PreNeural (Fig. A5E).

**The intermediate sub-landscape (Fig. A6):** contains the six clusters (ACs: p2, pMN, MNDiff, DP, and the transitioning states pMNTrans, DPTrans). By including p3, the corresponding LDA space is 6-dimensional. This sub-landscape contains a complex set of transitions: one from p2 to either pMN or DP followed by a binary choice decision from pMN (Olig2) to either MNDiff (Olig2, Neurog2) or DP (Olig2, Nkx2.9, Hes1). A cell in the pMN attractor transitions to one of these two options, depending on whether the pMN attractor is closer to the left or right saddle (Fig. A6C). The pMN-to-DP route up-regulate Nkx2.9 and maintains Hes1 (Notch target); the pMN-to-MNDiff route represses Hes1 and up-regulate Neurog2, Dll1 (both required for neuronal differentiation[29]).

The progressive loss of DP cells by D7 with convergence towards p3 supporting a direct DP-to-p3 transition (Fig. A7A-B). Transition analysis (Fig. A7C-D) reveals that the intermediate (yellow) and ventral (green) routes to p3 are mainly distinguished by markers such as Foxa2 and Olig2. Lineage studies support this connection: while FP cells rarely derive from Olig2-expressing precursors, a subset of p3 progenitors descend from cells with an Olig2 expression history[8].

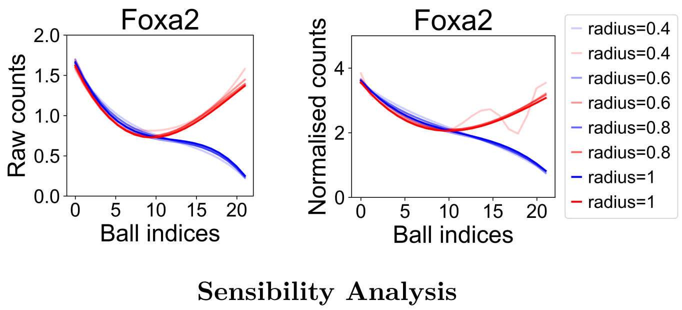

**Figure A3:** mRNA counts of the cells inside each ball are averaged to determine how gene expression changes along each unstable manifold (blue and red). The horizontal axis is labelled according to the ball indices $0$ to $N = 20$. Splines are for different ball raddii. (Left) Raw counts. (Right) Log-normalised counts. We note no major difference except that drop-outs lead to more noisy tendencies if the ball radius is taken too small (radius$= 0.4$).

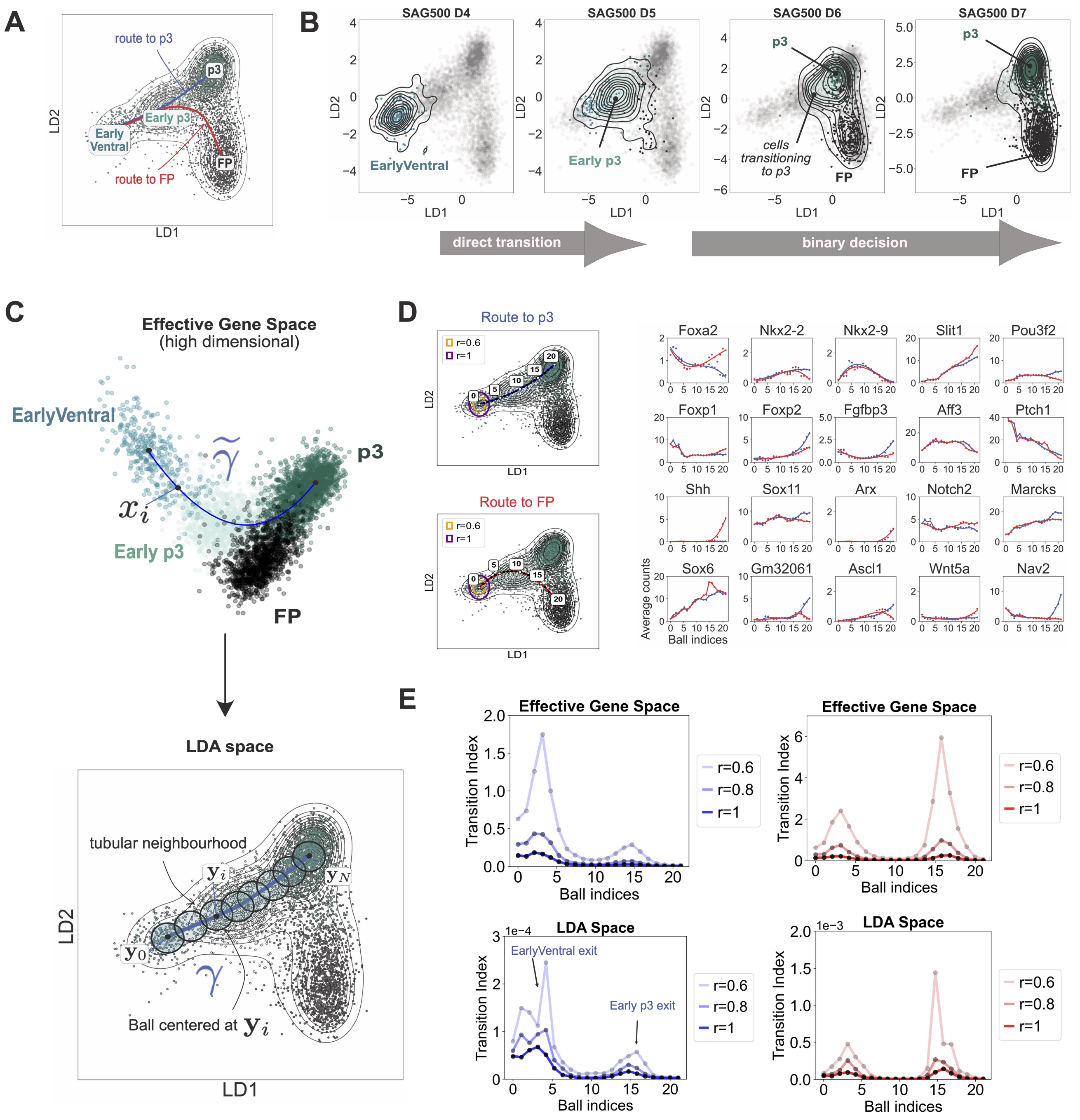

**Transitions in the Ventral Sub-landscape**

**Figure A4:** **A**. 2D LDA representation of the ventral sub-landscape using data from all time points. We highlight two distinct routes starting at the EarlyVentral state: one route $\gamma$ leads to p3 (blue) and one route leads to FP (red). **B**. Temporal progression of the ACs from D4 when cells populate EarlyVentral (Foxa2, Foxp1) to D7 when cells end up in the two different states p3 (Nkx2.2) and FP (Foxa2, Arx) via an intermediate state labeled Early p3 (Nkx2.9). At D4, cells briefly remain in EarlyVentral before transitioning to Early p3 at D5 (direct transition). Subsequently, at D6, some cells head towards p3 while others head towards FP (binary decision). Grey dots represent the cells at all time points, coloured dots indicate cells present in the indicated samples with the colour indicating their cluster assignment, contour lines the density of cells at the indicated time points. **C**. Approximation of an invariant manifold $\widetilde{\gamma}$ in effective gene space from EarlyVentral to p3 projected onto $\gamma$ in 3D LDA space. The projected curve $\gamma$ is divided into equal intervals marked by ticks $y_0, y_1, \ldots, y_N$. Each ball is centred at one point $y_i$ and has fixed radius $r$ which is chosen such that we cover the invariant manifold by sufficiently many balls containing enough data points. Together, these balls form a tubular neighbourhood of the projected unstable manifold. **D**. (Left) Projected splines for the two distinct routes, from EarlyVentral to either p3 or FP ACs, with ticks $y_i$ (numbered) in LDA space. (Right) Balls (here of radius $r = 0.8$) are placed along the projected curves in LDA space and raw mRNA counts of the cells inside each ball are averaged to determine how gene expression changes along each unstable manifold. The horizontal axis is labelled according to the ball indices $0$ to $N = 20$. **E**. Critical Transition index[24] $I_c(B)$ computed in each ball $B$ along the approximated unstable manifolds to p3 (blue) and to FP (red) show peaks as cells transition from one AC to another. Colour variations indicate different ball radii. $I_c$ computed in effective gene space (Top) and in 2D LDA space (Bottom).

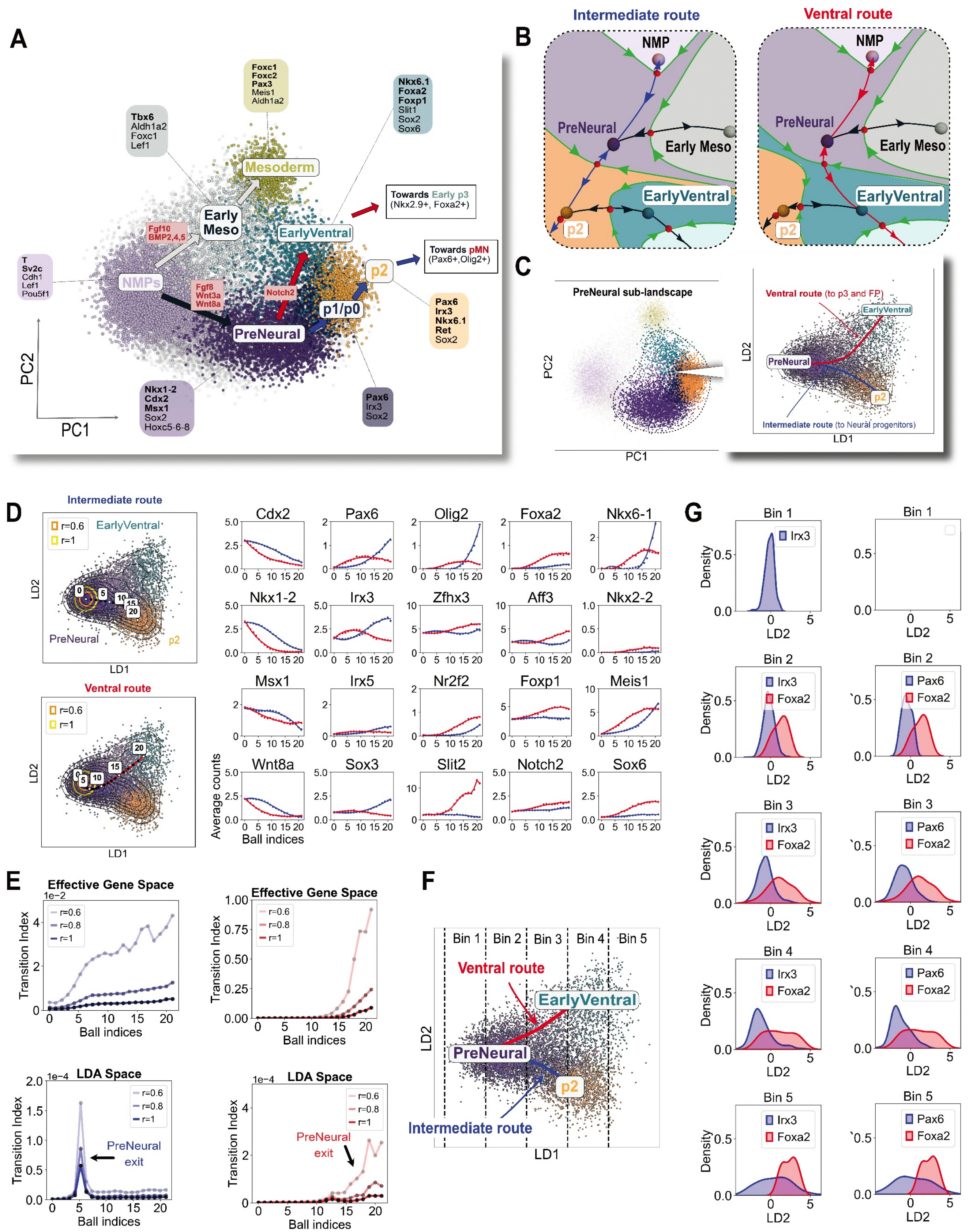

**Transitions in the PreNeural Sub-landscape**

**Figure A5:** (Caption next page)

**Figure A5:** **A**. 2D PCA projection of the combined NMP and PreNeural sub-landscapes (D3-D4). **B**. Phase plots of local dynamical systems. The stable manifolds (green) partition the plane into distinct basins of attraction, each containing one attractor. The two systems differ in the configuration of their unstable manifolds: in the left panel, the unstable manifold (blue) directs a PreNeural cell towards p0/p1, whereas in the right panel, the unstable manifold (red) directs the cell towards EarlyVentral. **C**. We extracted the PreNeural sub-landscape, including p2, and projected it in 2D LDA space. We distinguish two distinct routes that cells can take as they differentiate. **D**. Projections in 2D LDA space of the approximated unstable manifolds in effective gene space corresponding to the two routes. Markers of the dorsal identities such as Pax6 and Irx3 are up-regulated along the intermediate route (blue) while ventral markers such as Foxa2 and Foxp1 get up-regulated along the ventral route (red). Balls of distinct radii $r$ are shown in orange and yellow. **E**. Critical Transition index[24] computed in each ball along the approximated unstable manifolds in effective gene space (top) and in 2D LDA space (bottom). Colour variations indicate different ball radii. **F**-**G**. The cells are divided into bins along the LD1 component (F). They are further assigned to one samples: cells expressing Pax6, Irx3 and/or Foxa2. A cell requires at least 3 mRNA counts of the corresponding gene to belong to a sample. Distributions of the LD2 scores for the found samples show a progressive divergence as cells differentiate along the unstable manifolds (G).

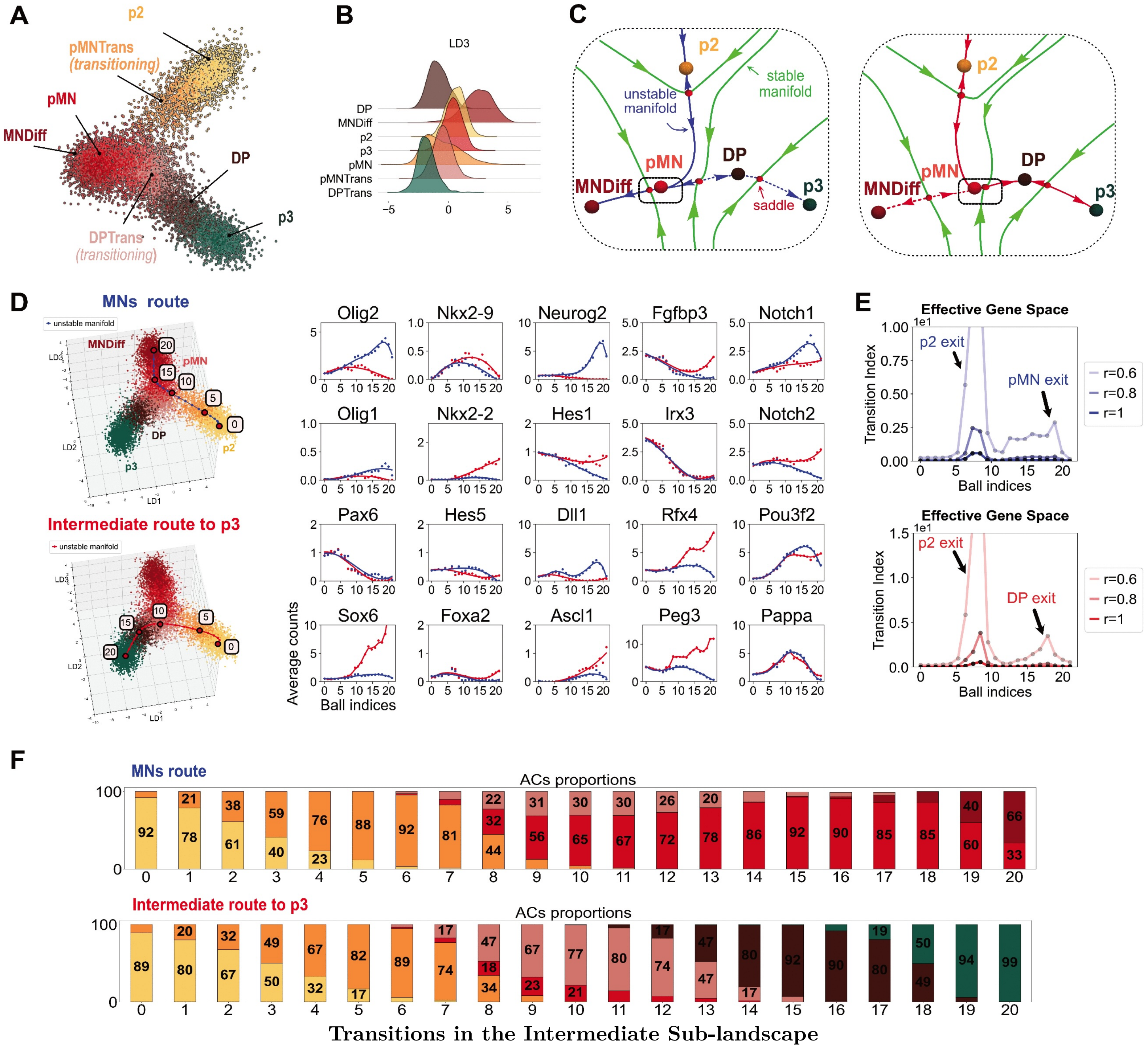

**Figure A6:** **A**. 2D LDA representation of the intermediate sub-landscape (ACs p2, pMN, MNDiff, DP, and transitioning clusters pMNTrans, DPTrans), including the p3 AC. The ACs MNDiff and pMN are not well separated using only two LDs. **B**. The third LD separates the ACs pMN and MNDiff. **C**. Binary choice decision: the pMN attractor can bifurcate with either its left saddle (left panel) leading a pMN cell to transition to MNDiff, or with its right saddle (right panel) and the pMN cell transitions to DP. We postulate that the proximity of pMN with its right saddle correlates with activation of Nkx2.9 while maintaining Hes1 expression. **D**. (Left) 3D LDA representation of the intermediate sub-landscape, including p3. The two fitted cubic splines (blue and red) indicate the routes towards MNDiff and p3, respectively. (Right) Gene variations along the two routes show that the branching is characterised by up-regulation of Neurog2 on the motor neuron differentiation route and up-regulation of Hes1, Nkx2.9 on the intermediate route to p3. **E**. Critical Transition index[24] computed in each ball along the approximated unstable manifolds (in effective gene space). Colour variations indicate different ball radii. **F**. Proportions of the distinct cell types in each ball along the tubular neighbourhood for each of the two routes in D. The x-axes correspond to the ball indices.

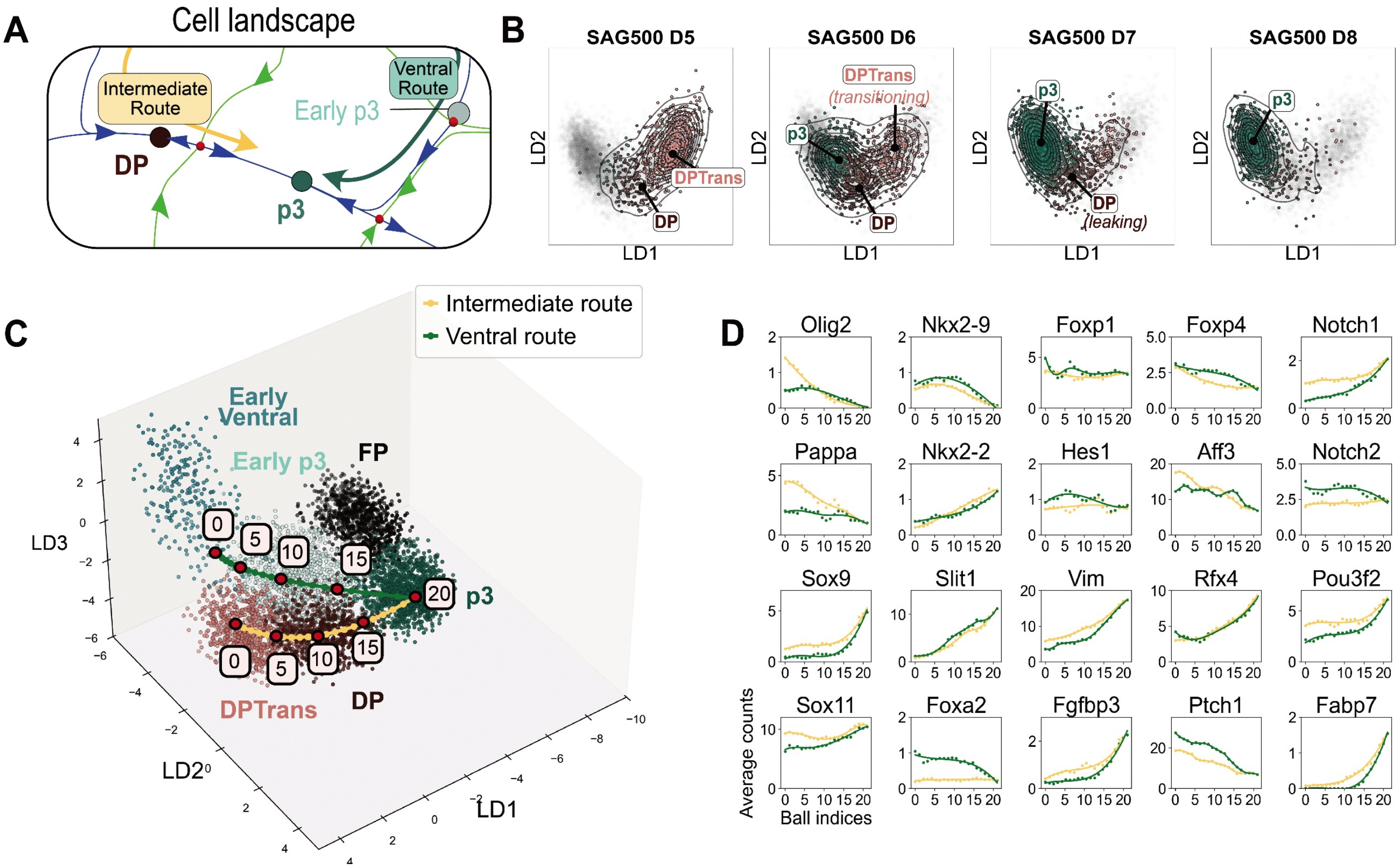

**The DP to p3 Transition**

**Figure A7:** **A.** Local view of a cell's landscape around the DP-to-p3 transition. The circular topology allows cells to access p3 via two distinct routes (ventral and intermediate). **B.** Transition from DP to p3 in the scRNA-seq dataset at 500nM SAG that we extended until D8. Plots indicate distribution of cells at the indicated time points in LDA space. **C.** 3D LDA representation of the ventral ACs with approximation of the two routes ventral (green) and intermediate (yellow) using the transition analysis. Both start in an AC at D5 (Early p3 for ventral and DPTrans for intermediate). **D.** Gene expression dynamics along the two routes to p3, ventral (green) and intermediate (yellow). It shows that cells that reach p3 along the intermediate route have an history of Olig2 expression.

### A3. FACs analysis

To further validate the clusters found in the scRNA-seq dataset and to better understand their persistence and stability under varying signalling conditions, we quantified the allocation of cells to these clusters over time in different signalling conditions using single-cell flow cytometry data (FACs). We identified a set of five markers - Sox2, Pax6, Olig2, Nkx6.1, and Nkx2.2 - as sufficient to classify most ventral progenitors[7]. A sixth marker Tubb3 was included to exclude neurons. Using these, we generated timeseries FACs data, collected at 3 time points corresponding to days D4, D5 and D6, where cells were continuously exposed to four SAG concentrations (0nM, 10nM, 100nM, 500nM). We conducted 5 experimental series named *March 12*, *April 16*, *October 16*, *Nov 5*, *June 5* and selected the Sox2 expressing cells, removing the neurons. The replicates *Nov 5* and *June 5* were only exposed to 0nM and 500nM SAG. As the data revealed a consistent level of Tubb3 across all cells we did not use it in further analysis.

#### A3.1. Clustering using Gaussian Mixtures Models

For each individual experimental replicate (e.g. *March 12*), we used the methodology of[28] to identify clusters in the resulting 5-dimensional gene expression space by applying a probabilistic Gaussian Mixture Model (GMM)[23] using the algorithm implemented in the Python package *sklearn.mixture*[26] or the Matlab function *fitgmdist*[21]. Multivariate Gaussian distributions were fitted to each experimental replicate separately, including all SAG concentrations and all time points. To ensure comprehensive coverage of attractor clusters, we applied a high-resolution clustering, yielding approximately 25 weighted Gaussian distributions. We validated the cluster assignments by verifying that each distribution's projection onto individual marker coordinates exhibited unimodality, indicative of a stable cell state. We merged clusters with similar gene expression profiles based on the Kullback-Leibler divergence metric (see below), thereby establishing a unified GMM representation.

#### A3.2. Assigning cells identities

We found all the expected cell states (Fig. A8 A): PreNeural at D4 (Sox2), p0/p1 (Pax6), p2 (Pax6, Nkx6.1), pMN (Olig2), DP (Olig2, Nkx2.2) and p3 (Nkx2.2). Without the markers Dbx1 and Dbx2, we cannot distinguish the progenitors p0 and p1 based on Pax6 expression levels, so we label this cluster p0/p1. We also identified the ACs that refine the p3 landscape with EarlyVentral (Nkx6.1) and Early p3 which has a lower expression of Nkx2.2 than p3 (Fig. A8B). The MNDiff cluster (co-expressing Olig2 and Neurog2 in the scRNA-seq dataset) could also be distinguished from pMN by its higher level of Olig2 and lower levels of Pax6 and Sox2 (Fig. A8B). Several clusters were labelled as transitioning (e.g p2Trans) and these choices are justified in Sect. A3.3.

By contrast with the scRNA-seq dataset, we did not identify neurons because they were excluded by selecting Sox2$^+$ cells. Although the FP marker gene Foxa2 is lacking, we expect FP cells to co-express Sox2 and Nkx6.1. The FP cluster should only appear at high SAG concentrations (100nM, 500nM) at D6 and later. The *March 12* experiment included unclassified cells (labelled 'other') that are candidates for FP cells (Fig. A8B), but this observation was not reproduced in other experimental replicates. For each experimental replicate we computed the proportion of cells in each cluster for each time point and for each SAG concentration (Fig. A8C). This shows that the presence or absence of specific cell states are dependent upon the level of SAG. Finally, some cells were left as unclassified as these were not consistently identified in all the experimental replicates. One of these unclassified clusters (labelled *NeuronTrans*) contains cells with low Pax6, low Sox2 and high Olig2 and appear at D4.

#### A3.3. Clusters quality

For FACs data we use similar criteria as for the scRNA-seq to assess the quality of the clusters.

1. *Unimodality:* In each cluster, the expression level of each of the individual markers follows a unimodal distribution. To merge two clusters $A, B$ together, given by multivariate Gaussian distributions $\alpha_A, \alpha_B$ in 5-dimensional gene expression space, we use the formulation of the KL-divergence $D_{KL}(\alpha_A, \alpha_B)$ (1) in dimension $m = 5$. This quantity does not define a distance function as it is not symmetric. We thus define the *KL-metric* by taking the symmetrisation of it:

$$d_{KL}(\alpha_A, \alpha_B) = D_{KL}(\alpha_A, \alpha_B) + D_{KL}(\alpha_B, \alpha_A).$$

   When two GMM clusters have distributions sufficiently close to each other with respect to this metric they can be merged together so that the resulting distribution remains unimodal in each component (Fig. A9A). This distinguishes clusters based on the expression level of the markers. The Gaussian structure of the clusters is also revealed by projecting them onto the expression levels of a pair of markers (e.g. Fig. A9D-E).

2. *Temporal progression:* Neighbouring clusters are extracted and their temporal dynamics analysed in LDA coordinates for each SAG concentration. For example, Fig. A9B (10nM SAG) shows a direct transition between the ACs p0/p1 and p2 with transitioning cells in between (cluster p2Trans). The KL-metric defined above can be used to quantify the variation between the distribution of an AC at a given time (given by its mean and covariance matrix) and the same AC at a later time.

3. *Local gene-gene correlations:* Distinct cell identities are further validated by testing for distinct correlations within each cluster (Fig. A9C).

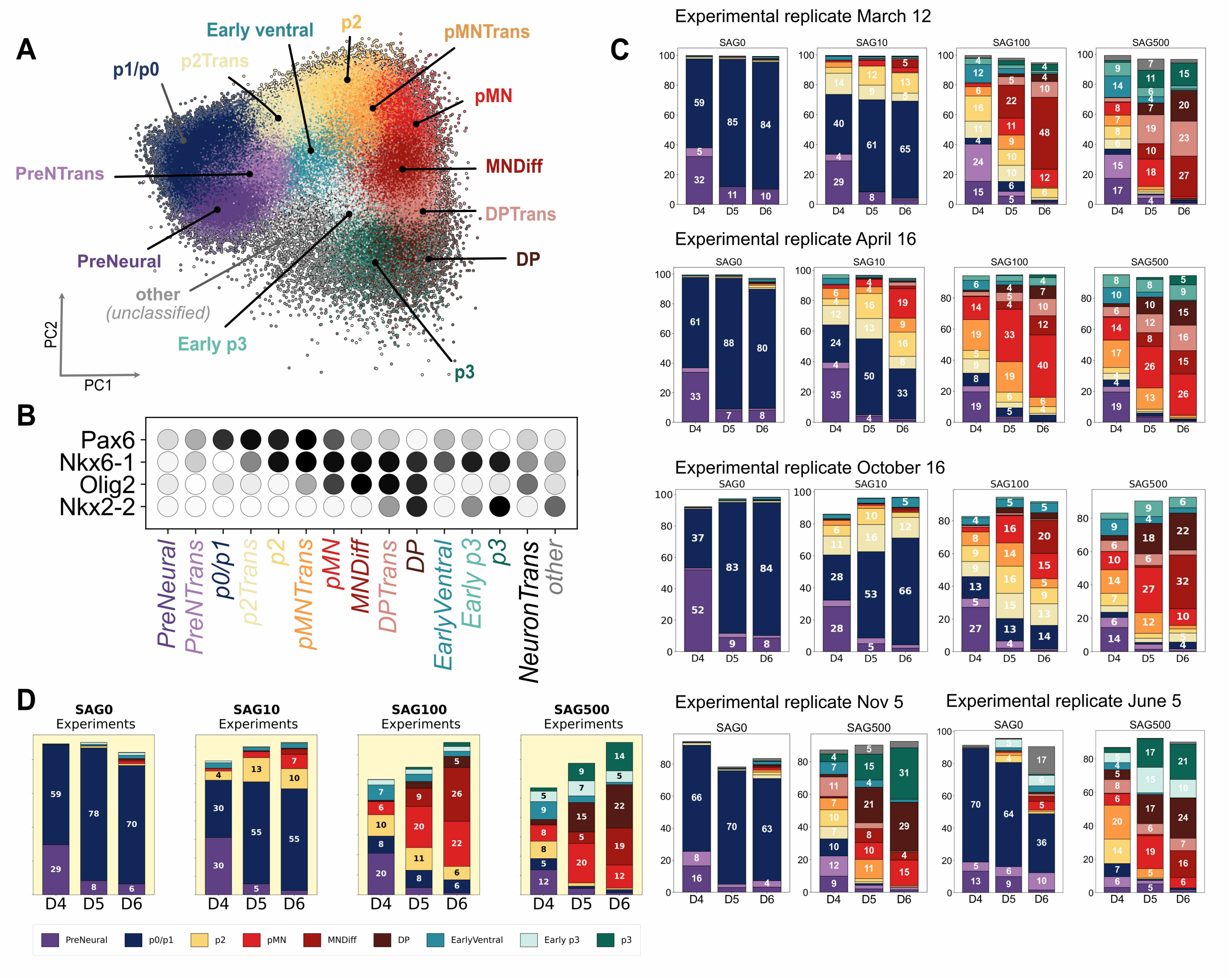

**Attractor Clusters in FACs**

**Figure A8:** **A**. PCA representation of all the GMM clusters (*March 12* replicate) after being amalgamated based on the KL-divergence metric. We find all the expected cell states PreNeural, p0/p1, p2, pMN, DP and p3. In addition we also identify states that refine the p3 lineage such as EarlyVentral (Nkx6.1$^+$, Pax6$^-$) and Early p3, characterised by a lower expression of Nkx2.2 than p3. We also identify MNDiff cells, which are distinguished from pMN cells by lower levels of Pax6 and Sox2, and higher levels of Olig2. The clusters of cells transitioning between two ACs are labelled as *-Trans*. **B**. Dot plot indicating the expression of the markers used to identify the ACs and transitioning clusters (*March 12* replicate). Darker shading indicates higher levels of expression of the indicated marker (z-score). **C**. Proportions of cells (in percent) in each GMM cluster as a function of time and SAG concentration for each of the 5 experimental replicates. The gaps that can be observed in the different experiments correspond to unclassified cells that have low expression of Pax6 and higher expression of Olig2 (labeled 'NeuronTrans'). **D**. ACs proportions obtained by averaging all the experimental replicates in C for each day and each SAG concentration.

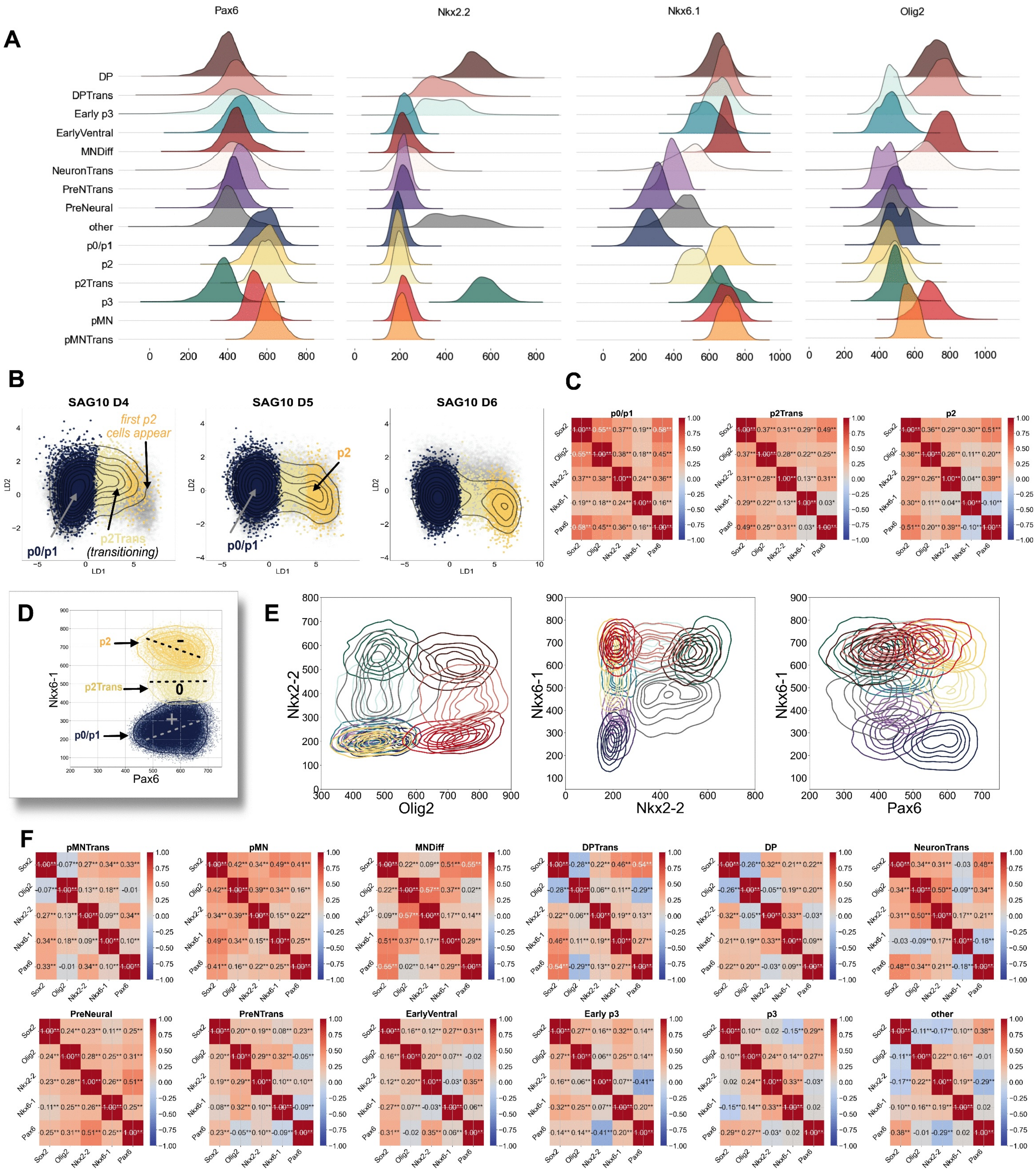

**Quality Criteria for Attractor Clusters in FACs Data**

**Figure A9:** (Caption next page)

**Figure A9:** The Figure represents data from the *March 12* replicate: the other datasets are similar. **A**. Distributions of the marker genes, across time points and SAG concentrations in each cluster distinguish the different cell states. Note that p0/p1 is bimodal at 0nM SAG due to variability of Pax6 and Olig2 in these cells over time. This may be due to these cells becoming V0/V1 neurons under continuous exposure to low SAG concentration. For simplicity, we kept only one AC. The unimodality criterion is satisfied for the ACs PreNeural (Sox2), p0/p1 (Pax6), EarlyVentral (Nkx6.1), Early p3 (Nkx2.2$^{+}$) and p3 (Nkx2.2$^{++}$). A cluster labelled 'other' corresponds to unclassified cells, but it does not appear in all the experimental replicates. Similarly PreNTrans is not densely populated in all experimental replicates. It is however distinct from PreNeural by a higher level of Nkx6.1 and Pax6 suggesting that it contains a mix of cells that are transitioning towards either p0/p1 or EarlyVentral (also suggested by the bimodality of the Olig2 marker). **B**. Temporal progression of the triad p0/p1, p2Trans (transitioning) and p2 at 10nM SAG in LDA coordinates. Cells are gradually transitioning from p0/p1 towards p2. Grey dots represent the cells at all time points, coloured dots indicate cells present in the indicated samples with the colour indicating their cluster assignment, contour lines the density of cells at the indicated time points. **C**. Correlations between markers within each cluster for all time points and SAG concentrations. There is a change in regulatory interaction between Nkx6.1 and Pax6 as cells transition from p0/p1 to p2. **D**. Projected clusters in the gene space components Nkx6.1 and Pax6 for all days at 10nM SAG with correlations between Nkx6.1 and Pax6. **E**. Projected clusters in some of the gene space components (NeuronTrans and pMNTrans not included for clarity). **F**. Gene-gene correlations for each AC and transitioning cluster across all time points and SAG concentrations. This provides further validation that the ACs are distinct. The correlation between Pax6 and Olig2 in pMN ($0.16^{**}$) and MNDiff ($0.02^{**}$) are different supporting their distinct identities. Their gene expression pattern also shows that they do not share the same level of Pax6, Olig2 and Sox2. In the scRNA-seq these clusters are distinct because MNDiff shows higher levels of Neurog2 and Elavl3. Finally, the correlation between Olig2 and Pax6 marks a difference between the DP AC ($0.20^{**}$) and the transitioning cells DPTrans ($-0.29^{**}$).

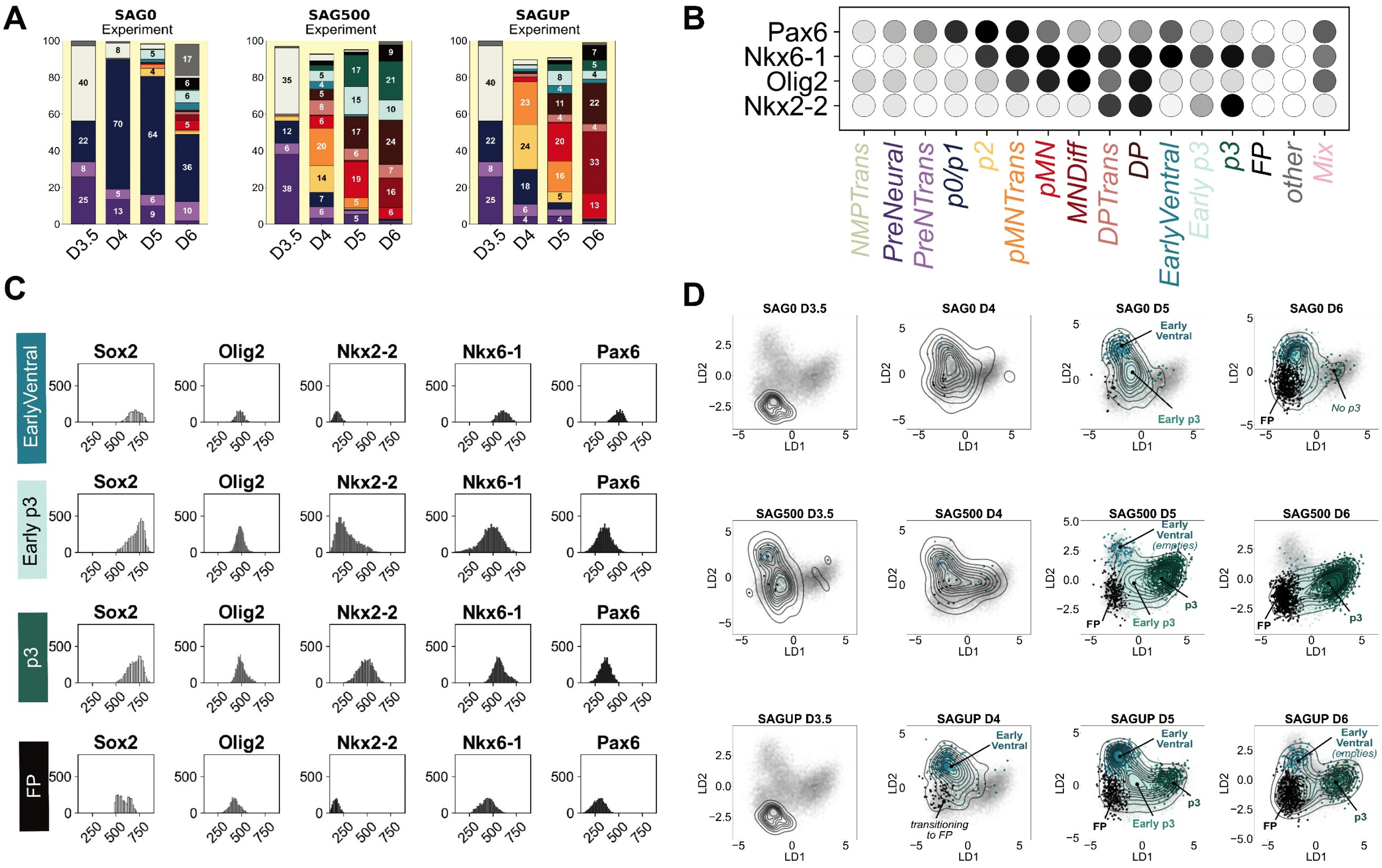

**FP Cluster in Early Pulse Experiment**

**Figure A10:** **A**. Replicate *June5*: experimental proportions of cell types (in percent) for fixed SAG concentrations 0nM, 500nM, and the pulse experiment SAGUP: 0nM SAG from D3 to D3.5, 500 nM SAG from D3.5 to D4 and then 0nM SAG until D6. **B**. Dot plot indicating the expression of the markers used to identify the ACs and transitioning clusters. Darker shading indicates higher levels of expression of the indicated marker (z-score). **C**. Unimodal distributions of the FACs markers for the 4 ventral ACs. We identified a FP-like cluster with co-expression of Nkx6.1 and Sox2. **D**. Temporal progression of the ventral ACs for the constant SAG concentrations 0nM, 500nM and the pulse experiment SAGUP. We observe a binary decision at D5 from Early p3 to p3 and the FP-like AC.

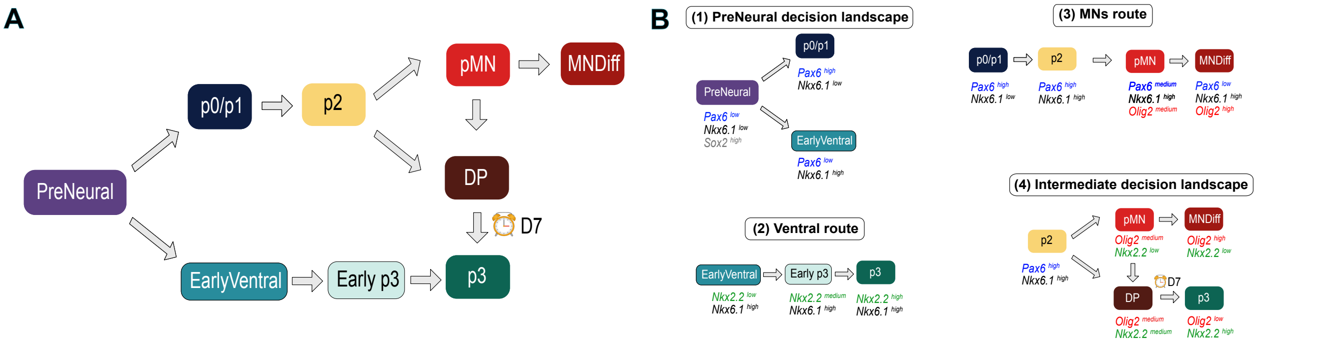

**Decision Landscape Schematic from FACs Analysis**

**Figure A11:** **A.** Schematic representation of the decision landscape as suggested by the FACs datasets. It displays all possible transitions we observe in this dataset. The scRNA-seq dataset indicates further connections between DP and p3 and a binary decision from Early p3 to either p3 or FP. The FP AC is not visible in the FACs, but it is clearly observed in the scRNA-seq dataset. For this reason we refer here to the *ventral route* rather than the *ventral sub-landscape*. **B.** Schematic representation of the decision sub-landscapes and routes that will be analysed in LDA coordinates.

##### A3.4. Early pulse experiment June5

The replicate *June5* includes an experiment labelled "SAGUP" in which 0nM SAG was applied to cells from D3 to D3.5 (12h), followed by 500nM SAG from D3.5 to D4 (12h) and then cells were returned to 0nM SAG until D6. Constant exposure to 0nM and 500 nM where compared as controls. The proportions of cells in each cluster by time point are shown in Fig. A10A and their gene expression profile in Fig. A10B. At the early collection stage D3.5, we identified cells referred to as NMPTrans (white) that we hypothesised are transitioning from NMP (they do not appear at later time points). These cells do not express Sox2 and cannot therefore be classified as PreNeural. A distinct cluster with floor plate-like properties emerged at D6 across all experiments (Fig. A10C). Cells in this cluster co-express Sox2 and Nkx6.1 but remain distinct from the EarlyVentral population and the temporal progression of the ventral sub-landscape (Fig. A10D), analysed in LDA coordinates, matches the patterns seen for the corresponding p3 transition in the scRNA-seq dataset (Fig. A4B). This cluster was also seen at low SAG concentration 0 nM. Cells appear to reach this state from EarlyVentral at D5, bypassing the p3 state. However, when an increased concentration to 500nM SAG is applied to the system, p3 cells emerge at D5 (Fig. A10D).

#### A4. Decision landscape and transitions in FACs data in response to SAG perturbations

This section provides a detailed analysis of how the SAG perturbations affect the different sub-landscapes by analysing their temporal progression in LDA space. This enables a schematic representation of the decision landscape and its sub-components (Fig. A11). This diagram can be further refined and expanded as more data are incorporated. For example, the scRNA-seq dataset revealed extra ACs NMP, Early Mesoderm, Mesoderm and FP, and a transition route between DP and p3 at D7.

We observed branching governed by binary flips for the transition from PreNeural to either p0/p1 or EarlyVentral and potentially for the transition from p2 to either pMN or DP. We focus on specific transition routes that certain groups of cells follow, which allows us to determine the presence or absence of attractors along these routes. The figures presented in this section for illustration are derived from the *March 12* replicate, but our conclusions apply to all replicates.

##### A4.1. The PreNeural decision landscape

The FACs data at D4 reveal a binary decision from PreNeural towards either p0/p1 or EarlyVentral (Fig. A11B (1)) consistent with a recent study[7]. Even in the absence of the characteristic markers Foxa2 and Foxp1, the expressions of Pax6 and Nkx6.1 alone are sufficient to distinguish EarlyVentral from p0/p1 (Fig. A9A). Cells in PreNeural are characterised by high expression of Sox2 and low expression of the other markers.

We observe the following SAG-dependent bifurcations (Fig. A13A-C):

- *Low SAG (0nM, 10nM):* the transition to EarlyVentral is absent and all cells transition to an intermediate fate (p0-p1, pMN). Cells that appear to transition to EarlyVentral, which can be attributed to stochastic fluctuations, remain confined to EarlyVentral without any further progression to p3 (Fig. A13D). This is a further indication that EarlyVentral can be viewed as an AC rather than a transitioning cluster.

- *High SAG (100nM, 500nM)*: two distinct pathways emerge (ventral and intermediate). We thus postulate that, for each cell, while the p0/p1 and EarlyVentral attractors are present and deep at low SAG, they bifurcate at high SAG, a prediction validated by the model in Appendix B Sect. B5.1 (Fig. B7A Panel (3)). The PreNeural does not persist for all time points and declines rapidly at high SAG concentrations. This suggests that, when we fit a mathematical model, the resulting parameters should be distributed close to a fold-flip point (Fig. A12), a prediction verified below in Appendix B Sect. B5.1 (Fig. B7A Panel (2)).

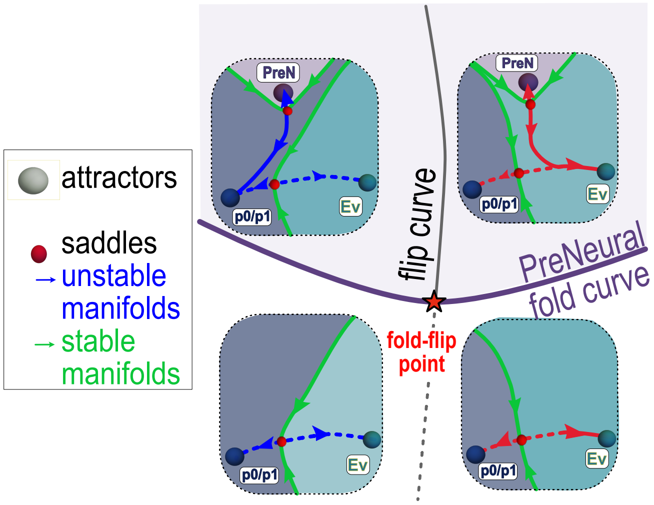

**Figure A12:** Flip bifurcation of the PreNeural sub-landscape. It shows sub-landscape model representatives in distinct regions of parameter space separated by fold and flip bifurcation curves intersecting at a fold-flip point (red asterisk). The unstable manifold from PreNeural can connect to p0/p1 or to EarlyVentral (Ev), depending on which side of the flip curve the parameters are located.

##### A4.2. The ventral route

The transition from EarlyVentral to the more ventral states Early p3 and p3 is marked by the increase in expression of Nkx2.2. This route emerges when the SAG concentration is 100nM or higher (Fig. A13D,F). Here, Early p3 appears to be a transient state for two reasons: first, there is a large variance in the Nkx2.2 expression (Fig. A9A), and second, unlike EarlyVentral and p3, this cluster lacks a well-defined structure in LDA space (Fig. A13D). This prediction is confirmed by fitting the model in Appendix B Sect. B5.1 (Fig. B7A Panel (6)). However, this outcome depends on the dataset as the fit of the model to two other scRNA-seq datasets (mouse and human) suggests that Early p3 is an unbifurcated, shallow AC (Appendix B Sect. B7 and Sect. B9.1).

Due to the lack of markers in the FACs panel, we could not identify FP cells (Nkx6.1, Sox2, Foxa2, Arx); however, these cells are predicted to appear at D6 in response to high SAG concentrations (500nM) as observed in the scRNA-seq dataset. The *March 12* replicate included unclassified cells (labeled ’other’) that are candidates for FP cells (Fig. A8A-B), but this observation was not reproduced in other experimental replicates (Fig. A8C). We examined the correlations in the *March 12* replicate at 500nM SAG (Fig. A13 G). The PreNeural cluster persists at D6 but displays a different correlation between Nkx6.1 and Sox2 than observed at D4. Given the high SAG concentration, we would not expect this cluster to remain at D6, suggesting it may represent a different cell type. The additional FACs datasets *June5* (Sect. A3.4), are more likely to contain FP cells (Fig. A10).

##### A4.3. MN route and intermediate decision landscape

We observe the following SAG-dependent bifurcations (Fig. A14):

- *0nM SAG*: Only PreNeural and p0/p1 cells are present and both remain stable across time points.
- *10nM SAG*: Olig2 is up-regulated in p2 cells and some take a pMN identity. The prediction that p2 is a shallow is validated by the model in Appendix B Sect. B5.1 (Fig. B7A Panel (4)).
- *High SAG (100nM, 500nM)*: p0/p1 and p2 does not persist; Olig2 is up-regulated and we observe two distinct transitional pathways: in one, a group of cells up-regulates Nkx2.2, co-expressed with Olig2, forming the double-positive (DP) AC. This results in Pax6 down-regulation. In the alternative pathway, Olig2-positive cells retain moderate Pax6 expression without expressing Nkx2.2. These pMN cells differentiate into MNDiff cells and subsequently into motor neurons (MNs). The scRNA-seq dataset showed that these cells belong to a lineage in which Neurog2 is up-regulated. Nkx2.2 is not expressed along this pathway.

Note that the DP AC was also identified in the scRNA-seq dataset and was characterised by the early expression of Nkx2.9/Gm38103 in Olig2 cells, preceding Nkx2.2 expression. This may explain why the cells expressing exclusively Olig2 (pMN) seem to appear before the DP cells in the FACs dataset, although they are more synchronised in the scRNA-seq dataset (Fig. A2G). Additionally, we identified a group of cells transitioning between pMN and DP, labelled DPTrans, which can be distinguished by their gene expression (Fig. A9A) and local correlations between Olig2 and Pax6 (Fig. A9F). These DPTrans cells were also observed in the scRNA-seq data and the temporal progression of this group of cells supports their transitioning status.

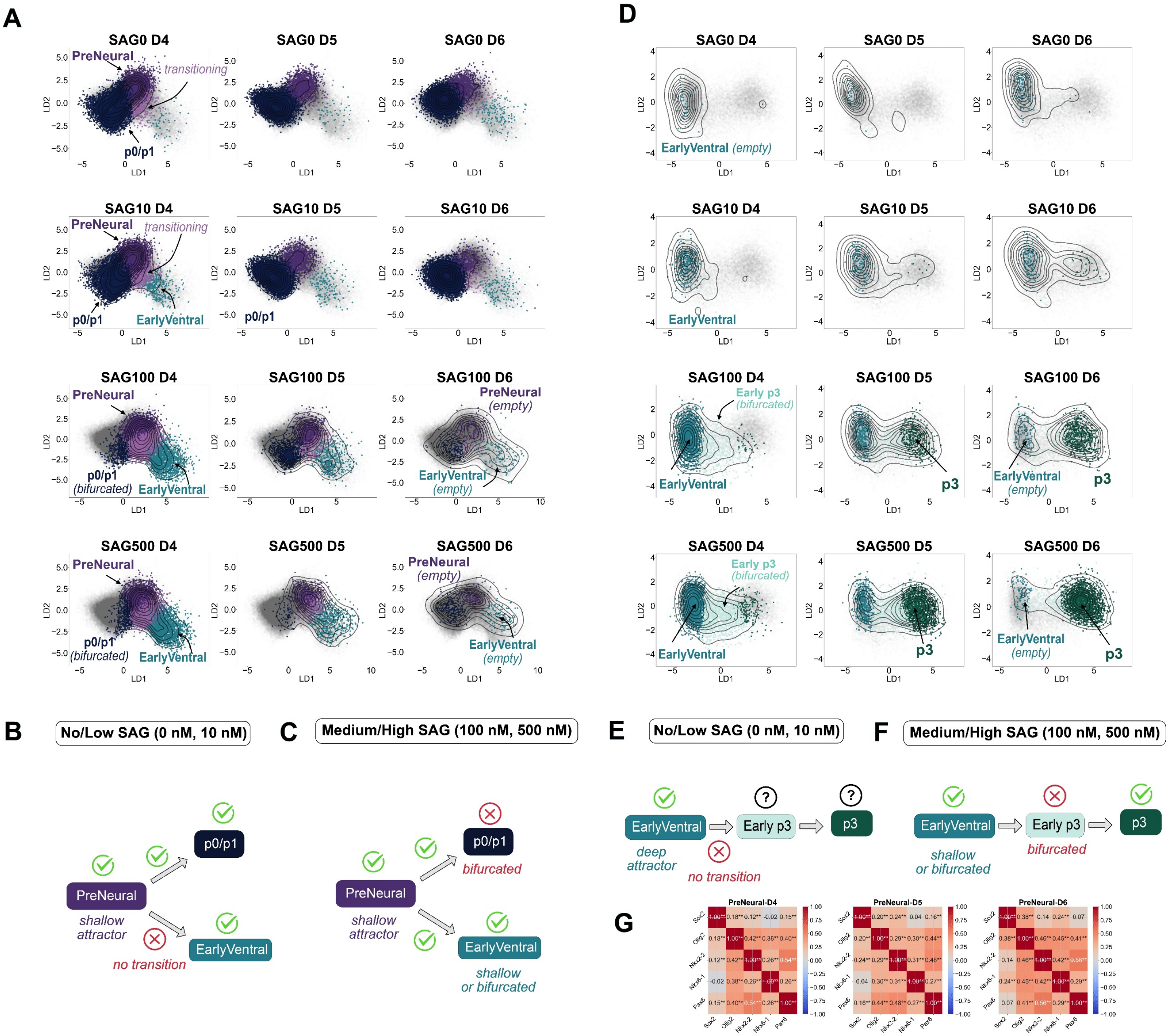

**The PreNeural and Ventral Sub-landscapes in FACs Data**

**Figure A13: A.** The temporal progression for the flip binary decision from PreNeural to p0/p1 or EarlyVentral in LDA coordinates as a function of SAG. The EarlyVentral cluster only appears at higher SAG concentrations (10nM, 100nM, 500nM) while it remains absent at low SAG concentration (0nM). Grey dots represent the cells at all time points, coloured dots indicate cells present in the indicated samples with the colour indicating their cluster assignment, contour lines the density of cells at the indicated time points. **B.** Schematic representation of the PreNeural decision landscape under low SAG exposure. **C.** Similar schematic for high SAG. **D.** Temporal progression of the ACs along the ventral route in each SAG concentration. At SAG 10nM, EarlyVentral persists without cells transitioning to the p3 AC. This supports the idea that EarlyVentral is an AC rather than a transient state. By contrast, Early p3 appears as a transient state in this dataset, which will be later supported by fitting the model. The emergence of p3 cells requires greater than 100nM SAG. Grey dots represent the cells at all time points, coloured dots indicate cells present in the indicated samples with the colour indicating their cluster assignment, contour lines the density of cells at the indicated time points. **E.** Schematic representation of the ventral route under low SAG exposure. **F.** Same as E for high SAG. **G**. Gene-gene correlations between marker expression for the PreNeural AC at 500nM SAG by day. Notably, the correlation between Nkx6.1 and Sox2 (FP markers) shifts between D4 and D6, suggesting that these cells may have changed over time. However, the lack of markers prevents the definitive identification of these cells at D6.

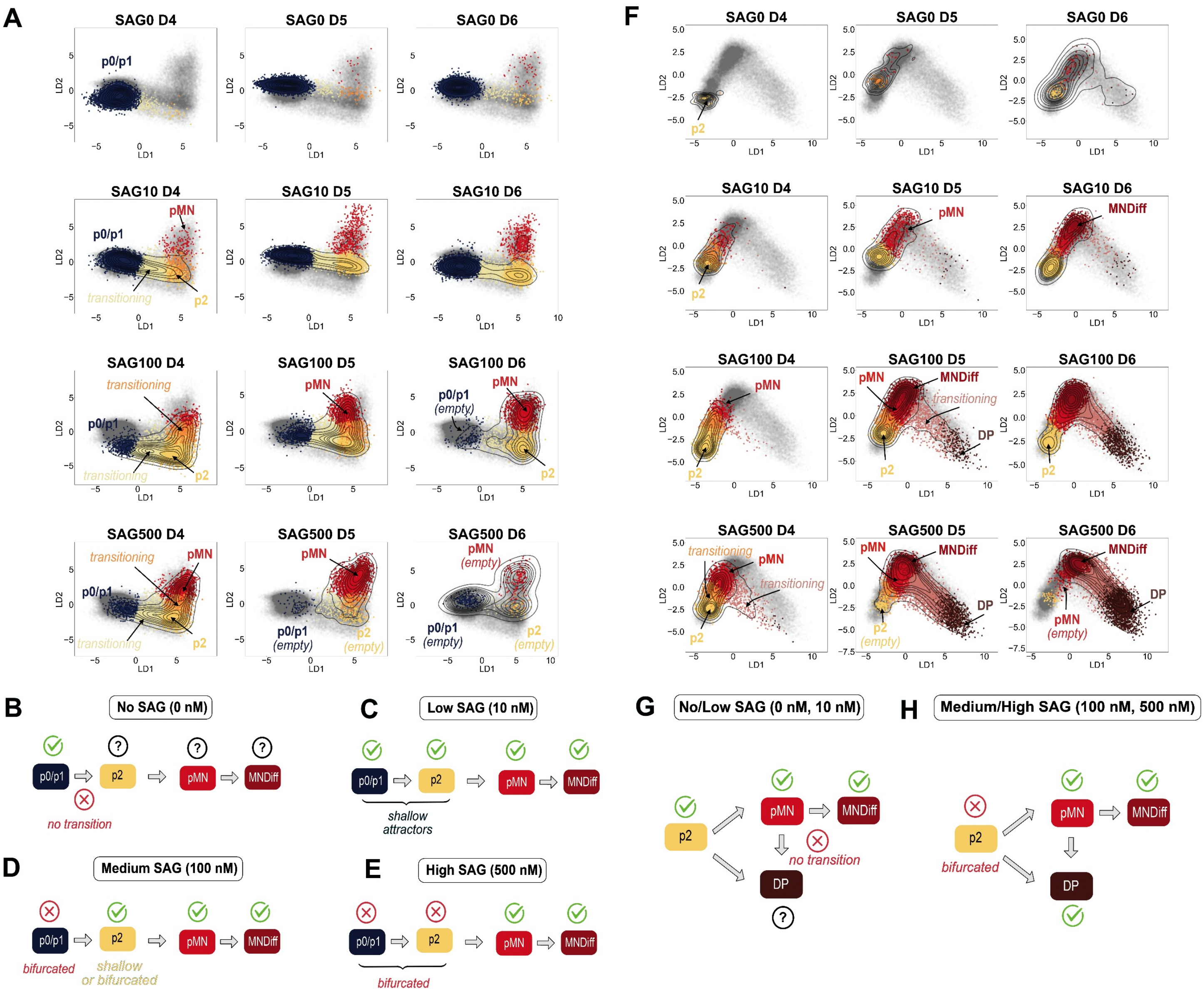

**MN Route and Intermediate Decision Landscape in FACs Data**

**Figure A14: A.** Temporal progression of the ACs along the MN route in LDA coordinates. We observe a linear transition from p0/p1 towards p2 at 10nM SAG and higher concentrations, as well as pMN cells emerging from the p2 cluster. At 0nM SAG, cells remain confined to p0/p1 and never transition to p2 or pMN. This also suggests that the pMN attractor exists across all the conditions we explored but varies in its accessibility. As p2 is always a shallow attractor under SAG exposure, cells can only reach the pMN state if they leave the p0/p1 attractor which requires a concentration greater than 10nM SAG. Grey dots represent the cells at all time points, coloured dots indicate cells present in the indicated samples with the colour indicating their cluster assignment, contour lines the density of cells at the indicated time points. **B.-E.** Schematic representations of the MN route in the indicated SAG concentrations. At 0nM SAG, it is not possible to determine whether the p2, pMN, MNDiff ACs exist as no cells escape from p0/p1. **F.** Temporal progression of the ACs of the intermediate decision landscape in LDA coordinates. The pMN cluster (red) gradually appears when SAG concentration is increased and the DP branch (brown) requires high levels of SAG (100nM, 500nM) to emerge. Grey dots represent the cells at all time points, coloured dots indicate cells present in the indicated samples with the colour indicating their cluster assignment, contour lines the density of cells at the indicated time points. **G-H.** Schematic representations of the intermediate decision landscape. The p2 AC appears stable but shallow at 10nM SAG and bifurcates at higher SAG concentrations.

#### A5. Testing the model on human data

To test whether the network topology and cell fate decisions identified in this study are conserved across species, we analysed an scRNA-seq dataset from human embryonic stem cells[27] in which embryonic tissues such as ventral neural progenitors, mesoderm, and notochord cells were identified. The notochord cells provide a natural source of endogenous SHH signalling, driving the patterning and differentiation of the ventral progenitors (p0/p1, p2, pMN, p3) and the floor plate (FP). As SHH in this case is endogenously produced, cells may hypothetically receive a changing rather than a constant concentration of this signal over time, depending on their position in the tissue[2]. The study of Rito et al.[27] contains data from conditions that produced different proportions of notochord cells, achieved by altering the timing and duration of the transforming growth factor-$\beta$ (TGF$\beta$) signalling, with a 24h delay yielding more notochord cells than an 18h delay. Transcriptome data were collected at three time points D3, D5, and D7 of differentiation for each condition.

##### A5.1. Attractor clusters

To find the ACs we proceeded as in Sect. A1 and applied the outward clustering method starting with 4 markers NKX2.2 (p3), OLIG2 (pMN), SHH (FP) and FOXC1 (Mesoderm).

After filtering out the mitochondrial and ribosomal genes, we selected one-marker samples (including all time points) and selected, for each sample, the top 40 genes with a score greater than 10 obtained from the Scanpy Wilcoxon rank-sum test[34]. This yielded a list of 135 differentially expressed genes (Table A4). We found that, for this dataset, not dividing the samples by time point resulted in better cluster separation. Next, we added to the gene module 20 genes (Table A5) identified from the mouse dataset analysis for comparative analysis. Since these genes were not differentially expressed between the chosen samples, incorporating them before or after performing the Leiden clustering did not make any difference. We performed Leiden clustering by restricting the dataset to this module of genes after PCA reduction (5 PCs). Data were clustered using all time points and both experimental conditions.

The clustering revealed distinct cell types NMP (TBXT, SOX2), Mesoderm (FOXC1), Notochord (TBXT, SHH, NOTO) and the neural progenitors (SOX2). Notochord cells form the SHH signalling centre (Fig. A15A). The neural progenitors exhibited ACs similar to those in the mouse dataset and were evaluated using the criteria of Sect. A1.3 (Fig. A15B-M). The p3 AC co-expressed NKX2.8 (the equivalent of Nkx2.9 for the mouse) and NKX2.2 while in the mouse Nkx2.9 was only expressed in Early p3. FOXP1 is expressed in the ACs that form the ventral sub-landscape with higher expression in FP whereas, in the mouse dataset, it was higher in EarlyVentral.

##### A5.2. Sub-landscapes and gene expression

Due to the presence of several ACs at D3, including NMP and notochord, we could not delineate the order of transitions at this early stage. We focussed our analysis on the transitions involving neural progenitors as there were sufficient time points to capture the developmental progression, starting with cells in the PreNeural state. We therefore extracted the corresponding sub-landscapes identified in the mouse dataset, specifically the *PreNeural sub-landscape* (PreNeural, PreNTrans, p0/p1, EarlyVentral), the *ventral sub-landscape* (EarlyVentral, Early p3, p3, FP), and the *intermediate sub-landscape* (p2, pMN, DP) leaving out MNDiff for better 2D visualisation in LDA coordinates. Using the transition analysis outlined in Sect. A2, we examined gene expression dynamics along the branching routes and compared these patterns with those observed in the mouse dataset (Fig. A16-A17). While several genes exhibited similar expression profiles across both species, some differences were noted. If a gene found in the mouse dataset was missing from the human gene module, we included it for this analysis.

We first compared the two routes leading to p3 (the ventral and intermediate routes) with starting points at the same day (Fig. A16). We observe similar patterns for some genes, for example FOXA2, NOTCH2, HES1 and FOXP4 show higher expression along the ventral route in both species whereas OLIG2, POU3F2 and SOX11 are higher along the intermediate route.

We note that both routes do not intersect at equal angle in the LDA space (a smooth connection for human and non-smooth for mouse). However, the angles are affected by the projection and in effective gene space they are very similar (the subspace defined by the genes modules).

In the PreNeural sub-landscape, the hallmark of the transition is the down-regulation of CDX2 and NKX1.2 as cells exit the PreNeural state. Upregulation of PAX6 and IRX3 guide the cells towards the intermediate route leading to p0/p1 and then p2. By contrast, the ventral route is characterised by the up-regulation of FOXA2 and FOXP1 (Fig. A17A-B). In the ventral sub-landscape, several genes displayed expression patterns similar to those observed in the mouse dataset: NKX2.2 and POU3F2 were up-regulated as cells transitioned to p3, while FOXA2, ARX and FOXP1 were up-regulated as cells reached FP (Fig. A17C-D).

However, NKX2.8 —the human equivalent of mouse Nkx2.9 — was notably up-regulated in the p3 state in human, whereas in the mouse, its expression is transient, appearing briefly in Early p3 before being replaced by Nkx2.2. Additionally, SOX6 did not show significant expression in the human ventral sub-landscape, despite being a clear marker in the mouse analogue.

The remaining transitions are part of the intermediate sub-landscape, with the route followed by cells exiting p2 characterised by the down-regulation of IRX3 and the up-regulation of OLIG1/OLIG2 (Fig. A17E-F). NEUROG2 did not have high expression in pMN but is expressed later in MNDiff, as in the mouse dataset. Similar dynamics were observed with the HES genes, which inhibit neuronal differentiation and maintain cells in a progenitor state[16]. Specifically, HES1, HES4 were up-regulated during the transition to the DP state (HES4 was absent from the mouse dataset). Additionally, RFX4, NKX2.8/Nkx2.9 and NKX2.2 were also up-regulated in the cells transitioning to DP in both datasets.

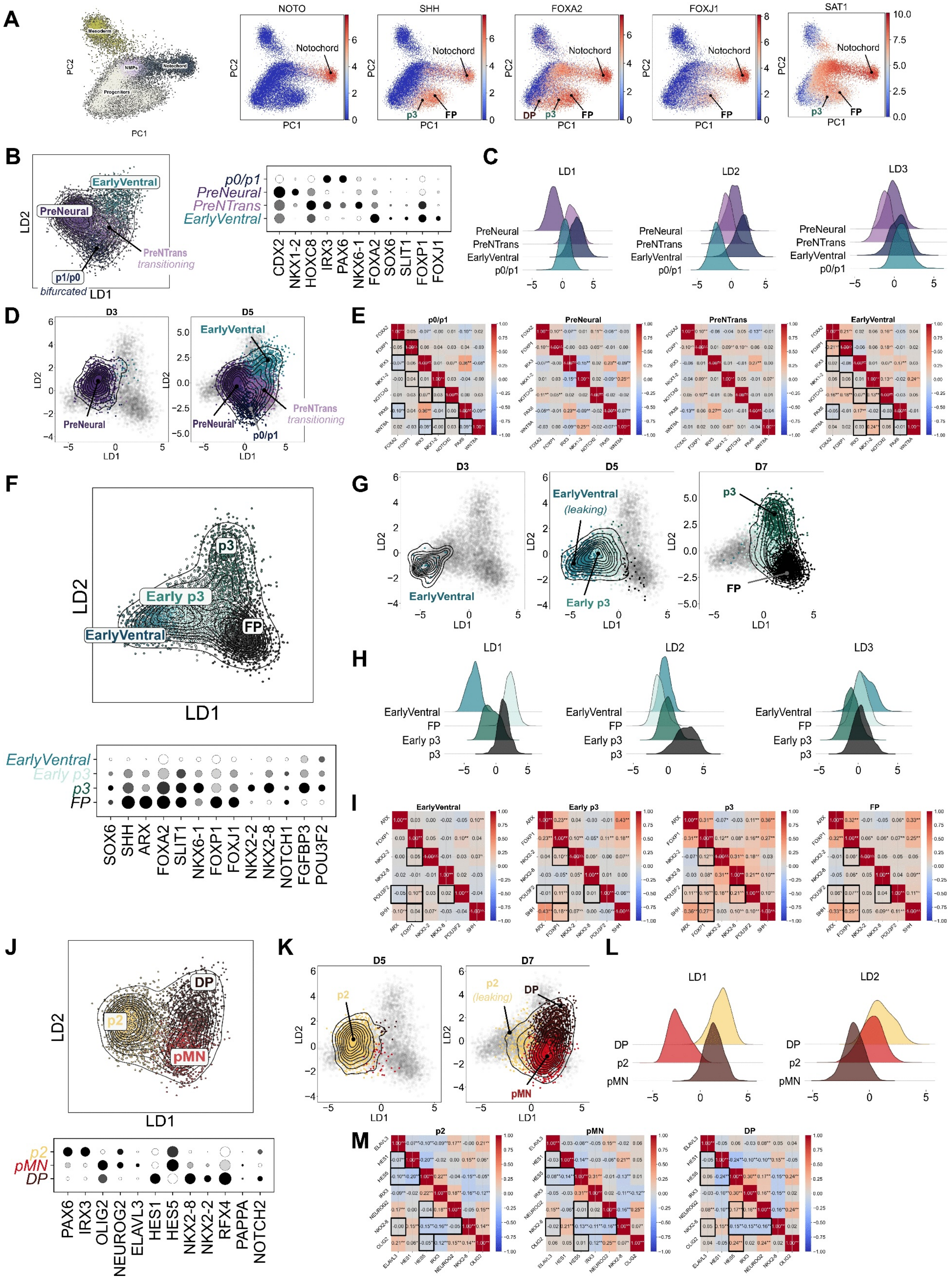

Attractor Clusters in the Human scRNA-seq Data

**Figure A15:** (Caption next page)

**Figure A15:** **A.** Notochord as a signalling center. The notochord expresses ventral markers that diffuse over time along the ventral route and retain high expression in the ventral states p3, FP, DP. **B.** (Left) 2D LDA representation of the PreNeural sub-landscape that includes the ACs PreNeural, p0/p1, EarlyVentral and the transitioning cells PreNTrans. The later expresses a mix of all markers characteristic of the two transition routes as cells exit PreNeural. (Right) Dot plot indicating the expression of the markers used to identify the ACs (D3-D5). **C.** Unimodal distributions of the LDA scores within each AC of the PreNeural sub-landscape. **D.** Temporal progression of the ACs of the PreNeural sub-landscape (no cells at D7). As for the mouse dataset (Fig. A1G), we identified a flip binary decision from PreNeural to either p0/p1 or EarlyVentral. **E.** Gene-gene correlations within each AC for all time points showing the same genes as for the mouse dataset (Fig. A1I). Statistical significance is given by a p-value $< 0.01$ (**) and a p-value between $0.01$ and $0.05$ (*). Distinct correlations for a pair of genes between two ACs suggest a difference in the regulatory interactions. We marked the same gene pairs as those for the mouse dataset in Fig. A1I. While some pairings still mark a distinction between the ACs p0/p1 and EarlyVentral (like Pax6-Foxa2 or Foxp1-Foxa2), others do not (e.g. Irx3-Notch2). **F.** (Top) 2D LDA representation of the ventral sub-landscape that includes the ACs EarlyVentral, Early p3, p3 and FP. (Bottom) Dot plot indicating the expression of the markers used to identify the ACs (D3-D7). **G.** Temporal progression of the ACs of the ventral sub-landscape. As for the mouse dataset (Fig. A2B) we observe a direct transition from EarlyVentral to Early p3, followed by a binary decision from Early p3 to either p3 or FP. **H.** Unimodal distributions of the LDA scores within each AC. As for the mouse dataset, the first 2 LDs are sufficient to separate the ACs. **I.** Gene-gene correlations within each AC for all time points showing the same genes as for the mouse dataset (Fig. A2D). **J.** (Top) 2D LDA representation of the intermediate sub-landscape that includes the ACs p2, pMN and DP (MNDiff not shown for figure clarity). (Bottom) Dot plot indicating the expression of the markers used to identify the ACs (we use the same markers as for the mouse dataset). **K.** Temporal progression of the ACs of the intermediate sub-landscape (no cells at D3). The figure suggests a binary decision from p2 to either pMN or DP and also indicates that pMN cells can transition to DP. **L.** Unimodal distributions of the LDA scores within each AC. **M.** Gene-gene correlations within each AC for all time points showing the same genes as for the mouse dataset (Fig. A2I).

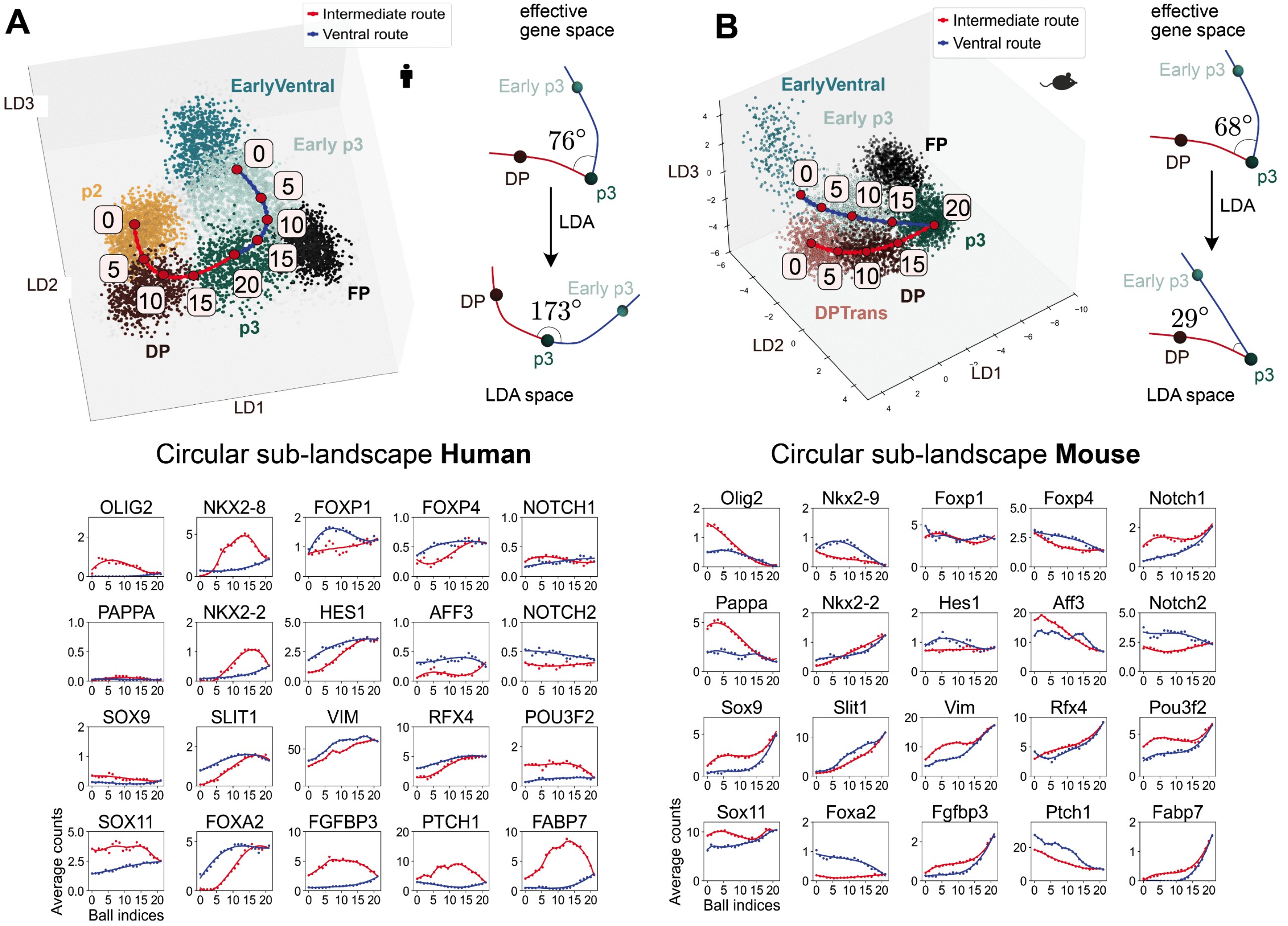

**Circle Sub-landscape and Transitions Analysis Comparison between Human and Mouse**

**Figure A16:** **A.** Circular sub-landscape transition analysis. Ventral route (blue) starts at Early p3 at D5 and the intermediate route starts at p2 at D5. The two routes join smoothly (horizontal tangents) in LDA space but not in effective gene space. We observe similar patterns to B, for example FOXA2, NOTCH2, HES1 and FOXP4 show higher expression along the ventral route in both species whereas OLIG2, POU3F2 and SOX11 are higher along the intermediate route. **B.** Same as A for the mouse dataset. The starting points (balls of index $0$) are Early p3 at D5 (ventral route) and DPTrans at D5 (intermediate route). We computed the angle at the junction of the two routes in effective gene space and found a similar angle than for the mouse.

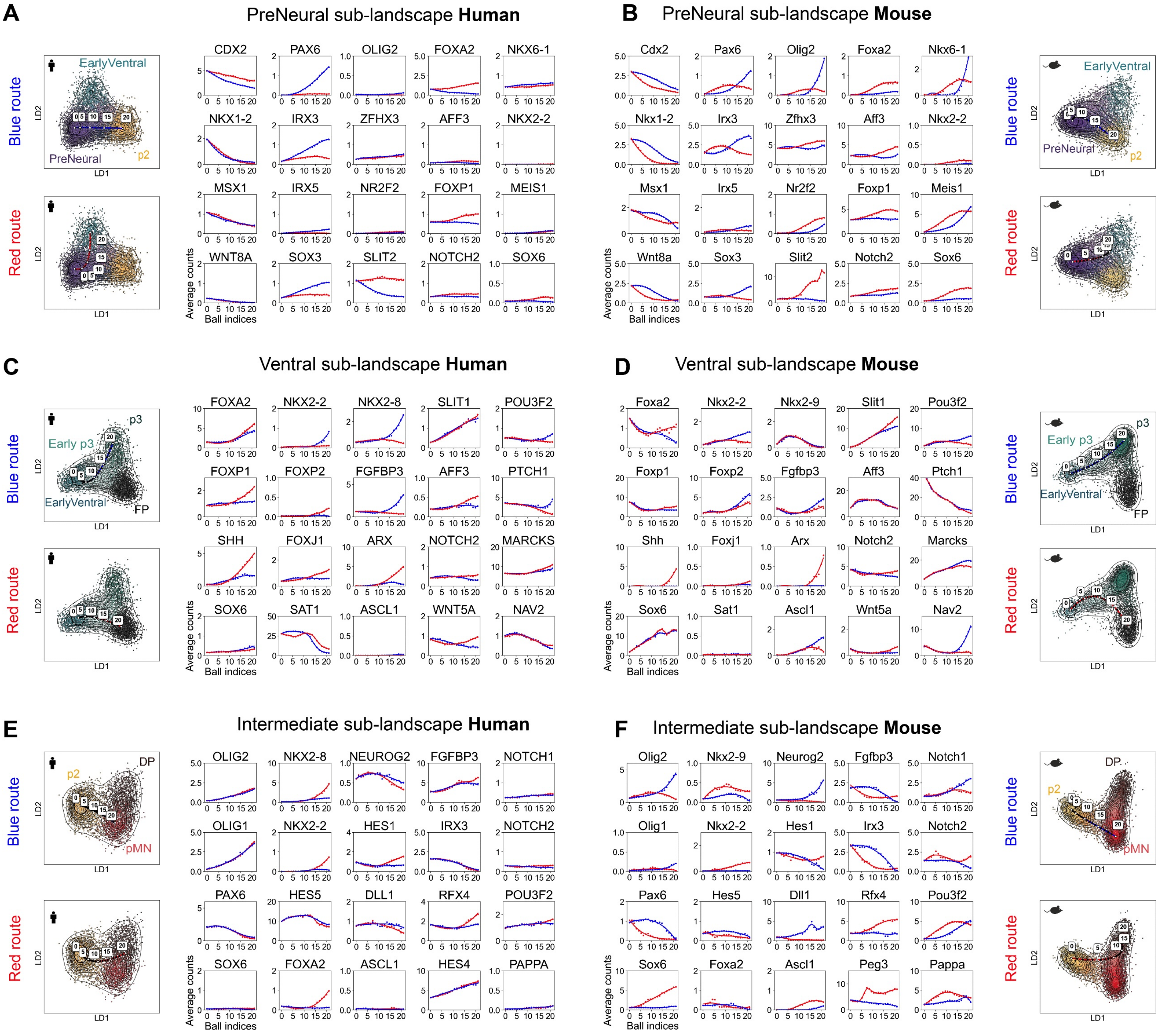

**Sub-landscapes and Transitions Analysis Comparison between Human and Mouse**

**Figure A17: A-F.** Transition analysis for the main sub-landscapes PreNeural, ventral and intermediate from the Human scRNA-seq dataset (left column) compared to the Mouse scRNA-seq dataset (right column). All sub-landscapes contain a binary flip decision and are visualised in 2D LDA space. Unstable manifolds were estimated as detailed in Sect. A2, and transitions of key genes were analysed along these, using the same gene set as in the mouse dataset, and their corresponding human homologues for cross-species comparison. **A.** PreNeural sub-landscape (human). Similarly to the mouse dataset in B, genes CDX2, NKX1.2, MSX1 down-regulate as cells exit PreNeural. Genes characteristic of the intermediate route (blue) such as PAX6, IRX3 AND SOX3 get up-regulated as cells transition to p0/p1 and p2. Marker genes such as FOXA2, FOXP1 and SLIT2 up-regulate along the ventral route (red). **B.** PreNeural sub-landscape (mouse). **C.** Ventral sub-landscape (human). The route to p3 (blue) is characterised by up-regulation of genes such as NKX2.2 and FGFBP3 as in the mouse dataset in D. NKX2.8, the human homologue of mouse Nkx2.9, also shows an increase along this route, in contrast to the mouse, were expression was restricted to Early p3. We did not observe an increase in Ascl1 expression as cells transitioned to p3, unlike what was seen in the mouse in D. Markers such as SHH, FOXA2, ARX and FOXP1 characterise the route to FP (red). **D.** Ventral sub-landscape (mouse). **E.** Intermediate sub-landscape (human). We excluded MNDiff and p3 for better 2D visualisation, otherwise three LDs are required to separate the clusters (Fig. A6B). The p2 exit is demarcated by rising expression of pMN markers OLIG1, OLIG2 and decrease of IRX3 expression. NEUROG2 increases in cells transitioning to pMN but the trend is less pronounced than for the mouse in F. Up-regulation of NKX2.8, NKX2.2, RFX4 and HES1 marks the route to DP (red), consistent with observations in mouse. In this dataset FOXA2 is expressed in all ventral states, including DP. PEG3 was not available in this dataset. Instead, we highlight HES4, whose up-regulation in both pMN and DP coincides with the down-regulation of HES5 and IRX3. **F.** Intermediate sub-landscape (mouse). Hes4 was not available in the dataset so we show Peg3 instead which mark the route to DP.

### A. Appendix: Gene modules

| $G_{early}$ | | | | | | | |
|---|---|---|---|---|---|---|---|
| 9030622O22Rik | Actn2 | Adgrl2 | Adgrv1 | Ankrd11 | Ankrd28 | Anp32e | B430203G13Rik |
| B4galnt1 | Braf | Calr | Ccnd2 | Cdh1 | Cdh6 | Cdkn1a | Celsr1 |
| Cfap100 | Chaf1a | Chka | Colec12 | Cracd | Cul1 | Dgkh | Dll1 |
| Dok6 | Dst | Efnb2 | Eif4a3 | Epcam | Fgd4 | Fgfbp3 | Fn1 |
| Foxa2 | Foxc1 | Foxc2 | Foxo3 | Frem1 | Frem2 | Gli2 | Gm13456 |
| Gm6477 | Gnai3 | Gspt1 | Gtf2a1 | Hipk2 | Hmgcr | Hoxaas3 | Hoxb5os |
| Hoxb9 | Hspa5 | Ier5l | Il17rd | Ildr2 | Irx3 | Itga6 | Jarid2 |
| Kalrn | Kdm6b | Kif26b | Lama1 | Lamb1 | Lef1 | Limch1 | Lrig3 |
| Lyar | Map1b | Mast2 | Mat2a | Mcc | Mcm4 | Meg3 | Msi2 |
| Nid2 | Nin | Nkx6-1 | Npm1 | Nrg2 | Olig2 | Pak1 | Pax6 |
| Pcdh19 | Pcdh7 | Pde4d | Pdzrn3 | Pgm2 | Pid1 | Pla2g7 | Ppp1r9a |
| Prickle1 | Psmd2 | Ptch1 | Ptk7 | Ptp4a2 | Ptpn2 | Pum1 | Rftn1 |
| Rfx4 | Rnf213 | Rnf220 | Robo1 | Rpl18 | Rspo3 | Rtf1 | Sall2 |
| Sema3e | Sema6a | Sgip1 | Slc2a1 | Slc38a2 | Smoc1 | Sox2 | Spred2 |
| Stam2 | Stat5a | Sulf1 | T | Tarsl2 | Tcerg1 | Tcf20 | Thsd4 |
| Tns3 | Tut4 | U2surp | Utrn | Vps13c | Ywhaz | Zc3h7a | Zdbf2 |
| Zfp462 | | | | | | | |

**Table A2:** List of $108$ genes contained in the gene module $G_{early}$ used to perform Leiden clustering of $X_{early}$ (D3–D4) of the mouse scRNA-seq dataset. Obtained by fixing the threshold of significance $\alpha = 99.95$.

$G_{late}$

| | | | | | | | |
|---|---|---|---|---|---|---|---|
| 9030622O22Rik | Adgrv1 | Aff3 | Ankrd28 | Baz2b | Btbd17 | Ccnd2 | Cdc42bpa |
| Cdh6 | Chd3 | Chd7 | Col18a1 | Col4a1 | Cpeb4 | Cracd | Crim1 |
| Ctnnd2 | Cux1 | Dach2 | Dchs2 | Dpysl3 | Ednrb | Elavl3 | Elavl4 |
| Epb41l4b | Fat3 | Fbxl7 | Fndc3b | Foxa2 | Foxn3 | Frmd4a | Glra1 |
| Gm32061 | Gm38103 | Gpc3 | Greb1l | Heg1 | Hmga2 | Hoxb4 | Igsf9b |
| Iqgap2 | Kalrn | Kank1 | Kazn | Kbtbd11 | Kif21a | Lgr4 | Map1b |
| Mir100hg | Mir99ahg | Myh9 | Myo16 | Nectin3 | Neurod4 | Neurog2 | Nkx2-2 |
| Notch2 | Npas3 | Nrp2 | Ntn1 | Olig2 | Pde4d | Peg3 | Phactr2 |
| Plekha7 | Pou2f2 | Pou3f2 | Pou3f3 | Ppp2r2b | Prmt8 | Ptch1 | Ptprn2 |
| Ptpru | Rcor2 | Rfx4 | Rgma | Rhbdl3 | Rnf165 | Robo2 | Ror2 |
| Rtn4 | Sdk2 | Setbp1 | Shh | Slc2a1 | Slit1 | Slit2 | Slit3 |
| Soga3 | Sox11 | Sox5 | Sox6 | Spag9 | Spon1 | Spsb4 | Srrm4 |
| Ssbp3 | Stxbp6 | Sulf1 | Susd6 | Tbc1d9 | Tcf7l1 | Tenm3 | Thsd7a |
| Tns3 | Tox2 | Trabd2b | Trp53i11 | Tspan5 | Ttc28 | Zfp503 | Zswim6 |

**Table A3:** List of 112 genes contained in the gene module $G_{late}$ used to perform Leiden clustering of $X_{late}$ (D5–D8) of the mouse scRNA-seq dataset. Obtained by fixing the threshold of significance $\alpha = 99.9$.

| $G_{human}$ | | | | | | | |
|---|---|---|---|---|---|---|---|
| AL359091.1 | AMOT | ANXA2 | APLNR | BASP1 | BTG1 | C4orf48 | C5orf49 |
| CALM1 | CAMK2N1 | CCND1 | CD63 | CD99 | CDH11 | CETN2 | CKB |
| CMTM8 | COL18A1 | COL1A2 | COL2A1 | COL3A1 | COL4A1 | COL4A2 | COL5A2 |
| COTL1 | DLK1 | DNALI1 | DPYSL5 | EFCAB1 | EFHC1 | ENKUR | FABP7 |
| FAT3 | FBLN1 | FBN2 | FGFBP3 | FOXA2 | FOXC1 | FOXC2 | FOXD1 |
| FOXJ1 | FZD3 | FZD7 | GMDS | GPC3 | GPM6B | GYPC | H3F3A |
| HAS2 | HES4 | HES5 | ID2 | IFT22 | IFT57 | IGDCC3 | IGFBP5 |
| ITM2C | JAM2 | KCNG1 | KIF9 | KRT18 | KRT8 | LANCL2 | LAPTM4A |
| LRP2 | LY6E | MAGED1 | MALAT1 | MAML2 | MAP1B | MARCKS | MEOX1 |
| MGST3 | MLF1 | MLLT1 | MORN2 | MXRA8 | NAP1L1 | NES | NKX2-2 |
| NKX2-8 | NKX3-2 | NKX6-1 | NKX6-2 | NR2F1 | NR2F2 | NTRK2 | OLIG1 |
| OLIG2 | PCOLCE | PDGFRA | PGF | PIFO | PLEKHA5 | PLTP | POU3F2 |
| PPIL6 | PRTG | PTCH1 | PTMA | RBMS1 | RBP1 | RFX4 | RGMB |
| RSPH4A | S100A11 | SAT1 | SEPT11 | SEPT6 | SERPINF1 | SFRP1 | SHH |
| SLC2A1 | SLIT2 | SOX11 | SOX2 | SOX3 | SOX9 | SPAG1 | SPARCL1 |
| TAGLN2 | TCF15 | TFDP2 | TFF3 | TLE4 | TMEM97 | TPM2 | TPPP3 |
| TSC22D1 | TTYH1 | TUBA1A | TUBA1B | TUBB2B | TWIST1 | VIM | |

**Table A4:** List of $135$ genes contained in the gene module used to perform Leiden clustering of the scRNA-seq human dataset (D3,D5,D7). These are obtained by taking the first $40$ differentially expressed genes with score higher than $10$ for each sample.

| | | | | | | | |
|---|---|---|---|---|---|---|---|
| SOX6 | SLIT1 | NEUROG2 | SPON1 | PAPPA | PAX6 | NKX1-2 | IRX3 |
| SOX1 | FGF8 | CDX2 | ALDH1A2 | TBXT | LEF1 | TBX6 | SOX17 |
| TUBB3 | ELAVL3 | SIM1 | HES1 | | | | |

**Table A5:** Genes missing from Table A4 added to the gene module for clustering.

#### References

[1] T. A. Alexander, R. A. Irizarry, and H. C. Bravo. Capturing discrete latent structures: choose lds over pcs. *Biostatistics*, 24(1):1–16, 2022. doi: 10.1093/biostatistics/kxab030.

[2] N. Balaskas, G. R. Azevedo, J. K. Panov, E. Marti, J. Briscoe, and K. R. M. Anderson. Gene regulatory logic for reading the sonic hedgehog signaling gradient in the vertebrate neural tube. *Cell*, 148(1-2):273–284, 2012. doi: 10.1016/j.cell.2011.10.047.

[3] P. N. Belhumeur, J. P. Hespanha, and D. J. Kriegman. Eigenfaces vs. fisherfaces: Recognition using class specific linear projection. *IEEE Transactions on Pattern Analysis and Machine Intelligence*, 19(7):711–720, 1997. doi: 10.1109/34.598228.

[4] A. Butler, P. Hoffman, P. Smibert, E. Papalexi, and R. Satija. Integrating single-cell transcriptomic data across different conditions, technologies, and species. *Nature Biotechnology*, 36:411–420, 2018. doi: 10.1038/nbt.4096.

[5] T. M. Cover. *Elements of information theory*. John Wiley & Sons, 1999.

[6] I. Csiszár. On information-type measure of difference of probability distributions and indirect observations. *Studia Sci. Math. Hungar.*, 2:299–318, 1967.

[7] M. J. Delás, C. M. Kalaitzis, T. Fawzi, M. Demuth, I. Zhang, H. T. Stuart, E. Costantini, K. Ivanovitch, E. M. Tanaka, and J. Briscoe. Developmental cell fate choice in neural tube progenitors employs two distinct cis-regulatory strategies. *Developmental Cell*, 58(1):3–17.e8, 2023.

[8] E. Dessaud, L. Yang, K. Hill, B. Cox, F. Ulloa, A. Ribeiro, A. Mynett, B. G. Novitch, and J. Briscoe. Interpretation of the sonic hedgehog morphogen gradient by a temporal adaptation mechanism. *Nature*, 450 (7170):717–720, 2007. doi: 10.1038/nature06347.

[9] K. Fukunaga. *Introduction to Statistical Pattern Recognition*. Academic Press, Boston, 2 edition, 1990. ISBN 0122698517.

[10] C. Gambella, B. Ghaddar, and J. Naoum-Sawaya. Optimization problems for machine learning: A survey. *European Journal of Operational Research*, 290(3):807–828, 2021. doi: 10.1016/j.ejor.2020.08.045.

[11] M. Gouti, A. Tsakiridis, F. J. Wymeersch, Y. Huang, J. Kleinjung, V. Wilson, and J. Briscoe. In vitro generation of neuromesodermal progenitors reveals distinct roles for wnt signalling in the specification of spinal cord and paraxial mesoderm identity. *PLoS Biology*, 12(8): e1001937, 2014.

[12] M. Gouti, J. Delile, D. Stamataki, F. J. Wymeersch, Y. Huang, J. Kleinjung, V. Wilson, and J. Briscoe. A gene regulatory network balances neural and mesoderm specification during vertebrate trunk development. *Developmental Cell*, 41(3):243–261, 2017.

[13] C. Hafemeister and R. Satija. Normalization and variance stabilization of single-cell rna-seq data using regularized negative binomial regression. *Genome Biology*, 20 (1):296, 2019. doi: 10.1186/s13059-019-1874-1.

[14] M. Jacobsen. Homogeneous gaussian diffusions in finite dimensions. Preprint No. 3, Institute of Mathematical Statistics, University of Copenhagen, 1991.

[15] A. Javali, A. Misra, K. Leonavicius, D. Acharyya, B. Vyas, and R. Sambasivan. Co-expression of tbx6 and sox2 identifies a novel transient neuromesoderm progenitor cell state. *Development*, 144(24):4522–4529, 2017.

[16] R. Kageyama, T. Ohtsuka, and T. Kobayashi. Roles of hes genes in neural development. *Development, Growth & Differentiation*, 50(s1):S97–S103, 2008. doi: 10.1111/j.1440-169X.2008.00993.x. URL `https://pubmed.ncbi.nlm.nih.gov/18430159/`.

[17] T. G. Kurtz. *Approximation of Population Processes*. Society for Industrial and Applied Mathematics, 1981. doi: 10.1137/1.9781611970333.

[18] M. I. Love, W. Huber, and S. Anders. Moderated estimation of fold change and dispersion for RNA-seq data with DESeq2. *Genome Biology*, 15(12):550, 2014. doi: 10.1186/s13059-014-0550-8.

[19] M. D. Luecken and F. J. Theis. Current best practices in single-cell rna-seq analysis: a tutorial. *Molecular Systems Biology*, 15(6):e8746, 2019. doi: 10.15252/msb.20188746.

[20] R. J. Maizels, D. M. Snell, and J. Briscoe. Reconstructing developmental trajectories using latent dynamical systems and time-resolved transcriptomics. *Cell Systems*, 15(5):411–424.e9, 2024.

[21] *MATLAB Statistics and Machine Learning Toolbox*. The MathWorks, Inc., Natick, Massachusetts, 2023. URL `https://www.mathworks.com/help/stats/fitgmdist.html`.

[22] L. McInnes, J. Healy, N. Saul, and L. G. berger. Umap: Uniform manifold approximation and projection. *Journal of Open Source Software*, 3(29):861, 2018. doi: 10.21105/joss.00861. URL `https://doi.org/10.21105/joss.00861`.

[23] G. McLachlan, S. Ng, and D. Peel. On clustering by mixture models. In *Exploratory Data Analysis in Empirical Research: Proceedings of the 25 th Annual Conference of the Gesellschaft für Klassifikation eV, University of Munich, March 14–16, 2001*, pages 141–148. Springer, 2003.

[24] M. Mojtahedi, A. Skupin, J. Zhou, I. G. Castaño, R. Y. Y. Leong-Quong, H. Chang, K. Trachana, A. Giuliani, and S. Huang. Cell fate decision as high-dimensional critical state transition. *PLOS Biology*, 14 (12):e2000640, 2016. doi: 10.1371/journal.pbio.2000640.

[25] B. G. Novitch, A. I. Chen, and T. M. Jessell. Coordinate regulation of motor neuron subtype identity and pan-neuronal properties by the bHLH repressor Olig2. *Neuron*, 31(5):773–789, 2001. doi: 10.1016/s0896-6273(01)00407-x.

[26] F. Pedregosa, G. Varoquaux, A. Gramfort, V. Michel, B. Thirion, O. Grisel, M. Blondel, P. Prettenhofer, R. Weiss, V. Dubourg, J. Vanderplas, A. Passos, D. Cournapeau, M. Brucher, M. Perrot, and E. Duchesnay. Scikit-learn: Machine learning in Python. *Journal of Machine Learning Research*, 12:2825–2830, 2011.

[27] T. Rito, A. R. G. Libby, M. Demuth, M.-C. Domart, J. Cornwall-Scoones, and J. Briscoe. Timely tgf$\beta$ signalling inhibition induces notochord formation from human pluripotent stem cells. *Nature*, 637, 2025.

[28] M. Sáez, R. Blassberg, E. Camacho-Aguilar, E. D. Siggia, D. A. Rand, and J. Briscoe. Statistically derived geometrical landscapes capture principles of decision-making dynamics during cell fate transitions. *Cell Systems*, 2022.

[29] H. Shimojo, T. Ohtsuka, and R. Kageyama. Oscillations in notch signaling regulate maintenance of neural progenitors. *Neuron*, 58(1):52–64, 2008. doi: 10.1016/j.neuron.2008.02.014.

[30] T. Takemoto, M. Uchikawa, M. Yoshida, D. M. Bell, R. Lovell-Badge, V. E. Papaioannou, and H. Kondoh. Tbx6-dependent sox2 regulation determines neural or mesodermal fate in axial stem cells. *Nature*, 470(7334):394–398, 2011.

[31] V. A. Traag, L. Waltman, and N. J. van Eck. From Louvain to Leiden: guaranteeing well-connected communities. *Scientific Reports*, 9(1):5233, 2019.

[32] L. van der Maaten and G. Hinton. Visualizing data using t-sne. *Journal of Machine Learning Research*, 9(86):2579–2605, 2008.

[33] Y. Wang, H. Huang, C. Rudin, and Y. Shaposhnik. Understanding how dimension reduction tools work: An empirical approach to deciphering t-sne, umap, trimap, and pacmap for data visualization. *Journal of Machine Learning Research*, 22(201):1–73, 2021.

[34] F. A. Wolf, P. Angerer, and F. J. Theis. Scanpy: large-scale single-cell gene expression data analysis. *Genome Biology*, 19(1):15, 2018. doi: 10.1186/s13059-017-1382-0.
