## Appendix B for "Dynamic Landscape Analysis of Cell Fate Decisions: Predictive Models of Neural Development From Single-Cell Data"

#### Modelling & estimation

**Overview:** This appendix provides comprehensive mathematical details for the dynamical landscape modelling approach presented in the main text. Section B1 introduces the theoretical foundation linking gene regulatory networks to dynamical landscapes and binary flip bifurcations. Section B2 describes the construction of parameterised sub-landscape models using catastrophe theory normal forms. Section B3 details the simulation methodology and noise implementation. Section B4 describes the general ABC fitting procedure for the two first binary decisions. Section B5 presents parameter estimation procedures using FACs data. Section B6 and B8 demonstrate model predictions for SAG pulse experiments. Section B7 validates the model against mouse scRNA-seq data across development. Section B9 applies the framework to human data for cross-species validation.

**Connection to Main Paper:** The theoretical framework explains how Shh signaling remodels the decision landscape through bifurcations (**Main Fig. 5-7**), and the fitted parameters enable quantitative predictions for dynamic signaling perturbations (**Main Fig. 8-10**). Human data validation (**Main Fig. 11**) demonstrates cross-species conservation of the landscape topology and quantifies how notochord-derived Shh affects cell decision making.

### Part II
### Modelling & estimation

#### B1. Dynamical landscapes and GRNs

An underlying hypothesis of our approach is that cellular decision making is due to a gene regulatory network i.e. a stochastic dynamical system involving the production, modification, interaction, and decay of the mRNAs and proteins of a set of genes[8]. This dynamical system is subject to signals that modify the parameters and rates associated with these processes and so can cause equilibria of the dynamical system to bifurcate and change its behaviour.

In the context of cell decision making, the attractors of this system play a special role in that they correspond to stable cell states[9,10]. However, this idea requires discussion since most of these cell states are transitory, a fact that seems in opposition to the notion of an attractor in a deterministic dynamical system because, in such a system, the state cannot escape an attractor. In our approach this is not contradictory because cells can escape an attractor by any of the following processes.

(i) Since the dynamics are stochastic, a cell can escape an attractor because of a stochastic fluctuation. This is more likely if the basin of the attractor is small/shallow meaning that the attractor is close to undergoing a bifurcation by colliding with a nearby index-1 saddle (called a *saddle-node* or *fold bifurcation*).

(ii) A change in a signal can cause an attractor to either (a) bifurcate and disappear so that all cells escape its hold or (b) become small/shallow, as in (i), so that the cells escape more readily.

(iii) On arriving at an attractor, cells change the expression of a gene(s) that acts on the dynamical system in a similar fashion to the signal in (ii) so as to destabilise the attractor and allow the cell to move on to the next state.

We can consider that there is a global GRN comprising all the genes involved in the decision-making producing the set of cell types that we consider. However, it is likely that only a subset of these are active in the decision making and the transitions to and/or from any cell state. Therefore, when considering any particular transition or attractor, it is reasonable to focus on the active GRN by identifying that part of the global GRN that is active in this local process. In this context, genes that are differentially expressed between this attractor and others, and genes that change along the transition paths into and out of this attractor, are of special interest as candidates for the GRN. A particular advantage of this approach arises from this observation because once one has candidate *Attractor Clusters* (ACs), which are groups of cells representing stable cell states, one can investigate the genes that are differentially expressed between the cells in different

ACs and the genes that change their expression when they transition between ACs.

##### B1.1. The GRN is associated with individual cells

In this manuscript, the active GRN belongs to the single cell[4,21,26], and not the set of all cells in the same cell state. Thus, for example, two cells currently in the same cell state, might have slight differences in their molecular state or the signals they see. That means that the parameters governing the GRNs are not identical and their exit from the attractor might follow different escape routes. An example of this is two cells in a cell state that is at the head of a binary flip that make different choices between the two routes associated with the binary flip (Fig. B1A).

##### B1.2. Dynamical Landscapes

The parameterised dynamical system determined by the GRN and the signals it receives has associated with it a well defined and parameter/signal dependent geometric structure determined by the attractors and the saddles with 1-dimensional unstable manifolds joining pairs of attractors. The latter define the transition paths followed by cells when they move between these attractors. This structure we refer to as a *dynamical landscape.* By a *sub-landscape* in the dynamical landscape we mean a subset of neighbouring attractors together with the connecting 1-dimensional unstable manifolds. A sub-landscape in the data is a group of neighbouring clusters from which some clusters correspond to stable cell states (ACs) and other to cells transitioning between stables cell states that we hypothesise are positioned close, in gene expression space, to an unstable manifold. When we talk about a *route* we mean such a transition route and when we discuss a *decision or cell decision* we mean the branching that happens when, due to variation in the signals, we observe more than one transition route out of an AC that links to other ACs.

It is important that in a dynamical system, for fixed parameters, the transition route out of an attractor must go to another specific attractor. This means that in our picture when the signals are fixed the cells transitioning from an AC will, with a certain probability, go to a specific AC. This probability approaches 1 as stochasticity reduces. Without this property there would be no certainty that cells in a given state that start to transition would go to one of a small number of well-defined states.

##### B1.3. The binary flip sub-landscape

The binary flip sub-landscape (Fig. B1A-B) contains three attractors and two saddles. There are simple 3-gene regulatory network models that produce it[5,15]. In the developmental context one of the attractors (termed the *head* $H$) is distinguished in that it will be the attractor at which cells first arrive. The other two attractors (labelled $A$ and $B$) are in a bistable pair together with a saddle $s_{AB}$ joined by an unstable manifold. The head attractor $H$ is joined to one of $A$ and $B$ by the unstable manifold of the other saddle $s_H$. Associated with such a landscape is the heteroclinic flip bifurcation where the unstable manifold of the saddle $s_H$ changes its connection from $A$ to $B$ (Fig. B1A). The presence of a head attractor is important as this is the way the sub-landscape captures the output from the ACs supplying it with cells.

The landscape depends essentially on two-parameters because we combine the curve of saddle-nodes that destroy the head attractor (*fold curve*) with the curve of homoclinic connections (*flip curve*) where the unstable manifold of the saddle $s_H$ hits the saddle $s_{AB}$ (Fig. B1C). In this way we obtain a normal form for the binary flip. There is a special point (*fold-flip point*) in parameter space where the flip curve ends on the curve of saddle-nodes.

In the data presented in Appendix A, we see cells following both routes and filling both attractors $A$ and $B$. There are three mechanisms by which this can occur:

1. *close pass to a forward saddle* $s_{AB}$: although cells are effectively identical, the unstable manifold passes close to the saddle and stochasticity during this transition results in some cells reaching one attractor and some cells the other (Fig. B1C).
2. *cellular heterogeneity*: the cells or the signals they are responding to are heterogeneous and hence have slightly different landscapes.
3. *feedback*: there is some feedback that alters the route of the unstable manifold. For example, cells might fill one attractor at first and in doing so turn on a feedback signal that causes the unstable manifold to flip to the other target[14] (Fig. B1D).

##### B1.4. Binary choice decisions

A *binary choice decision*[15] arises when an attractor $H$ bifurcates through one or both of two distinct saddles $s_A, s_B$ (Fig. B1E). The simultaneous bifurcation through both saddles is known as a dual cusp bifurcation and is less common than the close pass to a forward saddle described above because dual cusps are singularities of codimension 2. While a configuration of an attractor being close to two distinct saddles is typically rarer, cells taking distinct routes in this scenario is more common under intrinsic cellular heterogeneity. In this context, different cells may experience their attractor as being closer to one saddle or the other, opening two distinct transition paths.

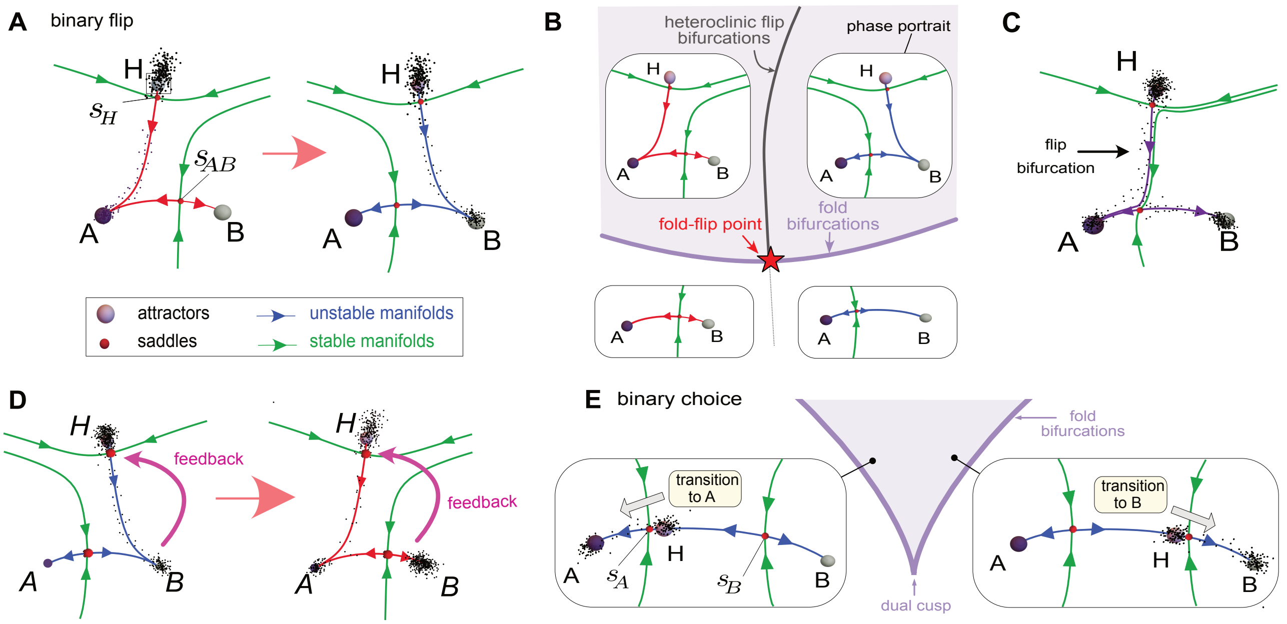

**Binary Flip Landscape Model of Cell Decisions**

**Figure B1:** **A.** Binary flip landscape. Cells start in the progenitor state $H$ which, in the figure, corresponds to a shallow basin. Noise is sufficient for cells to transition into the more committed states and they follow the red escape route. The same applies to the situation where $H$ has disappeared due to a saddle-node bifurcation. The cells proceed along the escape route towards state $A$. In response to a change in a signal, the escape route flips towards $B$ and cells instead transition towards it. Such a change is called a heteroclinic bifurcation or a flip bifurcation. **B.** The normal form for the bifurcation set of the binary flip in parameter space. There is a curve (purple) of saddle-node bifurcations destroying the head attractor $H$ and a curve of heteroclinic flips (grey) changing the connection from $A$ to $B$ or vice-versa. The two curves intersect at a fold-flip point (red asterisk) organising the dynamics. **C.** Close pass to a forward saddle $s_{AB}$. Stochasticity results in cells reaching different attractors. **D.** Feedback. In this example as B fills it causes a feedback that flips the escape route to A. **E.** The normal form for the bifurcation set of a binary choice landscape in parameter space. Two curves (purple) represent saddle-node bifurcations where the head attractor $H$ collides with one of two distinct saddles, $s_A$ and $s_B$. These curves meet at a dual cusp singularity. Stochastic fluctuations can lead a cell in $H$ to transition to either attractor $A$ or $B$ depending on the closeness of $H$ to one of the two saddles. When $H$ is close to both saddles, the parameters of the system are close to a dual cusp singularity.

#### B2. Landscape model

##### B2.1. Strategy: modular construction from sub-landscapes models

Our approach to producing a generative dynamical model of the decision making system follows that discussed in [18] but is challenging here because of the number of decisions the system makes. To address this we break the phase space into smaller decision regions containing one or two decisions, associate a gradient-like sub-landscape model to each of these, and then glue them together. The justification of using gradient-like models and the gradient normal forms from catastrophe theory is given in [15,18]. The goal is a parameterised model that explains the effects of extrinsic signals. Using normal forms from catastrophe theory efficiently minimises the number of parameters needed to reproduce the bifurcations suggested by the data. An advantage of normal forms is that although they are defined in arbitrary dimensions, for the transition types and bifurcations identified in this system, only two dimensions suffice to reproduce observed dynamics. The key features are the structures formed by equilibria and 1-dimensional unstable manifolds.

**Definition B2.1** (Sub-landscape model) A *sub-landscape model* is a parameterised system of ODEs of the form: $\dot{\mathbf{x}} = L(\mathbf{x}, \boldsymbol{\theta})$ where $\mathbf{x} \in \mathbb{R}^n$ are the state variables, $\boldsymbol{\theta} \in P$ is a vector of parameters taken from an open domain $P \subset \mathbb{R}^d$ and, for each $\boldsymbol{\theta}$, the vectorfield $L(\cdot, \boldsymbol{\theta})$ is gradient-like away from its bifurcation set. Here we only used $n, d = 2$.

When fixing a vector of parameters $\boldsymbol{\theta} \in P$ away from bifurcation curves, we refer to the resulting vectorfield or phase portrait as a *sub-landscape instance* (or *representative*) to emphasise choosing a particular representative in the parameterised family. In particular, the topology formed by the complex of its equilibria and stable/unstable manifolds is fixed. Being gradient-like, such a system has only a finite number of non-degenerate equilibria that move smoothly if $\boldsymbol{\theta}$ changes [20]. Such systems can be treated as gradient since only small changes near equilibria transform the system into a topologically equivalent gradient system [13]. Furthermore, it is structurally stable: slightly altering parameters $\boldsymbol{\theta}$ without crossing the bifurcation set leaves the dynamics qualitatively unchanged - we can continuously map trajectories of the unaltered system to those of the altered one. Structural changes only occur as $\boldsymbol{\theta}$ crosses the bifurcation set, an inherent property of biological systems [23].

Here, the parameters are functions of the relevant signals/morphogens. A cell $\mu$ exposed to a particular signalling environment is associated with a parameter $\boldsymbol{\theta}^\mu \in P$ and the corresponding landscape instance $\dot{\mathbf{x}} = L(\mathbf{x}, \boldsymbol{\theta}^\mu)$ governs the deterministic dynamics as the cell differentiates.

To build a global model, we constrain the state variables

$\mathbf{x}$ of $L$ to a decision region $B \subset \mathbb{R}^n$ that captures all incoming trajectories and is fixed such that the equilibria of $L$ belong to $B$ for each instance $\boldsymbol{\theta} \in P$. We recall the definition of[15] (SI.A):

**Definition B2.2** (decision region) A *decision region* $B$ for $L$ is a compact domain in $\mathbb{R}^n$ with a smooth topologically spherical boundary $\partial B$ with the property that at all points in $\partial B$, the vectorfield points transversally into the interior of $B$.

Since we only require $n = 2$, we model these decision regions as disks, define a sub-landscape model in each, then glue them to obtain a global model. We assume $\partial B$ is smooth except possibly at a finite number of points. The most important property is that a decision region is globally attracting for the flow of $L(\cdot, \boldsymbol{\theta})$ and contains all the equilibria for each $\boldsymbol{\theta} \in P$.

##### B2.2. Example: NMP and PreNeural sub-landscapes

We start by constructing the first two decisions: from NMP to either PreNeural or Early Mesoderm/Mesoderm (Fig. B2A) and from PreNeural to either p0/p1 or EarlyVentral (Fig. B2B). Because we are mainly interested in neural fates, we treat Early Mesoderm and Mesoderm as a single attractor

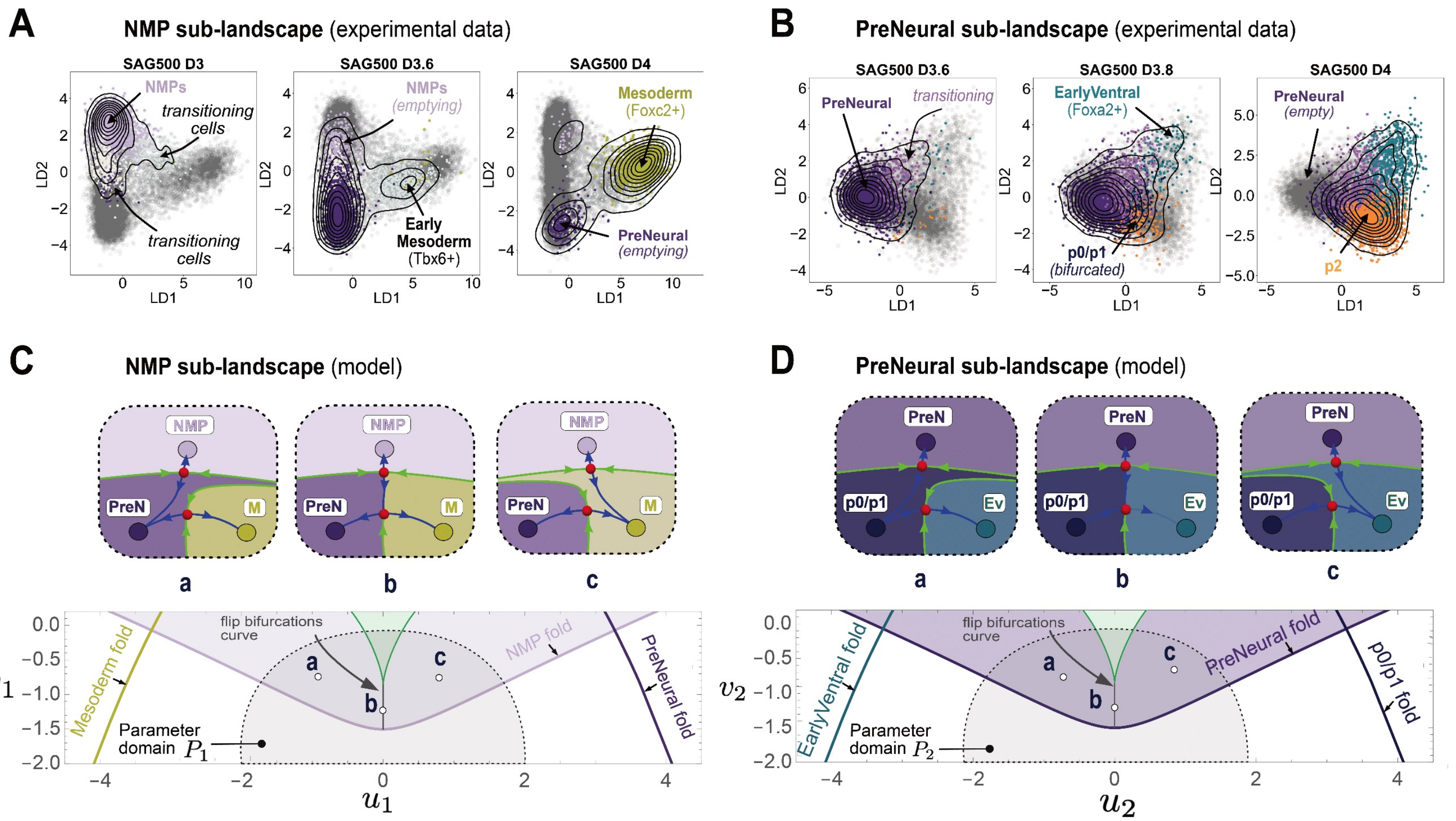

**NMP and PreNeural Sub-landscapes Construction**

**Figure B2:** **A**. 2D LDA representation of the experimental data (scRNA-seq) showing the temporal progression of the ACs NMP, PreNeural and Early Mesoderm/Mesoderm from day D3 to D4. The binary decision from NMP to PreNeural and Early Mesoderm/Mesoderm is modelled by a binary flip sub-landscape. **B**. Same as A for the ACs PreNeural, EarlyVentral, p0/p1. Since the p0/p1 appears bifurcated we also included p2 (yellow). **C**. (Top) Landscape instances (phase portraits) driving the trajectory of a cell escaping the NMP attractor. The attractors are labelled as cells states and the saddles are shown in red. The configuration of the unstable manifolds (blue) is crucial to determine where a cell goes. In (a) the NMP attractor is connected to the PreNeural attractor by an unstable manifold (blue), in (b) the situation is not generic, two saddles are connected by the unstable manifold, in (c) the situation is reversed and NMP is connected to the Early Mesoderm/Mesoderm attractor. Each landscape instance is constrained to a decision region (the same region is fixed for all) which attracts and captures all the incoming trajectories that traverse the boundary of this region transversally. The stable manifolds (green) divide the region into basin of attractions coloured according to the attractor they contain. (Bottom) Bifurcation set in the plane of parameters $\boldsymbol{\theta_1} = (u_1, v_1)$ for a gradient system with compact elliptic umbilic potential (1), modelling the decision from NMP to PreNeural and Mesoderm. For a parameter $\boldsymbol{\theta_1}$ chosen as one of the marked points a, b, c, the corresponding dynamical system is equivalent to the respective landscape instances shown above. The fold bifurcation curves in the parameter plane indicate where the corresponding attractors NMP, PreNeural and Early Mesoderm/Mesoderm undergo a fold bifurcation with a neighbouring saddle; for example, if $\boldsymbol{\theta}_1$ is chosen outside the purple shaded region, the head NMP attractor is bifurcated. The gray curves represent the loci where a flip bifurcation occurs as in instance b. This situation is not generic. For parameters in the central deltoid-shaped region (green) the landscape instances have an extra saddle and repeller. **D**. (Top) Landscape instances for the binary decision from PreNeural to p0/p1 and EarlyVentral. (Bottom) Bifurcation set in the plane of parameters $\boldsymbol{\theta_2} = (u_2, v_2)$ for a gradient system with compact elliptic umbilic potential (1), modelling the decision from PreNeural to p0/p1 and EarlyVentral.

(labelled Mesoderm). We will define sub-landscape models for each decision and constrain their phase space into disk-like decision regions. Next, we will glue their trajectories as parameterised families by using smoothing at the boundary of their decision regions as explained below. This construction can be generalised to allow the systematic coupling of multiple sub-landscapes (Sect. B2.5). This facilitates the 'Lego block' construction from[1] also applied in[17] that took into account more global structures to model two decisions.

The dynamics of an NMP cell will be governed by a sub-landscape instance with a phase portrait of the types shown in Fig. B2C. The second decision from PreNeural to p0/p1 and EarlyVentral follows a similar pattern with the possible sub-landscape instances given in Fig. B2D. Cells transition in accord with one of the processes described in Sect. B1.3: cellular heterogeneity, close pass to a forward saddle, or feedback. Since we see no obvious signs of feedback (such as one state filling before the other) we hypothesise that what we see in this case is due to one of the former two effects.

##### B2.3. Binary flips as sub-landscape

The compactification of Thom's elliptic umbilic catastrophe[15,22] contains a normal form for the binary flip given by a potential function obtained by perturbing Thom's umbilic $y^3 - 3yx^2$ with an additional term[1]. The resulting potential belongs to the double cusp family[25]. This gives a 3-attractor sub-landscape model with a flip bifurcation in a family of gradient vectorfields. To model the first decisions, we take as potential functions

$$f_{\boldsymbol{\theta}_i}(\mathbf{x}) = x^4 + y^4 - (x^2 + y^2) - (y^3 - 3yx^2) - u_i x - v_i y \quad (1)$$

for $i = 1, 2$, one for each sub-landscape, with vector of parameters $\boldsymbol{\theta}_i = (u_i, v_i) \in \mathbb{R}^2$ and state variables $\mathbf{x} = (x, y)$. A choice of gradients yields the bifurcation sets as in Fig. B2C-D. Other candidates for the potential functions and gradients are valid, and we can vary other parameters (eventually adapting the size of the decision regions), as long as we can produce the sub-landscape instances of Fig. B2C-D. What matters initially is that altering the parameters, placing the system in specific regions of the parameter space (*Morse-Smale components*), allows the resulting dynamical system to replicate the qualitative form of the transitions we have observed. Hence, we can validate the model by estimating parameters that give a quantitative fit to data. The binary flip sub-landscape models for the NMP decision and the PreNeural decision are respectively given by $\dot{\mathbf{x}} = L_i(\mathbf{x}, \boldsymbol{\theta}_i)$, for $i = 1, 2$, where the vectorfields $L_i$ are defined as the gradients of the potential functions (1), shifted such that they appear as in Fig B3A. Explicitly these are

$$L_i(\tau_i(\mathbf{x}), \boldsymbol{\theta}_i) = -5\nabla f_{\boldsymbol{\theta}_i}(\tau_i(\mathbf{x})) \quad (2)$$

where $\tau_1(\mathbf{x}) = (x - 1.5, y - 6)$ and $\tau_2(\mathbf{x}) = (x, y - 4)$ translate the coordinates in the plane. The decision regions $B_1, B_2$ are taken to be the disks of radii $r_1 = 2.5$ and $r_2 = 1.9$ centred at $\mathbf{c}_1 = (1.5, 6)$ and $\mathbf{c}_2 = (0, 3.4)$ (Table B2). [1] The vector of parameters are denoted $\boldsymbol{\theta}_i = (u_i, v_i)$ where $i = 1, 2$ and are restricted to abstract domains $P_i \subset \mathbb{R}^2$, located in the region surrounding the fold-flip point as shown in Fig. B2C for $i = 1$ and Fig. B2D for $i = 2$.

##### B2.4. Joining sub-landscapes along their trajectories

To join the vectorfields, the flow lines coming from $B_1$ must traverse the boundary of $B_2$ into the intersection of $B_1$ and $B_2$ as in Fig. B3A. For each $i$, we take a smooth positive bump function $\phi_i : \mathbb{R}^2 \to \mathbb{R}$ that equals 0 inside the decision region $B_i$ and equals 1 at points that have distance greater than $\varepsilon$ from $B_i$. Here $\varepsilon$ is a small positive number. Consider the vectorfield given by

$$\begin{aligned} \mathcal{L}(\mathbf{x}, \boldsymbol{\theta}, vel_A) &= (1 - \phi_1(\mathbf{x}))\phi_2(\mathbf{x})L_1(\tau_1(\mathbf{x}), \boldsymbol{\theta}_1) \\ &\quad + vel_A(1 - \phi_2(\mathbf{x}))L_2(\tau_2(\mathbf{x}), \boldsymbol{\theta}_2). \end{aligned} \quad \text{(FDL)}$$

Then $\mathcal{L}$ is a smooth vectorfield that is smoothly parameterised by $\boldsymbol{\theta} = (\boldsymbol{\theta}_1, \boldsymbol{\theta}_2)$ and $vel_A > 0$ (the latter does not affect the bifurcations). If $N$ is the strip of points in $B_1$ that are at distance less than $\varepsilon$ from the boundary of $B_2$ then $\mathcal{L}$ equals $L_2$ on $B_2$ away from the strip, equals $L_1$ on $B_1 \setminus (N \cup B_2)$, and smoothly interpolates between these two regions on $N$. The parameters $\boldsymbol{\theta}_1$ and $\boldsymbol{\theta}_2$ are constrained (by restricting $P_1, P_2$) so that at no time do any attractors or saddles involved in bifurcations pass into $N$. Then in this constrained parameter space, the bifurcations observed that take place wholly within $B_1 \setminus ((B_1 \cap B_2) \cup N)$ or $B_2 \setminus (B_2 \cap N)$ correspond exactly to bifurcations of $L_1$ and $L_2$. For example, in the case discussed, to ensure that the upper attractor of $B_2$ (PreNeural) captures the unstable manifold coming from $B_1$, the decision regions should be positioned so that the lower left attractor of $L_1$ is close to PreNeural.

##### B2.5. Assembling the global model

The global landscape $\mathcal{L}$ is a combination of 6 sub-landscapes $\dot{\mathbf{x}} = L_i(\tau_i(\mathbf{x}), \boldsymbol{\theta}_i)$, summarised in Table B1, with state variables $\mathbf{x}$ taken within their decision region $B_i \subset \mathbb{R}^2$. We can combine these using bump functions $\phi_i(\mathbf{x})$ (Table B2) that allow smooth transitions between the different regions as in Fig. B3B. The global landscape is

$$\begin{aligned} \mathcal{L}(\mathbf{x}, \boldsymbol{\theta}, \mathbf{vel}) &= (1 - \phi_1(\mathbf{x}))\phi_2(\mathbf{x})L_1(\tau_1(\mathbf{x}), \boldsymbol{\theta}_1) \\ &\quad + vel_A(1 - \phi_2(\mathbf{x}))\phi_3(\mathbf{x})L_2(\tau_2(\mathbf{x}), \boldsymbol{\theta}_2) \\ &\quad + vel_B(1 - \phi_3(\mathbf{x}))\phi_4(\mathbf{x})\phi_6(\mathbf{x})L_3(\tau_3(\mathbf{x}), \boldsymbol{\theta}_3) \\ &\quad + (1 - \phi_4(\mathbf{x}))\phi_6(\mathbf{x})(\phi_5(\mathbf{x})L_4(\tau_4(\mathbf{x}), \boldsymbol{\theta}_4) \\ &\quad + vel_C(1 - \phi_5(\mathbf{x}))L_5(\tau_5(\mathbf{x}), \boldsymbol{\theta}_5)) \\ &\quad + (1 - \phi_6(\mathbf{x}))L_6(\tau_6(\mathbf{x}), \boldsymbol{\theta}_6) \end{aligned} \quad \text{(GL)}$$

where the landscape vector of parameters

$$\boldsymbol{\theta} = (\boldsymbol{\theta}_1, \boldsymbol{\theta}_2, \boldsymbol{\theta}_3, \boldsymbol{\theta}_4, \boldsymbol{\theta}_5, \boldsymbol{\theta}_6) \quad (3)$$

is restricted to a domain $P \subset \mathbb{R}^{12}$ because each $\boldsymbol{\theta}_i = (u_i, v_i) \in P_i \subset \mathbb{R}^2$ are restricted to their own domains $P_i$.

The additional parameters

$$\mathbf{vel} = (vel_A, vel_B, vel_C) \in \mathbb{R}^3_+ \quad (4)$$

are (positive) velocity parameters that are used to adapt the timing of transition in some decision regions. They do not alter the topology of the trajectories.

Sub-landscape $L_5$ contains a repeller (Fig. B3C) which will allow cells to transition to ventral states via two distinct pathways. A landscape instance with all the attractors present is shown in Fig. B3D-E.

[1]We may wish to make the vectorfields (2) generic by slightly perturbing them but this would not bring much to the analysis.

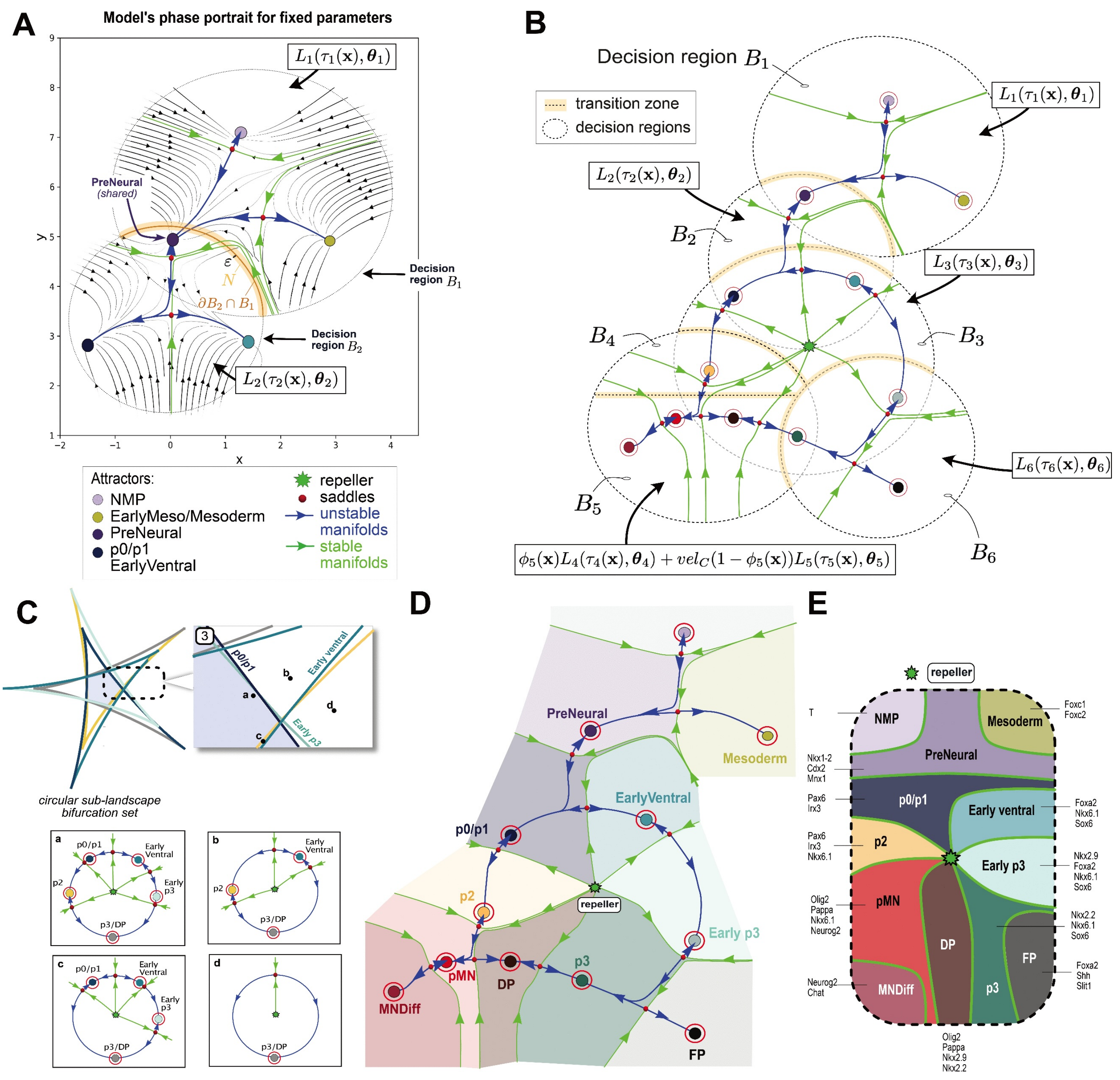

Global Landscape Model Construction

**Figure B3:** **A**. Connecting the First Decision Landscapes. Gluing the two sub-landscape models $L_1, L_2$ along the strip $N$ creates the illustrated compound vectorfield for a fix set of parameters $\boldsymbol{\theta}_1, \boldsymbol{\theta}_2$. Unstable manifolds (blue), stable manifolds (green), saddles (red) and flow lines (arrowed curves) characterise this vectorfield. **B**. Landscape instance for the global model with the sub-landscapes decision regions $B_i$. **C.** Full bifurcation set of the circular sub-landscape $L_3$ and the sub-landscape instances (a, b, c, d) that appear for the parameter $\boldsymbol{\theta}_3$ in a specific region $P_3$ of the bifurcation set. After the gluing construction, the fold bifurcations of the p2 and Early p3 attractors are controlled by other parameters. The grey attractor represents the combined ventral fates (DP, p3). **D.** Landscape instance for the global model with coloured basin of attractions that contain each of the attractors. **E.** Schematic representation of the basins of attraction shown in D, separated by stable manifolds (green) and the corresponding genes expressed in each basin. The central asterisk denotes a repeller (index-2 saddle).

| $L_i$ | Vectorfield $L_i(\mathbf{x}, \boldsymbol{\theta}_i)$ with $\mathbf{x} = (x, y)$ and $\boldsymbol{\theta}_i = (u_i, v_i)$ | translation $\tau_i(\mathbf{x})$ |
|---|---|---|
| $L_1$ | $\dot{x} = 5(-2x + 6xy + 4x^3 - u_1)$ <br> $\dot{y} = 5(-2y + 3x^2 - 3y^2 + 4y^3 - v_1)$ | $(x - 1.5, y - 6)$ |
| $L_2$ | $\dot{x} = 5(-2x + 6xy + 4x^3 - u_2)$ <br> $\dot{y} = 5(-2y + 3x^2 - 3y^2 + 4y^3 - v_2)$ | $(x, y - 4)$ |
| $L_3$ | $\dot{x} = 50(-2x + 6x(x^2 + y^2)^2 + x^3y - xy^3 + \frac{1}{100}v_3)$ <br> $\dot{y} = 50(-2y + 6y(x^2 + y^2)^2 + \frac{1}{4}x^4 - \frac{3}{2}x^2y^2 + \frac{1}{4}y^4 + \frac{1}{100}u_3)$ | $(0.4x, 0.4y - 0.6)$ |
| $L_4$ | $\dot{x} = -2x + 12xy + 4x^3 - u_4$ <br> $\dot{y} = -2y + 6x^2 - 6y^2 + 4y^3 - v_4$ | $(2x + 3.5, 2y - 1)$ |
| $L_5$ | $\dot{x} = 4x^3 + 4xy^2 - 6x - 12xy + v_5$ <br> $\dot{y} = -4yx^2 + 6y - 6y^2 + 6x^2 - 204y^3 - 1.4x - u_5$ | $(x + 1.8, -y - 0.2)$ |
| $L_6$ | $\dot{x} = 4x^3 + 4xy^2 - 8x + 6x^2 - 6y^2 + u_6$ <br> $\dot{y} = 4y^3 + 4yx^2 - 8y - 12yx + v_6$ | $(2x - 2.6, 2y + 0.5)$ |

**Table B1:** Sub-landscape vectorfields. The vectorfields are then evaluated at $\tau_i(\mathbf{x})$.

| i | bump function $\phi_i(\mathbf{x})$ | Decision region $B_i = B_{\delta_i}(\mathbf{c}_i)$ |
|---|---|---|
| **1** | $\frac{1}{2}(\tanh(10((x - 1.5)^2 + (y - 6)^2 - 2.5^2)) + 1)$ | $\{\mathbf{x} \in \mathbb{R}^2 : (x - 1.5)^2 + (y - 6)^2 \leq 2.5^2\}$ |
| **2** | $\frac{1}{2}(\tanh(10(x^2 + (y - 3.4)^2) - 1.9^2) + 1)$ | $\{\mathbf{x} \in \mathbb{R}^2 : x^2 + (y - 3.4)^2 \leq 1.9^2\}$ |
| **3** | $\frac{1}{2}(\tanh(10(x^2 + (y - 1.6)^2) - 2.4^2) + 1)$ | $\{\mathbf{x} \in \mathbb{R}^2 : x^2 + (y - 1.6)^2 \leq 2.4^2\}$ |
| **4** | $\frac{1}{2}(\tanh(10((x + 2)^2 + y^2) - 2^2) + 1)$ | $\{\mathbf{x} \in \mathbb{R}^2 : (x + 2)^2 + y^2 \leq 2^2\}^*$ |
| **5** | $\frac{1}{2}(\tanh(10(y - 0.5)) + 1)$ | $\{\mathbf{x} \in \mathbb{R}^2 : (x + 2)^2 + y^2 \leq 2^2\}^*$ |
| **6** | $\frac{1}{2}(\tanh(10((x - 1.2)^2 + (y + 0.5)^2) - 1.6^2) + 1)$ | $\{\mathbf{x} \in \mathbb{R}^2 : (x - 1.2)^2 + (y + 0.5)^2 \leq 1.6^2\}$ |

**Table B2:** Bump functions and decision regions. *Include the decision regions $B_4, B_5$.

#### B3. Parameterising and simulating the landscape

##### B3.1. Stochastic differential equation

At this stage the dynamics of the global landscape

$$\dot{\mathbf{x}} = \mathcal{L}(\mathbf{x}, \boldsymbol{\theta}, \mathbf{vel}) \tag{5}$$

is deterministic with $\boldsymbol{\theta}$ in the domain $P$ and $\mathbf{vel}$ is the velocity adjustment as in (4). To account for the stochastic nature of the dynamics we consider instead the stochastic differential equation:

$$d\mathbf{x}(t) = \mathcal{L}(\mathbf{x}(t), \boldsymbol{\theta}, \mathbf{vel}, t)dt + \eta(\mathbf{x}(t), \boldsymbol{\sigma}, t)d\mathbf{w}(t) \tag{6}$$

where $\mathbf{w}(t)$ is a 2-dimensional Wiener process, $\eta(\mathbf{x}(t), \boldsymbol{\sigma}, t)$ is a diffusion coefficient that depends on the space variable and a noise-intensity parameter vector $\boldsymbol{\sigma}$, $\mathcal{L}(\mathbf{x}(t), \boldsymbol{\theta}, \mathbf{vel}, t)$ is the deterministic part representing the global landscape (GL).

Since the global landscape is composed of several normal form models, some having "flatter" potential function than others, using a constant diffusion coefficient would yield poor results, as it would disproportionately affect certain sub-landscapes compared to others. Therefore, we apply distinct diffusion coefficients in specific regions of the phase space. While a different diffusion coefficient could be defined within each decision region of each sub-landscape, this would introduce an excessive number of unnecessary parameters. Thus, we use only four distinct scalar noise parameters $\boldsymbol{\sigma} = (\sigma_A, \sigma_B, \sigma_C, \sigma_D)$ across specific regions of the phase space and combine them to form a global, space-dependent diffusion coefficient:

$$\begin{aligned} \eta(\mathbf{x}(t), \boldsymbol{\sigma}, t) \quad = \quad & (1 - \phi_1(\mathbf{x}(t))\phi_2(\mathbf{x}(t)))\phi_3(\mathbf{x}(t))\sigma_A \\ & +(1 - \phi_3(\mathbf{x}(t)))\phi_4(\mathbf{x}(t))\phi_6(\mathbf{x}(t))\sigma_B \\ & +(1 - \phi_4(\mathbf{x}(t)))\phi_6(\mathbf{x}(t))\sigma_C \\ & +(1 - \phi_6(\mathbf{x}(t)))\sigma_D \end{aligned} \tag{7}$$

with the bump functions $\phi_i(\mathbf{x})$ listed in Table B2. The stochastic differential equation depends on parameters of the form $\mathbf{p} = (\boldsymbol{\theta}, \mathbf{vel}, \boldsymbol{\sigma})$, in which $\boldsymbol{\theta} \in P \subset \mathbb{R}^{12}$ as in (3) controls the bifurcations of each sub-landscape, $\boldsymbol{\sigma} \in \mathbb{R}^4_+$ is a noise parameters vector and $\mathbf{vel} \in \mathbb{R}^3_+$ modifies the velocity of some of the sub-landscapes. Increasing velocity (or noise) can influence bifurcations and higher velocity parameters may allow cells to more easily bypass saddle points as exemplified in [3]. To avoid overly noisy simulations we restrict the parameters $\mathbf{p}$ of the SDE to a subset $D \subset P \times \mathbb{R}^3_+ \times \mathbb{R}^4_+$.

##### B3.2. Simulations of cell's trajectories

We simulate $m = 500$ trajectories of the SDE (6) with (GL) as the deterministic part, one for each cell $\mu$ on its transition pathway, using using the Euler–Maruyama method [7,12]. Each simulated cell's trajectory $t \mapsto \mathbf{x}^\mu(t)$ evolves according to its own SDE, characterised by a unique set of parameters $\mathbf{p}^\mu$ that are close but not identical to each other. The small stochastic variation in the parameters represents the cell heterogeneity discussed in the paper.

The implementation of the simulation algorithm proceeds as follows: we start by fixing a "mean" parameter vector $\mathbf{p}$. We then take $m$ perturbed parameters $\{\mathbf{p}^\mu\}_{\mu=1}^m$ obtained from $\mathbf{p}$ by adding a Gaussian perturbation with zero mean and a small variance to each component resulting in $m$ parameters covering a small region of parameter space.

To simulate $m$ cell trajectories we need to provide initial conditions in phase space: we fix a coordinate $\mathbf{x}_0 \in B$ in the decision region of the global landscape, corresponding to the initial location of the NMP population. We then generate a 2D Gaussian cluster of $m$ initial conditions $\mathbf{x}^\mu(0)$ with indices $\mu = 1, \ldots, m$ in $B$ corresponding to the initial positions of the NMP cells in phase space. The mean vector of the cluster of points are the coordinates $\mathbf{x}_0$, and the covariance matrix is diagonal with entries 0.05. To simulate the SDE (6) with the global landscape (GL) as the deterministic part (5), we used the time interval $[0, 4]$ with timestep $dt = 0.01$. We thus obtain a total of 399 discrete time points $t_i$ and a position of each cell $\mathbf{x}^\mu(t_i)$ at each time point with initial condition $\mathbf{x}^\mu(0)$.

##### B3.3. Example: simulation of the combined NMP and PreNeural sub-landscapes

As an example to illustrate the simulation procedure, we restrict ourselves to the first two binary decisions between D3 and D4 discussed in Sect. B2.3 and B2.4. We use the SDE (6) with the first decisions landscape (FDL) as deterministic part $\mathcal{L}$ and diffusion coefficient

$$\eta(\mathbf{x}(t), \sigma_A, t) = (1 - \phi_1(\mathbf{x}(t))\phi_2(\mathbf{x}(t)))\sigma_A \tag{8}$$

with $\sigma_A \in \mathbb{R}_+$, focusing exclusively on the two first decision regions $B_1 \cup B_2$. For the simulation, we use a shorter time interval $[0, 1]$ and timestep $dt = 0.01$. Fig. B4A shows the synthetic data generated by simulating $m = 500$ cells trajectories, retaining the positions of the cells in phase space at 6 discrete time points (Fig. B4B). Clearly, each cell follows a distinct trajectory as they progress towards one of the attractors p0/p1, EarlyVentral or Mesoderm. A cell $\mu$ differentiating towards p0/p1 follows the blue route in Fig. B4A along a simulated trajectory $\mathbf{x}^\mu(t)$ of the SDE (6) with landscape parameters $\boldsymbol{\theta_1}^\mu$ (resp. $\boldsymbol{\theta_2}^\mu$) which are marked by a blue dot in Fig. B4C (resp. Fig. B4D). The corresponding landscape instance is therefore as in Fig. B4E: the unstable manifold connected to the PreNeural attractor is directed towards p0/p1. By contrast, a cell transitioning towards EarlyVentral evolves according to a landscape instance as in Fig. B4F with the unstable manifold from PreNeural directed towards EarlyVentral. Finally, a cell transitioning towards Mesoderm will follow a landscape instance as in Fig. B4G, avoiding the neural part of the landscape.

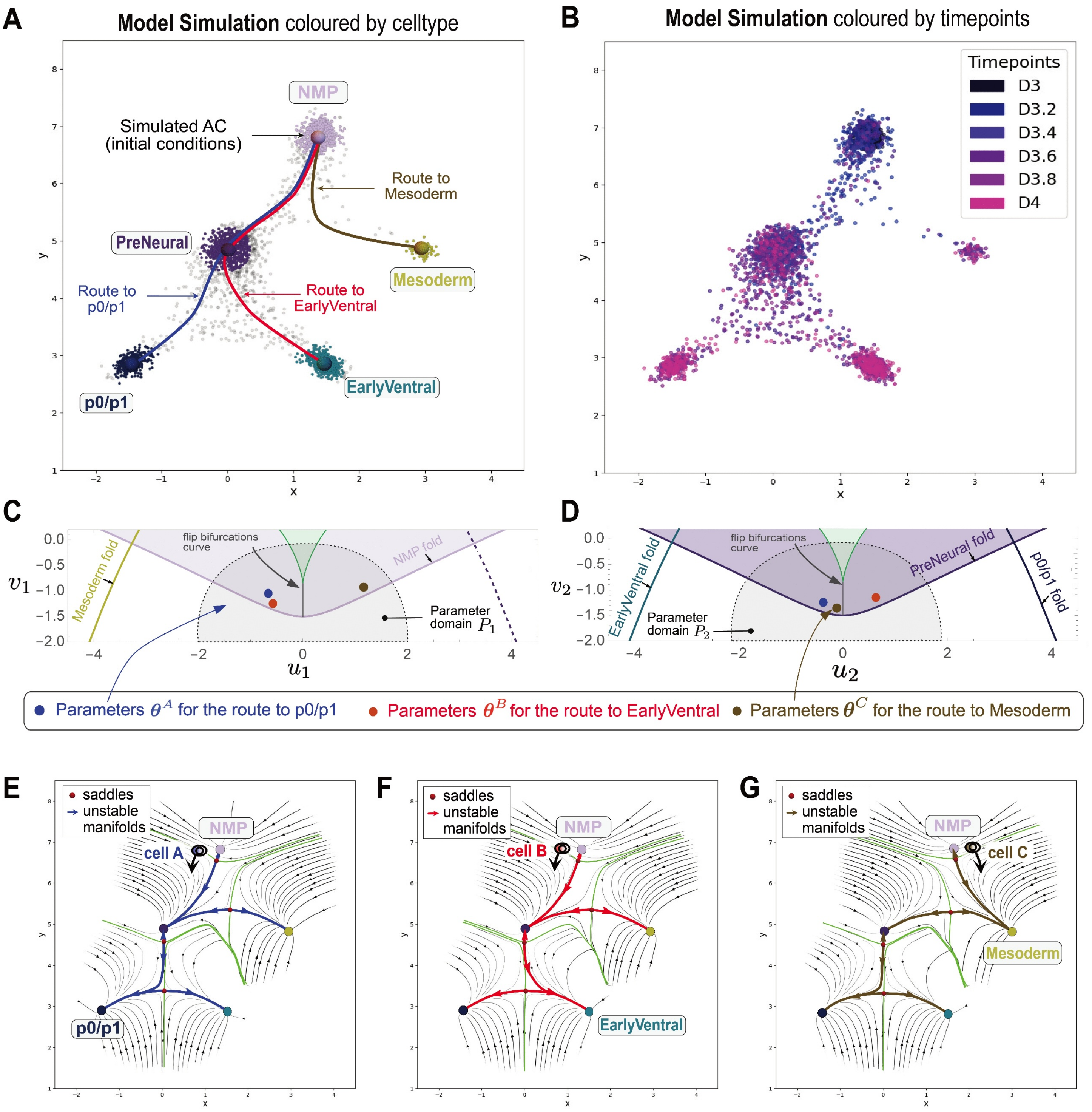

**Simulation Example of Cell Trajectories in the Landscape Model**

**Figure B4:** **A**. Synthetic data generated by simulating $m = 500$ cell trajectories starting in a Gaussian cloud of points near the marked-point NMP. Each cell trajectory $t \mapsto \mathbf{x}^{\mu}(t)$ is derived from its own SDE (6) defined by a unique vector of parameters $\mathbf{p}^{\mu}$. **B**. Simulated cells coloured by 6 discrete intermediate time points between D3 and D4. **C**. Parameter domain $P_1$ that determines the dynamics of the NMP sub-landscape model plotted on top of the bifurcation set. The coloured dots represent distinct parameters $\boldsymbol{\theta}_1^{\mu} = (u_1, v_1)$ of the NMP sub-landscape $L_1$, with $\mu = A, B, C$ corresponding to the labels of different cells $\mu$. **D**. Same as C for the PreNeural sub-landscape $L_2$ with sub-landscape parameters $\boldsymbol{\theta}_2^{\mu} = (u_2, v_2)$ taken within the parameter domain $P_2$. **E**. A landscape instance for a cell transitioning towards p0/p1. The landscape parameters $\boldsymbol{\theta}_1^{A} \in P_1$ in C and $\boldsymbol{\theta}_2^{A} \in P_2$ in D to obtain this configuration are marked by a blue dot in C and D, respectively. The NMP and PreNeural attractors are close enough to their saddle such that the cell can easily escape over them. The unstable manifold connecting PreNeural is directed towards p0/p1. In this scenario, the cell can also potentially escape towards EarlyVentral due to stochastic fluctuations but is more likely to transition to p0/p1. **F**. A landscape instance for a cell transitioning towards EarlyVentral with corresponding parameters $\boldsymbol{\theta}_1^{B}$, $\boldsymbol{\theta}_2^{B}$ marked by red dots in C and D. **G**. A landscape instance for a cell transitioning directly from NMP towards Mesoderm. Note that $\boldsymbol{\theta}_1^{C} \in P_1$ in C (indicated by a brown dot) is far to the right of the flip bifurcation curve (gray) forcing cell C to transition from NMP to Mesoderm.

#### B4. Fitting the model to simulated proportions

In this section we discuss fitting the landscape model to the experimental data and how this makes the abstract parameter domains $P_i$ of the sub-landscapes explicit. We continue using the guiding example of the first two binary decisions in the scRNA-seq dataset to illustrate the method. As explained in Sect. B3.3 and shown in Fig. B4A, we simulate $m = 500$ cells trajectories starting near the NMP-marked point along the time interval $[0, 1]$, with timestep $dt = 0.01$, using the SDE (6) with deterministic part (FDL) and diffusion coefficient (8). The domain of parameters $\mathbf{p} = (\boldsymbol{\theta}, vel_A, \sigma_A)$ of the SDE in this case is a subset $D \subset P \times \mathbb{R}_+ \times \mathbb{R}_+$ where $P = P_1 \times P_2$ is the joined domain of parameters of the two sub-landscapes (Sect. B2.4).

##### B4.1. Computing proportions of cells in each attractor

For each simulation of $m$ cell trajectories, we retain the coordinates of these cells at the 6 time points $t_0, t_{20}, t_{40}, t_{60}, t_{80}, t_{99}$, corresponding respectively to days D3, D3.2, D3.4, D3.6, D3.8 and D4 in the experiment. This gives the synthetic data of Fig. B5A. We then compute the proportions of simulated cells, at each of these time points, located within a distance of 0.5 of the "mean" locations of the attractors that represent stable cell states obtained by uniformly sampling parameters within the region, computing the position of the attractors, and averaging. These locations are identified as regions of high cell density, meaning that, at each time point of the simulation, the simulated cells cluster more densely in certain areas of phase space, indicating the mean locations of the systems attractors. We keep in mind that, since each cell responds to its own SDE, the precise locations of the attractors in its landscape slightly differ from that of another cell, as the parameters fixing the dynamics vary between cells. However, given the choice of landscape parameters that can vary and because we restricted them to a small region $P = P_1 \times P_2$, these variations are not substantial and it is reasonable to consider the mean location of attractors. Since an attractor may undergo bifurcation, with trajectories passing through the remnants, we also aim to capture these "ghost" states. If the parameters are taken in a region where a particular attractor is not present, these ghost states do not appear clearly as high-density points but rather as trails of cells moving through them, which will result in smaller proportions (for example PreNeural can be bifurcated, yet cells still traverse it).

##### B4.2. Comparing simulated proportions to data

Each simulation produces a $6 \times 5$ matrix $X_{sim}$ whose entries $(X_{sim})_{ij}$ are the simulated proportions of cells, at time point $t_i$, within a small distance of each of the high density points $j$ named after the five cell states NMP, PreNeural, p0/p1, EarlyVentral and the combined Early Mesoderm/Mesoderm. We want to compare the simulated proportions matrix $X_{sim}$ with the matrix of proportions $X_{data}$ of the experimental data at fixed SAG concentration $s = 500$nM (Fig. B5B left). Since there are also p2-like cells at D4 in the experimental data, we combined these proportions with those of p0/p1, for the sake of this example. To assess how closely the simulated proportions match the real proportions, we use the standard $L_1$-*distance*, calculated as the sum of the absolute pointwise differences between the matrices entries:

$$d(X_{sim}, X_{data}) = \sum_{i,j} |(X_{sim})_{ij} - (X_{data})_{ij}|. \tag{9}$$

##### B4.3. Fitting using ABC approach

Ideally we would like to determine a (signal-dependent) domain $D_\varepsilon(s) \subset D$ containing all the parameters $\mathbf{p} = \mathbf{p}(s)$ of the SDE (6) for which the simulated proportions most closely align with those of the experimental data. By that we mean that, for any $\mathbf{p} \in D_\varepsilon(s)$, the proportion matrix $X_{sim}$ obtained by simulating $m = 500$ trajectories with parameters $\mathbf{p}^\mu$ around $\mathbf{p}$ satisfies $d(X_{sim}, X_{data}) < \varepsilon$ with $\varepsilon$ as small as possible. At this stage we have only one SAG concentration, $s = 500$nM. However, we will introduce multiple concentrations that will require us to determine a domain of parameters for each of them individually. This is why we indicate the dependence on $s$ in the notation.

For fitting the model, we use Approximate Bayesian Computation (ABC) based on[24] (and references therein) following the methodology of[1,2]. ABC approximates the posterior distribution giving the probability of the parameters $\mathbf{p}$ conditional on the data $P(\mathbf{p}|X_{data})$. We take advantage of the PyABC package developed for Python in[11,19]. The procedure begins by defining priors distributions for the parameters $\mathbf{p}$. In our case these are uniform distributions supported in the parameter domain $D$ of the SDE (Table B3 ).

| Parameter | Priors | Parameter | Priors |
|---|---|---|---|
| $\mathbf{u_1}$ | $\mathcal{U}([-1, 1])$ | $\mathbf{v_1}$ | $\mathcal{U}([-2.5, -1])$ |
| $\mathbf{u_2}$ | $\mathcal{U}([-1, 1])$ | $\mathbf{v_2}$ | $\mathcal{U}([-2.5, -1])$ |
| $\sigma_A$ | $\mathcal{U}([0.2, 0.8])$ | $\mathrm{vel}_A$ | $\mathcal{U}([0.05, 1.2])$ |

**Table B3:** Priors used for the ABC fitting for the sub-landscape parameters $\boldsymbol{\theta}_i = (u_i, v_i)$ for $i = 1, 2$, noise magnitude $\sigma_A > 0$ and velocity parameter $\mathrm{vel}_A > 0$.

The algorithm is iterative: PyABC first samples N parameters from these priors distributions and runs N simulations, each of them generating a matrix of proportions, which are then compared to the averaged proportions of the experimental data using the distance function (9). Parameters producing a distance smaller than a initial acceptance threshold $\varepsilon_0$ are accepted, while others are dismissed. The sampling continues until N parameters are accepted. This defines a distribution of accepted parameters and a new generation starts by resuming sampling within the distribution defined by the previously accepted parameters. At each new generation, a smaller threshold $\varepsilon$ is calculated as the $\alpha$-quantile of the distances obtained at the previous step with $\alpha = 0.2$.

To sample new parameters, PyABC uses a transition kernel at each step of the algorithm to perturb the parameters accepted at the previous generation. The transition kernel implemented in pyabc.LocalTransition uses multivariate normal kernels with an adaptive covariance matrix[6]. By iteratively sampling from an increasingly accurate approximation of the posterior, PyABC gradually reduces $\varepsilon$ at each generation. The sampling stops either because a maximal number of generations is reached or a 0.1 $\varepsilon$-threshold is attained.

To fit the stochastic landscape model, we chose $N = 1500$ and a maximal number of 8 generations. We also tested greater values such as $N = 5000, 3000, 2000$ and the results remained similar. In fact, because the hypercube region defined by our priors is relatively restricted, iterative samplings of $N = 1500$ parameters is sufficient to fully explore the entire space. Fig. B5B (right) shows the mean of the simulated proportions for each cell state at each time point for the N accepted parameters after 8 generations. With each generation we observe a decrease in the $\varepsilon$ threshold with a minimal threshold value $\varepsilon_{min} = 0.85$ (Fig. B5C). Note that whether this value is considered sufficiently small depends on the number of entries of the matrix $X_{data}$ as adding a small error to each entry increases the overall distance (9). The proportions retained for the accepted parameters form relatively tight distributions around their mean (Fig. B5D), which may be an indication that the number of generations is set too high.

##### B4.4. Explicitly defining the parameter domains of the landscape

The result of the ABC fitting is a distribution of parameters $\mathbf{p} = \mathbf{p}(s)$ within the priors parameter domain $D$ of the SDE. We can define $D_\varepsilon(s) \subset D$ as the 80% confidence ellipsoid of this distribution. We are mainly interested in the domains $P_i(s) \subset P_i$ of the accepted parameters components $\boldsymbol{\theta}_i = \boldsymbol{\theta}_i(s)$ corresponding to each sub-landscape $L_i$ for $i = 1, 2$. This information reveals the different type of model instances that are most likely to emerge when cells are exposed to SAG concentration $s$ (Fig. B2C-D). For each sub-landscape model $L_i$, we consider the plane that supports the parameters $\boldsymbol{\theta}_i$. We then project the parameters $\mathbf{p} \in D_\varepsilon(s)$ to its sub-landscape components $\boldsymbol{\theta}_i$ in this plane. The image of this projection is represented by a 2D smooth density which defines the boundary of the domains $P_i(s)$ for $i = 1, 2$. The result is shown in Fig. B5E: Panel 1 shows the parameter domain $P_1(s)$ (blue) for the NMP sub-landscape that were found within the prior region taken inside the initial abstract domain $P_1$. The parameters $\boldsymbol{\theta_1} = (u_1, v_1)$ converged sharply indicating that the model is approaching overfitting (Sect. B4.5). On the other hand, the parameter domain $P_2(s)$ (blue) for the PreNeural sub-landscape in Panel 2 covers a broader region of the prior domain inside $P_2$. It remains close to the flip bifurcation curve (grey) near the fold-flip point but within the region where the PreNeural attractor has just bifurcated and where it is connected to p0/p1 as shown below.

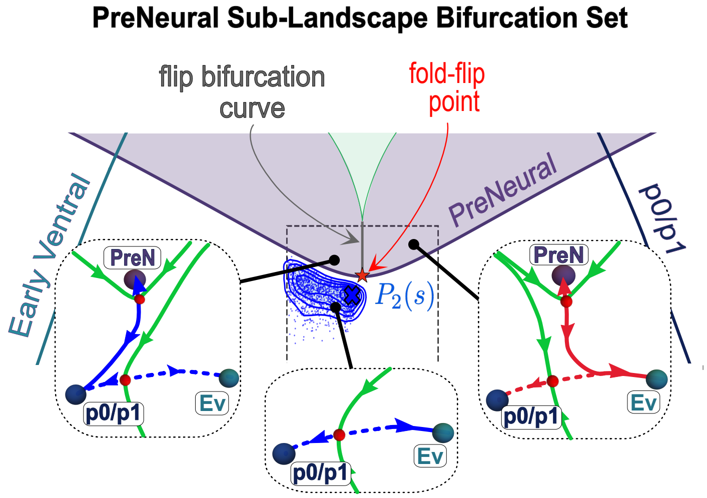

This means that the trajectories of the cells at this signal concentration $s$ are driven by dynamical systems close to the one of the left above (blue unstable manifolds) with PreNeural and its nearby saddle removed (middle).

The synthetic data used to compute the simulated proportions at each retained time point are shown in Fig. B5F. They show the progression of the cells in phase space as they follow their landscape.

##### B4.5. Limitations

Overfitting occurs when the learned model unduly fits the intricacies (and even noise) found in the training data instead of capturing the underlying pattern in a way that allows for better generalisation to future unseen data. When fitting ABC to stochastic systems, the sampling process at each generation continues until $N$ parameters producing distances within a small threshold are accepted. However, if the allowed number of generations is set too high or the final tolerance is set too low, then such overfitting may occur, and the parameter distribution found may be very different from the true posterior. To guard against this, we ensure that the final tolerance is not too small. Usually, one uses a training and test approach to detect overfitting. However, for data like ours, generating test datasets is too expensive. On the other hand, in general, overfitting will produce a posterior distribution that is too compact, and this can be detected by comparing across the different datasets that we have. It can also be detected by perturbing the datasets computationally, as this should destroy the excessive fit to the original dataset.

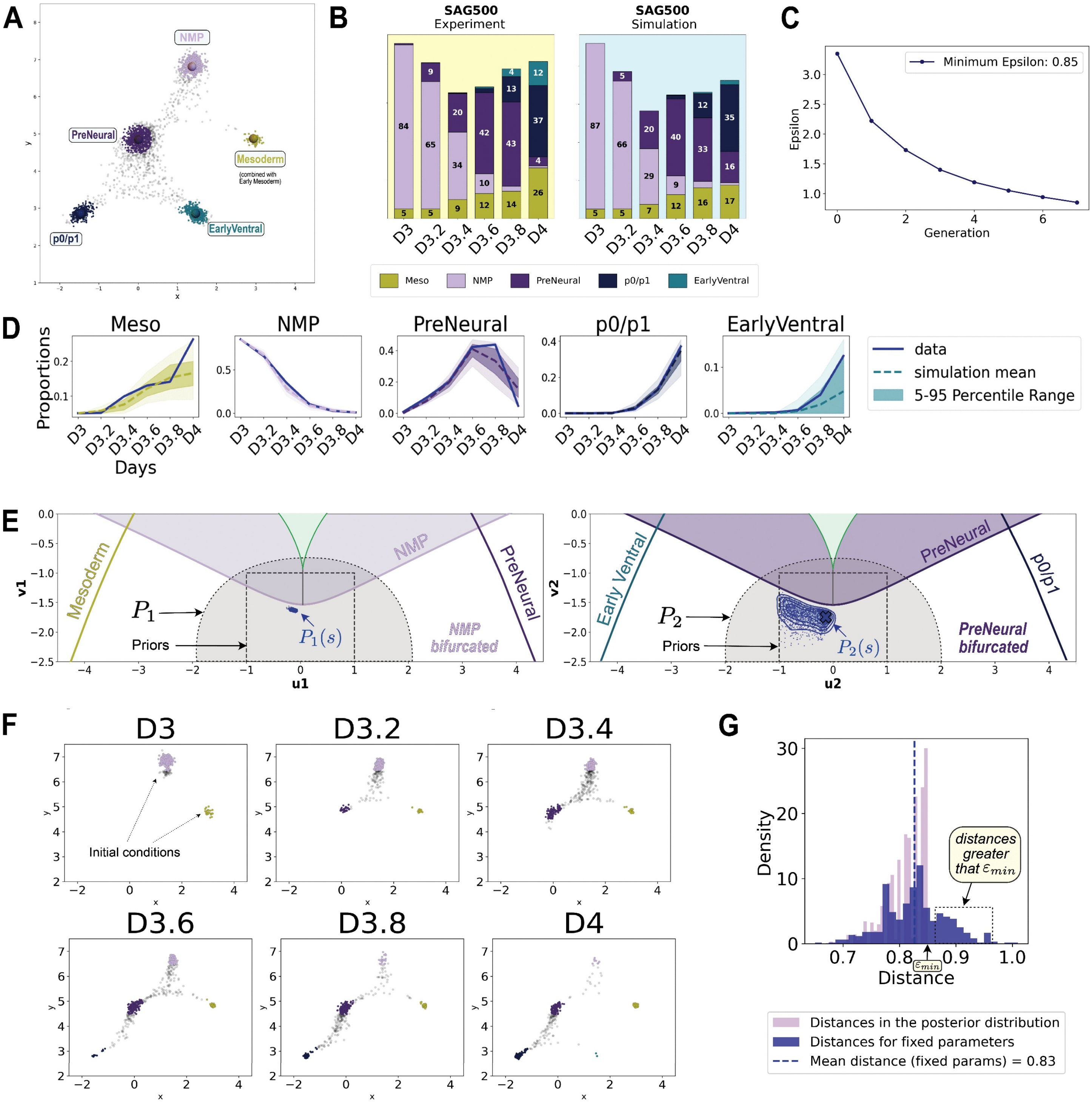

**Parameter Fitting Example for Sub-landscape Model**

**Figure B5:** (Caption next page)

**Figure B5:** **A**. Stochastic simulation of $m = 500$ cells trajectories for the First Decisions Landscape model (FDL), retaining positions of cells at 6 time points. The coloured clusters represent the simulated attractor clusters (ACs). **B**. (Left) The proportions of cells (in percent) in each AC at the 6 measurement time points from the experimental data. For this example, we combined the proportions of p0/p1 and p2 cells, and also the proportions of Early Mesoderm and Mesoderm. The blank spaces represent the proportions of transitioning cells: NMPTrans and PreNTrans (not shown). (Right) Mean of the simulated proportions for the $N = 1500$ parameters accepted by the ABC fitting after 8 generations. **C**. Decrease in $\varepsilon$-threshold over the 8 ABC generations. **D**. Distributions over time for the $N = 1500$ simulated proportions accepted at the final generation of the ABC fitting (percent values in decimal form). The mean proportions of these simulations are represented by dashed curve and the experimental data proportions by thick blue curves. The NMP and p0/p1 proportions appear close to overfitted, whereas the experimental proportions of EarlyVentral are further away from the means of the simulated proportions, yet still in the confidence interval. The variances at each time point for the EarlyVentral are wider. **E**. Bifurcation sets of the first two binary decisions: NMP sub-landscape (left) and PreNeural (right). The abstract parameter domains $P_1$ and $P_2$ were chosen so that the regions can be glued and are sufficient to reproduce the landscape instances needed to model these decisions. The initial sample of $N = 1500$ parameters for the ABC fitting are uniformly distributed within the priors domains (dashed rectangles), chosen for the ABC fitting initialisation. These parameters converged to form well-defined distributions (blue), making the parameter domains explicit. **F**. Synthetic data of simulated cells at each time point. To ensure we start with appropriate initial conditions, we also initialised with a proportion of cells in Mesoderm and NMPTrans (gray) at D3. The clusters of cells within a distance 0.5 of their respective mean attractor are used to compute the simulated proportions. **G**. Distribution of distances of the N parameters accepted by the ABC fitting at the 8th generation (purple). The minimal threshold $\varepsilon_{min}$ attained is 0.85. However, if we pick a parameter $\mathbf{p}$ among the N accepted parameters and run a simulation multiple time with the same $\mathbf{p}$, we obtain the distribution of distances in blue. In particular, the upper quantile goes beyond the $\varepsilon_{min}$ value. This illustrates that whether ABC fitting accepts or rejects a parameter is a matter of chance and the minimal $\varepsilon_{min}$ is not definitive.

#### B5. Fitting the model to the FACs data

We demonstrated the approach of simulating and fitting the first two binary decisions to the scRNA-seq data at early time points (Sect. B4.3). We now fit the global model to the proportions of cell states derived from the flow cytometry data. In this case we have datasets for 4 different SAG concentrations $s \in \{0, 10, 100, 500\}$. For each $s$, we simulate proportions matrices as above and compare them with the matrix of proportions $X_{data}$ of the biological data, obtained by averaging the proportions of the 3 experimental replicates *March 12*, *October 16*, *April 16* at SAG concentration $s$, and the SAG concentrations 0nM and 500nM from two extra experiments *Nov5*, *June 5* (Appendix A Fig. A8C).

We simulate $m = 500$ cells trajectories as explained in Sect. B3.2 over the time interval $[0, 4]$ and timestep $dt = 0.01$. For each simulation of $m$ cells trajectories, we retain the coordinates of these cells at the 5 time points $t_0, t_{100}, t_{200}, t_{300}, t_{399}$, corresponding respectively to days D3, D4, D5, D6, D7 in the experiment. In this case, each simulation produces a $5 \times 10$ matrix $X_{sim}$ the entries of which are the simulated proportions of cells, at time point $t_i$, within a distance 0.5 of each of the high density points $j$ named after the 10 cell states NMP, PreNeural, p0/p1, p2, pMN, DP, EarlyVentral, Early p3, p3 and FP. Cells belonging to the Mesoderm were not generated in the FACs experiments because NODAL and BMP inhibitors had been applied. Although our model includes Mesoderm, cells do not reach this state.

Since the FACs datasets did not contain the first and last time points D3 and D7 and specific markers for FP were not included, we excluded the corresponding matrix entries at this stage. We denote by $X'_{sim}$ the proportions matrix obtained from $X_{sim}$ by removing the column corresponding to FP and the rows corresponding to D3 and D7. The resulting simulated FP proportions will be used as predictions. We used the priors listed in Table B4. For each SAG concentration $s$, the result of the ABC fitting is a distribution of $N = 1500$ parameters, $\mathbf{p}(s)$ in $D \subset \mathbb{R}^{19}$, drawn from the approximated posterior which, after a few generations started converging into (possibly disconnected) elliptical regions. The simulated proportions $X'_{sim}$ for each of these parameters satisfy $d(X'_{sim}, X_{data}) < \varepsilon$ for some $\varepsilon$ - that is in practice greater than the minimal threshold $\varepsilon_{min}$ found at the last ABC generation as shown in Fig. B5G.

For each signal concentration $s$, the proportions retained for the accepted parameters form relatively tight distributions around their mean and align well with the proportions observed in each individual experiment (Fig. B6A-B).

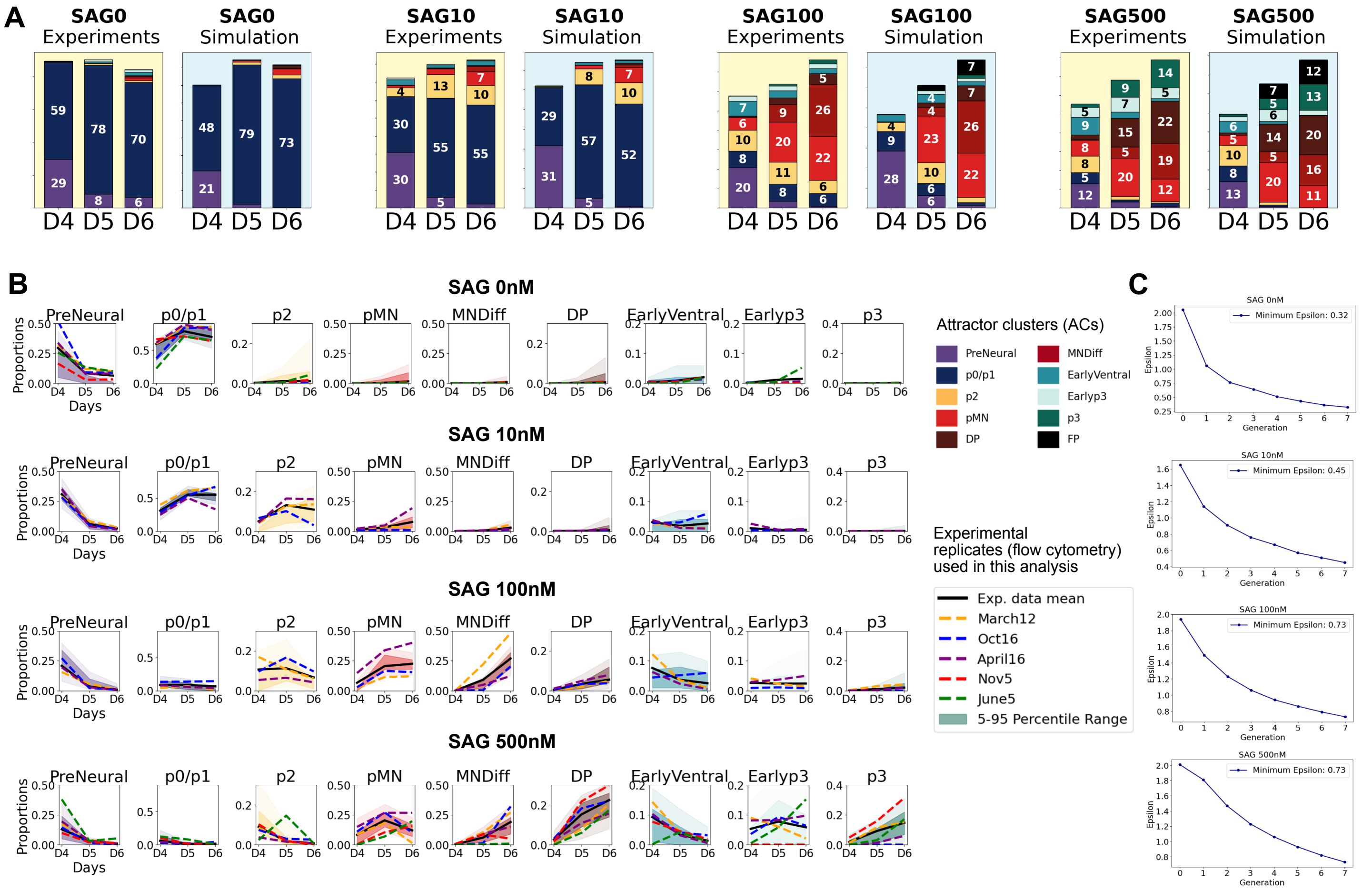

**Cell Proportions in FACs Datasets Across Experimental Replicates**

**Figure B6:** **A**. Stacked barplots showing averaged experimental proportions (yellow background) of ACs next to the mean of the simulated proportions (blue background) for the $N = 1500$ parameters retained from the ABC fitting. The FP proportions in the SAG100 and SAG500 Simulation panels are retained as model predictions. **B**. Simulated proportions by day using the $N = 1500$ parameters accepted by the ABC fitting for each signal concentration (percent values in decimal form). The dashed curves correspond to the experimental proportions *March 12*, *October 16*, *April 16* (for all SAG concentrations) and *Nov 5*, *June 5* (for only 0nM and 500nM SAG). The black thick curves correspond to the averaged experimental proportions as shown in A (yellow panels). **C**. Decrease of the $\varepsilon$-threshold after 8 generations of the ABC fitting.

**SAG 0 nM, SAG 10nM**

| Parameter | Priors | Parameter | Priors |
|---|---|---|---|
| $\mathbf{u_1}$ | $\mathcal{U}([-2.5, 0])$ | $\mathbf{v_1}$ | $\mathcal{U}([-2, -0.5])$ |
| $\mathbf{u_2}$ | $\mathcal{U}([-3, 0.5])$ | $\mathbf{v_2}$ | $\mathcal{U}([-2.2, -0.5])$ |
| $\mathbf{u_3}$ | $\mathcal{U}([2, 14])$ | $\mathbf{v_3}$ | $\mathcal{U}([-4, 10])$ |
| $\mathbf{u_4}$ | $\mathcal{U}([-12, 0.3])$ | $\mathbf{v_4}$ | $\mathcal{U}([-9, -1])$ |
| $\mathbf{u_5}$ | $\mathcal{U}([6, 8])$ | $\mathbf{v_5}$ | $\mathcal{U}([-1.2, 0.8])$ |
| $\mathbf{u_6}$ | $\mathcal{U}([2, 8])$ | $\mathbf{v_6}$ | $\mathcal{U}([6, 14])$ |
| $\sigma_i$ | $\mathcal{U}([0.2, 0.8])$ | $\text{vel}_i$ | $\mathcal{U}([0.05, 1.2])$ |

**SAG 100 nM, SAG 500nM**

| Parameter | Priors | Parameter | Priors |
|---|---|---|---|
| $\mathbf{u_1}$ | $\mathcal{U}([-2.5, 0])$ | $\mathbf{v_1}$ | $\mathcal{U}([-2, -0.5])$ |
| $\mathbf{u_2}$ | $\mathcal{U}([-3, 0.5])$ | $\mathbf{v_2}$ | $\mathcal{U}([-2.2, -0.5])$ |
| $\mathbf{u_3}$ | $\mathcal{U}([6, 23])$ | $\mathbf{v_3}$ | $\mathcal{U}([-4, 10])$ |
| $\mathbf{u_4}$ | $\mathcal{U}([-12, 0.3])$ | $\mathbf{v_4}$ | $\mathcal{U}([-9, -1])$ |
| $\mathbf{u_5}$ | $\mathcal{U}([6, 8])$ | $\mathbf{v_5}$ | $\mathcal{U}([-1.2, 0.8])$ |
| $\mathbf{u_6}$ | $\mathcal{U}([2, 8])$ | $\mathbf{v_6}$ | $\mathcal{U}([6, 14])$ |
| $\sigma_i$ | $\mathcal{U}([0.2, 0.8])$ | $\text{vel}_i$ | $\mathcal{U}([0.05, 1.2])$ |

**Table B4:** Priors used for the ABC fitting of FACs at constant SAG concentration. All noise and velocity parameters had same priors.

##### B5.1. Bifurcations associated with the sub-landscapes

As before we take $D_\varepsilon(s) \subset D$ as the 80% confidence ellipsoid of this distribution and the corresponding sub-landscapes parameter domains $P_i(s)$ for the different $s$ are represented by 2D density plots in each parameter plane that supports the bifurcation set of the corresponding sub-landsdcape $L_i$. These domains are displayed in Fig. B7A where they are labelled as SAG0 (purple), SAG10 (yellow), SAG100 (orange), SAG500 (red) according to the SAG concentration $s$. The highest density points in each is marked by a $\times$. The bifurcation sets divide the planes into components, each of which produces qualitatively different dynamics (sub-landscape instances). The fold bifurcation curves are coloured according to the attractor they bifurcate (saddle-node or dual cusp bifurcation). The flip bifurcations curve, indicating where an unstable manifold directly connects two saddles, is shown in gray. In Fig. B7A, the panels $(i)$ are labelled according to the sub-landscape index $i$. Because at 0nM SAG we mainly see p0/p1 cells, the domain SAG0 is only shown in Panels (2) and (3) as the parameters did not converge for the other sub-landscapes as all cells remain stuck in the PreNeural and the p0/p1 state. We comment on each panel individually:

(1) **NMP sub-landscape** $L_1$**:** This binary flip sub-landscape governs the transitions from the NMP head attractor to Early Mesoderm/Mesoderm or PreNeural (Fig. B2A). In the scRNA-seq dataset, the NMP cells (TBXT/Bra-expressing) at D3 do not remain at D4 and we thus assume that, at this stage, they already adopted a PreNeural identity. Accordingly, all the parameter domains are located below the NMP fold curve where the NMP attractor undergoes a fold bifurcation with its neighbouring saddle. With NODAL and BMP inhibitors applied, Mesoderm cells were not expected to be generated, and the fitting results selected the best candidates for the parameter domains to the left of the flip bifurcation curve but below the NMP fold bifurcation curve. This results in cells adopting a neural identity rather than a mesoderm identity.

(2) **PreNeural sub-landscape** $L_2$**:** The transition from PreNeural is highly dependent upon the SAG concentration, with cells more likely to transition to the EarlyVentral state at high concentrations (100nM, 500nM). This is reflected by the proximity of the corresponding parameter distributions SAG100 (orange) and SAG500 (red) to the flip bifurcations curve (gray). This indicates that a proportion of cells adopt a ventral identity (EarlyVentral/Early p3/p3/FP) in high SAG concentrations but are constrained to intermediate fates (p0/p1, p2 or pMN) at lower concentrations (0nM, 10nM).

(3) **Circular sub-landscape** $L_3$**:** The circular sub-landscape primarily controls the bifurcations of p0/p1 and EarlyVentral (Fig. B3D). For the transition from p0/p1 to p2, the parameter distributions SAG0 (purple) and SAG10 (yellow) are positioned below and relatively far from the p0/p1 bifurcation fold curve in the shaded blue region. This indicates that, at these SAG concentrations, the p0/p1 attractor is far enough from its neighbouring saddle that it retains cells until D6. However, the distribution is still close enough to the fold curve to allow some cells to escape due to stochasticity. As a result, at 10nM SAG, we observe a mix of p0/p1 and p2 cells at D6. The parameter distributions for SAG100 (orange) and SAG500 (red) are well above the p0/p1 bifurcation curve (blue), resulting in the bifurcation of the p0/p1 attractor. In this scenario, all p0/p1 cells eventually transition to p2. The parameters of the circular sub-landscape enable tuning of the EarlyVentral attractor position relative to its neighbouring saddle, facilitating cell transitions towards Early p3, p3 and FP. The EarlyVentral attractor appears to be bifurcated (or close to bifurcation) for SAG100 and SAG500 while it remains well established for SAG10 meaning that cells are unlikely to escape. This is consistent with the experimental observations in Appendix A Fig. A13D (SAG10). This sub-landscape alone also controls the bifurcations of p2 and Early p3 (Fig. B3A) but the sub-landscapes were only glued along their decision regions and not along their bifurcation sets. However, further deformation of the vectorfields that define this sub-landscape could deform the p2 and Early p3 fold bifurcation curves of Panel 3 such that they coincide with those of Panel 4 and Panel 6 relatively to the positions of the parameter distributions.

(4) **p2 sub-landscape** $L_4$**:** The key observation here is that the p2 attractor is either very shallow (SAG10), indicated by the closeness of the domain of SAG10 parameters to the p2 fold bifurcation curve, or absent (SAG100 and SAG500). This indicates that, even with a low level of SAG, the p2 attractor is shallow and cells quickly transition towards the Olig2 positive attractors (pMN and DP). On the other hand, the distribution SAG500 (red) is located below the p2 fold bifurcation curve which implies that, by increasing the SAG concentration from 10nM to 100nM or 500nM SAG, the p2 attractor bifurcates and a cell that was stuck in this attractor transitions to pMN or DP. Although the model was designed to allow the unstable manifold connecting p2 to flip to either pMN or DP, the domains of accepted parameters appeared biased in favour of a connection to pMN rather than DP. Hence the transition towards DP are due to stochastic fluctuations or from a direct transition from pMN as we further explain below in (5).

(5) **pMN sub-landscape** $L_5$**:** This panel shows the bifurcation set for the sub-landscape governing the transitions MNDiff-pMN-DP-p3. At the core of the parameter space lies a dual cusp singularity where the two pMN fold bifurcations curves meet. These enclose a shaded red region where the 4 attractors exist and are arranged on a line, smoothly connected by unstable manifolds (see below).

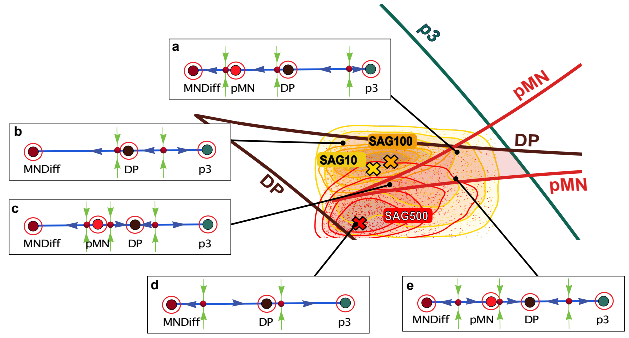

From this region (c), the pMN attractor can bifurcate with two distinct saddles, modelling a binary choice decision between either MNDiff or DP. The parameter domain SAG10 (yellow) did not converge sharply but appears skewed towards the upper region, extending beyond the DP bifurcation curve. This suggests that the DP attractor is close to bifurcation for many parameters so even if cells reach it, they return to the pMN attractor (a). This aligns with the experimental data where we observed that the DP cluster does not fully form at 10nM SAG (Appendix A Fig. A14F) .

The parameters in the domain SAG100 are constrained to the region where the DP attractor is present and more DP cells form at this concentration. Both high density points × (SAG10) and × (SAG100) lie just above the bifurcation curve where pMN has bifurcated. In this case the cell transitions directly to MNDiff (b). However, many other parameters lie within the red shaded region of the bifurcation set where the pMN attractor is present.

The domain SAG500 contains many parameters that lie below the red shaded region, and primarily beyond the fold curve where the pMN attractor bifurcates, leading more cells into the DP attractor (d). Indeed we observed that there are very few pMN cells at D6 compared to D5 suggesting that cells rapidly differentiate to MNDiff or DP.

(6) **Early p3 sub-landscape** $L_6$**:** The fit of this sub-landscape to the FACs dataset is not as precise as it would be if a FP cluster was defined (as we will do later in Sect. B7 for the scRNA-seq). However, we can use the proportions of Early p3 and p3 cells to make predictions about the proportions of FP cells as simulated cells end up in this attractor. The domain of parameters for SAG10 is broadly dispersed across the rectangular region defined by the priors, as there are almost no cells transitioning into the ventral sub-landscape. The parameter domains for SAG100 and SAG500 on the other hand are biased towards the upper region and on top of the Early p3 fold curve, suggesting that this attractor is bifurcated and cells directly transition towards p3 or FP.

As previously explained in Sect. B3.2, for each parameter vector $\boldsymbol{\theta}$, as in (3), taken in the domains of Fig. B7A, there is a corresponding global landscape instance that governs the dynamics of a cell. In Fig. B7B we show 3 of these instances, one for each of the 3 set of parameters marked by a cross × (SAG10), × (SAG100), × (SAG500), indicating the high density points for each domain. For a parameter domain such as SAG10, multiple dynamical system instances are possible as the parameters are spread across distinct components of the bifurcation sets, each of which give a topologically distinct system. The ones shown in Fig. B7B are specific examples. The two panels in Fig. B7C-D provide a closer view of the intermediate sub-landscape for a parameter inside the red shaded region of Panel 5 where a binary choice occurs.

This analysis also suggests that the distribution of cell states at an intermediate signal regime between 100nM and 500nM SAG (for example SAG200) will generate cell states proportions close to the ones observed at SAG500 as the corresponding landscape representatives in that case are qualitatively equivalent.

##### B5.2. Notes on the accuracy of the fitting and the interpretation of the result when dealing with stochastic systems

The concept of a shallow attractor is to be taken with some care as this is highly dependent on the type of diffusion and its magnitude, which is fixed by the priors[3]. In our case we chose a space-dependent diffusion coefficient and we constrained the magnitude of the noise parameters to remain relatively close to zero. We argue that, even if the resulting distributions of parameters are not exact, and naturally depend on several choices we have made, they remain highly informative and allow conclusions that align well with experimental observations. For example, it is known that the progenitors p0, p1 are not fully generated - the markers Dbx1, Dbx2 do not appear - when the SAG concentration is high, which the simulation successfully reflects (p0/p1 appears bifurcated at SAG 100nM and 500nM). This observation is replicated in a different scRNA seq dataset (Sect. B9.1). Furthermore this analysis demonstrates that p2 is always shallow even under low SAG exposure (10nM). This is consistent with the difficulty of generating a large number of p2 cells in vitro without additional "contaminating" cell types such as Olig2-expressing cells.

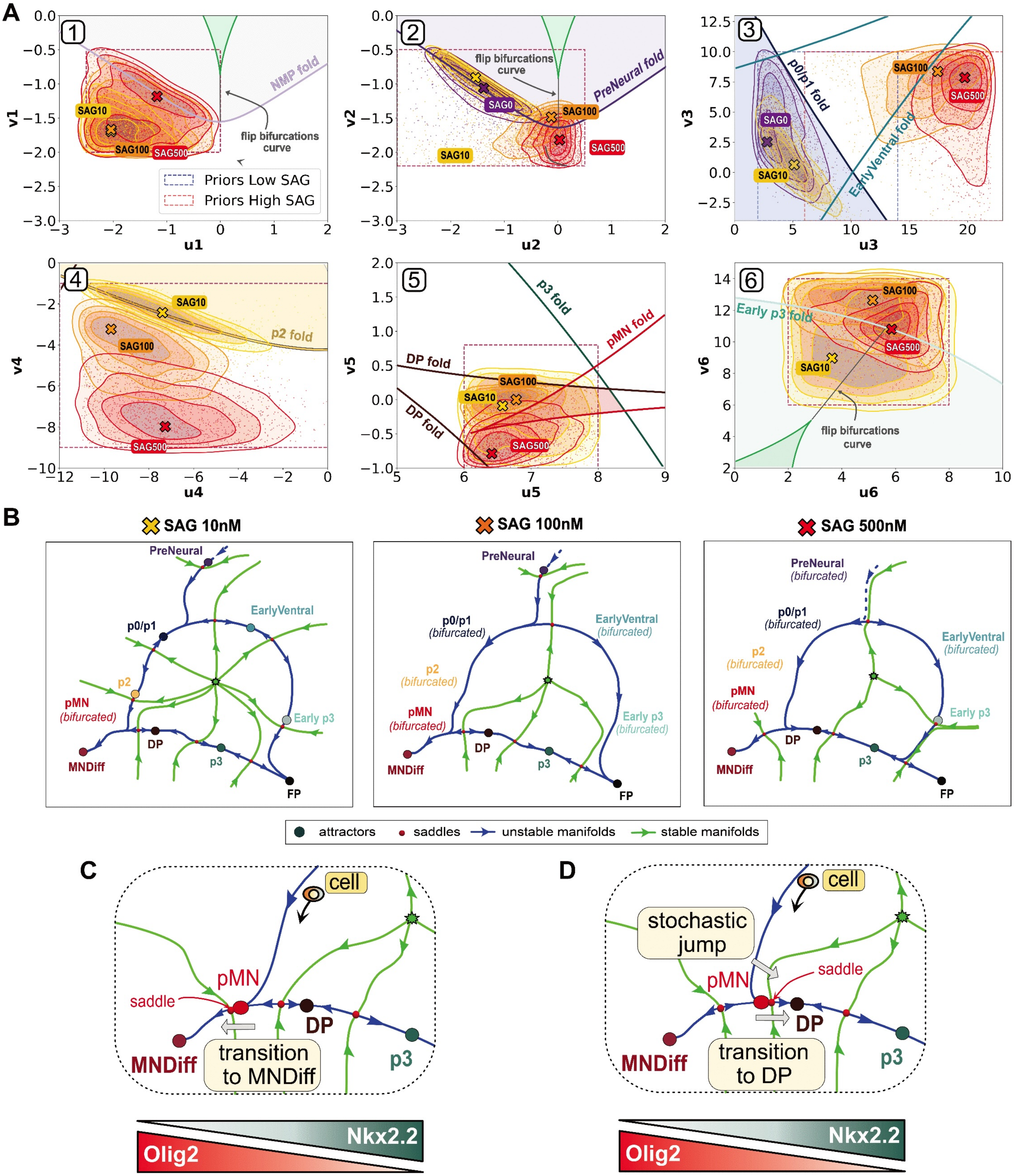

**SAG Concentration-Dependent Bifurcations in Neural Progenitor Development**

**Figure B7:** **A**. Accepted parameter domains (shaded contour lines) within their prior regions (dashed rectangle) drawn on the bifurcation set of each sub-landscape. The points marked by a cross × (SAG0), × (SAG10), × (SAG100), × (SAG500) indicate the point of highest density for each domain. The function of each parameter on the sub-landscape bifurcations are summarised in Table B5 below. **B**. Global landscape instances corresponding to the sub-landscapes parameters $\boldsymbol{\theta}_i$ marked by a cross for the domains SAG10, SAG100 or SAG500. Each parameter set taken within these domains for a fixed SAG concentration will give a slightly different representative. The ones shown here for SAG10 and SAG100 have their pMN attractor bifurcated driving the cell directly to MNDiff. Other parameters $\boldsymbol{\theta}_5$ in Panel 5 taken within the shaded red region give rise to landscape instances with a pMN attractor. **C**. Sub-landscape instance with parameters $\boldsymbol{\theta}_5$ in Panel 5 within the shaded red region where the pMN attractor is close to its left saddle and a cell reaching pMN can easily transition to MNDiff. **D**. Same as C but the pMN cell transitions to DP.

| Parameter | Bifurcations controlled by the parameter |
|---|---|
| $u_1$ | Flip bifurcation of the unstable manifold from NMP to either PreNeural or Mesoderm |
| $v_1$ | Fold bifurcation of the NMP attractor |
| $u_2$ | Flip bifurcation of the unstable manifold from PreNeural to intermediate (p0-p2) or ventral (p3, FP) fates |
| $v_2$ | Fold bifurcation of the PreNeural attractor |
| $u_3, v_3$ | Jointly control the fold bifurcations of the p0/p1 and EarlyVentral attractors |
| $u_4$ | Flip bifurcation of the unstable manifold from p2 to pMN or DP. |
| $v_4$ | Fold bifurcation of the p2 attractor |
| $u_5, v_5$ | Jointly control the fold bifurcations of the pMN, DP, p3 and MNDiff attractors |
| $u_6, v_6$ | Jointly control the fold bifurcation of Early p3 and the flip bifurcation from Early p3 to either p3 or FP |

**Table B5:** Type of bifurcations controlled by the sub-landscapes parameters $\boldsymbol{\theta}_i = (u_i, v_i)$.

#### B6. Model predictions using FACs data

In the previous section we obtained a landscape SDE model (6) with explicit parameter domain $D(s)$ for each fixed SAG concentration $s$ using ABC fitting. We use the explicit parameter domains $D(0)$ and $D(500)$ to test the predictive potential of our model by simulating untested conditions involving changes in the SAG concentration during the experiment.

##### B6.1. Early pulse experiment

We simulated an experimental protocol "SAGUP" in which 0nM SAG was applied to cells from D3 to D3.5 (12h), followed by 500nM SAG from D3.5 to D4 (12h) and then cells were returned to 0nM SAG until D6 (Fig. B8A). Constant exposure to 0nM and 500 nM where compared as controls. The corresponding FACs dataset comprising these 3 experiments is named *June 5* in Appendix A Sect. A3.4.

To compare the experimental proportions of cells in each attractor cluster (AC) with simulated proportions, we excluded the proportions corresponding to the designated FP cluster. In this case, the matrix $X_{data}$ is a $4 \times 9$ matrix, with rows corresponding respectively to D3.5, D4, D5, D6, and columns corresponding to the cellstates PreNeural, p0/p1, p2, pMN, MNDiff, DP, EarlyVentral, Early p3 and p3.

##### B6.2. Model simulation and prediction

To simulate 500nM SAG, we first take the $N = 1500$ parameters $\mathbf{p}^\mu(500)$ accepted by the particle filter in Sect. B5 and use these to simulated $N$ cells trajectories that generate simulated ACs. As done before, we retain the FP proportions as a prediction. The simulated proportions show good agreement with the experimental data (Fig. B8B). Additionally, the FP proportions predicted by the simulation at D6 (13%) are consistent with the experimental proportions of FP-like cells (9%) at D6. To simulate the SAG pulse we proceeded slightly differently from Section B3.2 because we wanted to use the domain of parameters found by the ABC fitting and take into account the change of signal concentration. We take the $N = 1500$ accepted parameters $\mathbf{p}^\mu(0)$ of the ABC fitting within the distribution $D(0)$ together with the $N$ accepted parameters $\mathbf{p}^\mu(500)$ within the distribution $D(500)$. As done previously, we choose the time interval $[0, 4]$ with timestep $dt = 0.01$ and divide it into 400 discrete time points $t_i$ to simulate the trajectory of a single cell $\mu$. The cell will follow its own SDE (6) with a parameter $\mathbf{p}^\mu$ that will be updated at each time point $t_i$. This is necessary to account for the change in signal concentration $s$ from 0 to 500 at time $T_1 = 50$ (index of $t_{50}$), corresponding to D3.5, followed by a change from 500 to 0 at $T_2 = 100$ (index of $t_{100}$), corresponding to D4. To do this, we use a combination of 2 sigmoid functions: $\phi_1(t_i) = \frac{1}{2}(\tanh((t_i - T_1)) + 1)$ and $\phi_2(t_i) = \frac{1}{2}(\tanh(0.1(t_i - T_2)) + 1)$. Together they define a bump function $b_{T_1,T_2}(t) = \phi_1(t)(1 - \phi_2(t))$. The SDE parameters of the cell are updated at each time point $t_i$ as

$$\mathbf{p}^\mu(t_i) = (1 - b_{T_1,T_2}(t_i))\mathbf{p}^\mu(0) + b_{T_1,T_2}(t_i)\mathbf{p}^\mu(500). \tag{10}$$

The bump function $b_{T_1,T_2}$ allows a sharp change in concentration $s$ from 0 to 500 at D3.5, followed by a slower exponential decrease from 500 to 0 starting at D4 (Fig. B8C). We thus simulated $m = 1500$ cells trajectories following this methodology and using, for each cell $\mu$, a pair of parameters vectors $\mathbf{p}^\mu(0)$ and $\mathbf{p}^\mu(500)$ taken from the distributions $D(0)$ and $D(500)$ respectively. We retain the position of each cell at the 4 time points $t_{50}, t_{100}, t_{200}, t_{399}$ corresponding respectively to the experimental time points D3.5, D4, D5, D6. We then computed the proportions of cells located within a distance of 0.5 from the mean location of the attractors. As in Sect. B5 we removed the column corresponding to FP in the simulated proportions matrix and used the $4 \times 9$ truncated matrix $X'_{sim}$ to compare to the experimental proportions $X_{data}$. The resulting simulated proportions are given in Fig. B8D. The model predicted a significant increase of p0/p1 cells at D4, peaking around D5 and persisting through D6. The simulated proportions of p0/p1 cells remained high across all days as reducing SAG concentration to 0 deepens the p0/p1 attractors, effectively trapping the cells in the p0/p1 state. The experimental proportions (thick blue curve) however do not follow this tendency as p0/p1 proportions keep decreasing over time despite removal of the signal.

We thus refitted the model keeping the SDE parameters in $D(0)$ and $D(500)$ fixed but allowing the time points $T_1$ and $T_2$, when a signal change occurs, to vary. The priors were taken as uniform distribution in $[50, 100]$ for $T_1$ and in $[100, 399]$ for $T_2$. The $N = 500$ parameters accepted by the particle filter are shown in Fig. B8E. The accepted $T_1$ parameters are sharply distributed around their mean which is located between D3.5 and D4. This indicates that the cells responded to the increase of signal concentration from 0 to 500 within a few hours of D3.5, which is experimentally plausible. However, the accepted $T_2$ parameters appear to be clustered towards the end of the timeline around D6, suggesting that the cells did not respond to the decrease of signal from 500 to 0. The corresponding bump function $b_{T_1,T_2}$ for the means of the fitted parameters $T_1$ and $T_2$ in Fig. B8F illustrates this. The simulated proportions based on these fitted parameters now show good alignment with the experimental data (Fig. B8G-H). We also note that the simulated proportions of FP cells (4%) are consistent with the experimental proportions of FP-like cells (7%).

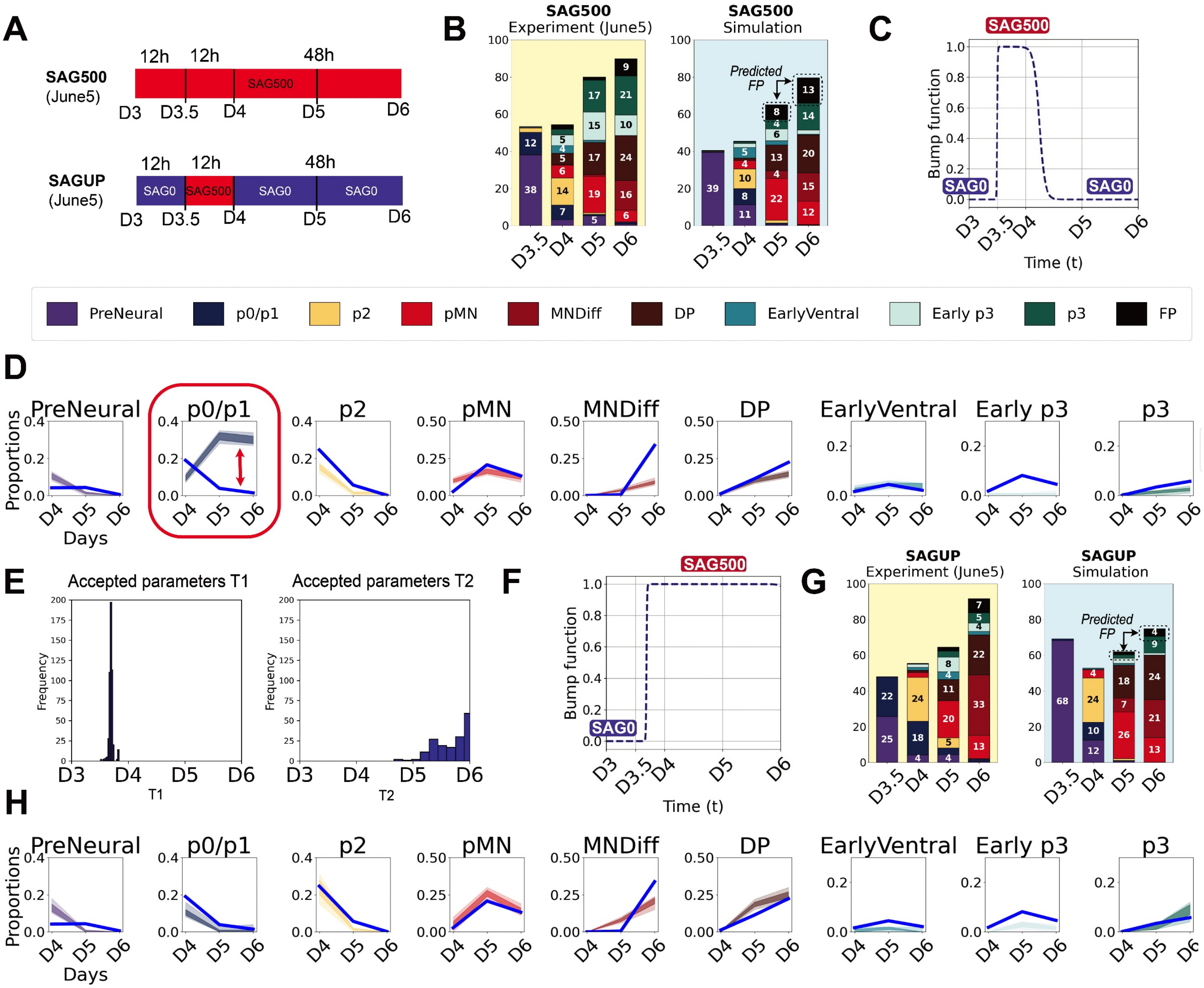

**Modelling SAG perturbation**

**Figure B8:** (Caption next page)

**Figure B8:**

**A**. Summary of two experimental protocols for the series of experiments *June 5*. (Top) Cells are exposed to constant 500nM SAG for 72h. (Bottom) Cells are exposed to 0nM SAG for the first 12h, followed by a pulse of 500nM SAG for 12h and then back to 0nM SAG for the remaining 48h.

**B**. Proportions of cells in the ACs from experimental data (left) compared to simulated proportions (right) for the 500nM SAG condition. The simulation used the previously fitted parameter distribution $D(500)$ obtained in Sect. B5. The FP proportions predicted by the simulation at D6 $(13\%)$ are consistent with the experimental proportions of FP-like cells $(9\%)$ at D6.

**C**. Bump function $b_{T_1,T_2}$ as a function of time $t$ models a sharp increase in signal concentration at D3.5 and an exponential decrease at D4. Here we take $T_1 = 50$ and $T_2 = 100$.

**D**. Distribution of the simulated proportions (coloured by cell state) compared with the experimental proportions (percent values in decimal form). Shaded region indicates $95\%$ confidence interval for the simulated proportions. For each cell state, the thick blue curve represents the experimental proportions and the dashed curve represents the mean of the simulated proportions. These distributions are obtained by pooling $N = 1500$ parameters from the previously fitted parameter distributions $D(0)$ and $D(500)$ from Sect. B5. Then simulate proportions by changing the parameters using the bump function in C. The simulated proportions of p0/p1 did not align with the experimental data as the simulation predicts a sharp increase of p0/p1 cells that remain in equilibrium from D5 to D6 (red rectangle). The data however showed that the proportion of p0/p1 cells decreased despite the removal of the signal.

**E**. Distribution of the $T_1, T_2$ parameters found by the ABC fitting (here $N = 500$). The distribution of $T_1$ parameters are tightly distributed a few hours after D3.5 suggesting that the cells responded to the increase of signal from 0 nM to 500 nM. The distribution of $T_2$ parameters however are broadly dispersed at the end of the timeline showing that the cells did not respond to the signal change at D4.

**F**. Bump function $b_{T_1,T_2}$ as a function of time with the mean parameters $T_1, T_2$ accepted by the ABC fitting. The fitting suggests that the cells did not respond to the decrease of signal at D4.

**G**. Proportions of cells in the ACs from experimental data (left) compared to simulated proportions (right) for the SAGUP condition. The simulation used the previously fitted parameter distributions $D(0)$ and $D(500)$ as in B but the change of signal is modelled using the function in F. The proportions from D4 to D6 now align with the experimental data and the FP proportions predicted by the simulation at D6 $(4\%)$ are consistent with the SAGUP experimental proportions of FP-like cells $(7\%)$.

**H**. Same as D but using the bump function $b_{T_1,T_2}$ in F to model the change in signalling and all the fitted parameters $T_1, T_2$ in E to obtain a distribution of simulated proportions. The distributions recapitulate the tendencies observed in the experimental proportions (thick blue lines).

#### B7. Fitting the model to the scRNA-seq data

##### B7.1. Fitting scRNA-seq excluding Mesoderm

We tested the model on the scRNA-seq (SAG 500nM) dataset (Fig. B9A) using the same priors as those for the FACs at the same concentration (Table B4 right) with the exceptions of $u_5, v_5$ for which we had to extend the priors region (see Sect. B7.3 below). We compared the results to those obtained for the FACs at the same SAG concentration (Fig. B9B). The experimental protocols differ: firstly, the scRNA-seq dataset contains Mesoderm cells, as no inhibitor preventing the formation of these cells was used; secondly, it contains MNs and V3 neurons whereas these were excluded in the FACs (Sox2 cells were selected). To test the model and compare the results, we selected only the neural progenitors in the scRNA-seq, excluding the Early Mesoderm and Mesoderm ACs. We then computed the proportions of cells in each AC, at each time point. We categorised motor neurons, MN, as part of MNDiff, and V3Trans/V3 as part of p3 by summing their corresponding proportions at each time point (Fig. B9A left barplot). The timing in the FACs dataset differs slightly as we are comparing the production of proteins against the production of mRNA. However, we observe a similar pattern: the p0/p1 and p2 states are present at D4 but do not persist at D5 at 500nM SAG. Substantial proportions of pMN cells appear at D5 in both datasets, decreasing at D6 as they transition to MNDiff. Meanwhile, p3 cells emerge at D5 in both datasets. There are significantly more p3 cells in the scRNA-seq dataset because these proportions include the V3 neurons (which were excluded in the FACs) and the largest proportions of p3 cells are seen at D7.

For the simulation, we assumed that the initial cluster of points, used as initial conditions for the SDEs driving the trajectories of the cells, represents cells that are already committed to PreNeural (e.g cells previously labelled NMPTrans in Appendix A Fig. A1A). Fig. B9A (right barplot) shows the mean of the simulated proportions retained for the accepted parameters by the ABC fitting. The simulated proportions successfully replicated the trends observed in the experimental proportions at each time point as the data proportions (blue) mostly belong within the 95% confidence interval of the $N = 1500$ simulated proportions retained by the accepted parameters of the ABC (Fig. B9C).

##### B7.2. Transition from DP to p3

The simulation did not produce the large number of p3 cells simply by flipping the unstable manifold from PreNeural towards EarlyVentral. Instead the circular landscape topology was crucial with DP cells transitioning to p3 (Fig. B9D). This was not observed in the FACs dataset at D6 but is supported by a subsequent FACs experiment with a D7 time point that took advantage of a pulse of SAG exposure (Sect. B7.4). In this experiment, some DP cells at D7 downregulated Olig2 while retaining Nkx2.2 expression, suggesting these cells took on a p3 identity. Fig. B9D shows the simulated cells in phase space at the 4 retained time points (corresponding to D4-D7 in the experiment) for the $N = 1500$ parameters $\mathbf{p}$ accepted by the ABC fitting. The simulation at D4 accurately reproduced the early states with a trail of transitioning cells (PreN-Trans) from PreNeural and the emergence of p2 cells transitioning towards pMN (we labelled them p2/pMNTrans). At D5 we observe the emergence of the most ventral states, along with the DPTrans transitioning cluster, which bridges pMN and DP (these cells had significant counts of Olig2 but also Nkx2.9). At this stage, MNDiff cells (Neurog2$^+$) begin to appear, along side some p3 and FP cells that arise from the Early p3. At D6 the DP cells transition towards p3 following the unstable manifold connecting them, given by the circular topology of the landscape and the DP cluster continues to be filled by cells coming from DPTrans. This phenomenon persisted at D7.

##### B7.3. Comparison with FACs fitting results: parameter domains analysis

Fig. B9F illustrates the domains of accepted parameters $P_i(500)$ of the sub-landscapes for the two datasets, labelled respectively RNAseq (blue) and FACs (red). These are similar to the panels of Fig. B7 except that we extended the priors domain in Panel 5 for the scRNA-seq fitting to allow the parameter search closer to the DP fold bifurcation curve which facillitates the transition from DP to p3. We observe that, even given the differences in experimental protocols, most of the qualitative topological features are preserved. In Panel 1, both parameter domains are far to the left of the flip bifurcation curve, forcing the cells to adopt a neural identity rather than a mesoderm identity - since we ignored cells committed to Mesoderm in the scRNA-seq to focus only on the neural progenitors. The parameters involved in the PreNeural binary decision in Panel 2 are similar for both datasets. This supports the assumption that the high SAG concentration (500nM) influences the flip towards EarlyVentral. Panel 3 and 4 show that, in both datasets, the EarlyVentral, p0/p1 and p2 attractors are bifurcated. In Panel 5, the domain of the scRNA-seq (blue) straddles on both side of the DP bifurcation curve which means that some cells will remain in the DP attractor (i.e. those with parameters located to the right of the DP bifurcation curve) and some will transition to p3 (i.e. those with parameters located close or to the left of the DP fold curve). Panel 6 highlights a distinction between the two datasets regarding the p3 and FP proportions. The FACs domain (red) lies to the left of the flip bifurcation curve, favouring FP over p3, whereas the opposite is observed for the RNAseq domain (blue). This effect is a consequence of the increased number of p3 cells in the scRNA-seq data, as these were merged with V3Trans and V3 neurons.

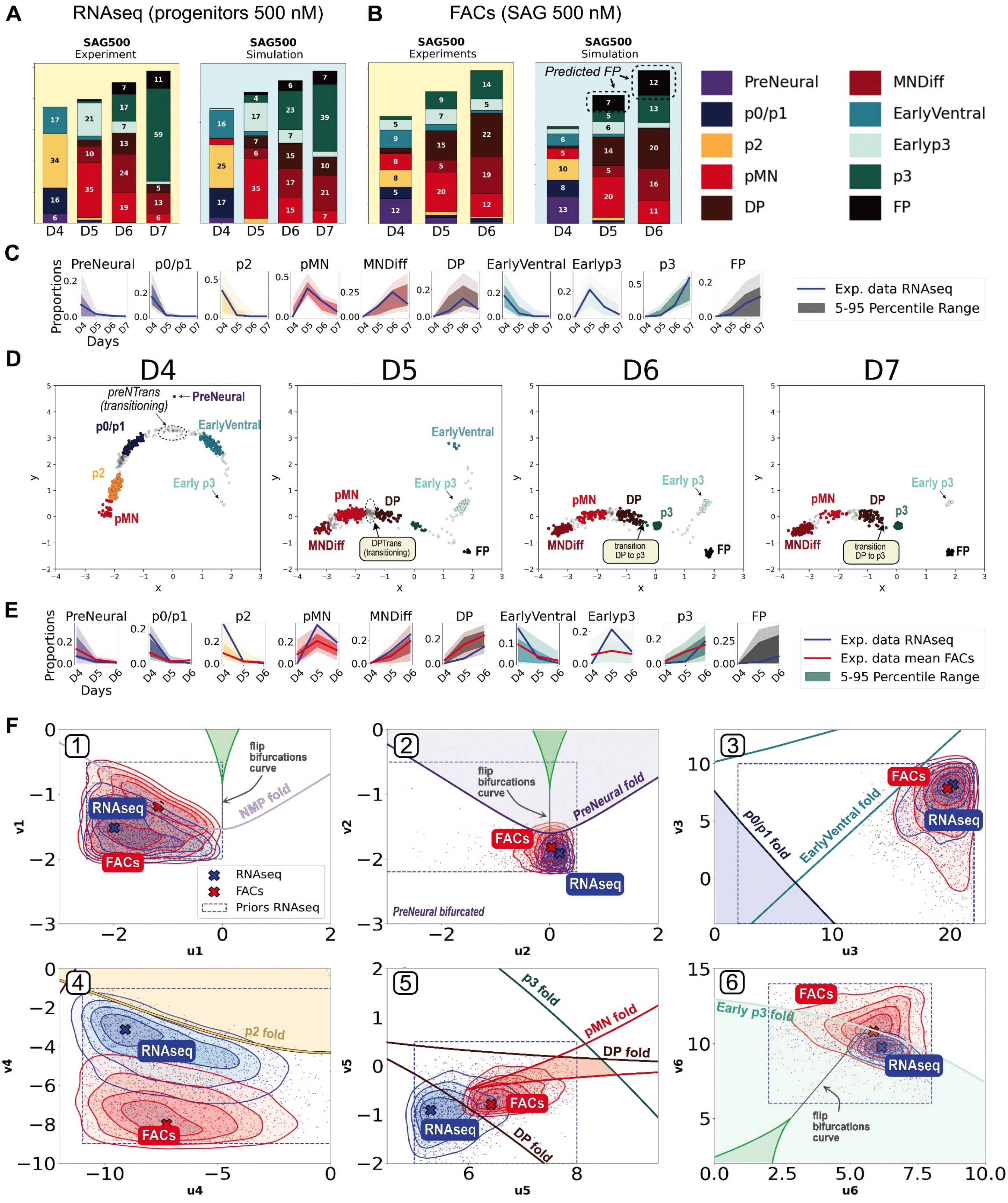

**scRNA-seq Data Fitting and Comparison with FACs at 500nM SAG**

**Figure B9:** (Caption next page)

**Figure B9:** **A**. Proportions of cells (in percent) in the indicated ACs by time point for the scRNA-seq dataset (500nM SAG) (left) compared to the means of the fitted simulated proportions for each ACs (right). **B**. Same as A for FACs (500nM). **C**. Distributions of the simulated proportions (percent values in decimal form) for each AC by time point using the accepted parameters after fitting the model to the scRNA-seq dataset (500nM SAG). The darker bands indicate the $95\%$ confidence interval. The blue line indicates the observed experimental proportions of cell types. **D**. Simulated cells by time point for each of the accepted parameters accepted by the ABC particle filter (scRNA-seq dataset). The model results in a direct transition from DP to p3. **E**. Distributions of the simulated proportions (percent values in decimal form) for each AC by time point using the accepted parameters after fitting the model to the FACs dataset (500nM). The last panel (FP) is a prediction. The scRNA-seq experimental proportions are indicated by a thick blue line. For comparison, the FACs experimental averaged proportions are represented by the red line (except in the FP panel because we did not identify FP in the FACs). **F**. Parameter domains (contour lines) within their prior regions (dashed rectangle) for parameters fit to the scRNAseq (blue) and FACS (red) projected on the bifurcation set of each sub-landscape. The points marked by a cross × (RNA-seq), × (FACs) indicate the point of highest density for each domain. Most distributions generate topologically equivalent landscape instances for each cell, as they belong to the same Morse-Smale component of parameter space, delineated by fold and flip curves. However, an exception occurs in Panel 5, where the blue distribution (RNA-seq) overlaps the DP fold curve, forcing a DP bifurcation and driving many DP cells towards a transition to p3. In contrast, the red distribution (FACs) indicates no bifurcation but DP cells can still transition to p3 due to stochastic fluctuations. This is because the FACs dataset had a lower proportion of p3 cells at D6 as V3 neurons where excluded. Panel 6 shows that the unstable manifold linked to Early p3 is primarily directed towards p3 in the RNA-seq data, whereas it also connects to FP in the FACs. This bias arises because a higher proportion p3 cells where captured in the RNA-seq at D6 as these include V3 neurons.

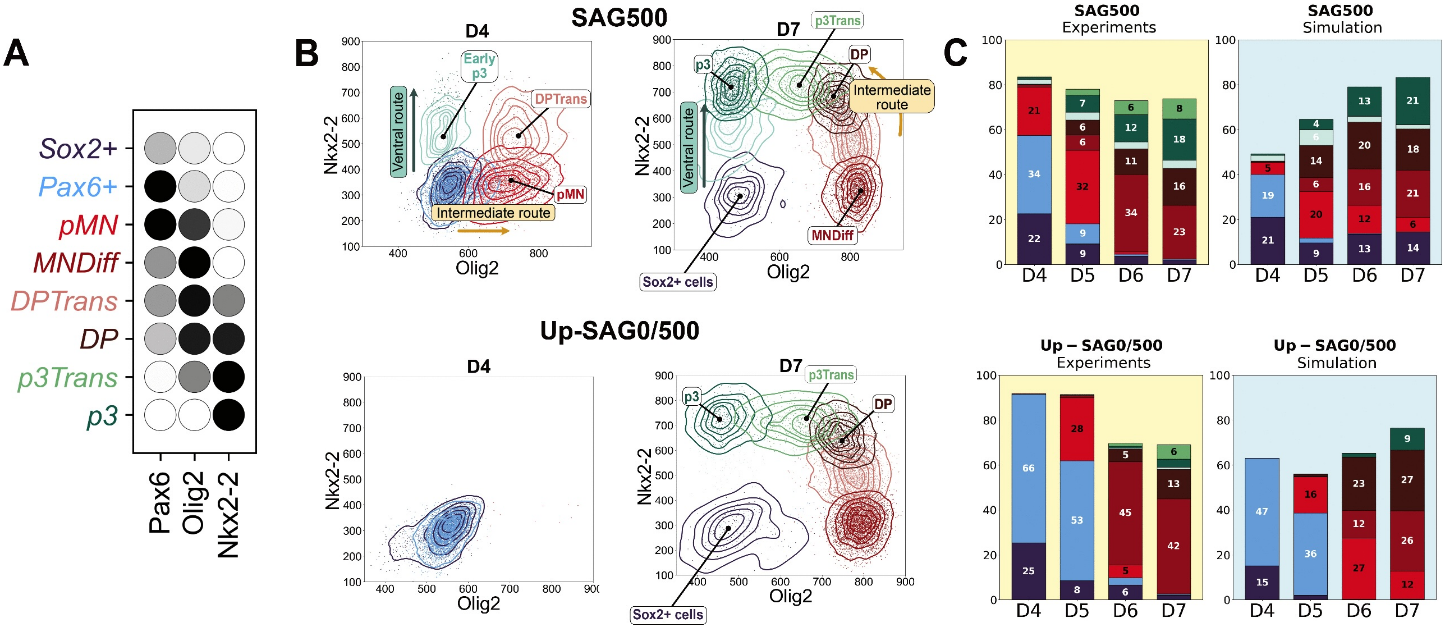

**Validation of the DP to p3 Transition**

**Figure B10:** **A.** Dot plot indicating the expression of the markers used to identify the clusters identified in the FACs dataset. Without the Nkx6.1 marker we could not distinguish p0/p1 from p2 (Pax6$^+$ cells) and PreNeural from EarlyVentral and FP (Sox2$^+$ cells). **B.** (Top) 2D plots of gene expression (Olig2 and Nkx2.2) in cell clusters at 500nM SAG distinguish the ventral route from the intermediate route. (Bottom) In the UPSAG0/500 condition, the ventral route is inaccessible at D4 and cells transition to p3 via the intermediate route from DP but this takes longer as cells transition through shallow attractors. **C.** Proportions of cells (in percent) in ACs from the experimental data (left) compared to simulated proportions (right) for both signalling conditions. The simulation used the previously fitted parameter distributions $D(0)$ and $D(500)$ as in Sect. B6. This experiment was not included in the fitting of the model to obtain the above parameter distributions.

##### B7.4. Validation of the DP to p3 Transition

The SAG 500nM scRNA-seq dataset and the fitting of the model (Sect. B7) support the idea that cells can access p3 via two distinct routes: a ventral route via EarlyVentral and Early p3, and an intermediate route via a direct transition DP-to-p3. To further test this, we extended our flow cytometry to D7, using 4 markers Sox2, Pax6, Olig2 and Nkx2.2 in two signalling conditions: a constant exposure of 500nM SAG from D3 to D7, and a delayed SAG exposure regime (24h at 0nM SAG, followed by 72h at 500nM) referred to as Up-SAG0/500. We identified the relevant cell states (Fig. B10A). Without the Nkx6.1 marker we could not distinguish p0/p1 from p2 and PreNeural from EarlyVentral and FP. We thus grouped these cells and labelled the resulting clusters Pax6$^+$ and Sox2$^+$, respectively . The temporal analysis in gene space supported a direct transition DP-to-p3 that is captured under the Up-SAG0/500 condition (Fig. B10B) as the ventral route in this case is not accessible.

To test the mathematical model with these data, we proceed as in Sect. B6 and use the parameter domains $D(0)$ and $D(500)$ obtained from previous fitting to the FACs data (Sect.

B5.1). The 5 timesteps of simulation $t_0, t_{100}, t_{200}, t_{300}$ and $t_{399}$ here correspond to the experimental time points D3, D4, D5, D6 and D7. To model the pulse, we let the individual cells parameters vary over time using sigmoids as in (10) but modified the changing time of the signal from 0nM to 500nM by setting $T_1 = 100$ (corresponding to 24h) and $T_2 = 399$ (no further signal change). At each timestep we retained the proportions of cells in each AC and combined the proportions p0/p1 and p2 ($Pax6^+$ cells) and PreNeural, EarlyVentral and FP ($Sox2^+$ cells) to compare with the experimental proportions. The result is shown in Fig. B10C. The simulation captured the larger proportions of p3 cells coming from the ventral route at 500nM SAG. In the Up-SAG0/500 simulation, the majority of p3 cells at D7 come from the intermediate route via the DP-to-p3 transition (see Main text).

#### B8. SAG pulses analysis

##### B8.1. Fitting local model results

Once the global topology of the system had been validated, we set out to test specific transitions. We focus on the transitions involving the flip between PreNeural to intermediate (p0-p2) or ventral (p3, FP) fates, and the later transition from p0/p1 towards p2 and pMN. We build a new simplified model that considers these two sub-landscapes and aggregates EarlyVentral, Early p3 and p3 into a state called p3*; and pMN, MNDiff and DP into a state called pMN*. The two pieces of the model correspond to a flip and a linear transition of states joined at the p0/p1 attractor.

The fitting of the signalling regimes 0nM, 10nM, 100nM, 500nM SAG can be achieved in a similar way to those with the complete model topology. The main goal of this simplified model is to understand the effect of different signalling pulses in the state proportions of the relevant cell types.

##### B8.2. Butterfly normal form as sub-landscape model

The butterfly normal form model discussed in [17,18] is used to produce a three attractor configuration in which the attractors and the saddles between them lie on a smooth connector consisting of the unstable manifold of the saddles (Fig. B11A). It allows a sequence of direct transitions between the states p0/p1, p2 and pMN*.

##### B8.3. Assembling the simplified model

Since this model is simpler and has only two pieces, we take a sigmoidal function to join the two decision regions. We follow an approach similar to the one for the global landscape for its construction. A normal form for the elliptic umbilic (Fig. B11B) is

$$f_{\boldsymbol{\theta}_1}(\mathbf{x}) = x^4 + y^4 + 2yx^2 - y^2 - y^3 - u_1 x - v_1 y$$

and for the butterfly, we take a two dimensional normal form inside the double cusp to ease the gluing of the two parts and we use

$$g_{\boldsymbol{\theta}_2}(\mathbf{x}) = x^4 + y^4 - 4xy^2 + x^2 + x^3 + u_2 y + v_2 x$$

with $\boldsymbol{\theta}_i = (u_i, v_i)$ for $i = 1, 2$. We translate the coordinates of the elliptic umbillic gradient form using $\tau(\mathbf{x}) = (x-2, y-2)$ and glue the two parts together with $\phi(\mathbf{x}) = \frac{1}{2}(\tanh(10(x - 1.5)) + 1)$ considering an horizontal separation between the two decision regions. The assembled model is therefore

$$\mathcal{L}(\mathbf{x}, \boldsymbol{\theta}, \mathbf{vel}) = \mathrm{vel}_1 \phi(\mathbf{x}) L_1(\tau(\mathbf{x}), \boldsymbol{\theta}_1) + \mathrm{vel}_2 (1 - \phi(\mathbf{x})) L_2(\mathbf{x}, \boldsymbol{\theta}_2)$$

where $L_1(\mathbf{x}, \boldsymbol{\theta}_1) = -\nabla f_{\boldsymbol{\theta}_1}(\mathbf{x})$ and $L_2(\mathbf{x}, \boldsymbol{\theta}_2) = -\nabla g_{\boldsymbol{\theta}_2}(\mathbf{x})$. The 6 parameters in $\boldsymbol{\theta} = (\boldsymbol{\theta}_1, \boldsymbol{\theta}_2)$ and $\mathbf{vel} = (\mathrm{vel}_1, \mathrm{vel}_2)$ depend on the SAG concentration considered.

##### B8.4. Simulations

Similar to above, the simulations are stochastic following (6). In this case we take the diffusion coefficient $\eta$ to be constant throughout the landscape and equal to the noise parameter $\sigma$. Hence, we have 6 parameters per experimental condition to fit plus one noise parameter common to all of them.

In each simulation, we simulate 500 cells. Here we assume that the landscape determining the trajectories of all cells in one simulation are the same, that is, we use the same parameters vector for all of them. Stochasticity makes them take different specific trajectories. To generate the initial condition, we simulate a first version of the landscape where the PreNeural attractor is a large deep attractor and then obtain a cloud of points in the basin of attraction of the initial state. Then, we apply the specific parameters for the condition and let the points evolve according to the SDE. We use $dt = 0.001$, with time in the interval $[0, 3 \cdot 24]$ to reproduce the real timing of the experiments. We save the positions of the points at times $t_i \in \{0, 24, 48, 72\}$ corresponding to the samples in our experimental data. Once we have simulated all conditions of interest, we aggregate all simulated points in a single dataset and apply a predefined Gaussian clustering model to these points, assuming 8 clusters corresponding to the 5 attractors and the 3 transitions between them (Fig. B11C). The proportions of points assigned to each cluster at each time point and for each experimental condition give us the simulated proportions. We aggregated the proportions for each attractor together with the cells transitioning out of it, in order to obtain 5 attractor cluster proportions that can be compared to the experimental data.

##### B8.5. Fitting using ABC approach

To fit the model to the data we used the MatLab ABC algorithm [1] based on Toni et al. [24] as presented in [2,17]. We acquired 10000 accepted particles at each iteration, and used the distance defined in (9) to compare experimental and simulated data. We reclustered the FACs data to obtain the proportions of the aggregated ACs we have defined above. We combined all the data for a specific experimental replicate together and fit a GMM with 7 components (PreNeural, p0/p1, p2, pMN, DP, p3* and transitioning).

We defined priors for the parameter regions based on the topology of the landscape and the observation of the experimental data being fitted as shown in Table B6. We kept the level of noise constant, with $\sigma = 0.02$.

##### B8.6. Constant SAG

The fitting results for constant SAG concentrations gave good agreement between experimental and simulated proportions (Fig. B11D).

We observed that for the 0nM SAG, the most important feature that defines the accepted parameters (Fig. B11E middle) is that the p0/p1 attractor is far from bifurcation, defining a stable state that captures the cells. For 10nM SAG, parameters move the system closer to the bifurcation lines of both p0/p1 and p2 attractors, in such a way that stochasticity is sufficient for the transition of a substantial number of cells towards pMN*. For 500nM SAG, the landscape parameters appear beyond the p0/p1 fold curve leading cells to rapidly transition to p2 and pMN*.

##### B8.7. Pulses analysis

We considered the effect of delayed addition of SAG. To this end we cultured cells for 24h without SAG followed by a 48h period in 500nM SAG. We termed these conditions Up-SAG0/500. In a second experiment, called Down-SAG500/0, an initial 24h period of exposure to 500nM SAG was followed by a 48h period with no SAG. The simplest hypothesis was to assume that the landscape depends only on the SAG concentration that cells are receiving, and hence that it changes at the times that the signal is added or removed. Examining the data we saw that this hypothesis is consistent with the Up-SAG0/500 experiment but is violated by the Down-SAG500/0 results. It was clear from the Down-SAG500/0 data that the p0/p1 attractor remained bifurcated when the SAG level was reduced from 500nM SAG to 0nM SAG. To further test this hypothesis, we fitted the simple model with 4 parameter regimes corresponding to the 4 concentrations (using the 4 constant SAG conditions plus the two described pulses), with the landscape being agnostic to previous treatment of the cells. The fitting was unsuccessful, with no parameter sets able to get sufficient close to all 6 experimental conditions to pass the acceptance thresholds.

We therefore hypothesised that there is an irreversible effect on cells that have seen 500nM SAG for 24 hours so that when the signal is removed, the landscape does not return to the form corresponding to 0nM SAG. We included a 5th set of parameters in the fitting corresponding to the irreversible effect of SAG and fitted the model accordingly. The result showed that the parameters accepted for post-high-SAG were very similar to those for 500nM SAG (see Fig. B11E-F), confirming this irreversible effect.

| Parameter | SAG0 | SAG10 | SAG100 | SAG500 | Up-SAG0/500 |
|---|---|---|---|---|---|
| $u_1$ | $\mathcal{N}(-0.5, 0.2)$ | $\mathcal{N}(-0.5, 0.2)$ | $\mathcal{N}(-0.2, 0.2)$ | $\mathcal{N}(-0.1, 0.2)$ | $\mathcal{N}(0, 0.2)$ |
| $v_1$ | $\mathcal{N}(-1.2, 0.2)$ | $\mathcal{N}(-1.2, 0.2)$ | $\mathcal{N}(-1.2, 0.2)$ | $\mathcal{N}(-1.2, 0.2)$ | $\mathcal{N}(-1.2, 0.2)$ |
| $u_2$ | $\mathcal{N}(0.6, 0.1)$ | $\mathcal{N}(0.75, 0.1)$ | $\mathcal{N}(1.1, 0.2)$ | $\mathcal{N}(0.9, 0.2)$ | $\mathcal{N}(0.9, 0.2)$ |
| $v_2$ | $\mathcal{N}(0.6, 0.1)$ | $\mathcal{N}(0.6, 0.2)$ | $\mathcal{N}(1.5, 0.4)$ | $\mathcal{N}(1, 0.2)$ | $\mathcal{N}(1.1, 0.2)$ |
| $\mathrm{vel}_1$ | $\mathcal{N}(0.2, 0.2)$ | $\mathcal{N}(0.4, 0.2)$ | $\mathcal{N}(0.2, 0.2)$ | $\mathcal{N}(0.2, 0.2)$ | $\mathcal{N}(0.3, 0.2)$ |
| $\mathrm{vel}_2$ | $\mathcal{N}(0.2, 0.2)$ | $\mathcal{N}(0.2, 0.2)$ | $\mathcal{N}(0.2, 0.2)$ | $\mathcal{N}(0.2, 0.2)$ | $\mathcal{N}(0.2, 0.2)$ |

**Table B6:** Priors Target Model FACs data.

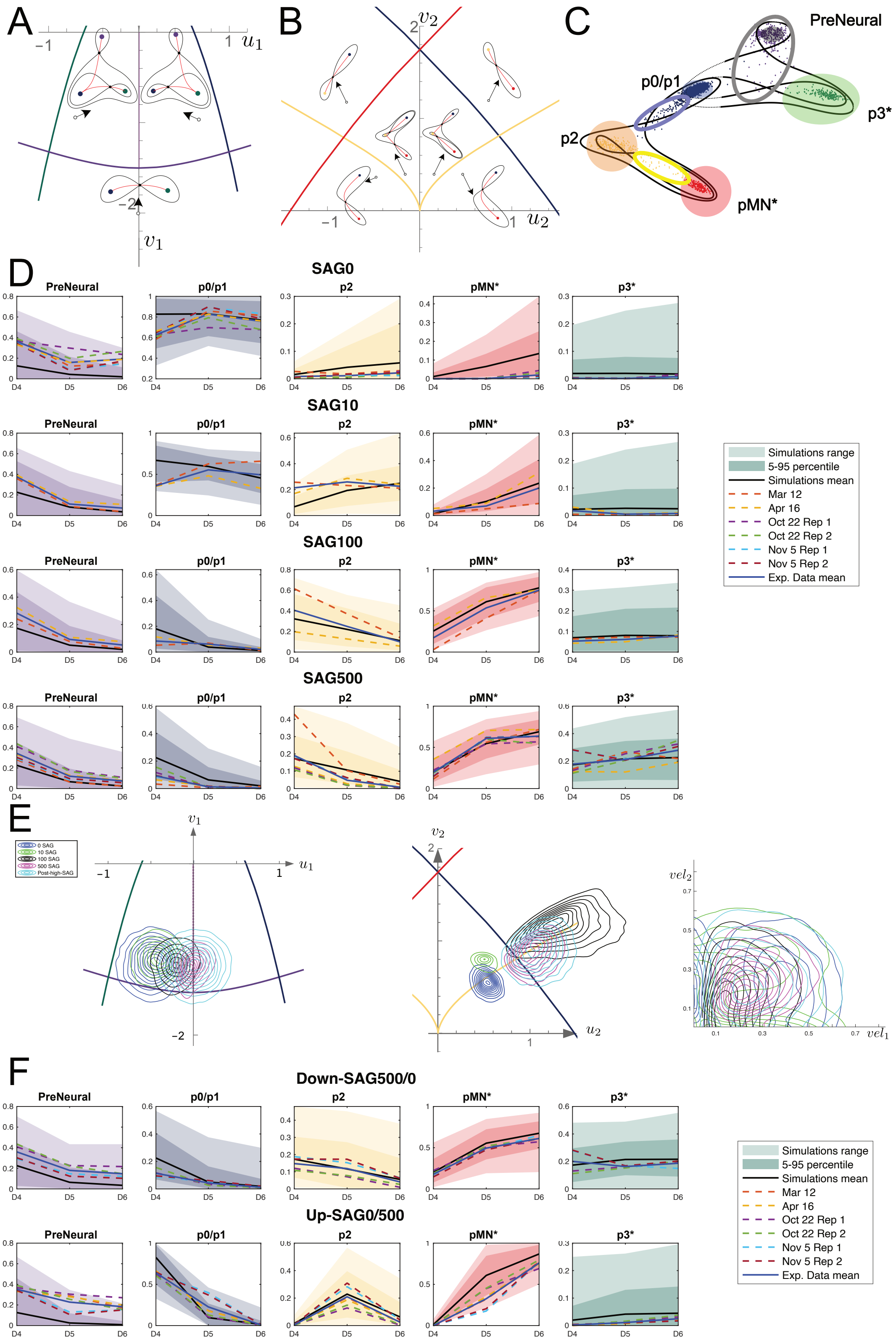

**Model Predictions through Signalling Perturbations**

**Figure B11:** **A.** Binary flip landscape. **B.** Binary choice sub-landscape with a dual cusp. **C.** Simplified model with fate regions shown. **D.** Results for constant SAG. **E.** Parameters obtained in constant SAG. **F.** Results pulses.

#### B9. Model Parameters fit to the human dataset

We fitted the same model (GL) to the human dataset [16] (Fig. B12A), following the procedure outlined in Sect. B4 with the same discretised time points and priors of Table B7. In this case, the parameter domain $D$ of the SDE (6) is not determined by the concentration of a specific morphogen but rather by whether the application of TGF$\beta$ is delayed by 18h or 24h. We focused on the neural progenitor landscape, initialising the simulations around PreNeural rather than NMP.

We retained the simulated proportions $X_{sim}$ across 5 time points, as described in Sect. B4.1, but only considered the 3 time points D3, D5, D7 for comparison with the experimental proportions $X_{data}$ by using the distance (9). The simulated cells across these 3 time points are shown in Fig. B12B next to the experimental data in 2D PCA space.

The simulated proportions $X_{sim}$, derived from the $N = 1500$ accepted parameters after 8 generations of the ABC algorithm, form tight distributions around their mean values, which closely align with the experimental proportions (Fig. B12C). As expected, we observe a higher proportion of p3 and FP at D7 with a 24h-delay compared to an 18h-delay. This quantitative analysis highlights how the presence of the notochord influences the patterning of the ventral progenitors.

##### B9.1. Bifurcation analysis

The parameter domains projected on the bifurcation sets of each sub-landscape were obtained as described in Sect. B5.1. This highlighted key insights from the two experimental conditions. The major qualitative difference between the two conditions appeared at the level of the PreNeural flip bifurcation (Fig. B12E). Specifically, the dynamics of the cells initially in the PreNeural state are predominantly governed by the model on the left (blue) for the 18h-delay condition, whereas in the 24h-delay condition model shifts to the dynamics of the model on the right (red). This interpretation is supported by the location of the parameter domains in Panel (2) of Fig. B12F: the blue distribution (18h-delay) is located on the left of the flip bifurcation curve, above the PreNeural fold curve, while the red distribution (24h-delay) lies on the right. Other bifurcations remain qualitatively unaffected as the two distributions (18h-delay and 24h-delay) are drawn in regions of parameter space with topologically equivalent sub-landscape instances (Fig. B12F Panels (3)-(6)). In Panel (3), both domains are away from the p0/p1 bifurcation curve indicating that the p0/p1 attractor is bifurcated in both cases. This is consistent with our previous analysis for high SAG concentrations (100nM and 500nM) demonstrating the destabilisation of the p0/p1 state (Sect. B5.1 (3)). The same conclusion applies for EarlyVentral in Panel (3) suggesting that the cells transition either straight to p2 or to Early p3 as indicated in Fig. B12E. In Panel (4) both domains appear near - but above - the p2 fold curve and hence this attractor is present but shallow for most cells. In Panel (5), the parameters did not converge sharply but the high density points (×-marked), and most of the parameters remain within the region of the bifurcation set in which the pMN attractor is present.

Similarly in Panel (6) the Early p3 appears close to bifurcation. The fact that both domains lie on top of the flip bifurcation curve near the fold-flip point reflects that p3 and FP cells appear synchronously in both experiments.

| Parameter | Priors | Parameter | Priors |
|---|---|---|---|
| $\mathbf{u_2}$ | $\mathcal{U}([-0.6, 0.6])$ | $\mathbf{v_2}$ | $\mathcal{U}([-1.8, -1.1])$ |
| $\mathbf{u_3}$ | $\mathcal{U}([10, 18])$ | $\mathbf{v_3}$ | $\mathcal{U}([-2, 4])$ |
| $\mathbf{u_4}$ | $\mathcal{U}([-4, 0])$ | $\mathbf{v_4}$ | $\mathcal{U}([-6, -2])$ |
| $\mathbf{u_5}$ | $\mathcal{U}([5.8, 7.8])$ | $\mathbf{v_5}$ | $\mathcal{U}([-0.8, 0.5])$ |
| $\mathbf{u_6}$ | $\mathcal{U}([4, 7])$ | $\mathbf{v_6}$ | $\mathcal{U}([8, 12.5])$ |
| $\sigma_i$ | $\mathcal{U}([0.2, 0.8])$ | $\text{vel}_i$ | $\mathcal{U}([0.2, 1])$ |

**Table B7:** Priors used for the ABC fitting of the human scRNA-seq dataset. The parameters $\mathbf{u_1}, \mathbf{v_1}$ were omitted as the NMP sub-landscape was not included. All noise and velocity parameters had same priors.

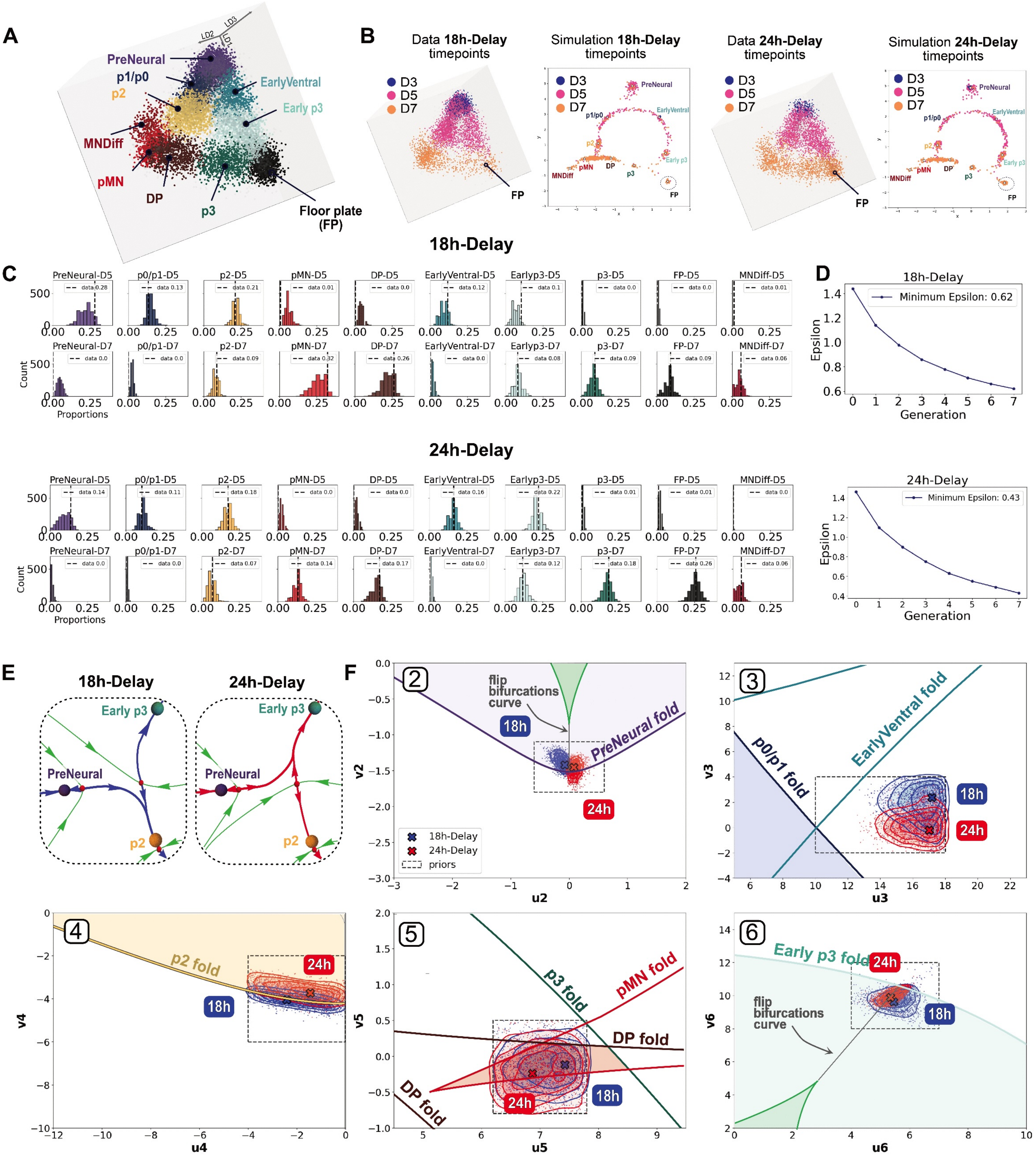

**Landscape Model for Human Organoid Datasets**

**Figure B12:** (Caption next page)

**Figure B12:** **A**. 3D LDA representation of the ACs corresponding to the neural progenitors in the dataset for all time points and both experimental conditions. **B**. 3D LDA representations of the data for each experimental condition, coloured by time points (D3,D5,D7) next to the simulated synthetic data coloured by the corresponding time points. More cells transition along the ventral route in the 24h-Delay condition, generating more FP cells. **C**. Histograms showing the distribution of the 1500 simulated proportions, corresponding to the accepted parameters of the ABC particle filter, for the time points D5 and D7 and each experiment. D3 is not shown as there are primarily PreNeural cells. The dashed black vertical lines correspond to the experimental proportions. **D**. Decrease in the $\varepsilon$ acceptance threshold for the 8 generations of the ABC algorithm (Sect. B4.3). **E**. Cells in the PreNeural attractor are most likely to follow the intermediate route towards p2 in the 18h-delay condition, with their trajectories governed by dynamical systems topologically equivalent to the one illustrated in blue on the left. By contrast, cells follow the ventral route in the 24h-delay condition, resulting in dynamical systems equivalent to the one shown in red on the right. **F.** Sub-landscape bifurcation sets. Each panel displays the parameter domains corresponding to an 18h-delay (blue) and a 24h-delay (red), drawn on top of the bifurcation sets of each sub-landscape (indexed as those in Fig. B7A). The points marked by a cross $\times$ (18h) or $\times$ (24h) indicate the highest density point for each domain. We removed the NMP sub-landscape as we were interested in the neural progenitors. The only qualitative differences appear at the level of the PreNeural flip bifurcation in Panel 2. The other parameter distributions produce equivalent cells landscapes as they cover the same regions in parameter space, delimited by the fold and flip curves.

#### References

[1] E. Camacho-Aguilar. *New mathematical methods for the study of stem cell differentiation.* PhD thesis, University of Warwick, Warwick, UK, 2018. URL `https://wrap.warwick.ac.uk/id/eprint/108668/`.

[2] E. Camacho-Aguilar, A. Warmflash, and D. A. Rand. Quantifying cell transitions in c. elegans with data-fitted landscape models. *PLoS Computational Biology*, 17(6): 1–28, 2021. doi: 10.1371/journal.pcbi.1009034.

[3] M. A. Coomer, L. Ham, and M. P. H. Stumpf. Noise distorts the epigenetic landscape and shapes cell-fate decisions. *Cell Systems*, 13(1):83–102.e6, 2022. doi: 10.1016/j.cels.2021.09.002.

[4] F. Corson and E. D. Siggia. Gene-free methodology for cell fate dynamics during development. *eLife*, 6:1–25, 2017. doi: 10.7554/eLife.30743.

[5] S. Farjami, K. C. Sosa, J. H. Dawes, R. N. Kelsh, and A. Rocco. Novel generic models for differentiating stem cells reveal oscillatory mechanisms. *Journal of the Royal Society Interface*, 18(182):20210442, 2021. doi: 10.1098/rsif.2021.0442.

[6] S. Filippi, C. P. Barnes, J. Cornebise, and M. P. H. Stumpf. On optimality of kernels for approximate bayesian computation using sequential monte carlo. *Statistical Applications in Genetics and Molecular Biology*, 12(1):87–107, 2013. doi: 10.1515/sagmb-2012-0069.

[7] D. J. Higham. An algorithmic introduction to numerical simulation of stochastic differential equations. *SIAM Review*, 43(3):525–546, 2001. doi: 10.1137/S0036144500378302.

[8] S. Huang, G. Eichler, Y. Bar-Yam, and D. E. Ingber. Cell fates as high-dimensional attractor states of a complex gene regulatory network. *Physical Review Letters*, 94(12):128701, 2005.

[9] S. A. Kauffman. Metabolic stability and epigenesis in randomly constructed genetic nets. *Journal of Theoretical Biology*, 22(3):437–467, 1969. doi: 10.1016/0022-5193(69)90015-0.

[10] S. A. Kauffman. *The Origins of Order: Self-Organization and Selection in Evolution.* Oxford University Press, New York, 1993. ISBN 9780195079517.

[11] E. Klinger, D. Rickert, and J. Hasenauer. pyabc: distributed, likelihood-free inference. *Bioinformatics*, 34 (20):3591–3593, 2018.

[12] P. E. Kloeden and E. Platen. *Numerical Solution of Stochastic Differential Equations.* Springer-Verlag, Berlin, 1992. doi: 10.1007/978-3-662-12616-5.

[13] S. Newhouse and M. M. Peixoto. There is a simple arc joining any two morse-smale flows. *Astérisque*, 31:15–41, 1976.

[14] A. Raju and E. D. Siggia. A geometrical model of cell fate specification in the mouse blastocyst. *Development*, 151(8):dev202467, 2024. doi: 10.1242/dev.202467.

[15] D. A. Rand, A. Raju, M. Sáez, F. Corson, and E. D. Siggia. Geometry of gene regulatory dynamics. *PNAS*, 118(38), 2021.

[16] T. Rito, A. R. G. Libby, M. Demuth, M.-C. Domart, J. Cornwall-Scoones, and J. Briscoe. Timely tgf$\beta$ signalling inhibition induces notochord formation from human pluripotent stem cells. *Nature*, 637, 2025.

[17] M. Sáez, R. Blassberg, E. Camacho-Aguilar, E. D. Siggia, D. A. Rand, and J. Briscoe. Statistically derived geometrical landscapes capture principles of decision-making dynamics during cell fate transitions. *Cell Systems*, 2022.

[18] M. Sáez, J. Briscoe, and D. A. Rand. Dynamical landscapes of cell fate decisions. *Interface Focus*, 12(4): 20220002, 2022.

[19] Y. Schälte, E. Klinger, E. Alamoudi, and J. Hasenauer. pyabc: Efficient and robust easy-to-use approximate bayesian computation. *Journal of Open Source Software*, 7(74):4304, 2022.

[20] S. Smale. On gradient dynamical systems. *Annals of Mathematics*, 74(1):199–206, 1961. doi: 10.2307/1970311. URL `https://www.jstor.org/stable/1970311`.

[21] M. P. Stumpf. Inferring better gene regulation networks from single-cell data. *Current Opinion in Systems Biology*, 27:100342, 2021. ISSN 2452-3100. doi: 10.1016/j.coisb.2021.05.003.

[22] R. Thom. Topological models in biology. *Topology*, 8(3): 313–335, 1969. doi: 10.1016/0040-9383(69)90018-4.

[23] R. Thom. *Structural Stability and Morphogenesis: An Outline of a General Theory of Models.* CRC Press, 1989.

[24] T. Toni, D. Welch, N. Strelkowa, A. Ipsen, and M. Stumpf. Approximate bayesian computation scheme for parameter inference and model selection in dynamical systems. *Journal of The Royal Society Interface*, 6(31): 187–202, 2009.

[25] E. C. Zeeman. The umbilic bracelet and the double-cusp catastrophe. In P. Hilton, editor, *Structural Stability, the Theory of Catastrophes, and Applications in the Sciences*, volume 525 of *Lecture Notes in Mathematics*, pages 328–366. Springer, Berlin, Heidelberg, 1976. ISBN 978-3-540-07791-6. doi: 10.1007/BFb0077854.

[26] S. Zhang and M. P. H. Stumpf. Learning cell-specific networks from dynamical single cell data. *bioRxiv*, 2023. doi: 10.1101/2023.01.08.523176. URL `https://www.biorxiv.org/content/10.1101/2023.01.08.523176v2`. Preprint.
